# Mapping the mammalian dark metabolome by *in vivo* isotope tracing

**DOI:** 10.64898/2026.03.31.713900

**Authors:** Michael R. MacArthur, Justine Raeber, Wenyun Lu, Hantao Qiang, Amelia V. Schueppert, Lucas B. Ayres, Ricardo A. Cordova, Michael D. Neinast, Edmundo Leiva, Vanha N. Pham, Jenna E. AbuSalim, Connor S.R. Jankowski, Laith Z. Samarah, Asael Roichman, Christian G. Peace, Daniil G. Ivanov, Gianna L. Renzo, Anna M. Oschmann, Julien F. Ayroles, Sarah J. Mitchell, Xi Xing, Kellen Olszewski, Hahn Kim, Joshua D. Rabinowitz, Michael A. Skinnider

## Abstract

Despite decades of biochemical study, a comprehensive map of the mammalian metabolome remains elusive. Mass spectrometry-based metabolomics detects thousands of small molecule-associated signals in mammalian tissues, but it is currently unclear how many of these reflect products of endogenous metabolism. Here, we leverage systematic *in vivo* isotope tracing to infer the biosynthetic origins of unidentified metabolites. We administered 26 different isotopically labelled nutrients to mice, measured circulating and tissue metabolite labelling by mass spectrometry, and developed a statistical framework to infer the number of carbon atoms incorporated from each of these precursors into more than 4,000 putative metabolites. We show this information can be harnessed for biosynthesis-aware structure elucidation using a multimodal AI model that co-embeds isotopic labelling patterns with chemical structures. This approach revealed several previously unrecognized families of mammalian metabolites, including cysteine-derived alkylthiazolidines, dithioacetal mercapturic acid derivatives, short-chain N-acyltaurines, acylglycyltaurines, and N-oxidized taurines. It further uncovered a family of mevalonate-derived isoprenoid metabolites that includes 2,3-dihydrofarnesoic acid, which is markedly depleted in both mouse and human aging. Age-related depletion of these isoprenoids is driven by impaired coenzyme A synthesis. Our work establishes the biosynthetic precursors for thousands of unidentified metabolites and reveals multiple previously unrecognized branches of mammalian metabolism.

---

Two decades after the initial sequencing of the human and mouse genomes, a reference map of the mammalian metabolome remains elusive. More than a century of biochemical study has produced detailed maps of mammalian metabolic pathways, but mass spectrometry-based metabolomics routinely detects thousands of peaks that cannot readily be superimposed upon these maps^1,2^. Mounting interest in elucidating this biochemical “dark matter” has motivated the development of artificial intelligence (AI) models to facilitate the discovery of previously unrecognized metabolites^3^. Current approaches aim to infer the structures of unknown metabolites from their tandem mass spectra (MS/MS)^4–11^. However, despite the proliferation of these methods, reports of their successful application to discover previously unrecognized metabolites have been limited. This disconnect suggests that MS/MS data alone may be insufficient for *de novo* structure elucidation at the scale and pace required to complete a draft map of the mammalian metabolome.

Isotope tracing is a technique in which cells or mice are treated with heavy labelled precursors (for instance, amino acids, sugars, or fatty acids containing carbon-13). Incorporation of those precursors into downstream metabolites is then quantified by mass spectrometry^12^. Isotope tracing is widely used to monitor flux through well-studied metabolic pathways, but can also be leveraged to establish the biosynthetic precursors of unknown metabolites, and thereby provide an orthogonal source of information for structure elucidation^13,14^. Here, we perform systematic *in vivo* experiments to trace the fates of 26 isotopically labelled nutrients across 18 mouse tissues (**Fig. 1a**). We show that this data enables identification of the biosynthetic precursors for thousands of unidentified metabolites, and integrate this information into an AI model for biosynthesis-aware structure elucidation. We leverage these approaches to uncover several previously unrecognized families of mammalian metabolites, including a series of isoprenoids that are markedly depleted in aging.

**Fig. 1.**
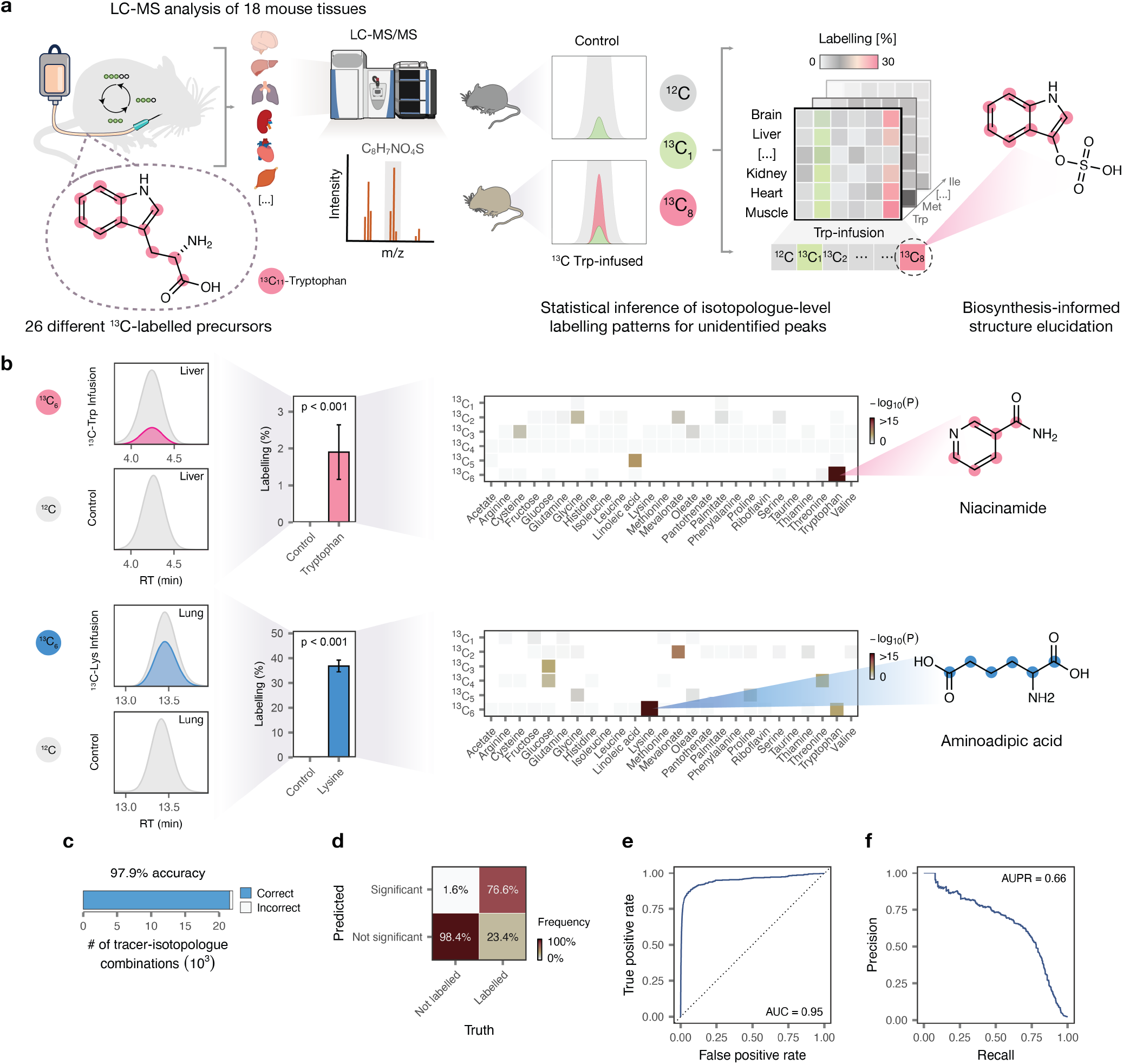
Metabolome-wide determination of the biosynthetic precursors of unidentified metabolites. **a**, Overview of the study design. The fates of 26 different ^13^C-labelled precursors were traced across 18 mouse tissues by LC–MS. The number of carbon atoms incorporated from each precursor into each detected peak was inferred from the resulting isotopologue distributions, which in turn facilitated structure elucidation of previously unrecognized metabolites. **b**, Example labelling patterns for peaks corresponding to known metabolites. The workflow converts raw extracted ion chromatograms, left, into isotopologue-level quantifications (fractional labelling after natural isotope abundance correction), center, which are used to generate a matrix of p-values that quantify the evidence for labelling with a given number of carbon atoms from each tracer, right. Cells with dark borders show significant labelling events after Bonferroni correction. **c**, Overall accuracy of the statistical model for inferring isotopologue-level labelling patterns, evaluated against manually curated ground-truth annotations for 123 known metabolites. **d**, Confusion matrix showing the concordance between isotopologues significant at a Bonferroni-corrected p-value of 0.01 versus ground-truth labelling patterns. **e**, Receiver operating characteristic (ROC) curve for the statistical model. Inset text shows the area under the curve (AUC). **f**, Associated precision-recall curve and area under the precision-recall curve (AUPR).

## Results

### Biosynthetic precursors of unidentified metabolites

We first leveraged liquid chromatography-negative mode electrospray ionization-mass spectrometry (LC–MS) to profile the metabolomes of 25 mouse tissues and biofluids. Mass spectrometric analysis of pooled extracts from these tissues detected 8,277 peaks, of which 3,771 could be rationalized as a mass spectrometry artifact (for instance, an adduct, dimer, isotopologue, or in-source fragment)^15^, whereas the remaining 4,506 represented putative metabolites (**Supplementary Table 1**). Of these, only 231 (5.1%) could be identified via comparison to a library of metabolite standards, while the remainder were unidentified.

To establish the biosynthetic precursors of these unidentified metabolites, we performed a series of *in vivo* isotope tracing experiments. A total of 26 isotopically labelled nutrients (^13^C-tracers), spanning a range of biochemically important precursors, were administered to mice. These included amino acids, sugars, fatty acids, vitamins, and other metabolic building blocks (**Supplementary Fig. 1a** and **Supplementary Table 2a**). We profiled a median of 17 tissues from each tracer condition (range, 13-18; *n* = 2-3 mice per tracer and 6 control mice) by LC–MS, for a total of 1,024 samples (**Supplementary Fig. 1b-c**).

We then sought to infer from this data the repertoire of precursors incorporated into each unidentified metabolite, and the number of carbon atoms incorporated from each of these precursors. Even with chromatographic separation and high-resolution mass spectrometry, accurate determination of isotopologue-level labelling patterns at the metabolome scale is challenging due to the number of peaks being monitored, dilution of signal across isotopologues, and the potential for interferences from other labeled or unlabeled metabolites. To achieve sufficient statistical power for metabolome-wide inference of labelling patterns, we developed a framework to jointly model the isotopologue-level data from all tissues and tracers simultaneously. This framework yields, for each metabolite, a matrix of p-values that quantify the evidence for labelling with a given number of carbon atoms from each tracer, which is reduced to a set of statistically significant labelling events following correction for multiple hypothesis testing (**Fig. 1b**).

To evaluate the performance of our model, we examined its predictions for a subset of peaks corresponding to known metabolites. Ground-truth labelling patterns were defined for each of these peaks through manual inspection of their extracted ion chromatograms (EICs), prior to the development of the statistical model itself (**Supplementary Fig. 2a-b**). The model accurately recapitulated these annotations, correctly classifying 97.9% of labelling events and achieving an area under the receiver operating characteristic curve (AUROC) of 0.95 (**Fig. 1c-e** and **Supplementary Fig. 2c-h**).

Examination of the most statistically significant labelling events revealed close agreement with canonical mammalian metabolic pathways (**Supplementary Fig. 3a**). For instance, among the most significant labelling events was three-carbon labeling of lactate from glucose, reflecting glycolytic activity. Other top-ranked labelling events likewise recapitulated well-characterized biosynthetic routes, such as the synthesis of niacinamide from tryptophan, 2-aminoadipate from lysine, and hypotaurine from cysteine (**Fig. 1b** and **Supplementary Fig. 3b**). In total, at least one significant labelling event was detected for 3,476 of the 4,506 putative metabolites (77.9%; **Supplementary Fig. 3c**).

Together, these results establish an experimental and computational framework to systematically determine the biosynthetic precursors of thousands of unidentified mammalian metabolites.

### Biosynthesis-aware structure elucidation

The incorporation of labelled carbon atoms from defined precursors places informative constraints on the potential structure of an unidentified metabolite. For instance, incorporation of seven carbons from tryptophan suggests the presence of an anthranilic or quinolinic acid moiety, whereas incorporation of six labelled carbons might instead imply a picolinic acid substructure. We hypothesized that a biochemical AI model could learn the patterns of precursor incorporation into known metabolites and leverage this understanding to guide the identification of previously uncharacterized metabolites.

Linking isotopic labelling patterns with chemical structures required a machine-learning framework capable of learning correspondences between these two different data modalities. Multimodal contrastive learning can represent different input data types in a shared low-dimensional embedding space (**Fig. 2a**)^16^. In this paradigm, a neural network is trained to map representations of the same underlying entity to nearby points within the embedding space, while pushing representations of different entities apart. A canonical example involves images and their captions, where an image is embedded close to its corresponding text description, but far from unrelated captions. Analogously, we sought to co-embed labelling patterns and chemical structures.

**Fig. 2.**
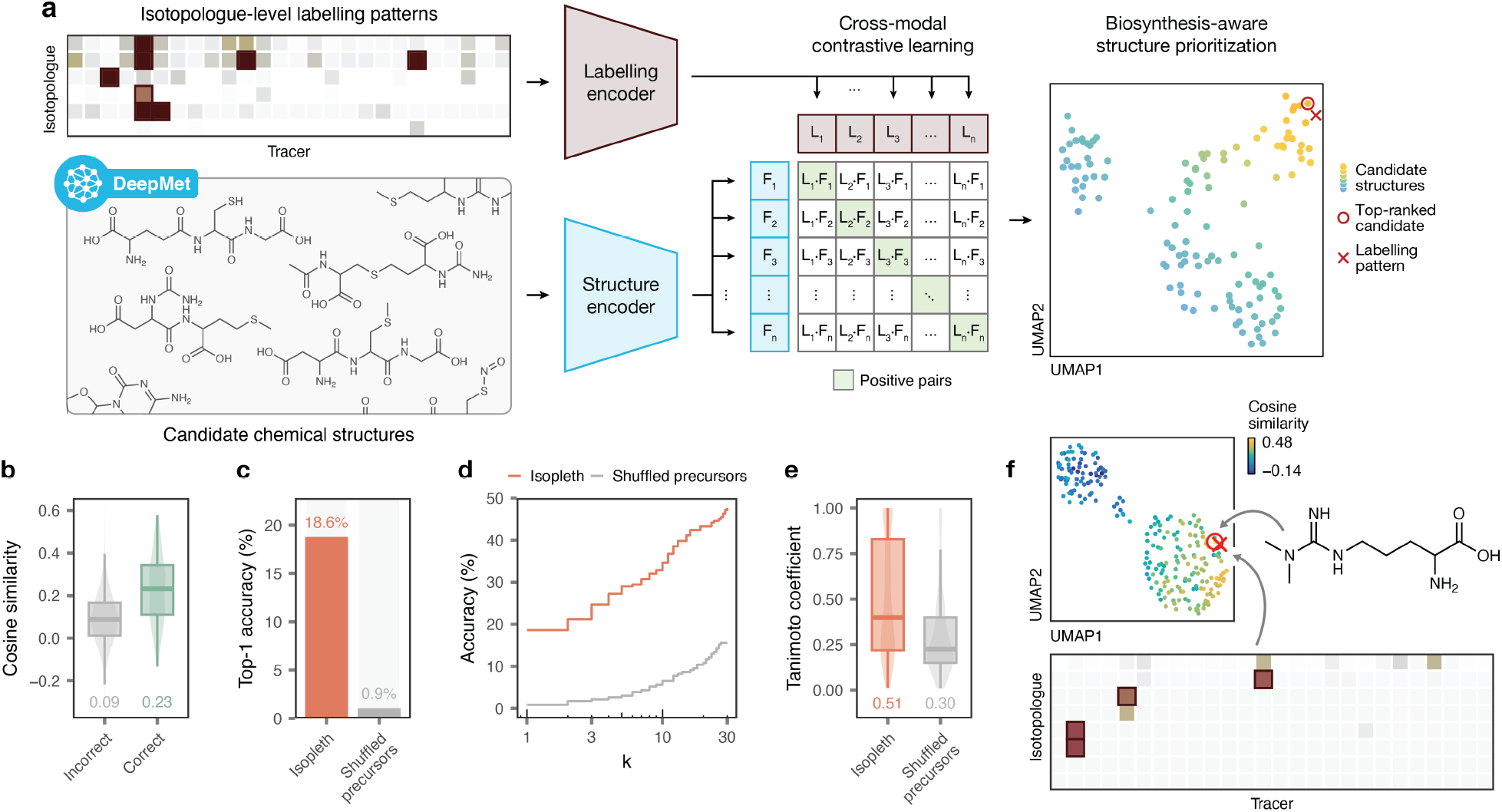
Biosynthesis-aware structure prioritization via multimodal contrastive learning. **a**, Schematic overview of Isopleth. Isotopologue-level labelling patterns and candidate chemical structures generated by DeepMet are mapped into a shared embedding space, placing labelling patterns for each metabolite near their corresponding structures while pushing unpaired patterns and structures apart using cross-modal contrastive learning. At inference time, candidate structures for an unidentified peak are ranked by their cosine similarity to the observed labelling pattern within this learned embedding space, allowing for structures to be scored based on their compatibility with labelling patterns revealed by systematic isotope tracing. **b**, Cosine similarities between labelling pattern and candidate structure embeddings for held-out metabolites, shown separately for correct versus incorrect candidates. **c**, Top-1 accuracy of biosynthesis-guided structure prioritization using Isopleth, compared to a control model trained on shuffled precursor labels. **d**, As in **c**, but showing the top-*k* accuracy (for *k* ≤ 30). **e**, Tanimoto coefficients between the structures of held-out metabolites and the top-ranked structures prioritized by Isopleth or a control model trained on shuffled precursor labels. **f**, Example of a held-out metabolite correctly prioritized by Isopleth. Top left, UMAP visualization of Isopleth embeddings for candidate structures, colored by their cosine similarity to the observed labelling pattern for this peak, shown at the bottom. The top-ranked candidate (red circle) corresponds to the correct structure, asymmetric dimethylarginine, shown at the right.

We trained a multimodal contrastive learning model on the structures and labelling patterns of peaks with known chemical identities. We named the resulting model Isopleth, after the cartographical term for a curve connecting points of equal value, to reflect our goal of aligning chemical structures with their biosynthetic precursors in a shared coordinate system. We envisioned that this model would allow for candidate structures to be ranked according to their compatibility with the labelling pattern of a given peak. To populate the search space of candidate structures for unidentified peaks, we leveraged DeepMet, a chemical language model that learns from the structures of known metabolites to propose metabolite-like structures that are absent from existing databases^17^.

To evaluate the performance of our approach, we simulated metabolite discovery by withholding known metabolites from the training sets of both DeepMet and Isopleth. Given no information other than an exact mass and labelling pattern, Isopleth correctly retrieved the exact chemical structure of held-out metabolites in 18.6% of cases (**Fig. 2b-c,f** and **Supplementary Fig. 4a-b**). In contrast, a model trained on randomized isotope labelling patterns achieved an accuracy of just 0.9%. Moreover, in cases where the top-ranked structure was not that of the held-out metabolite, the correct structure was often found within a short list of high-scoring candidates (top-3 accuracy, 24.7%; top-10, 34.6%; **Fig. 2d**). Additionally, the top-ranked structures, even when incorrect, were often structurally similar to the true metabolite, demonstrating an average Tanimoto coefficient of 0.51 to the true structure (compared to 0.30 for a model trained on randomized labelling patterns; **Fig. 2e**).

Finally, we asked whether Isopleth would complement existing approaches to metabolite annotation based on MS/MS spectrum and retention time prediction. To address this question, we trained classifiers to distinguish correct from incorrect annotations using predicted MS/MS spectra and retention times^17^, with or without including the cosine similarities emitted by Isopleth as an additional feature. Inclusion of Isopleth cosine similarities significantly increased the accuracy of metabolite annotation (**Supplementary Fig. 4c-e**).

Together, these results introduce a biochemical AI framework that harnesses the biosynthetic information embedded in isotope tracing experiments for metabolite structure elucidation.

### Biosynthesis-informed metabolite discovery

Having established the computational frameworks to infer the biosynthetic precursors of unidentified metabolites and to leverage this information for structure elucidation, we next sought to apply these approaches to elucidate the structures of unidentified peaks in our mouse tissue dataset. We studied a series of peaks that incorporated at least one of the precursors administered to mice, but could not readily be identified as a known metabolite.

We began by examining an unidentified peak that was inferred to incorporate five carbon atoms from arginine. Integration of Isopleth with MS/MS and retention time predictions placed an N,N-dimethylated derivative of the uremic toxin argininic acid^18^ among the top-ranked candidate structures, consistent with the incorporation of five arginine-derived carbons. To corroborate this assignment, we synthesized the proposed compound, which matched the peak in the mouse urine by both retention time and MS/MS (**Fig. 3a**).

**Fig. 3.**
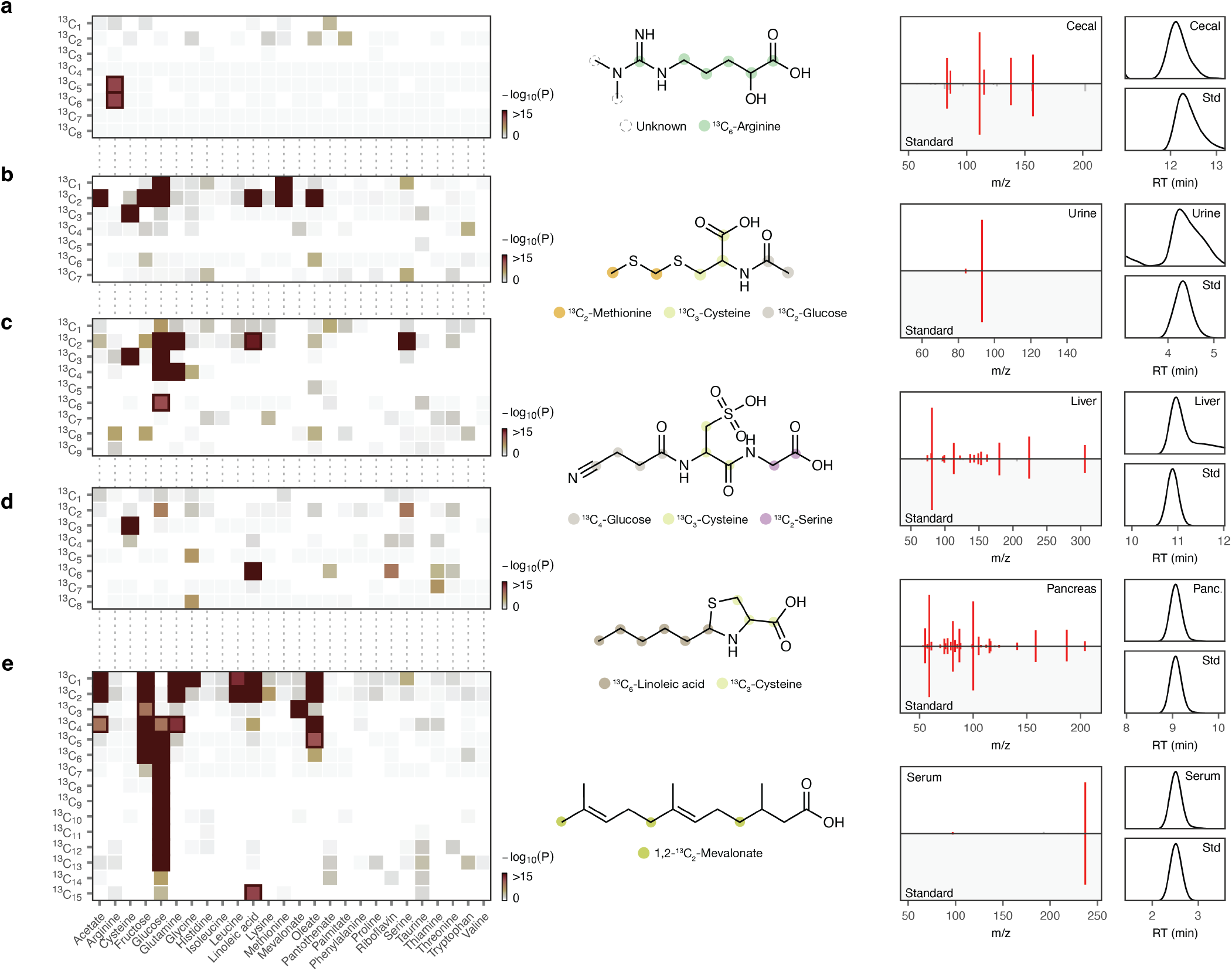
Metabolite discovery driven by isotope tracing. **a-e**, Examples of chemically diverse metabolites whose discovery was enabled by isotope tracing. Far left, heatmaps showing matrices of p-values that quantify the evidence for labelling with a given number of carbon atoms from each tracer; cells with dark borders show labelling events significant following Bonferroni correction. Center left, structures of previously unrecognized metabolites and positional labelling from ^13^C-labelled precursors (manually inferred from the structure and experimental isotope labelling patterns). Center right, mirror plots showing the similarity between MS/MS spectra from synthetic reference standards versus experimental peaks in mouse tissues. Far right, EICs for synthetic reference standards versus experimental peaks in mouse tissues. **a**, N,N-dimethylargininic acid; **b**, S-methylthiomethyl mercapturic acid; **c**, (3-cyanopropanoyl)(sulfo)alanylglycine; **d**, 2-pentylthiazolidine-4-carboxylic acid; **e**, 2,3-dihydrofarnesoic acid.

We next focused on an unidentified peak incorporating three carbons from cysteine, two from methionine, and two from various precursors to acetyl-CoA (glucose, acetate, linoleic acid, or oleic acid). Among the top-ranked candidate structures was an unusual mercapturic acid derivative bearing a dithioacetal moiety, which was prioritized in part based on the presence of a C_2_H_5_S_2_ fragment ion in the MS/MS spectrum. Chemical synthesis confirmed this peak to be S-methylthiomethyl mercapturic acid (**Fig. 3b**). To our knowledge, dithioacetal groups have not previously been described in the context of mammalian metabolism, although they occur in certain microbial natural products, such as the quinoxaline antibiotic echinomycin^19^. The identification of this metabolite highlights the potential of our approach to reveal unexpected metabolite structures.

Another unidentified peak incorporated three carbons from cysteine, two from serine, and up to four from glucose. This pattern of precursor incorporation was reminiscent of that observed for reduced glutathione, and indeed, the degree of labelling from each of these precursors correlated quantitatively with that observed for glutathione itself (**Supplementary Fig. 5a**). The tissue specificity of this unidentified peak also mirrored that of glutathione. These biosynthetic constraints, in combination with the inferred molecular formula and the SO_3_H fragment ion observed in the MS/MS spectrum, led us to hypothesize that this peak represented a nitrile-containing glutathione derivative comprising cysteinylglycine sulfonic acid conjugated to 3-cyanopropanoic acid. We synthesized this nitrile-containing structure, which matched the tissue peak by both retention time and MS/MS (**Fig. 3c**). Whereas nitrile groups are present in numerous plant secondary metabolites and microbial natural products, to our knowledge they have not been previously described in the context of endogenous mammalian metabolism^20^.

The tracing data also revealed other uncharacterized products of cysteine metabolism. One such peak incorporated three carbons from cysteine and six from linoleic acid. We tentatively annotated this peak as a thiazolidine derivative— a saturated five-membered ring containing one nitrogen and one sulfur atom—putatively arising from the condensation of cysteine with hexanal, a product of linoleic acid oxidation, in keeping with the established formation of thioproline (thiazolidine-4-carboxylic acid) from cysteine and formaldehyde^21^. Chemical synthesis confirmed this assignment (**Fig. 3d**). We reasoned that cysteine might react similarly with other aldehydes, and therefore queried our data for related species derived from aldehydes with varying chain lengths. This search yielded an additional six cysteine-derived alkylthiazolidines putatively derived from C4 to C10 aldehydes, each of which was validated by chemical synthesis (**Supplementary Fig. 5b-g**). This search also revealed a series of cysteine-derived peaks whose molecular formulas suggested one additional degree of unsaturation, which might represent thiazolines formed by condensation of cysteine with carboxylic acids rather than aldehydes. We confirmed the structure of one such thiazoline, formed by condensation of cysteine with isoleucine-derived 2-methylbutyrate (**Supplementary Fig. 5h**). Moreover, we confirmed the structures of three metabolites putatively formed by condensation of C6, C8, and C9 aldehydes with cysteinylglycine rather than cysteine (**Supplementary Fig. 5i-k**). The structures confirmed here likely represent a subset of a larger family, since an additional 19 peaks demonstrated labelling patterns and putative formulas consistent with cysteine-or cysteinylglycine-derived thiazolines or thiazolidines (**Supplementary Table 3a**).

Isotope tracing revealed incorporation of mevalonate-derived carbons into hundreds of unidentified peaks. One of the most significantly labelled species had the putative molecular formula C_15_H_26_O_2_. This formula was consistent with a C15:2 fatty acid, but typical linear fatty acids like palmitate do not label from mevalonate. Intriguingly, the peak labeled from glucose and acetate far more extensively than even the most rapidly synthesized long-chain fatty acid, palmitate, indicating it represented a high-flux metabolite from a different pathway (**Supplementary Fig. 5m**). The putative formula and mevalonate labeling suggested that this peak might represent a derivative of farnesoic acid, which is formed by oxidation of farnesyl pyrophosphate, an isoprenoid intermediate of the cholesterol biosynthetic pathway made by condensation of three mevalonate-derived isoprenyl units^22^. The formula further indicated that one of the three double bonds present in farnesoic acid was reduced. Chemical synthesis of three positional isomers confirmed the structure to be 2,3-dihydrofarnesoic acid (**Fig. 3e** and **Supplementary Fig. 5l**).

We considered the possibility that some of these metabolites might arise from microbial rather than host metabolism. To address this possibility, we reanalyzed metabolomics data from the serum and gastrointestinal tracts of mice treated with broad-spectrum antibiotics and untreated controls^23^. Of the metabolites described above, only S-methylthiomethyl mercapturic acid was reduced by more than two-fold after antibiotic treatment (**Supplementary Fig. 6a-b**). Thus, the microbiome is unlikely to be the sole source of these metabolites.

Together, these results illustrate the potential for systematic isotope tracing to guide the discovery of previously uncharacterized metabolites, including several bearing functional groups that have not previously been described in the context of mammalian metabolism.

### An expanded landscape of taurine metabolism

Taurine is among the most abundant metabolites in mammals. It is a precursor for bile acid conjugates, long-chain N-acyltaurines, and a handful of biotransformation products, such as N-acetyltaurine, taurocyamine, and isethionic acid^24–28^. Nevertheless, it is often viewed as an end product of sulfur amino acid metabolism that is cleared by renal secretion. The observation that over 300 peaks incorporated labelled taurine therefore came as a surprise and suggested that taurine metabolism might be considerably more extensive than previously appreciated. Detailed examination of these peaks led to the identification of 39 previously unrecognized taurine-derived metabolites that were validated by chemical synthesis (**Fig. 4** and **Supplementary Figs. 7-8**).

**Fig. 4.**
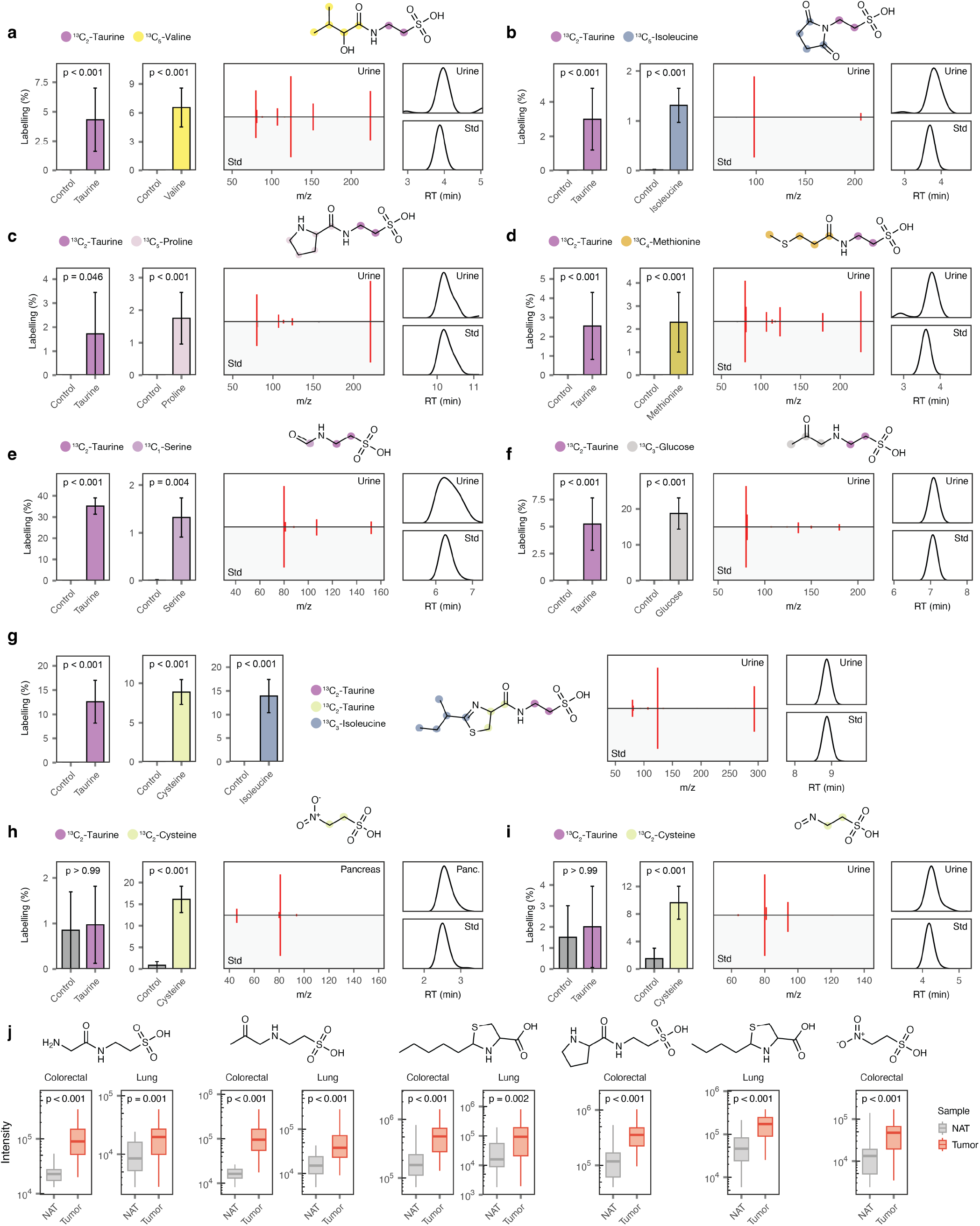
Taurine-related metabolites. **a-i**, Examples of previously unrecognized taurine-related metabolites. Bar charts show fractional labelling after natural isotope abundance correction for selected tracers; inset text shows Bonferroni corrected p-values. Mirror plots and EICs show data from synthetic reference standards versus experimental peaks in mouse tissues. Metabolite structures show positional labelling from ^13^C tracers (manually inferred from the structure and experimental isotope labelling patterns). **a**, *α*-Hydroxyisovaleryltaurine; **b**, taurine succinimide; **c**, prolyltaurine; **d**, 3-methylthiopropionyltaurine; **e**, formyltaurine; **f**, 2-(2-oxopropylamino)ethanesulfonic acid; **g**, (2-(*sec*-butyl)-4,5-dihydrothiazole-4-carbonyl)taurine; **h**, nitrotaurine; **i**, nitrosotaurine. **j**, Abundance of selected metabolites in human colorectal and lung adenocarcinoma versus matched normal adjacent tissue (NAT). Inset text shows p-values after Benjamini-Hochberg correction. From left to right: glycyltaurine; 2-(2-oxopropylamino)ethanesulfonic acid; 2-pentylthiazolidine-4-carboxylic acid; prolyltaurine; 2-butylthiazolidine-4-carboxylic acid; nitrotaurine.

The newly identified taurine metabolites fell into several classes. First, we identified several previously unrecognized N-acyltaurines. In contrast to the long-chain N-acyltaurines described previously, these species bore short-to medium-chain acyl groups (C4-C12). They included branched species such as leucine-derived isobutyryltaurine (C4:0) and valine-derived isovaleryltaurine (C5:0); oxygenated species such as *α*-and *β*-hydroxybutyryltaurine (C4:0;O1), valine-derived *α*-hydroxyisovaleroyltaurine (C5:0;O), *α*-and *β*-hydroxyhexanoyltaurine (C6:0;O1), *β*-hydroxycaprylyltaurine (C8:0;O1), and dodecanedioyltaurine (C12:1;O2); and conjugates to glucose-derived metabolites, including lactoyltaurine, acetoacetoyltaurine, fumaryltaurine, and 2-hydroxyglutaroyltaurine (**Fig. 4a** and **Supplementary Fig. 7**). Moreover, we identified a cyclic metabolite apparently formed by intramolecular condensation of succinyltaurine, in which the distal carboxylate cyclizes onto the amide nitrogen to yield a five-membered ring (**Fig. 4b**).

We also identified a series of amino acid-taurine conjugates. Aminoacyltaurines including *α*-and *γ*-glutamyltaurine, seryltaurine, and *β*-aspartyltaurine have been reported, but this class of metabolites has received little attention^29–31^. We identified seven previously unrecognized aminoacyltaurines, including prolyltaurine, glutaminyltaurine, alanyltaurine, glycyltaurine, threonyltaurine, N-acetylseryltaurine, and N-acetylphenylalanyltaurine (**Fig. 4c** and **Supplementary Fig. 8a-f**). Additionally, we uncovered a structurally related but previously undescribed class of taurine conjugates: acylglycyltaurines, in which taurine is conjugated to an acylglycine rather than a free amino acid. We confirmed the structures of acylglycyltaurines bearing C5:0, C6:0, C6:1, C8:0, and C10:1 acyl chains, as well as acetylglycyltaurine and phenylacetylglycyltaurine, by chemical synthesis (**Supplementary Fig. 8g-m**), and detected (but did not confirm) an additional 10 putative acylglycyltaurines (**Supplementary Table 3b**).

Beyond these major classes, our tracing data revealed taurine conjugates to a diverse range of other metabolites. For instance, a peak incorporating two carbons from taurine and six from tryptophan suggested a conjugate of picolinic acid from the kynurenine pathway; chemical synthesis confirmed this assignment (**Supplementary Fig. 8n**). A second peak incorporated two carbons from taurine and four from methionine, and was identified as 3-methylthiopropionoyltaurine, while a third incorporated a single carbon from serine and was identified as N-formyltaurine (**Fig. 4d-e**). A fourth peak that incorporated two carbons from taurine and three from glucose was identified as 2-(2-oxopropylamino)ethane-1-sulfonic acid, an isomer of propionyltaurine consistent with coupling between methylglyoxal and taurine via reductive amination (**Fig. 4f**). A fifth peak labelling with two carbons from taurine, three from cysteine, and five from isoleucine proved to be a taurine conjugate of the isoleucine-derived thiazoline described above (**Fig. 4g**). Still other taurine conjugates included the taurine conjugate of Smethylthiomercapturic acid and phloretoyltaurine, for which the taurine but not the phloretic acid moiety was detectably labelled (**Supplementary Fig. 8o-p**). The latter was among a handful of metabolites whose levels decreased in mice fed a purified diet lacking phytochemicals, consistent with incorporation of dietary phloretic acid (**Supplementary Fig. 6a,c**). A further 27 peaks demonstrated labelling patterns and putative formulas suggestive of additional N-acyltaurine conjugates (**Supplementary Table 3b**). Thus, the scope of taurine conjugation is far more extensive than previously recognized.

We also detected two peaks whose molecular formulas differed from taurine by the addition of one or two oxygen atoms, but which did not incorporate labelled taurine. Instead, they incorporated two carbon atoms from cysteine, suggesting that they are derived from hypotaurine or another upstream metabolite. Examination of the top-ranked candidate structures revealed taurine derivatives bearing nitroso and nitro functionalities, neither of which has previously been documented. Both predictions were confirmed by chemical synthesis (**Fig. 4h-i**). C-nitroso and aliphatic nitro groups on small polar metabolites are unusual in the context of mammalian metabolism, further underscoring the power of isotope tracing to reveal unexpected transformations of labelled precursors.

The vast majority of these taurine-related metabolites appeared to be host-derived. The lone exception was 2-(2-oxopropylamino)ethane-1-sulfonic acid, which was reduced by more than two-fold after antibiotic treatment, consistent with a microbial contribution to its biosynthesis (**Supplementary Fig. 6a-b**). In contrast, many taurine-related metabolites were elevated in antibiotic-treated mice, although taurine itself was unchanged. For instance, both taurine-6:0;O1(*α*) and taurine-6:0;O1(*β*) were increased by greater than threefold in the gastrointestinal tract, suggesting that the microbiome may consume these metabolites or their precursors.

Collectively, these findings substantially expand the known repertoire of taurine-derived metabolites and establish several previously undescribed metabolite families, including short-chain N-acyltaurines, acylglycyltaurines, and N-oxidized taurines.

### Metabolite conservation and cancer associations

We next sought to determine whether these metabolites are specific to mice or more broadly conserved across mammals. To address this question, we performed metabolomic analysis of urine samples from five mammalian species, including humans (**Supplementary Fig. 9a**). The vast majority of the metabolites detected in mouse urine were also found in other mammals, and several—such as taurine-C5:0, lactoyltaurine, and N-formyltaurine—were detected in all species (**Supplementary Fig. 9b**). Thus, with rare exceptions, the newly identified metabolites are not unique to mice but instead are conserved across mammalian metabolism.

The widespread detection of these metabolites led us to search MS/MS spectra for these metabolites against a compendium of published metabolomic experiments encompassing 152,743 samples from human tissues and cell lines. MS/MS spectral matching suggested that many of these metabolites had been detected, but not identified, in prior human metabolomics experiments (**Supplementary Fig. 9c-d** and **Supplementary Table 4**).

Despite their broad conservation, several of these metabolites showed pronounced sexual dimorphism in mice. In particular, a number of cysteine-, taurine-, and mevalonate-derived metabolites were considerably more abundant in males (**Supplementary Fig. 6a,d**).

Finally, we asked whether any of these metabolites are altered in human cancer. We performed metabolomic profiling of tumors and matched adjacent normal tissues from patients with colorectal (*n* = 41) or lung (*n* = 28) adenocarcinoma. Many of the newly identified metabolites were elevated in tumors (**Fig. 4j** and **Supplementary Fig. 6a,e**). Cysteine-derived thiazolines and thiazolidines, for instance, were upregulated in both colorectal and lung cancer. Most aminoacyltaurines were likewise elevated in these cancers. Only a single metabolite was decreased: (3-cyanopropanoyl)(sulfo)alanylglycine, in lung adenocarcinoma. These findings indicate that the metabolic pathways giving rise to these metabolites may be engaged in human malignancy.

### Depletion of mevalonate-derived metabolites in aging

Aging alters the levels of many metabolites. Integrating data from both tissues and serum, we assessed age-related changes in the levels of all 4,506 putative metabolite peaks. Across all putative metabolites, the two most significantly depleted metabolites with age were both previously undetermined structures that were elucidated through isotope tracing: S-methylthiomethyl mercapturic acid and 2,3-dihydrofarnesoic acid (**Fig. 5a**). These compounds are structurally unrelated but have similar tissue distributions, with levels highest in the kidney, and both decreased with age in every tissue examined (**Fig. 5b**).

**Fig. 5.**
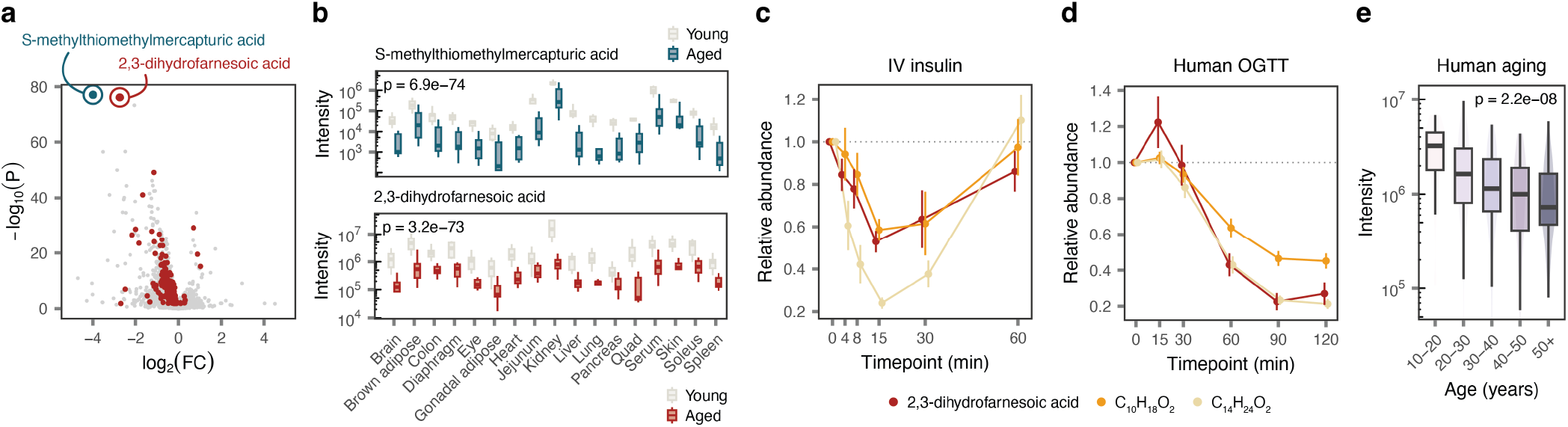
Mevalonate-derived metabolites are depleted in aging. **a**, Volcano plot comparing relative metabolite levels in young (4 months) versus aged (24 months) C57BL/6J mice (data are from 17 tissues with 9–15 independent mice per solid tissue and 30 for serum). The two most significantly depleted metabolites, 2,3-dihydrofarnesoic acid and S-methylthiomethyl mercapturic acid, are circled in red and blue, respectively; other age-responsive metabolites labelling with two to four carbons from ^13^C mevalonate are colored in red. **b**, Tissue levels of S-methylthiomethyl mercapturic acid (top) and 2,3-dihydrofarnesoic acid (bottom) across 17 tissues in young and aged C57BL/6J mice. Inset text shows p-values from the moderated t-test implemented in limma. **c**, Serum levels of mevalonate-derived metabolites in young mice following an intravenous bolus of insulin. Points and error bars show mean and standard error, respectively. Values are normalized to baseline measurements from each animal (*n* = 3). **d**, As in **c**, for an oral glucose tolerance test (OGTT) in humans (*n* = 34). **e**, Serum 2,3-dihydrofarnesoic acid levels across age from a cross-sectional human cohort study (*n* = 395). Inset text shows p-value from a two-sided test of Pearson’s correlation coefficient.

Because 2,3-dihydrofarnesoic acid is synthesized from mevalonate, we asked whether other mevalonate-derived metabolites were also affected by aging. More than 200 mevalonate-derived metabolites were significantly depleted in mouse aging, whereas only a handful were increased. Two prominent examples of abundant, strongly age-depleted compounds included 10-carbon (C_10_H_18_O_2_) and 14-carbon (C_14_H_24_O_2_) analogues of 2,3-dihydrofarnesoic acid whose structures remain under investigation (**Supplementary Fig. 10a**). These compounds, but not S-methylthiomethyl mercapturic acid, were readily detectable in human serum.

Beyond their depletion in aging, mevalonate-derived metabolites were acutely responsive to insulin and glucose. An intravenous bolus of insulin in mice rapidly and robustly suppressed the serum levels of 2,3-dihydrofarnesoic acid, C_10_H_18_O_2_, and C_14_H_24_O_2_ (**Fig. 5c**). Moreover, all three decreased in a human oral glucose tolerance test (**Fig. 5d**). In an observational human cohort, the C_10_H_18_O_2_ species did not change with age, but both 2,3-dihydrofarnesoic acid and the C_14_H_24_O_2_ species were significantly depleted (**Fig. 5e** and **Supplementary Fig. 10b**)^32^. Thus, the isoprenoid 2,3-dihydrofarnesoic acid and other mevalonate-derived metabolites decline with age across mice and humans.

### Age-related impairment of CoA synthesis

We next investigated the biochemical basis of 2,3-dihydrofarnesoic acid depletion in aging. We infused multiple stable isotope-labeled precursors in young and aged mice and found that generation of 2,3-dihydrofarnesoic acid and related species from major circulating nutrients (3-hydroxybutyrate, glucose and glutamine) was significantly decreased in aged mice. Upon infusion of labeled mevalonate, however, aged mice paradoxically produced more labeled 2,3-dihydrofarnesoic acid (**Fig. 6a**). These data suggested that the age-associated deficit in 2,3-dihydrofarnesoic acid generation lies between the major circulating nutrients and mevalonate.

**Fig. 6.**
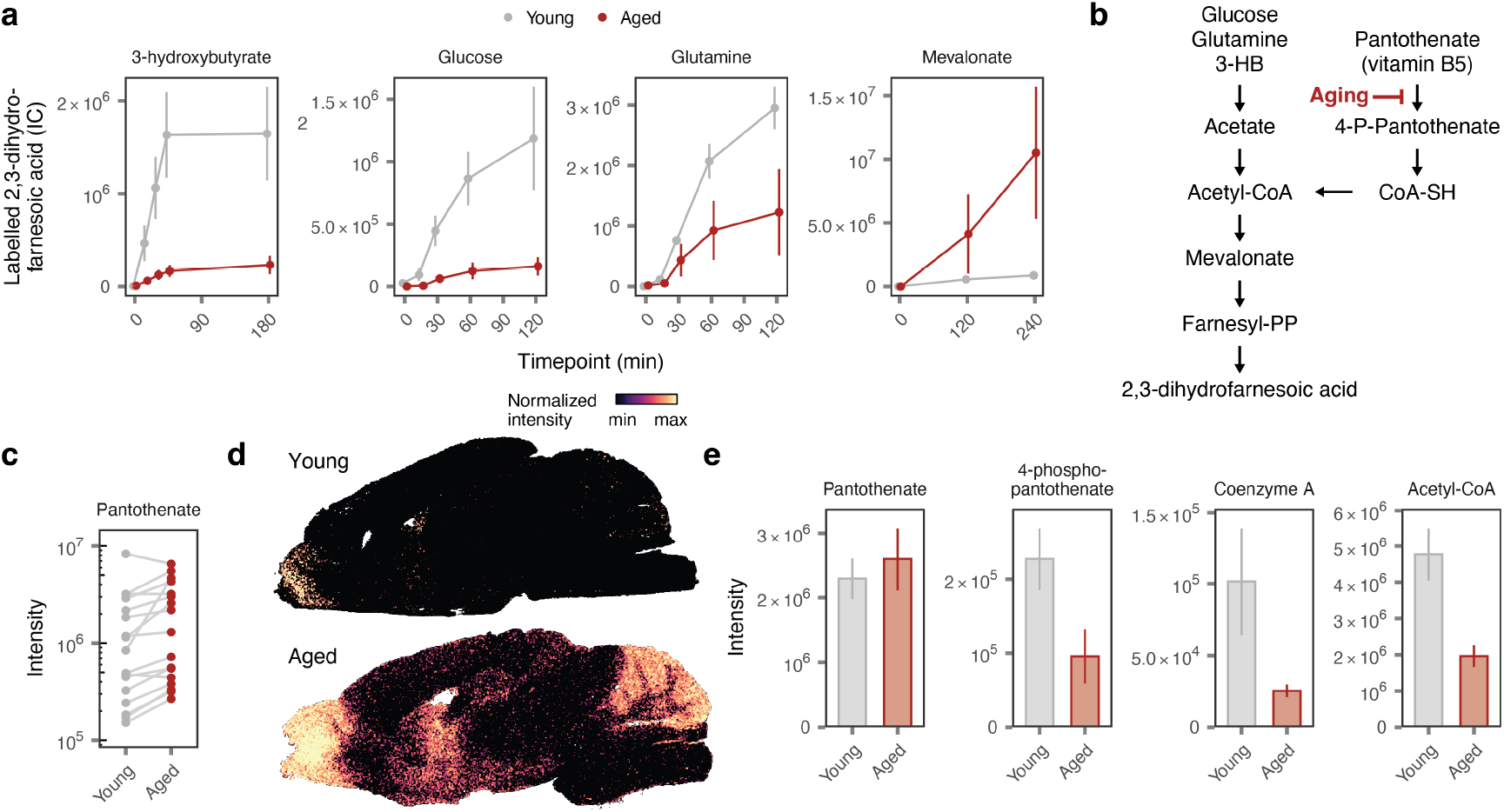
A deficit in CoA synthesis drives depletion of mevalonate family metabolites in aging. **a**, Serum levels of ^13^C-labeled 2,3-dihydrofarnesoic acid in young (3–6 month) and aged (20–24 month) C57BL6/J mice after infusion of ^13^C-labeled 3-hydroxybutyrate, glucose, glutamine or mevalonate. Points and error bars show mean and standard error of the mean, respectively (*n* = 3–11). **b**, Proposed pathway for 2,3-dihydrofarnesoic acid production. **c**, Pantothenate levels in young (4 month) and aged (24 month) C57BL/6J mouse tissues. Each point shows mean per tissue (*n* = 17 tissues). **d**, Spatial metabolomic measurements of pantothenate levels in representative young and aged C57BL/6J mouse brains. See **Supplementary Fig. 10** for additional replicates. **e**, Pantothenate, 4-phospho-pantothenate, coenzyme-A (CoA) and acetyl-CoA levels in liver samples of young and aged C57BL/6J mice. Bars and error bars show mean and standard error, respectively (*n* = 7–10). For associated pantothenate tracing data, see **Supplementary Fig. 10**.

Circulating nutrients are converted into mevalonate via acetyl-CoA (**Fig. 6b**). Synthesis of CoA begins with phosphorylation of the essential vitamin B5 (pantothenate), which is one of the most strongly increased metabolites in mouse aging^33^; its levels rose substantially in serum and most tissues, with focal brain accumulation in the aged olfactory bulb, striatum and cerebellum (**Fig. 6c,d** and **Supplementary Fig. 10c-d**). In contrast, downstream intermediates in the CoA biosynthesis pathway, beginning with 4-phosphopantothenate, were depleted in aging (**Fig. 6e**). Infusion of labeled pantothenate confirmed defective synthesis of 4-phosphopantothenate, CoA and acetyl-CoA in aged mice (**Supplementary Fig. 10e**).

Thus, working backwards from the discovery of 2,3-dihydrofarnesoic acid, we identified defective CoA synthesis as a hallmark of aging.

## Discussion

Metabolomic surveys of mammalian tissues routinely detect thousands of peaks that cannot be matched to known metabolites, but it has remained unclear how many of these unidentified peaks arise from endogenous mammalian metabolism or instead derive from exogenous sources. Here, we demonstrate the use of systematic *in vivo* isotope tracing to probe the biosynthetic origins of putative unknown metabolites on a metabolome-wide scale. By mapping the fates of 26 labelled precursors in mice, we generated a tracing resource of unprecedented scope that enabled annotation of the biosynthetic origins for approximately 3,500 putative metabolites. This resource provides a new lens through which to interrogate the mammalian metabolome and, as we demonstrate, a springboard for metabolite discovery.

Mounting interest in the dark metabolome has motivated efforts to expand reference MS/MS libraries and the development of new experimental and computational approaches for annotating MS/MS spectra in metabolomics datasets^3,34–39^. Isotopic labelling patterns, which encode the biosynthetic precursors of an unidentified metabolite, provide an orthogonal basis for structure elucidation. To harness this information, we developed a statistical framework to infer isotopologue-level patterns of precursor incorporation into unidentified metabolites, coupled with a multimodal contrastive learning model that co-embeds these patterns with chemical structures in a shared latent space. The latter approach allows for any arbitrary set of structures, whether drawn from existing databases or generated *in silico*, to be ranked according to their compatibility with a given set of biosynthetic precursors. The current performance of our model, Isopleth, is constrained by its relatively small training set of known metabolites, and should improve as a greater fraction of the metabolome is structurally characterized. Nonetheless, Isopleth already demonstrates a meaningful capacity for structure elucidation, even in the absence of any other analytical data, and complements established approaches based on MS/MS spectra and retention times.

Interrogation of the tracing data led to the discovery of 54 previously unrecognized mammalian metabolites spanning a number of families, including cysteine-derived alkylthiazolidines, dithioacetal mercapturic acid derivatives, short-chain N-acyltaurines, acylglycyltaurines, and N-oxidized taurines (**Supplementary Table 5**). Many are derived from simple, abundant precursors such as cysteine or taurine, whose metabolism has been studied for decades. That these metabolites have eluded discovery until now underscores how much of the metabolome likely remains to be characterized. Several of these species bear unexpected functional groups, such as nitriles, aliphatic nitro or nitroso groups, dithioacetals, and thiazolidines, that have rarely or never been described in the context of endogenous mammalian metabolism. These discoveries thus expand the known biochemical repertoire of the mammalian metabolome.

Many of these metabolites are enriched in particular tissues and dysregulated in aging or human cancer, suggesting active regulation of their synthesis or degradation. Among them, 2,3-dihydrofarnesoic acid stands out for its high flux, insulin-responsiveness, and marked depletion in aging in both mice and humans. Collectively, these observations suggest that 2,3-dihydrofarnesoic acid may play an important yet previously unappreciated role in mammalian metabolism, perhaps by providing a release valve for the cholesterol biosynthetic pathway. While the precise nature of this role remains to be determined, the structure elucidation of 2,3-dihydrofarnesoic acid shed immediate light on the deleterious metabolic effects of aging. In particular, our work revealed age-associated impairment in the synthesis of CoA, a fundamental cofactor required for both fat and cholesterol synthesis and oxidative energy generation. This example illustrates how systematic isotope tracing can guide the structure elucidation of metabolites with strong phenotypic associations and thereby reveal new opportunities for therapeutic metabolic intervention.

Some limitations of this work should be noted. In some cases, not all carbon atoms in a confirmed metabolite labelled from one of the administered precursors. Some of these cases likely represent false negatives, reflecting slow turnover or dilution from unlabeled precursor pools; others may reflect contributions from precursors not included in our tracing experiments. Conversely, our statistical approach to inferring isotopologue-level labelling patterns also has a nonzero falsepositive rate, which reflects the sheer number of isotopologues monitored and the potential for interference from unrelated signals with similar mass-to-charge ratios. Our tracing studies focused on male mice and negative ionization mode data, which likely overlooked additional metabolite families that are specific to females or ionize preferentially in positive mode. Our metabolite discovery campaign incorporated human oversight in prioritizing structures for synthesis, and we expect that the methods described here will continue to be used in collaboration with expert chemists, particularly given the resources required for chemical synthesis of reference standards. Finally, our work reveals previously unknown metabolites with unprecedented functional groups and disease associations, but the enzymes catalyzing the synthesis of these species remain to be elucidated. Further work will be needed to discover these enzymes and establish the roles of these metabolites in physiology and disease.

## Methods

### Experimental models

Animal studies were conducted in accordance with protocols approved by the Princeton University Animal Care and Use Committee. All mouse studies were performed using 10–14 week old C57BL/6N mice (Charles River Laboratories), except for aging studies where both aged animals and young controls were C57BL/6J mice (Jackson Laboratories). Mice used in aging studies were purchased from Jackson Laboratories at 8–24 weeks of age and aged in the Princeton University animal facility until used for experiments at 18–24 months of age. For isotope infusion studies, aseptic surgery was performed to insert catheters (Instech Laboratories) into the right jugular vein with access via button implanted under the skin on the back. Mice were allowed at least 5 days of recovery before experiments were performed. Catheterized mice were individually housed with enrichment including nestlets and houses. Mice had *ad libitum* access to food (LabDiet PicoLab Rodent 20 5053) placed on the floor and to water. Animals were kept on a forward light cycle (light period 8 am – 8 pm) in a housing facility maintained between 20–22 °C at 40–70% relative humidity. For aging studies not involving isotope infusion, mice were housed in groups of up to 5 per cage.

For infusions lasting 6 hours or less, mice were fasted for 2 hours starting at approximately 9 am before starting the infusion and remained fasting for the duration of the infusion. For infusions lasting longer than 6 hours, mice had *ad libitum* access to food throughout. Free fatty acids were infused with 1 mM essentially fatty acid-free bovine serum albumin. Tracer details and parameters for all experiments are listed in **Supplementary Table 2a**. Tracer infusates were prepared in sterile saline and sterile filtered before infusing. The saline concentration was adjusted to ensure that the final infusate was isosmotic with blood. Blood samples were taken via tail snip and collected into lithium heparin tubes. Samples were kept on ice for no more than 60 minutes before centrifuging at 10,000×g for 10 minutes at 4 °C. Plasma was transferred to a new tube and stored at –80 °C until analysis. At the end of the infusion period, mice were euthanized by cervical dislocation and tissues were rapidly dissected by teams of 3–4 researchers, wrapped in foil and clamped with a precooled Wollenberger clamp in liquid nitrogen. All tissue collections occurred at approximately 4 pm (ZT 8). Tissue samples were stored at –80 °C until analysis. Stable isotope tracers were delivered via intravenous infusions except fructose (oral gavage) and pantothenate, riboflavin, and thiamine (drinking water for 5 days). Studies involving stable isotope tracers were performed in male mice.

### Aging mouse studies

For analyses of metabolite levels in aging mouse tissues, a cohort of 4 month old (young) and 24 month (aged) C57BL/6J male mice were euthanized after 6 hours of fasting. Sixteen tissues (brain, brown adipose, colon, diaphragm, eye, gonadal adipose, heart, jejunum, kidney, liver, lung, pancreas, quadriceps, skin, soleus, and spleen) were rapidly collected, wrapped in foil and clamped with a precooled Wollenberger clamp in liquid nitrogen. Blood (~20 *µ*L) was collected from the tail vein before euthanasia, allowed to clot for 5 min at room temperature, centrifuged at 15,000×g for 15 minutes at 4 °C and serum was collected and stored at –80 °C. For aging infusion studies, protocols were followed as described in the section above but both aged and control mice were allowed to recover for at least 10 days to accommodate longer recovery time in aged mice. Infusion parameters for aging infusion studies are listed in **Supplementary Table 2b**.

### Human cancer samples

Pancreatic and colorectal cancer samples were obtained from the Rutgers Cancer Institute Biospecimen Repository and Histopathology Service (RCI BRHS). Lung cancer patient samples were acquired from the Fox Chase Cancer Center Biosample Repository Facility (FCCC BRF). Both institutions maintain longstanding Institutional Review Board (IRB)-approved protocols for the collection, banking, and distribution of de-identified biospecimens. Informed consent and authorization for unrestricted research use were obtained from all participants prior to specimen collection.

### Human serum samples

The human cohort used for plasma metabolomic analysis as a function of age involved non-pastoralist Northern Kenyan individuals from the Turkana Health and Genomics Project^32^. The human oral glucose tolerance test study was conducted in Australia following IRB-approved protocols^40^. Male and female participants between the age of 18 and 65 (median = 45.9) with BMI of 25.8–40 (median = 34.5) were selected with elevated triglycerides (triglyceride levels > 2.8 mmol/L), prediabetes, or well controlled diabetes. Exclusion criteria included pregnancy, breastfeeding, or terminal illness with less than 1 year expected survival. Participants were fasted overnight prior to a fasted blood draw. Following fasting blood draw participants were given an oral glucose solution and plasma was collected in heparin coated tubes at 15, 30, 60, 90, and 120 min post glucose ingestion.

### Metabolite extraction

Frozen solid tissue samples were weighed to approximately 40 mg aliquots per sample and transferred to 2 mL Eppendorf tubes on dry ice. Samples were homogenized to a fine powder using a cryomill (Retsch) cooled with liquid nitrogen. Metabolites were extracted by adding 1 mL of ice-cold 40:40:20 acetonitrile:methanol:water (v/v/v) with 0.5% formic acid per 30 mg of tissue. Samples were vortexed and incubated on ice for 10 min, after which 85 *µ*L of 15% ammonium bicarbonate (w/v) was added and vortexed to neutralize the acid. Following an additional 10 min incubation on ice, samples were centrifuged at 14,000 rpm for 25 min at 4 °C. The supernatants were transferred to fresh tubes and centrifuged once more under the same conditions, and the resulting supernatants were collected for analysis.

For serum and urine, frozen samples were thawed on ice. Protein was precipitated by adding 200 *µ*L of methanol to 10 *µ*L of sample, followed by vortexing for 10 seconds and centrifugation for 25 min. The supernatants were collected, dried under a nitrogen stream, and reconstituted in 200 *µ*L of 40:40:20 acetonitrile:methanol:water (v/v/v) prior to analysis.

### LC–MS analysis

Untargeted LC–MS analysis was performed on a Vanquish UHPLC system coupled with an Orbitrap Exploris 480 MS (Thermo Fisher Scientific). Chromatographic separation was achieved using a Waters XBridge BEH Amide column (2.1 × 150 mm, 2.5 *µ*m particle size) maintained at 25 °C. The injection volume was 5 *µ*L. Mobile phase A consisted of 95:5 water:acetonitrile (v/v) with 20 mM ammonium hydroxide and 20 mM ammonium acetate (pH 9.4), while mobile phase B was 100% acetonitrile.

The flow rate was set to 150 *µ*L/min, and the gradient program was set as follows: 0-2 min, 90% B; 2–3 min, 90%-75% B; 3–7 min, 75% B; 7–8 min, 75%-70% B; 8–9 min, 70% B; 9–10 min, 70%-50% B; 10–12 min, 50% B; 12–13 min, 50%–25% B; 13–14 min, 25% B; 14–16 min, 25%–0% B; 16–20.5 min, 0% B; 20.5–21 min, 0%–90% B; and 21–25 min, 90% B for re-equilibration.

For non-aging isotope tracing studies and analysis of tissue specificity and sexual dimorphism, the MS was operated in negative-scan mode. For other studies, dual-scan mode was used. Instrument settings were as follows: for the full scan, the resolution was set to 120,000 (except for tracing experiments, for which the resolution was set to 480,000) with a scan range of 70–1,000 m/z. The AGC target was set to 10^7^ and the maximum ion injection time (ITmax) to 200 ms. The spray voltage was set to 3,200 V and 3,000 V for positive and negative ionization mode, respectively. Source parameters included: Sheath gas at 35 Arb, Aux gas at 10 Arb, and Sweep gas at 0.5 Arb. The ion transfer tube was held at 300 °C and the vaporizer temperature at 35 °C. Internal mass calibration was enabled, and the RF lens was set to 60%. For data acquisition XCalibur (Version 4.4) was used and LC–MS data was stored as Thermo raw files. Quantification of coenzyme-A and acetyl-CoA used the same parameters, except only positive-scan mode was used with scan range of 700–950 and RF lens set to 100%.

### Mass spectrometry imaging and analysis

Mouse brains were excised from the skulls of 2-month (young, *n* = 3) and 26-month (aged, *n* = 3) C57BL/6J mice within 1 min after euthanasia and frozen on a weigh boat on liquid nitrogen. Frozen brain tissues were stored at –80 °C until slides were prepared. Tissues were equilibrated in a cryostat (Leica, CM3050S) at CT –18 °C/OT –16 °C for at least one hour before cutting into two hemispheres. A hemisphere was attached to the specimen disc with OCT and cut at 10 *µ*m thickness in the sagittal plane to obtain sections referring to image 14-15 of 21 bregma level from the Allen Mouse Brain Atlas ID 100042147. Sections were thaw mounted onto indium tin oxide (ITO)-coated slides (CG-90IN-S110, Delta Technologies Limited). Slides were dried in a vacuum desiccator for 15 min before spray coated using an automated HTX M3+ sprayer (HTX Technologies) with N-(1-naphthyl) ethylenediamine dihydrochloride (NEDC) (Sigma-Aldrich, 222488) matrix solution at 10 mg/mL in 70% v/v methanol/water for matrix density of 4.17 × 10^−3^ mg/mm^2^. The sprayer was set to nozzle temperature 75 °C, gas pressure 10 psi, flow rate 50 *µ*L/min, nozzle velocity 1200 mm/min, track spacing 2 mm, 20 passes, pattern CC, and drying time 10 s.

Samples were run in the timsTOF fleX MALDI-2 (Bruker Daltonics) with pixel area of 10 × 10 *µ*m^2^ resolution. Laser power and number of shots at a frequency of 10 kHz were optimized for signal intensity without oversampling. MS scan acquisition was performed in negative ion polarity and scan range 50-1050 m/z. MS Tune was set to Collision Cell energy 8 eV, Collision RF 500 Vpp, Quadrupole ion energy 5 eV, low mass 20 m/z, Focus Pre TOF transfer time 35 *µ*s, and pre-pulse storage 2.6 *µ*s.

Data was exported from Bruker SCiLS as pixel coordinates and intensities, which was then processed with R/RStudio. To remove batch effects across runs, a reference section from a refrozen homogenized 2-month-old mouse brain hemisphere was included with each run. Each feature was normalized to report intensities as relative to the mean intensity of the same reference section. K-means clustering was applied on pixels for each sample, and the cluster of off-tissue pixels were removed. Pantothenate was detected at 218.1034 m/z as an [M–H]^−^ adduct. The spatial image for pantothenate was produced after removing hotspots by capping the maximum intensity at the 99th percentile intensity across all six young and aged samples. Intensity was plotted using the exported (x, y) coordinates and normalized intensity using the magma color scale.

### Peak picking and untargeted data processing

Peak picking was performed in mzMine^41^ using a customized batch workflow. Mass detection was applied with a noise level of 2 × 10^4^, followed by the FTMS shoulder peaks filter (mass resolution 480,000, Gaussian). Chromatograms were constructed using the ADAP chromatogram builder (minimum consecutive scans 5, minimum intensity 2 × 10^4^, minimum absolute height 5 × 10^4^) and smoothed using a Savitzky-Golay filter (5 scans). Peaks were resolved using the local minimum feature resolver (chromatographic threshold 85%, minimum relative height 0%, minimum absolute height 5 × 10^4^, minimum peak-to-edge ratio 3, minimum scans 3). Gap-filling was performed using the peak finder (intensity tolerance 20%, retention time tolerance 0.1 min). Peak lists were subsequently filtered using the duplicate peak filter (retention time tolerance 0.3 min) and feature list rows filter, retaining only features present in all samples. These steps resulted in a data matrix for 8,277 peaks. Peak annotation was performed using NetID in R (https://github.com/LiChenPU/NetID) with default parameters^15^. The peak matrix generated by mzMine was used as an input, and no MS/MS spectra or external spectral libraries were used. NetID annotated features by constructing a global annotation network based on accurate mass differences, retention time proximity, and expected isotope, adduct, and biochemical transformation relationships. The Human Metabolome Database (HMDB) served as a library for the annotation of known metabolites^42^. No manual curation or parameter optimization was performed.

Isotope annotations were inferred from characteristic mass differences (for example, ^13^C, +1.0034 Da; ^34^S, +1.9959 Da) and expected natural abundance patterns. Elemental atom counts were inferred from the relative abundances of corresponding isotopologues.

Isotopologue matrices were constructed by extracting isotopologue intensities for each of the 8,277 signals detected in our mouse tissue dataset. For each signal, the number of carbon atoms was obtained from the molecular formula annotated by NetID. When formula annotation was unavailable, the carbon number was estimated from the measured m/z using an empirical approximation, with an upper bound of 30 carbons. Based on the inferred carbon count, the expected m/z values of the corresponding ^13^C isotopologues (M_0_ through M_n_, where n equals the number of carbons) were calculated and used for targeted signal extraction.

To account for retention time shifts between datasets, the monoisotopic (M_0_) signal was first located using a wide retention time window (± 1.0 min) after smoothing with a three point moving average. The observed retention time was used to refine the signal position, and isotopologue intensities were subsequently extracted using a smaller retention time tolerance (± 0.05 min). If the monoisotopic peak was not detected within the initial window, isotopologue extraction was not performed for that peak. Natural isotope abundance correction was then performed using the Autocorr package (https://github.com/xxing9703/Iso-Autocorr), as previously described^43^.

### Targeted MS/MS acquisition

For each peak of interest, signal intensities were extracted from the full-scan data across the 25 tissue extracts to identify the tissue with the highest abundance. Targeted MS/MS was then performed on the corresponding extract. Samples were analyzed in a single LC–MS run using a full scan followed by targeted MS/MS scans guided by an inclusion list. Full-scan parameters were set to a resolution of 60,000, a scan range of 70-1,000 m/z, an AGC target of 10^7^, and an ITmax of 200 ms. MS/MS parameters were an isolation window of 1.0 m/z, collision energies of 15, 20, and 30 eV, resolution of 15,000, AGC target of 10^6^, an ITmax of 100 ms, and a retention time (RT) window of 3 min.

Due to the complexity of biological samples and the presence of chimeric spectra containing fragments from multiple precursor ions within the same isolation window, fragment ions were deconvoluted. This was achieved by calculating the Pearson correlation between MS1 and MS/MS extracted ion chromatograms (EICs). A fragment ion was considered valid only if its MS/MS EIC correlated with the MS1 EIC of the precursor ion. Each precursor ion was associated with multiple MS/MS scans across a retention time (RT) window of up to 3 min. The scan corresponding to the maximum MS1 intensity was selected as the representative precursor scan. A chromatogram was constructed for each fragment ion in this MS/MS spectrum and correlated with the MS1 EIC of the precursor. Scan times were aligned via interpolation and restricted to a 0.3 min RT window centered on the MS1 peak. Fragment ions with a Pearson correlation coefficient below 0.8 were excluded.

### Metabolite standards

Synthetic standards for candidate metabolite structures were either purchased from commercial suppliers or obtained via chemical synthesis. Catalogue numbers for all commercially available compounds are provided in **Supplementary Table 5**, while detailed protocols of synthesized standards are outlined in **Supplementary Note 1**. Because stereoisomers are not expected to be distinguishable by our LC–MS/MS metabolomics approach, structures are drawn with undefined stereochemistry in both the main text and **Supplementary Note 1** except for chiral building blocks in the latter.

Standards were dissolved in 40:40:20 (v/v/v) acetonitrile:methanol:water to a concentration of 1 mg/mL. Stock solutions were subsequently diluted to 2 *µ*g/mL in the same solvent mixture for full-scan LC–MS analysis. For each standard, the representative mouse tissue extract (the tissue showing the highest intensity signal for a given metabolite) was analysed side-by-side. Retention times and MS/MS spectra were compared as described in the section above, with the exception of the ITmax being set to 100 ms.

To achieve unambiguous metabolite identification, we employed multiple orthogonal physicochemical criteria. High-confidence assignments were based on a combination of retention time matching across disparate stationary phases and MS/MS. We utilized both HILIC (BEH Amide) and reversed-phase liquid chromatography (RPLC) to exploit their distinct separation mechanisms. The HILIC method is described above. RP separation was achieved using a Waters Atlantis T3 column (2.1 × 150 mm, 3 *µ*m particle size) maintained at 25 °C. The injection volume was 5 *µ*L. Mobile phase A consisted of 98:2 water:acetonitrile (v/v) with 10 mM ammonium acetate and 0.1% acetic acid, while mobile phase B was 100% acetonitrile. The flow rate was set to 200 *µ*L/min, and the gradient program was set as follows: 0-3 min, 0% B; 3-9 min, 0%-70% B; 9-11 min, 70%-100% B; 11-19 min, 100% B; 19-20 min, 100%-0% B; and 20-25 min, 0% B for re-equilibration. Identification confidence was further supported by switching between positive and negative ionization modes where applicable and acquiring MS/MS. Positive mode MS/MS spectra are shown in the figures for the following compounds: 2-propylthiazolidine-4-carboxylic acid, 2-butylthiazolidine-4-carboxylic acid, 2-pentylthiazolidine-4-carboxylic acid, 2-hexylthiazolidine-4-carboxylic acid, 2-heptylthiazolidine-4-carboxylic acid, 2-octylthiazolidine-4-carboxylic acid, 2-nonylthiazolidine-4-carboxylic acid, and 2-(*sec*-butyl)-4,5-dihydrothiazole-4-carboxylic acid.

Thermo raw files were then analysed by the Xcalibur QualBrowser (with nine-point Gaussian smoothing) to determine the retention time for each synthetic standard and visualize the MS/MS spectra for the standards and corresponding metabolite peaks in the tissue samples. Raw data was transformed into mzML format using ProteoWizard (Version 3.0), and spectral quality was ensured by removing contaminating fragment ions from co-isolated precursors. Specifically, we filtered fragments that showed low correlation with the precursor m/z by applying the following criteria: (1) removal of internal mass calibration ions; (2) exclusion of fragments within the 1 m/z precursor to eliminate non-neutral loss signals; (3) manual adjustment of MS1/MS2 correlation thresholds based on the retention time, signal intensity and coeluting peaks; and (4) removal of MS2 fragment ions with absolute intensities exceeding 120% of the MS1 precursor intensity.

### Statistical analysis of isotopologue-level labelling patterns

To infer precursor incorporation into unidentified metabolites, we developed a statistical framework to test for differences in isotopologue intensities between mice given isotopically labelled nutrients and untreated controls. Specifically, we fit a linear model to the isotopologue-level matrix of corrected labelling percentages using limma^44^. For each isotopologue, we modelled the corrected labelling percentage as a function of the tracer condition and the tissue in which the measurement was obtained:

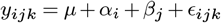

where *y*_*ijk*_ is the corrected labelling percentage for isotopologue *k* in tissue *i* from a mouse treated with tracer *j*, the intercept *µ* represents the expected corrected labelling percentage in a reference tissue from control mice, *α*_*i*_ is the effect of tissue *i, β*_*j*_ is the effect of tracer *j* relative to control mice, and *ϵ*_*ijk*_ is the residual error. The coefficients *β*_*j*_ were estimated using ordinary least squares via lmFit, then moderated using the empirical Bayes procedure implemented in eBayes, testing the null hypothesis *β*_*j*_ = 0. This procedure borrows information across isotopologues to obtain more stable variance estimates. Data from two non-^13^C-labelled tracers, ^15^N-uridine and ^15^N-inosine, was appended to the data matrix prior to model fitting in order to increase the number of observations per isotopologue and thereby increase the residual degrees of freedom for variance estimation.

To evaluate the performance of this approach, we curated a groundtruth reference set of isotopologue-level labelling events through manual inspection of extracted ion chromatograms (EICs) for 123 known metabolites identified by precursor m/z and retention time matches to authentic standards. EICs were extracted with a m/z tolerance of 3 ppm and a retention time window of ±1 min. We manually inspected the EIC for the ^12^C parent versus each possible isotopologue of these metabolites across all tissues and tracers simultaneously, in order to distinguish true labelling events from contaminating signals within the same m/z and retention time window, a process that entailed the inspection of approximately 1.2 million EICs. Particular attention was paid to contaminating signals that demonstrated subtle shifts in retention time relative to the ^12^C parent. Further evaluation criteria included chromatographic peak shape (symmetry, baseline resolution, and consistency across tracing experiments). The biochemical plausibility of a given labelling event was also accounted for, when applicable (for instance, labelling of kynurenine pathway metabolites from tryptophan is consistent with the established biochemistry of this well-studied pathway). This reference dataset was established in a blinded manner, prior to the development of the statistical model or any inspection of its output. We then computed p-values for each tracer-isotopologue combination for all 8,277 peaks in the dataset, converted p-values to binary labelling calls using a family-wise error rate threshold of 0.05, and computed the accuracy of these calls with respect to the ground-truth annotations for the 123 manually curated peaks. Additionally, we computed receiver operating characteristic (ROC) and precision-recall (PR) curves, ranking isotopologue-level labelling events by the negative logarithm of the uncorrected p-value.

To understand which aspects of data preprocessing, labelling quantification, and modelling decisions most impacted the performance of our statistical model, we carried out a series of ablation studies. We found that performance was maximized when using narrow retention time windows to extract isotopologue intensities and when modelling corrected labelling percentages rather than absolute intensities, corrected absolute intensities, or uncorrected labelling percentages. Performance was also maximized when fitting a model jointly to data from all tissues and tracers, relative to models fit to data from mice treated with a single tracer versus control, from individual tissues, or from individual tissue-tracer combinations. Filtering isotopologues with poor peak shape correlations to the ^12^C parent did not improve performance, likely because the comparison of treated versus control samples already distinguished true labelling events from contaminating signals and because of the presence of noisy EICs for low-intensity isotopologues.

### Biosynthesis-aware structure prioritization with Isopleth

We developed an approach to associate chemical structures with isotopic labelling patterns using contrastive learning. Our model, Isopleth, maps labelling patterns and chemical structures into a shared embedding space, placing labelling patterns for each metabolite near their corresponding structures while pushing unpaired patterns and structures apart. At inference time, candidate structures for an unidentified peak are ranked by their cosine similarity to the observed labelling pattern within this learned embedding space. Thus, unlike conventional approaches that rely primarily on the interpretation of MS/MS spectra, Isopleth scores candidate structures based on isotopologue-level patterns of biosynthetic precursor incorporation, as revealed through systematic isotope tracing experiments.

Isopleth consists of two encoder networks that project isotopic labelling patterns and molecular structures into a shared *d*-dimensional embedding space (**Fig. 2a**). The structure encoder takes as input a Morgan fingerprint representing the molecule as a fixed-length binary vector; here, Morgan fingerprints with a radius of 3 and a length of 2,048 bits were computed using RDKit^45^. The encoder consists of a two-layer fully connected network with ReLU activation and a hidden dimension of 1,024. The isotopic labelling pattern encoder takes as input a binary vector denoting which isotopologue-level patterns of precursor incorporation were statistically significant for a given peak. This encoder consists of a four-layer fully connected network with ReLU activation. Both encoders output 1,024-dimensional embeddings, which are then L2-normalized.

Isopleth is trained by minimizing a symmetric contrastive loss, inspired by approaches such as CLIP that jointly embed image-caption pairs in a shared embedding space^16,46^. Given a batch of *N* structure-pattern pairs, we compute a similarity matrix *S* where 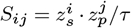 represents the scaled cosine similarity between structure embedding *i* and pattern embedding *j*, and *τ* is a temperature hyperparameter (set here to 0.1). The loss encourages diagonal entries (matching pairs) to have high similarity while off-diagonal entries (non-matching pairs) have low similarity:

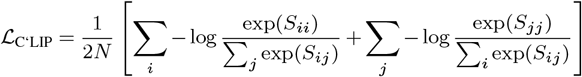

Models were trained using the Adam optimizer with a learning rate of 10^−4^ and default momentum parameters (*β*_1_ = 0.9, *β*_2_ = 0.999), and a batch size of 256. Gradient clipping with a maximum norm of 1.0 was used to stabilize training. Early stopping was triggered based on an exponential moving average of the validation loss, with smoothing parameter *α* = 0.1 and patience of 1,000 steps.

Isopleth was trained and tested in cross-validation on a dataset comprising the isotopologue-level labelling patterns and chemical structures of 231 metabolites that were identified by precursor m/z and retention time matches to authentic standards analyzed under identical LC–MS conditions. A binary labelling matrix was constructed for these metabolites using the statistical model described above, with entries set to 1 for tracer-isotopologue combinations passing a Bonferroni-corrected significance threshold of 0.01.

Given the small size of this ground-truth training dataset, we additionally experimented with two forms of data augmentation. First, we retrieved peaks that could readily be identified as mass spectrometric artifacts of these 231 metabolites (for instance, adducts or isotopologues), and which eluted within 0.5 min of the parent metabolite. We reasoned that, while these peaks should in principle exhibit the same labelling patterns as the parent metabolites themselves, measurement noise in isotopologue intensities might manifest in slightly different sets of labelling events passing thresholds for statistical significance, and we hypothesized that including these artifacts in the training set would encourage the model to learn more robust associations between labelling patterns and chemical structures. Each of these artifact peaks was labelled with the chemical structure of the parent metabolite and added to the training set.

Second, we leveraged putative structure annotations that were assigned by NetID to peaks that did not match a chemical standard. NetID annotates all experimentally observed peaks in an LC–MS experiment by constructing a network in which peaks are connected by edges representing mass differences that correspond to plausible biochemical transformations or MS artifacts (e.g., adducts, isotopes, or fragments)^15^. A subset of peaks in this network are annotated by comparison to a library of standards with known retention times and, optionally, MS/MS spectra. Candidate structures are then retrieved for the remaining peaks from a database of known metabolites (here, HMDB) and are scored via integer linear programming to ensure that connected peaks have chemically consistent annotations.

We found that both forms of data augmentation empirically improved the performance of Isopleth. The final training set therefore comprised isotopologue-level labelling patterns for 4,873 peaks and their associated chemical structures, including peaks identified directly through comparison to a library of chemical standards, mass spectrometry artifacts of those peaks, and peaks with putative structures assigned by NetID.

The performance of Isopleth was evaluated in ten-fold structure-disjoint cross-validation. A random subset comprising 10% of the training dataset was withheld as a validation set to monitor model convergence and trigger early stopping. At test time, candidate structures generated by DeepMet were retrieved for each peak in the test set, using a 5 ppm mass tolerance, and ranked by their cosine similarity to the observed labelling pattern in the embedding space learned by Isopleth. Performance was assessed by computing the top-*k* accuracy (the proportion of peaks for which the correct structure appears in the top *k* candidates) and the Tanimoto coefficient between the top-ranked candidate and the true structure. These metrics were calculated only for peaks identified via matches to chemical standards; artifacts and NetID annotations were used for data augmentation in the training set, but were excluded from the test set.

### Integrating analytical and isotope tracing data for structure annotation

Isopleth scores candidate structures based on their compatibility with the isotopic labelling pattern observed for a given peak. Conventionally, however, structural annotation of a mass spectrometric peak relies on other sources of analytical data, such as its MS/MS spectrum or retention time. We therefore sought to integrate the cosine similarities from Isopleth with orthogonal analytical data to rank candidate structures. We recently described a meta-learning approach in which a binary classifier is trained to distinguish true from false annotations using features derived from MS1-level mass errors, MS/MS spectra, and retention times^17^. This approach assigns each candidate structure a score that reflects its compatibility with the totality of the analytical data for a peak. Here, we extended this framework to additionally incorporate the cosine similarities from Isopleth.

We trained meta-learning classifiers to integrate MS1, MS/MS, and retention time data following our previously described approach. Briefly, we retrieved candidate structures generated by DeepMet for all 231 peaks with known identities, as described above. MS/MS spectra were predicted for the candidate structures using FraGNNet^6^, and retention times were predicted with a graph neural network^17^. Both the MS/MS and retention time prediction models were trained in cross-validation, simulating metabolite discovery by withholding known metabolites from the training sets of these models. Then, for each candidate structure, we computed a series of features, including (1) the frequency with which the candidate structure was generated; (2) the frequency normalized to the sum of frequencies of all candidate structures for that peak (i.e., the DeepMet confidence score); (3) the rank of the candidate structure, based on its sampling frequency, among all candidates generated by DeepMet; (4) the cosine similarity between the experimental and predicted MS/MS spectra for each candidate; (5) the number of fragment ions shared between the predicted and experimental spectra; (6) the mass error between the experimental and theoretical m/z, in parts per million; (7) the difference between experimental and predicted retention times; and (8) the cosine similarity between that structure and the isotopic labelling pattern for that peak, as computed by Isopleth. Missing feature values were imputed using the median of each feature computed across the training set, and features were standardized to zero mean and unit variance. A random forest classifier with 100 trees was then trained in tenfold cross-validation to predict whether a candidate structure was a correct or incorrect annotation for a given peak. To specifically assess the contribution of the isotope tracing information, we trained a second set of classifiers in cross-validation using only the first seven features (i.e., omitting the cosine similarities from Isopleth). The performance of the two meta-learning models was then assessed using the same metrics described above for the contrastive learning model: namely, the top-*k* accuracy and the Tanimoto coefficient between the top-ranked candidate and the ground-truth structure.

### Statistical analysis of metabolite abundance

We examined changes in abundance of the metabolites identified in this study across a range of physiological and pathophysiological contexts. The statistical analyses were performed using the limma R package. Peak intensities were log-transformed and no imputation of missing values was performed. Models were fit using ordinary least squares via lmFit and variance estimates were moderated via eBayes. Multiple hypothesis correction was applied using the Benjamini-Hochberg (BH) procedure over all 8,277 peaks.

Analyses of tissue specificity, sexual dimorphism, and response to purified diet were performed in the same mouse tissue dataset used to define the set of 8,277 peaks considered throughout the study. This dataset comprised metabolomic measurements from 25 mouse tissues in a total of 15 mice, including five male and five female mice fed standard chow diets and an additional five male mice fed purified diets. Tissue-enriched metabolites were identified by fitting a linear model to data from chow-fed mice, comparing samples from a given tissue to samples from all remaining tissues, following the approach of Melé et al.^47^ Specifically, we fit linear models with covariates for tissue (the focal tissue versus all others), repeating this analysis for each tissue in turn, with BH correction applied across all 25 tissue contrasts. Sexually dimorphic metabolites were identified by fitting a linear model to the same dataset that included indicator variables for each tissue as well as sex. Metabolites that responded to dietary perturbation were identified by extending this model with an additional coefficient for diet (purified relative to chow), and fitting the model to data from all 15 mice.

The responsiveness of metabolites to perturbation of the microbiome was evaluated by re-analyzing data from Roichman et al.^23^ This dataset comprised metabolomic measurements from the gastrointestinal tract (cecal contents, colonic contents, and feces) and serum (portal and tail vein, sampled in both fasted and fed states) of mice treated with broad-spectrum antibiotics and untreated controls. Intensities of the 8,277 peaks defined in the mouse tissue dataset were extracted using a retention time window of ± 1 min and an m/z tolerance of 3 ppm; only peaks with a mean intensity greater than 10^4^ were retained to exclude metabolites not reliably detected in these samples. Linear models were fit to log-transformed intensities separately within each combination of tissue and fasting state, with a single coefficient comparing antibiotic-treated to control mice.

Age-responsive metabolites were identified by re-analyzing the same non-isotope aging mouse samples described in the “Aging mouse studies” methods section (see also Jankowski et al.^33^). Mice were divided according to age into young (4 months) and old (24 months) groups, and a linear model was fit to the data with a coefficient for age. As in the analysis of the antibiotics dataset, only metabolites with a mean intensity greater than 10^4^ were included.

To identify metabolites altered in tumor tissue relative to matched normal adjacent tissue (NAT), we exploited the paired structure of the dataset by analyzing within-patient differences. Specifically, for each cancer type, a within-patient difference matrix was computed by subtracting intensities from NAT samples from the corresponding tumor samples to produce a matrix of within-patient differences in metabolite abundance, and a linear model with an intercept term only was fit to this matrix, again only for metabolites with a mean intensity greater than 10^4^.

### Reference MS/MS search against metabolomic repositories

To determine whether the metabolites described here had been detected in published human metabolomics experiments, we curated a large compendium of untargeted LC–MS data from the MetaboLights, Metabolomics Workbench, and GNPS repositories^48–50^.

We first identified human metabolomics experiments in each of these three repositories. An XML record of all studies deposited to MetaboLights was obtained (file ‘eb-eye_metabolights_studies.xml’) and filtered to only mass spectrometry-based metabolomics studies that included at least one sample from a human tissue or biofluid. Complete data depositions for this subset of studies were then downloaded from MetaboLights. The assay-level metadata (‘a_*’ files) were parsed to obtain a complete list of all mass spectrometric runs for all of the human metabolome studies and to exclude GC–MS, imaging MS, and targeted MS experiments, inspecting the relevant MTBLS pages and the corresponding publications as necessary to ensure that no LC–MS metabolomics studies were inadvertently removed and to correct manually any filenames that were discordant between the assay-level metadata and the deposited raw files. For Metabolomics Workbench, the REST API was queried first to retrieve all LC–MS studies (endpoint ‘/rest/study/study_id/ST/summary’) and then manually curated in order to subset these to experiments in human tissues and remove blanks. For GNPS, a record of all studies (file filtered_results.tsv) was retrieved from MassIVE and subset to publicly available human metabolomics studies. Each accession was then manually curated to discard runs not from human tissues or biofluids.

Complete data depositions for these subsets of studies were downloaded from the respective repositories. Compressed archives (.tar, .gz, .zip) were decompressed, and vendor-specific formats (.d, .raw, .wiff) were converted to mzML using the msconvert utility bundled with ProteoWizard^51^. MS/MS spectra were extracted from each mzML file and written to MGF format. MS/MS spectra from each run were then extracted and written to MGF files after ensuring the following quality control (QC) criteria were met: at least 50 unique precursor m/z values; at least 100 non-empty MS/MS spectra; both precursor m/z and fragment m/z recorded to at least four decimal places; and precursor m/z range > 200 Th. LC–MS/MS files that did not meet one or more of these filters, and the publications accompanying these depositions, were manually reviewed to determine why they did not pass these QC criteria. A handful of duplicate files, representing cases where the same mass spectrometry run was uploaded as part of more than one accession, were identified by their checksums and removed.

In total, these steps afforded a resource comprising 999 million MS/MS spectra from 152,743 LC–MS/MS experiments, spanning 512 distinct studies (i.e., accessions). Of these spectra, 85% were acquired in positive ionization mode.

Sample-level metadata was then curated for each file. For MetaboLights, metadata was obtained by parsing sample-level metadata (‘s_*’ files). For Metabolomics-Workbench, sample level-metadata was retrieved using the REST API (‘rest/study/study_id/ST00XXXX/factors’). For GNPS, the MassIVE page and (when available) accompanying publication for each individual dataset was manually inspected to extract associated sample information. Each of the collected LC–MS/MS runs from the three repositories was then further annotated by cross-referencing accessions and filenames to retrieve metadata annotated by Pan-ReDU^52^.

To maximize the likelihood of obtaining high-quality MS/MS matches to published datasets, MS/MS spectra were reacquired from the synthetic reference standards of confirmed metabolites in both ionization modes and at multiple collision energies (10, 20, 30, and 40 eV). These MS/MS spectra were then searched against the published human metabolomics data using a precursor m/z tolerance of 10 ppm, retaining matches with a cosine similarity ≥ 0.75 and at least three matching peaks above 1% of the base peak intensity.

### Terminology

Throughout the manuscript, we use the terminology of “previously unrecognized metabolite” to refer to a small molecule that, to the best of our knowledge, had not previously been recognized as a mammalian metabolite. We do so recognizing that it is challenging to assert a negative (i.e., that a molecule is unknown in the context of mammalian metabolism). We employed a multi-tiered process involving extensive manual review to support this categorization for all metabolites reported in the manuscript, in which three authors independently reviewed the literature and came to separate determinations regarding the status of each metabolite. First, we discarded any structures present in any version of the Human Metabolome Database, with any annotation status (quantified, detected, expected, predicted). Second, if the structure was present in PubChem or CAS SciFinder, we manually reviewed all of the associated literature references to establish whether any of these reported its detection in mammals. Third, we formulated potential common names or synonyms that we could envision describing the compound in question, and performed literature searches using Google Scholar and PubMed. Fourth, we searched for potential isomers on PubChem and SciFinder that could be envisioned to afford similar MS/MS spectra to identify whether these were known to be mammalian metabolites. This multi-tiered review procedure led us to discard a number of metabolites from the manuscript that, although apparently not broadly recognized as mammalian metabolites, had been described as such in at least one prior study. In **Supplementary Note 2**, we provide additional context for each of the previously unrecognized metabolites, including reports of their detection in non-mammalian species. Despite our best efforts, this review of the literature may have been incomplete for certain compounds.

### Visualization

Throughout the paper, box plots show the median (horizontal line), interquartile range (hinges) and smallest and largest values no more than 1.5 times the interquartile range (whiskers), and EICs were smoothed using a Gaussian kernel regression (bandwidth = 0.3).

## Acknowledgements

This work was supported by Ludwig Cancer Research; the Weill Cancer Hub East; the National Institutes of Health (DP5OD036960 to M.A.S., R50CA211437 to W.L., P30DK019525 to J.D.R., P30CA072720 to J.D.R.); the Searle Scholars Program (to M.A.S.); and the Hevolution Foundation (HF-GRO-23-1199192-1 to J.D.R. and M.R.M). The content is solely the responsibility of the authors and does not necessarily represent the official views of the National Institutes of Health. J.R. was supported by a Postdoc.Mobility fellowship from the Swiss National Science Foundation (SNSF, 235522). Services, results, and products in support of the research project were generated by the Rutgers Cancer Institute, Biospecimen Repository and Histopathology Service Shared Resource, supported, in part, with funding from the NCI-CCSG P30CA072770-6852. We thank Marea Therapeutics for providing human oral glucose tolerance test samples. We also thank Dr. Min Huang (Fox Chase Cancer Center) for assistance in acquiring samples from the Fox Chase Cancer Center Biosample Repository Facility.

## Data availability

Raw mass spectrometry data for isotope tracing experiments have been deposited to MassIVE.

## Code availability

Source code for Isopleth is available from GitHub at https://github.com/skinniderlab/Isopleth.

---

**Supplementary Fig. 1.**
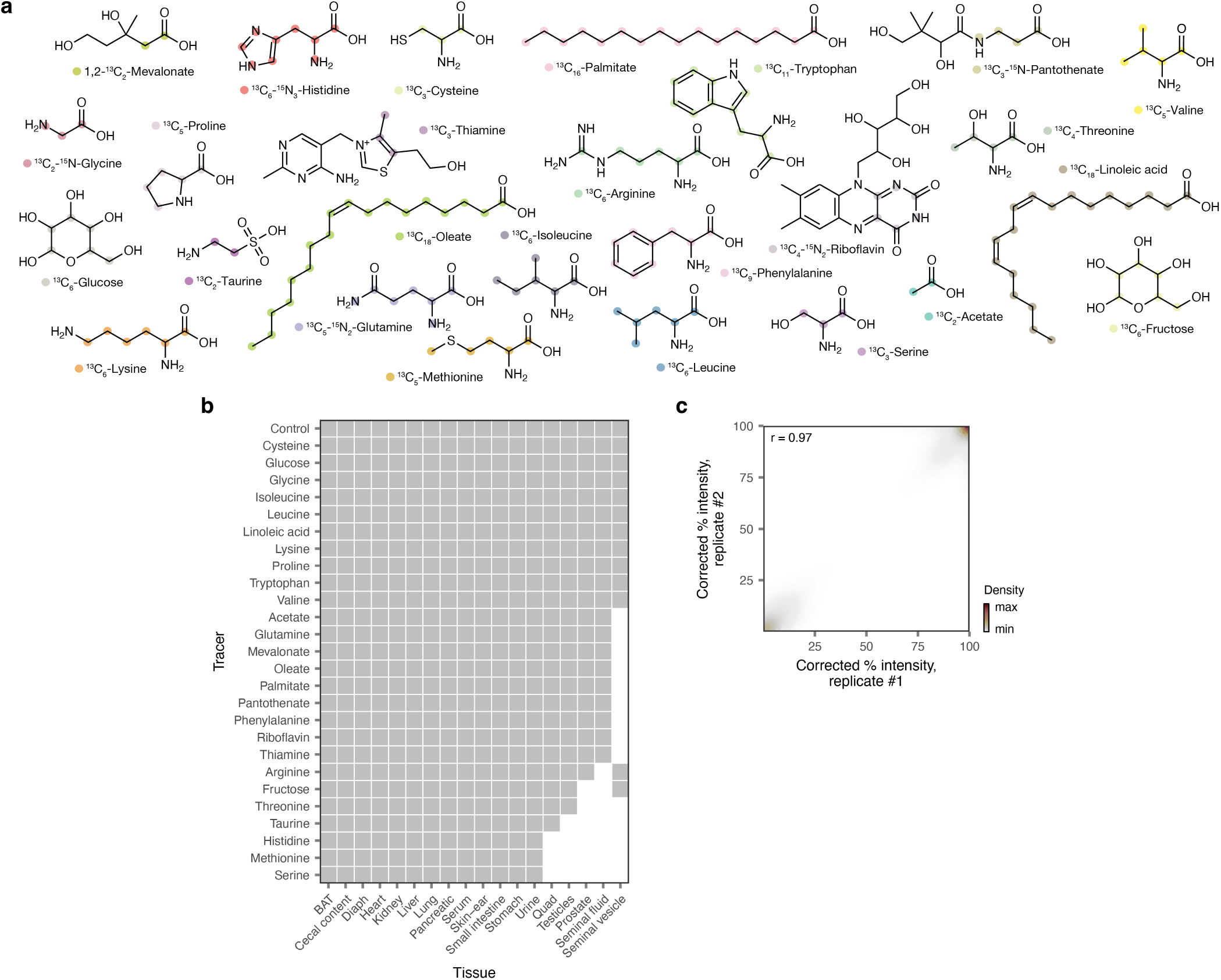
Tracing the fates of 26 isotopically labelled nutrients in mouse tissues. **a**, Chemical structures of nutrients administered to mice, with isotopically labelled atoms highlighted. **b**, Summary of mouse tissues profiled for each nutrient. **c**, Reproducibility of *in vivo* isotopic labelling experiments. Scatterplot shows the correlation in fractional labelling after natural isotope abundance correction for each of 113,039 isotopologues of 8,277 peaks between biological replicates; points are colored by local density to improve visualization of overlapping data points (omitting peaks without any detectable labelling, which comprise the majority of the dataset). Inset text shows the Pearson correlation coefficient.

**Supplementary Fig. 2.**
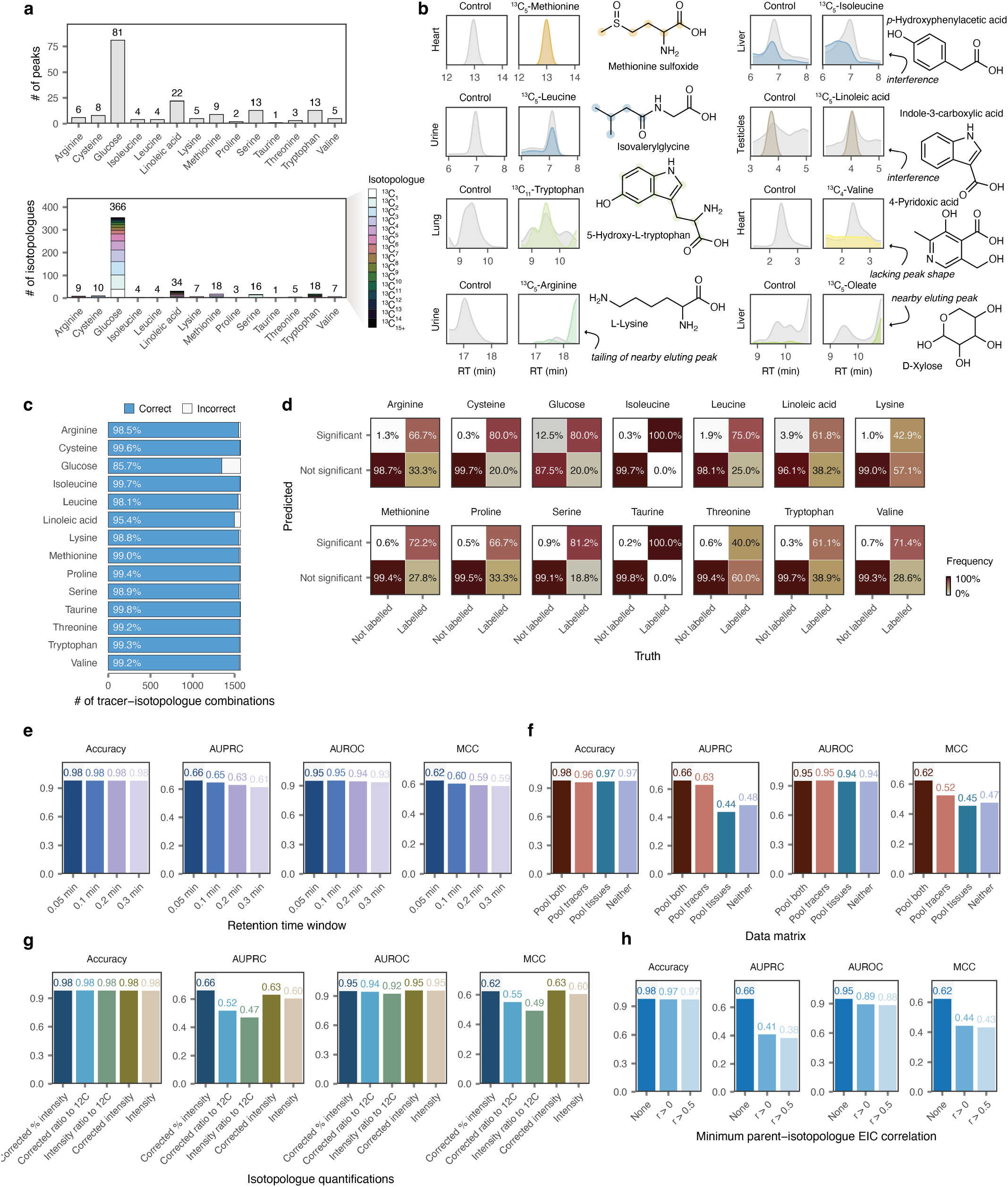
Statistical analysis of isotopologue-labelling patterns. **a**, Overview of the manually labelled ground-truth dataset against which the performance of the statistical model was evaluated. Ground-truth labelling patterns for a subset of nutrients (*n* = 14) were defined through manual inspection of their extracted ion chromatograms (EICs) by an experienced analyst, prior to the development of the statistical model itself. Top, number of peaks corresponding to known metabolites that were determined to label from each nutrient. Bottom, number of isotopologues from each of these peaks that were determined to label from each nutrient. **b**, Examples of manually annotated EICs from the ground-truth dataset showing credible or spurious labelling. **c**, Accuracy of the statistical model for determining isotopologue-level labelling patterns, as in Fig. 1c but here shown separately for each tracer. **d**, Confusion matrices, as in Fig. 1d but here shown separately for each tracer. **e-h**, Impact of key aspects of data preprocessing or model design. The accuracy, area under the receiver operating characteristic curve (AUROC), area under the precision-recall curve (AUPRC), and Matthews correlation coefficient (MCC) are shown for each. **e**, Effect of the retention time window used to extract isotopologue intensities. Narrower windows afford improved performance. **f**, Effect of the data matrix used for model fitting. Analyzing data from all tracers and tissues jointly yields the optimal performance. **g**, Effect of the isotopologue quantifications used for model fitting. Analyzing fractional labelling after natural isotope abundance correction outperforms other approaches to isotopologue intensity quantification. **h**, Effect of filtering isotopologue quantifications based on the Pearson correlation between parent and isotopologue extracted ion chromatograms (EICs). Filtering does not improve performance.

**Supplementary Fig. 3.**
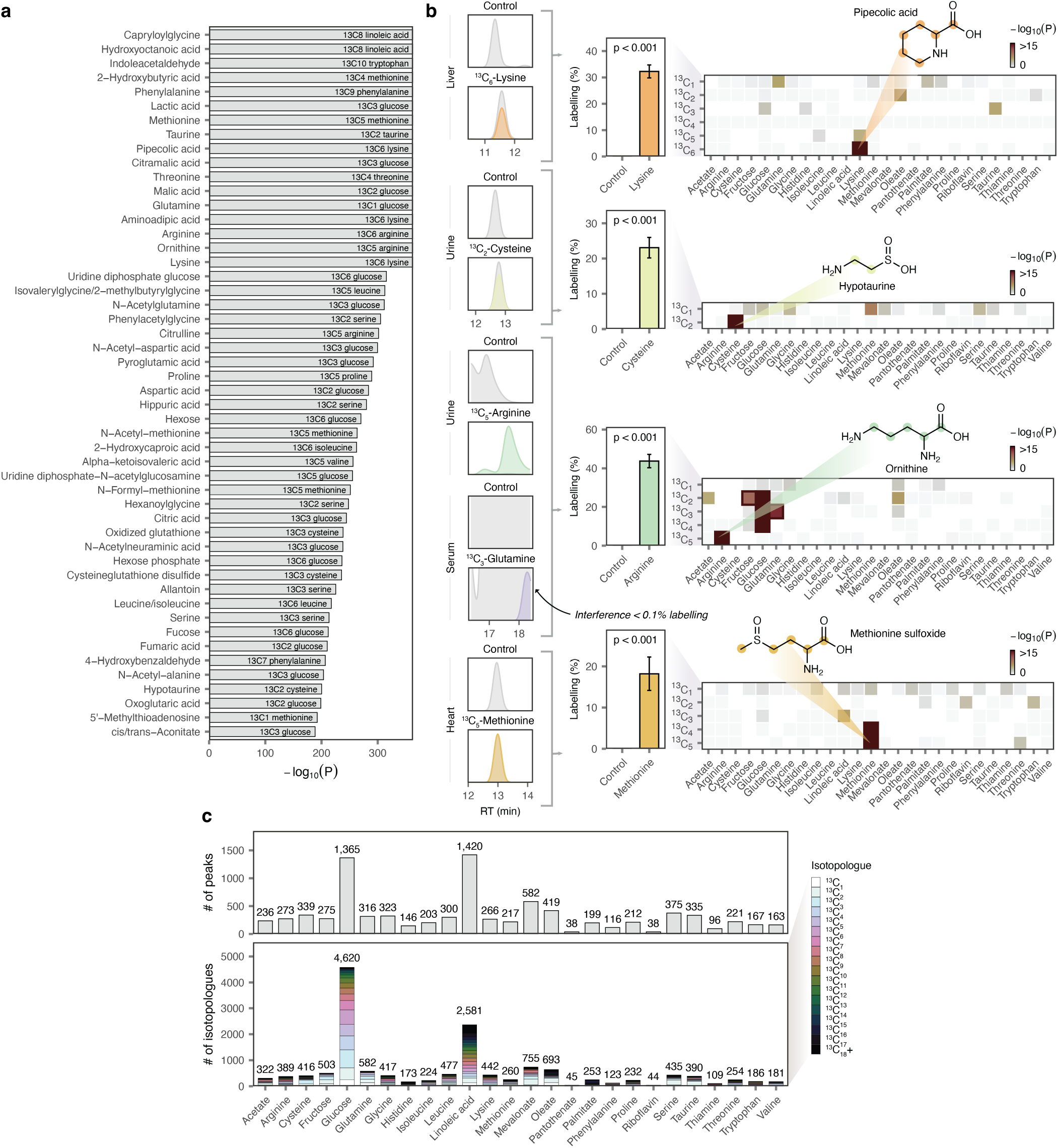
Precursor incorporation into known and putative metabolites. **a**, Examples of highly significant labelling events for peaks corresponding to known metabolites. Bars show −log_10_(P) values for individual labelling events; text indicates the number of carbon atoms incorporated from the relevant tracer. **b**, Additional examples of isotopologue-level labelling pattern inference for peaks corresponding to known metabolites, including examples of both true and false positives. **c**, Overview of precursor incorporation across 4,506 putative metabolites. Top, number of peaks inferred to incorporate at least one carbon atom from each precursor. Bottom, number of statistically significant labelling events (tracer-isotopologue combinations) from each precursor.

**Supplementary Fig. 4.**
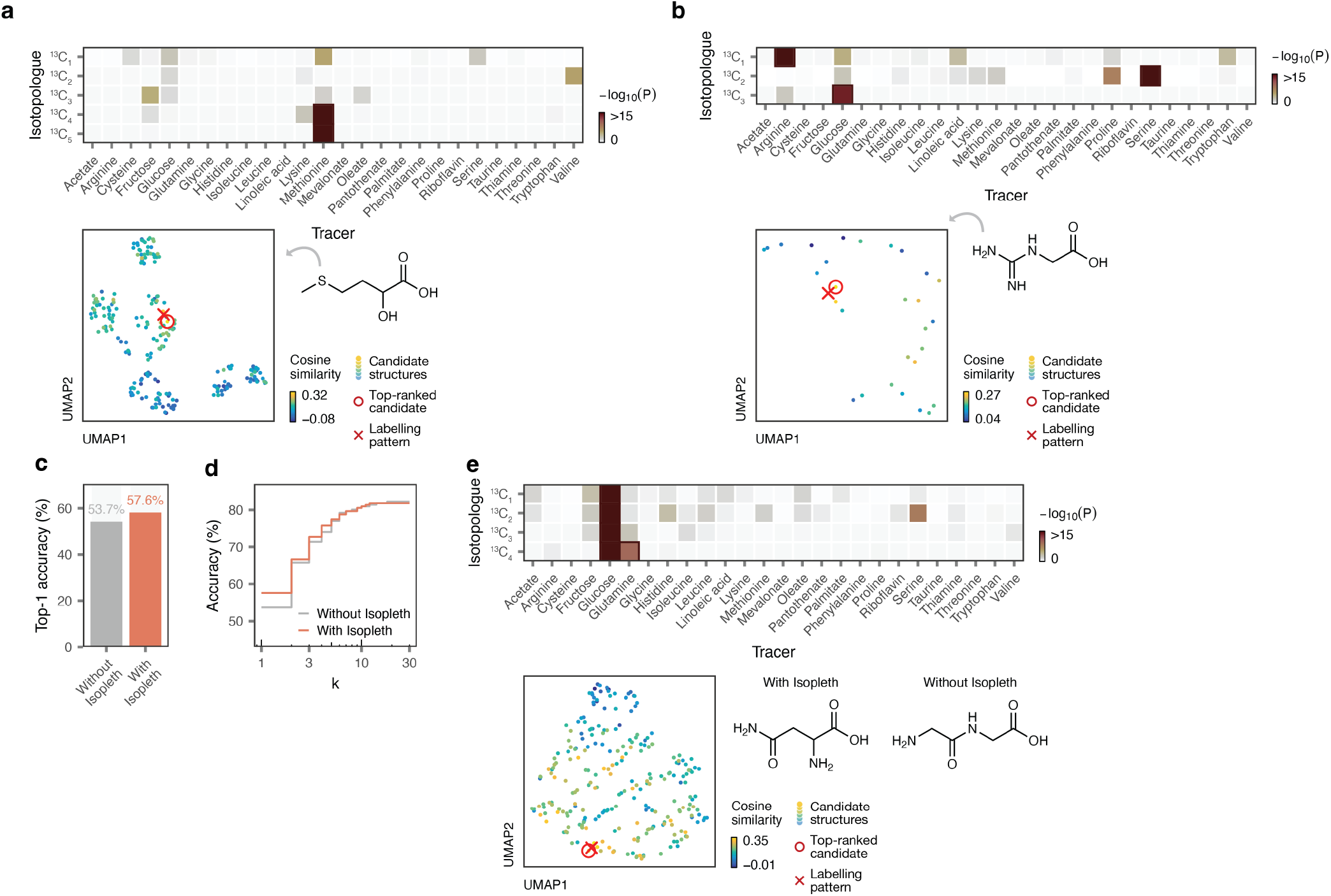
Biosynthesis-aware structure annotation of simulated unknown metabolites. **a-b**, Additional examples of held-out metabolites correctly prioritized by Isopleth. Top, isotopologue-level labelling patterns (heatmap shows Bonferroni-corrected −log_10_ p-values for each tracer-isotopologue combination); bottom left, UMAP visualizations of Isopleth embeddings for candidate structures, colored by their cosine similarity to the observed labelling pattern for each peak; bottom right, structures of the correctly prioritized metabolites. **a**, 2-hydroxy-4-(methylthio)butanoic acid; **b**, guanidinoacetic acid. **c**, Top-1 accuracy of structure annotations from meta-learning models, with or without the inclusion of cosine similarities from Isopleth as a feature in the model (p = 0.016, two-sided McNemar’s test). **d**, As in **c**, but showing the top-*k* accuracy (for *k* ≤ 30). **e**, Example of a held-out metabolite correctly prioritized by the meta-learning model with the inclusion of cosine similarities from Isopleth, but not without. Top, isotopologue-level labelling pattern; bottom left, UMAP visualizations of Isopleth embeddings for candidate structures, colored by their cosine similarity to the observed labelling pattern; bottom right, structure of the correctly prioritized metabolite (asparagine) and an isomer incorrectly prioritized by the meta-learning model without the inclusion of cosine similarities as a feature (glycylglycine).

**Supplementary Fig. 5.**
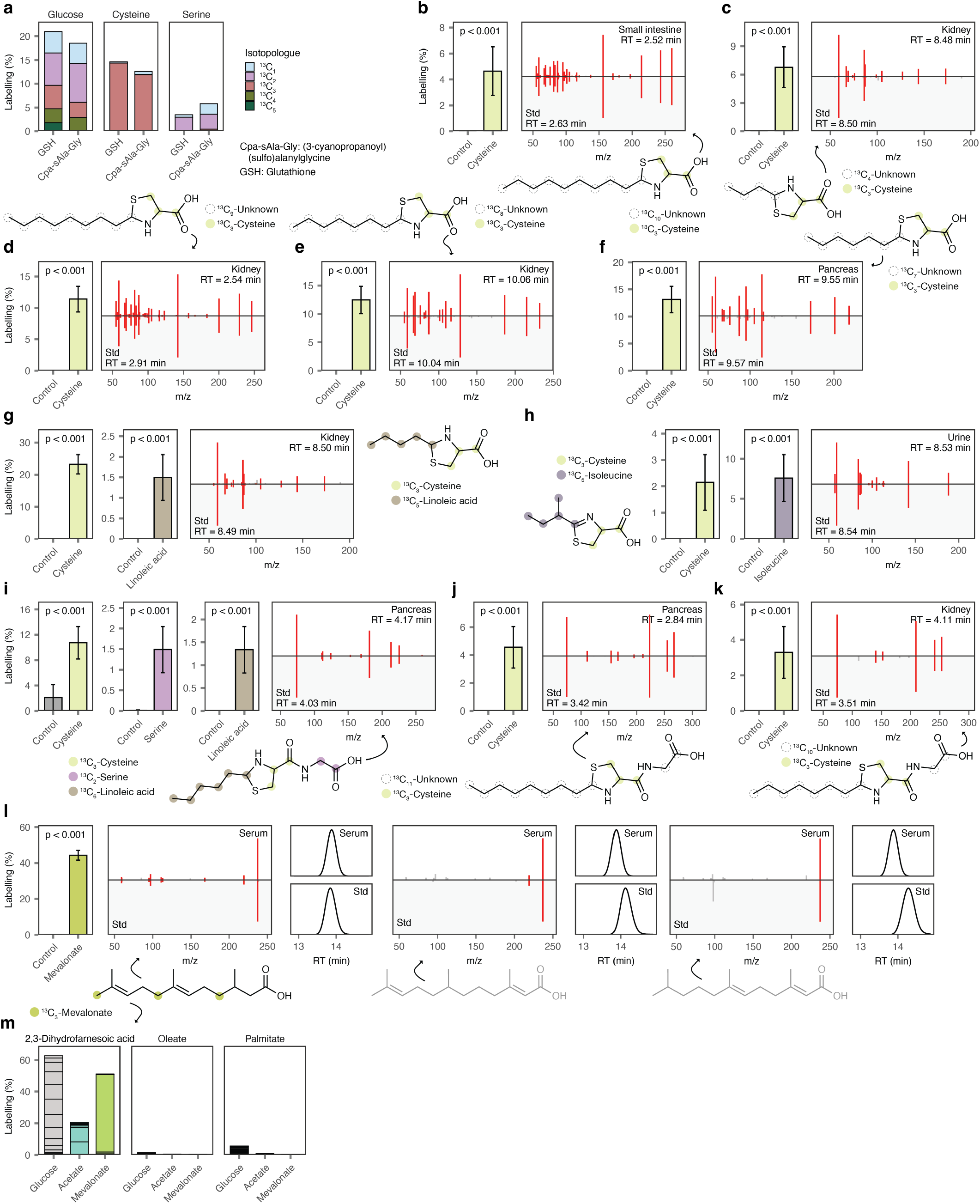
(3-cyanopropanoyl)(sulfo)alanylglycine, cysteine-derived alkylthiazolidines and 2,3-dihydrofarnesoic acid. **a**, Labelling of (3-cyanopropanoyl)(sulfo)alanylglycine and reduced glutathione from glucose, cysteine, and serine. Stacked bar charts show fractional labelling after natural isotope abundance correction for relevant isotopologues. **b-k**, Isotope tracing and MS/MS spectra for additional metabolites whose discovery was enabled by systematic isotope tracing data. Bar charts show fractional labelling after natural isotope abundance correction for selected labelling events; inset text shows p-values after Bonferroni correction. Mirror plots show the similarity between MS/MS spectra from synthetic reference standards versus experimental peaks in mouse tissues. **b**, 2-nonylthiazolidine-4-carboxylic acid. **c**, 2-propylthiazolidine-4-carboxylic acid. **d**, 2-octylthiazolidine-4-carboxylic acid. **e**, 2-heptylthiazolidine-4-carboxylic acid. **f**, 2-hexylthiazolidine-4-carboxylic acid. **g**, 2-butylthiazolidine-4-carboxylic acid. **h**, 2-(*sec*-butyl)-4,5-dihydrothiazole-4-carboxylic acid. **i**, (2-pentylthiazolidine-4-carbonyl)glycine. **j**, (2-octylthiazolidine-4-carbonyl)glycine. **k**, (2-heptylthiazolidine-4-carbonyl)glycine. **l**, Left, 2,3-dihydrofarnesoic acid; center and right, two positional isomers that were discredited based on retention time and MS/MS. Fragment ion intensities are shown after square-root transformation to accentuate low-intensity fragments. Retention times are shown here on reverse phase chromatography (versus HILIC in **Fig. 3e**). **m**, Labelling of 2,3-dihydrofarnesoic acid, palmitate, and oleate from glucose, acetate, and mevalonate. Stacked bar charts show fractional labelling after natural isotope abundance correction for each potential isotopologue from ^13^C_1_ to U^13^C.

**Supplementary Fig. 6.**
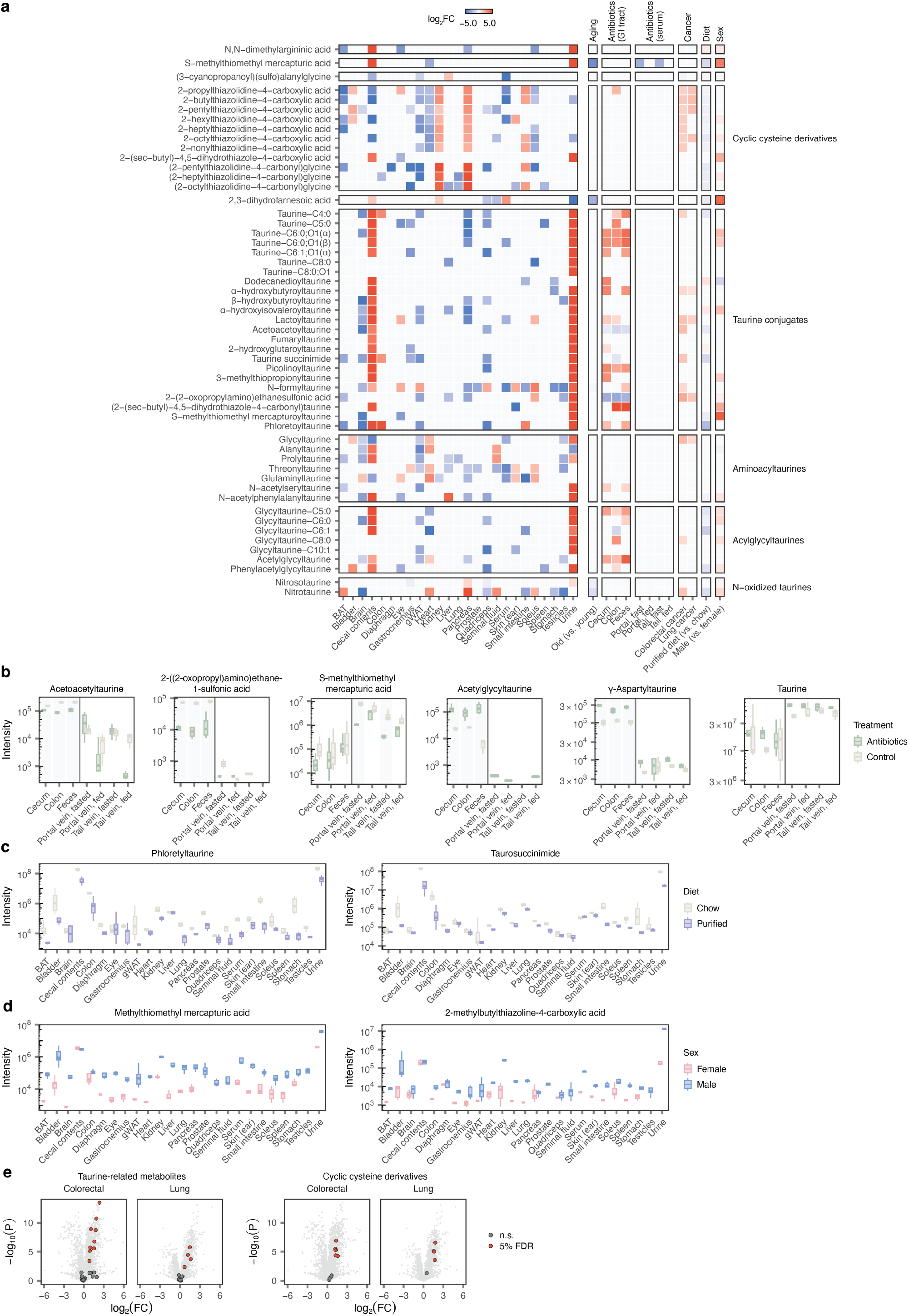
Metabolite origins and phenotypic associations. **a**, Tissue specificity and perturbation responsiveness of metabolites identified in this study. Left, tissue specificity. Log_2_ fold changes are shown for peaks significantly elevated or decreased in individual tissues at 5% FDR. Right, perturbation responsiveness. From left, columns show log_2_ fold changes for: aged versus young mice; antibiotic treatment versus untreated mice; cancer versus normal adjacent tissue (NAT); purified diet versus standard chow; male versus female mice. Log_2_ fold changes are shown for peaks with significant associations; non-significant changes are indicated in grey. Log_2_ fold changes were winsorized to a minimum and maximum of –5 and 5, respectively, for the purpose of visualization. **b**, Examples of peaks that are significantly elevated or decreased in mice treated with broad-spectrum antibiotics as compared to untreated controls. Taurine is shown for comparison with other taurine-derived metabolites. **c**, Examples of peaks that are significantly reduced in mice fed purified diets as compared to standard chow. **d**, Examples of peaks that are significantly elevated in male mice relative to female mice. **e**, Volcano plots comparing relative abundance of metabolite levels between colorectal adenocarcinoma, left, or lung adenocarcinoma, right, and adjacent normal tissue. Taurine-related metabolites or cyclic cysteine-derived metabolites, respectively, are highlighted. Metabolites significant at 5% FDR are shown in red.

**Supplementary Fig. 7.**
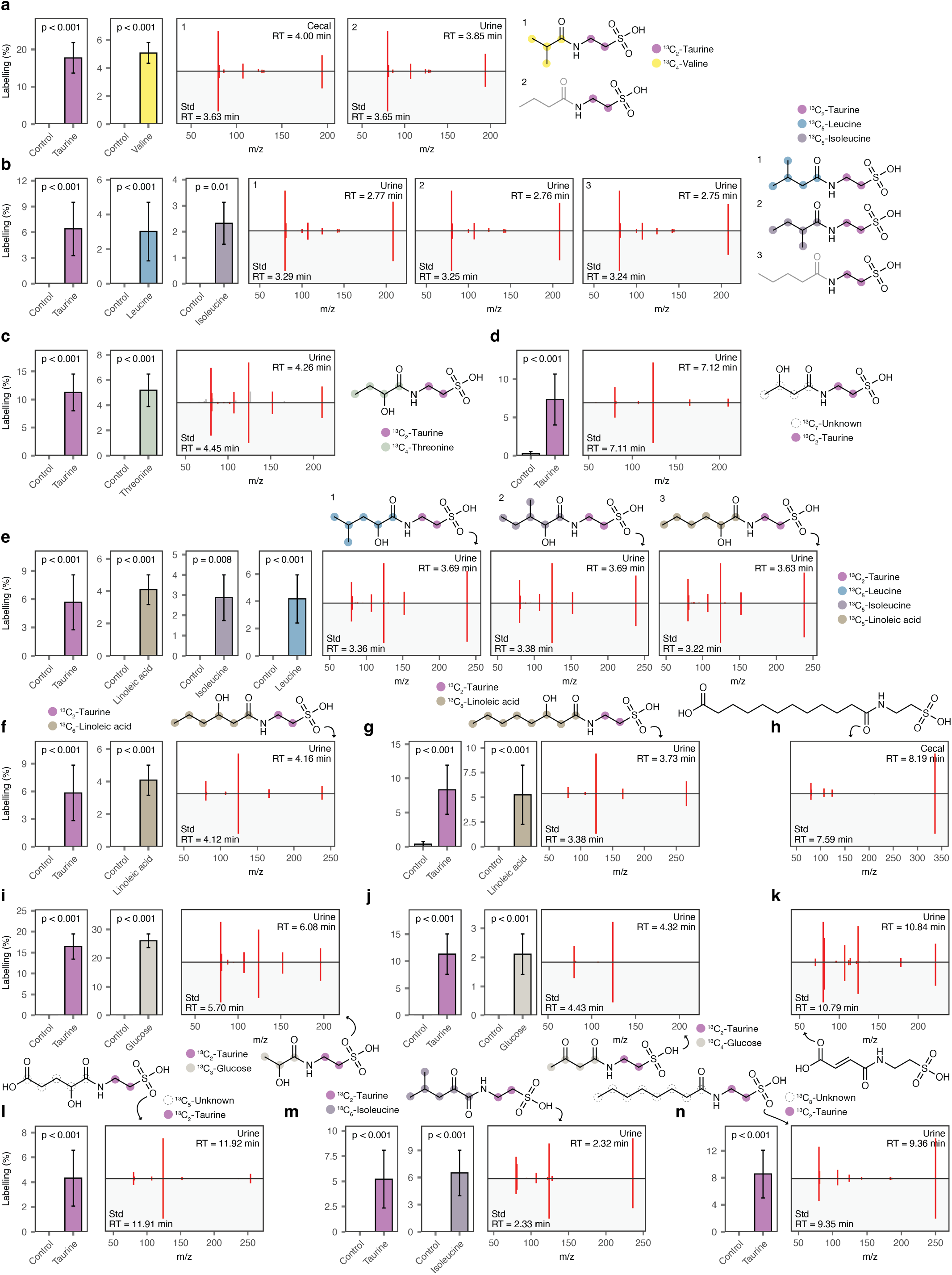
N-acyltaurines. **a-n**, Isotope tracing and MS/MS spectra for additional N-acyltaurines whose discovery was enabled by isotope tracing. Bar charts show fractional labelling after natural isotope abundance correction; inset text shows p-values after Bonferroni correction. Mirror plots show the similarity between MS/MS spectra from synthetic reference standards versus experimental peaks in mouse tissues. **a**, Taurine-C4:0. The peak was inferred to incorporate four carbons from valine, suggesting isobutyryltaurine (**1**); however, retention time and MS/MS data did not rule out the presence of the linear isomer butyryltaurine (**2**). **b**, Taurine-C5:0. The peak was inferred to incorporate five carbons from leucine or isoleucine (species **1–2**); however, retention time and MS/MS data did not rule out the presence of the linear isomer valeryltaurine (**3**). **c**, *α*-Hydroxybutyroyltaurine. **d**, *β*-hydroxybutyroyltaurine. **e**, Taurine-C6:0;O1(*α*). The peak was inferred to incorporate six carbons from isoleucine, leucine, or linoleic acid, suggesting a mixture of species **1–3** in mouse tissues. **f**, Taurine-C6:0;O1(*β*). **g**, Taurine-C8:0;O1. **h**, Dodecanedioyltaurine, which did not match to the originally hypothesized peak but matched another peak in the mouse tissue dataset. **i**, Lactoyltaurine. **j**, Acetoacetoyltaurine. **k**, Fumaryltaurine, which was not among the 8,277 peaks detected in mouse tissue extracts but was independently confirmed as a mouse metabolite. **l**, 2-hydroxyglutaroyltaurine. **m**, Taurine-C6:1;O1(*α*). **n**, Taurine-C8:0.

**Supplementary Fig. 8.**
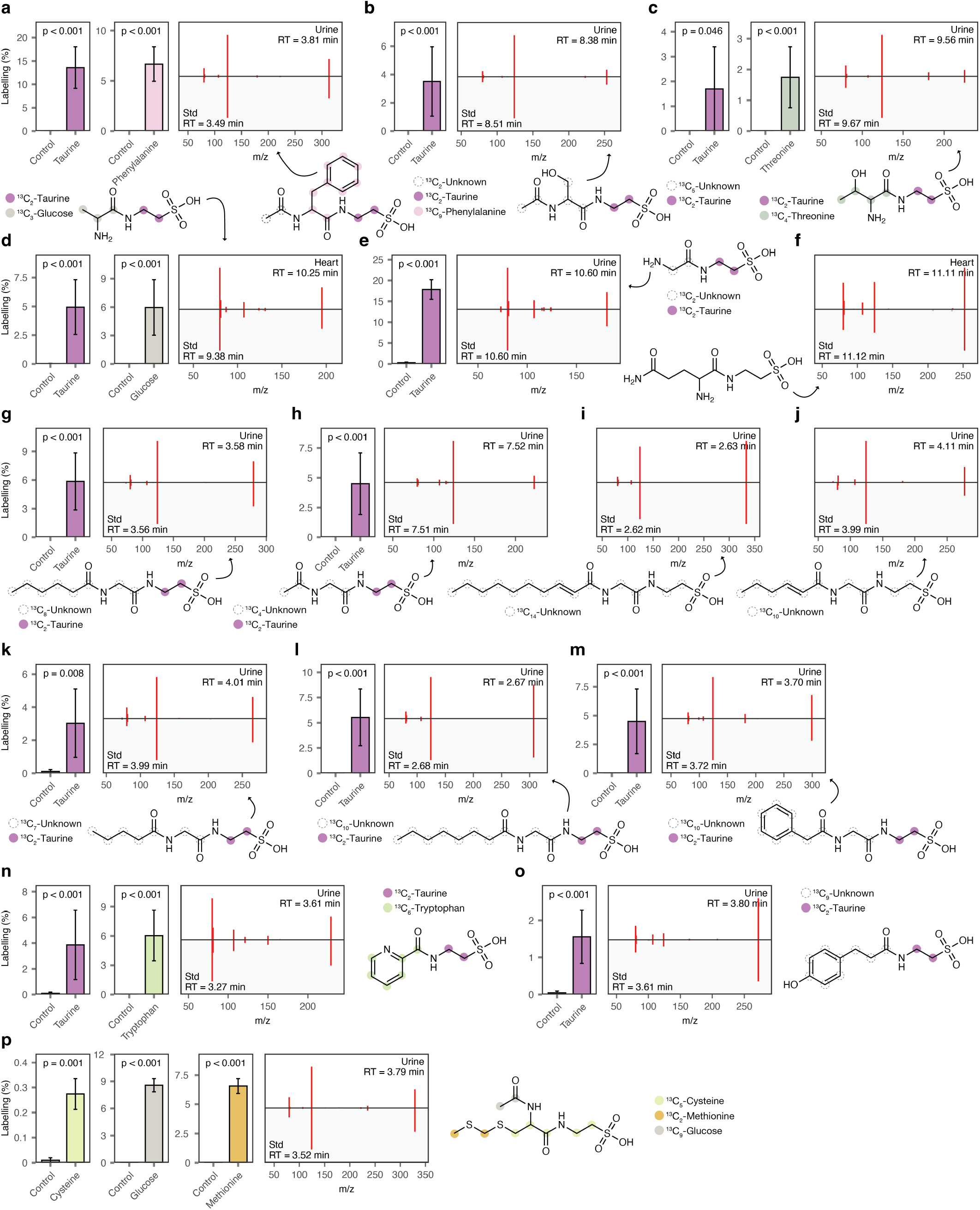
Aminoacyltaurines, acylglycyltaurines, and other taurine conjugates. **a**, N-acetylphenylalanyltaurine. **b**, N-acetylseryltaurine. **c**, Threonyltaurine. **d**, Alanyltaurine. **e**, Glycyltaurine. **f**, Glutaminyltaurine, which was not among the 8,277 peaks detected in mouse tissue extracts but was independently confirmed as a mouse metabolite. **g**, Glycyltaurine-C6:0. **h**, Acetylglycyltaurine. **i**, Glycyltaurine-C10:1. **j**, Glycyltaurine-C6:1. **k**, Glycyltaurine-C5:0. **l**, Glycyltaurine-C8:0. **m**, Phenylacetylglycyltaurine. **n**, Picolinoyltaurine. **o**, Phloretoyltaurine. **p**, S-methylthiomethyl mercapturoyltaurine.

**Supplementary Fig. 9.**
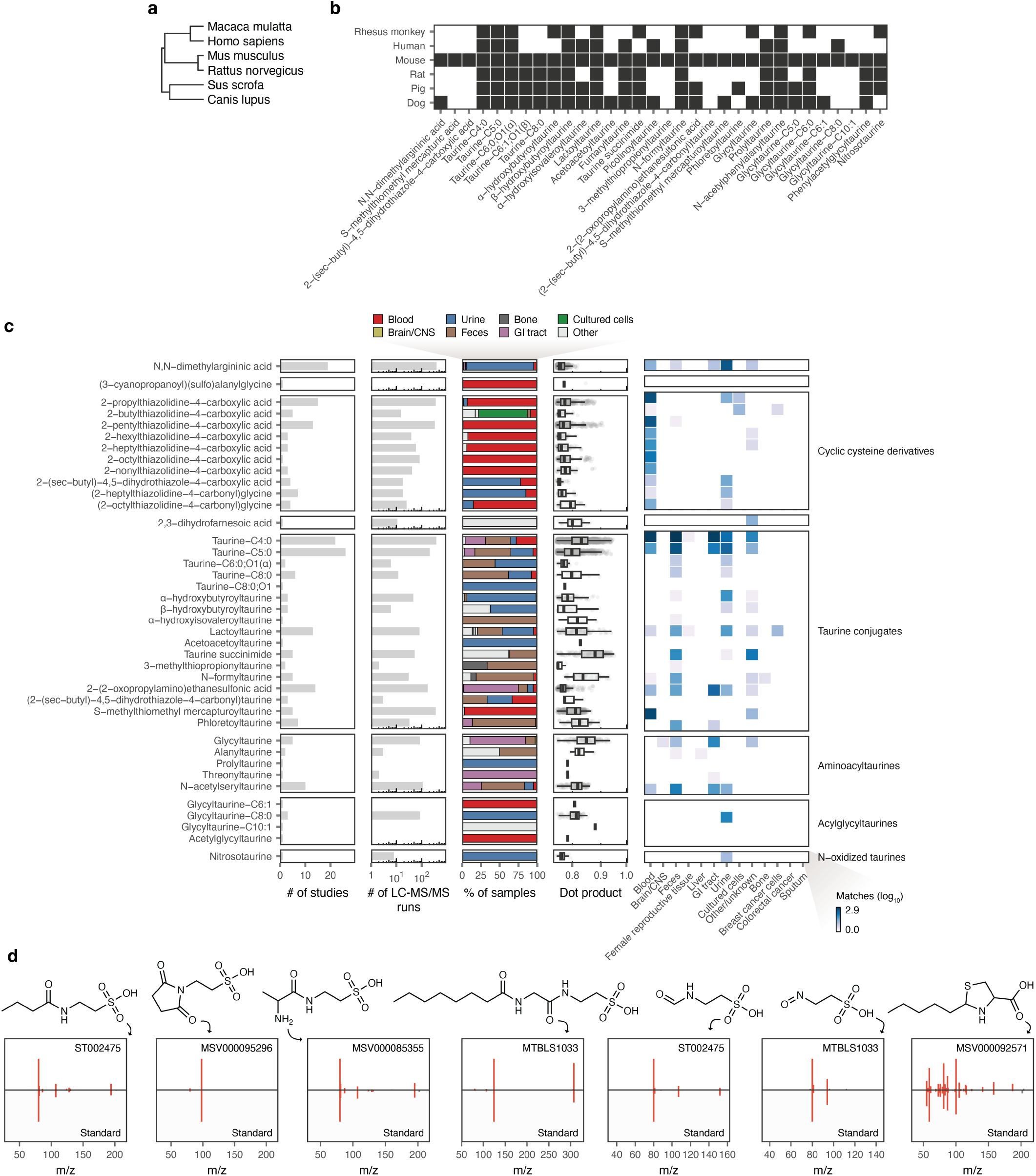
Metabolite distribution and evolutionary conservation. **a**, Phylogenetic tree of species analyzed in this study. **b**, Heatmap showing the presence or absence of metabolites that were detected in mouse urine across urine samples from five other mammals. **c**, Overview of MS/MS matches between reference spectra for metabolites identified in this study and published human metabolomics data. From the left, columns show: (1) number of studies (accessions) in which each metabolite was tentatively identified (minimum dot-product of 0.75, minimum 3 matching peaks, maximum 10 ppm precursor m/z error); (2) number of LC–MS/MS runs in which a metabolite was tentatively identified; (3) distribution of tissues and sample types in which each metabolite was tentatively identified; (4) distribution of dot-products between reference and experimental MS/MS spectra for each metabolite; (5) number of spectral matches (log_10_ scale) by tissue or sample type for each metabolite. **d**, Mirror plots showing representative matches between reference spectra for metabolites identified in this study and experimental spectra from published human metabolomics data.

**Supplementary Fig. 10.**
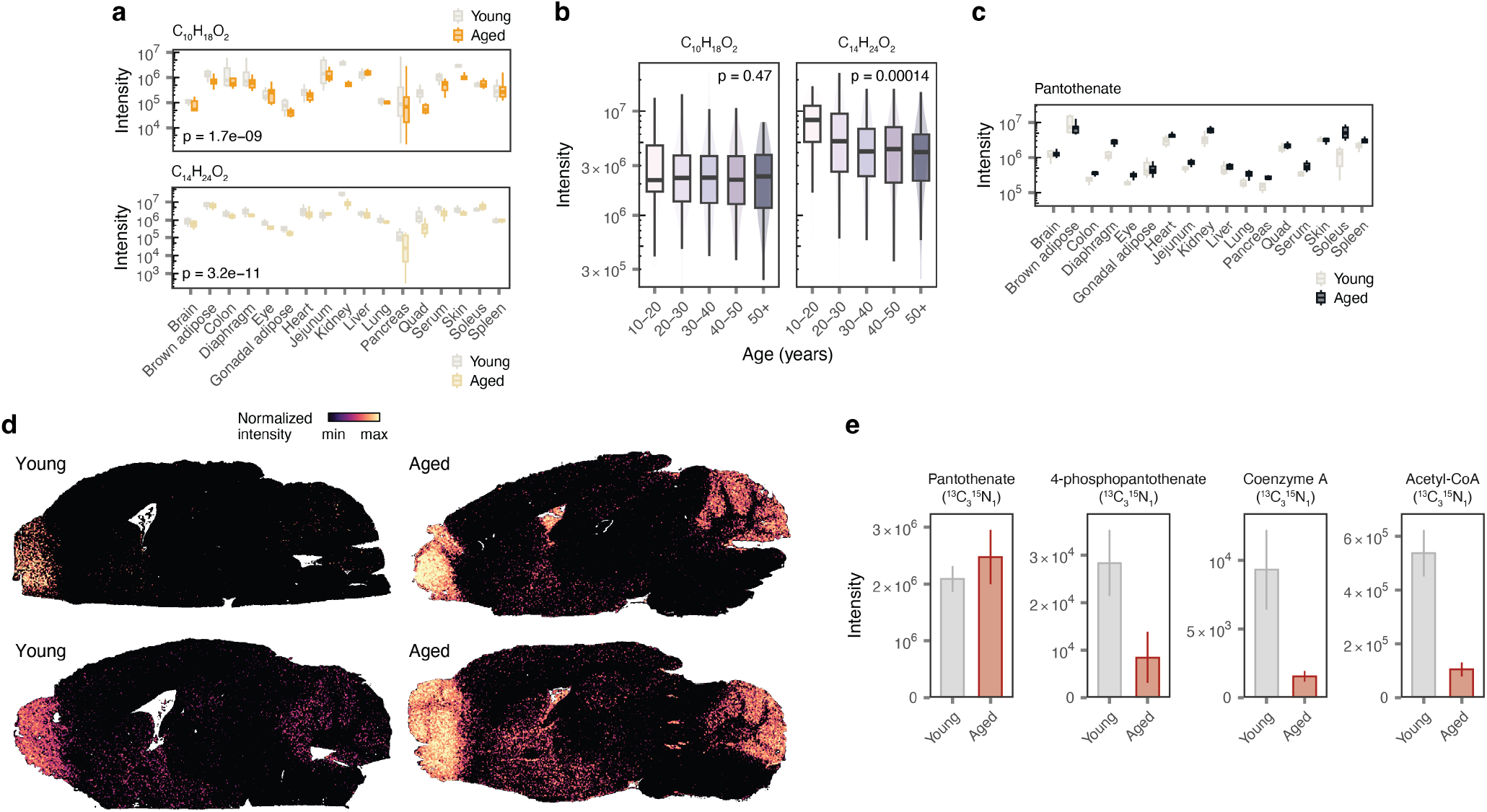
Supporting data for mevalonate-derived metabolites. **a**, Tissue levels of mevalonate-derived species C_10_H_18_O_2_ (top) and C_14_H_24_O_2_ (bottom) across 17 tissues in young and aged C57BL/6J mice (data are from 17 tissues with 9–15 independent mice per solid tissue and 30 for serum). Inset text shows p-values from the moderated t-test implemented in limma. **b**, Serum levels of C_10_H_18_O_2_ and C_14_H_24_O_2_ across age from a cross-sectional human cohort study. Inset text shows the p-value from a two-sided test of Pearson’s correlation coefficient (*n* = 395). **c**, As in **a**, but showing pantothenate. **d**, Images from additional biological replicates showing spatial metabolomic measurements of pantothenate levels in young and aged C57BL/6J mouse brains. **e**, Levels of ^13^C_3_-^15^N_1_-labelled pantothenate, 4-phospho-pantothenate, coenzyme-A (CoA) and acetyl-CoA in liver samples of young and aged C57BL/6J mice following infusion of ^13^C_3_-^15^N_1_-pantothenate. Bars and error bars show mean and standard error respectively (*n* = 4–10).

