## Supplementary Note 1 for "Mapping the mammalian dark metabolome by *in vivo* isotope tracing"

Instrument details

Below are the instruments used to collect the analytical data on the synthesized compounds:

NMR: Bruker AVANCE NEO 400MHz,  AVANCE III HD 400 MHz.

LCMS: Agilent 1260&6120, Agilent 1260&6125B, Agilent 1260&6135B.

Milligram-scale separation SFC instrument: Waters SFC 150AP/150Mgm, 80/80Q

Gram-scale prep separation SFC instrument: Waters SFC 350

LC-HRMS: Vanquish UHPLC system coupled with an Thermo Orbitrap Exploris 480.

**Synthesis of HKMS-0291:**

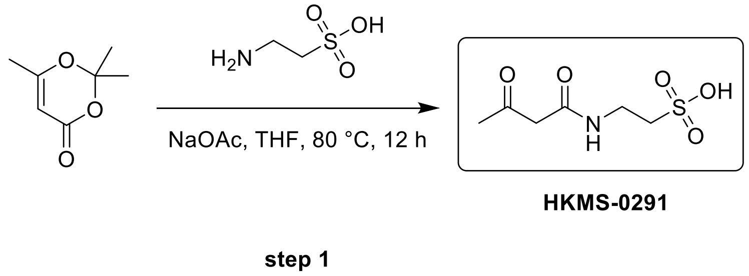


**2-(3-oxobutanamido) ethane-1-sulfonic acid**

To a solution of 2,2,6-trimethyl-1,3-dioxin-4-one (0.2 g, 1.41 mmol, 186.0 µL, 1.0 eq) and 2-aminoethanesulfonic acid (176.07 mg, 1.41 mmol, 175.4 µL, 1.0 eq) in THF (3 mL) was added NaOAc (230.83 mg, 2.81 mmol, 2.0 eq). The mixture was stirred at 80 °C for 12hr. The reaction mixture was concentrated under reduced pressure to give a residue. The residue was purified by prep-HPLC (column: Welch Utimate Hilic 100*25mm; mobile phase: [$H_2O$ (10mM $NH_4HCO_3$)-ACN]; gradient:95%-50% B over 15.0 min) 2-(3-oxobutanoylamino) ethanesulfonic acid (4.1 mg, 18.17 µmol, 1.29% yield, 92.72% purity) as a yellow oil.

LCMS: (M+H$^+$): 212.1 @ 4.036 min (97_80CD_10_HILIC_Amide_PH3_2)

$^1$H NMR: (400 MHz, DMSO-$d_6$) δ 7.98 (br s, 1H), 7.37 - 6.77 (m, 4H), 3.31 (br d, $J$ = 1.6 Hz, 2H), 3.27 (s, 2H), 2.57 - 2.53 (m, 2H), 2.14 (s, 3H)

$^{13}$C NMR: (100 MHz, DMSO-$d_6$) δ 203.61, 203.58, 166.86, 166.34, 115.53, 91.49, 51.74, 50.88, 50.62, 36.28, 36.00, 35.35, 30.42, 21.35, 19.43, 17.64

**Synthesis of HKMS-0300**

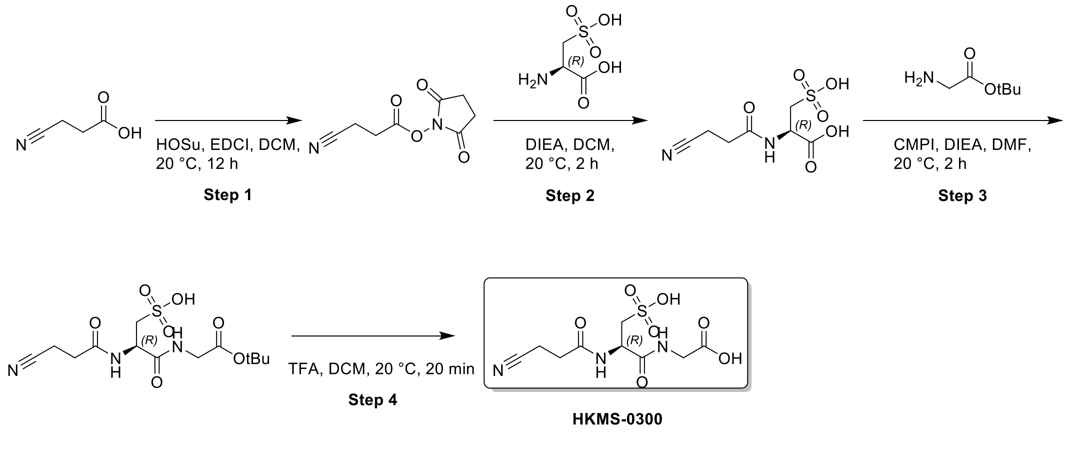

**Step 1:**

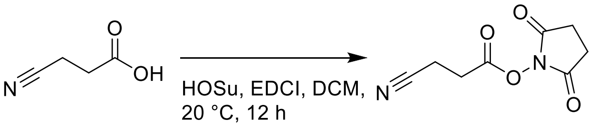


**Step 1**

**(2,5-dioxopyrrolidin-1-yl) 3-cyanopropanoate**

To a solution of 3-cyanopropanoic acid (800 mg, 8.07 mmol, 1 *eq*) in DCM (10 mL) was added EDCI (1.86 g, 9.69 mmol, 1.2 *eq*) and HOSu (1.02 g, 8.88 mmol, 1.1 *eq*) at 0°C. The mixture was stirred at 20°C for 12 h. The reaction mixture was diluted with $H_2O$ (40 mL) and extracted with DCM (20 mL * 3). The combined organic layers were washed with brine (40 mL), dried over $Na_2SO_4$, filtered and concentrated under reduced pressure to give a residue. Compound (2,5-dioxopyrrolidin-1-yl) 3-cyanopropanoate (750 mg, crude) was obtained as a yellow solid and used into the next step.

**Step 2 :**

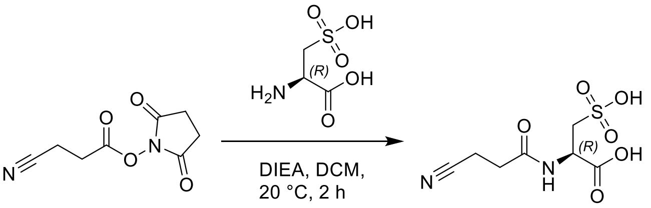


**Step 2**

**(2R)-2-(3-cyanopropanoylamino)-3-sulfo-propanoic acid**

To a solution of (2,5-dioxopyrrolidin-1-yl) 3-cyanopropanoate (750 mg, 3.82 mmol, 1 *eq*), (2R)-2-amino-3-sulfo-propanoic acid (517.40 mg, 3.06 mmol, 0.8 *eq*) in DCM (10 mL) was added DIEA (988.29 mg, 7.65 mmol, 1.33 mL, 2 *eq*). The mixture was stirred at 20 °C for 2 h. The reaction mixture was concentrated under reduced pressure to remove solvent. The residue was purified by prep-HPLC(TFA condition). (column: Kromasil C18(W) 150*30*10;mobile phase: [$H_2O$(0.1% TFA)-ACN];gradient:1%-8% B over 12.0 min ) Compound (2R)-2-(3-cyanopropanoylamino)-3-sulfo-propanoic acid (570 mg, 99%< purity) was obtained as a white solid and used into the next step.

$^1$H NMR: (400 MHz, DEUTERIUM OXIDE) δ 4.80-4.77 (m, 1H), 3.47-3.31 (m, 2H), 2.71 (s, 2H), 2.63-2.49 (m, 2H)

**Step 3 :**

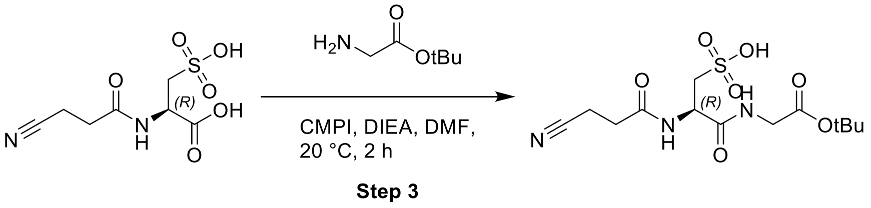


**(2R)-3-[(2-tert-butoxy-2-oxo-ethyl)amino]-2-(3-cyanopropanoylamino)-3-oxo-propane-1-sulfonic acid**

A mixture of (2R)-2-(3-cyanopropanoylamino)-3-sulfo-propanoic acid (300 mg, 1.20 mmol, 1 *eq*), tert-butyl 2-aminoacetate;hydrochloride (221.07 mg, 1.32 mmol, 1.1 *eq*), CMPI (398.18 mg, 1.56 mmol, 1.3 *eq*), DIEA (309.89 mg, 2.40 mmol, 417.6 µL, 2 *eq*) in DMF (5 mL) was degassed and purged with $N_2$ for 3 times, and then the mixture was stirred at 20 °C for 2 h under $N_2$ atmosphere. The reaction mixture was filtered and concentrated under reduced pressure to give a residue. The residue was purified by prep-HPLC(neutral condition). (column: Waters Xbridge BEH C18 100*30mm;mobile phase: [$H_2O$(10mM $NH_4HCO_3$)-ACN];gradient:1%-15% B over 15.0 min) Compound (2R)-3-[(2-tert-butoxy-2-oxo-ethyl)amino]-2-(3-cyanopropanoylamino)-3-oxo-propane-1-sulfonic acid (35 mg, 96.32 µmol, 8.03% yield, 99%< purity) was obtained as a white solid and used into the next step.

**Step 4:**

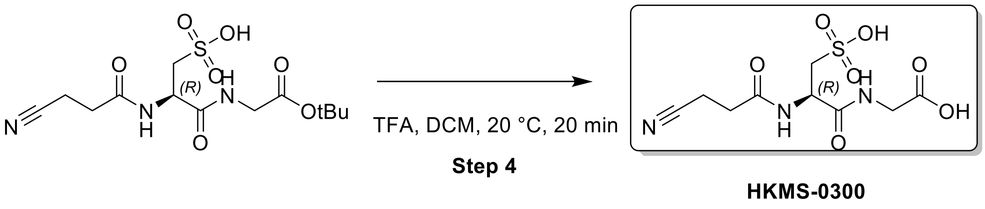


**2-[[(2R)-2-(3-cyanopropanoylamino)-3-sulfo-propanoyl]amino]acetic acid**

To a solution of (2R)-3-[(2-tert-butoxy-2-oxo-ethyl)amino]-2-(3-cyanopropanoylamino)- 3-oxo-propane-1-sulfonic acid (30 mg, 82.56 µmol, 1 *eq*) in DCM (1.5 mL) was added TFA (0.5 mL). The mixture was stirred at 20 °C for 20 min. The reaction mixture was concentrated by $N_2$ to remove solvent. The residue was purified by prep-HPLC(TFA condition). (column: Kromasil 150*30;mobile phase: [$H_2O$(0.1% TFA)-ACN];gradient: 1%-8% B over 12.0 min ) Compound 2-[[(2R)-2-(3-cyanopropanoylamino)-3-sulfo-propanoyl]amino]acetic acid (2.8 mg, 5.61 µmol, 6.79% yield, 84.34% purity, TFA) was obtained as a white solid.

LCMS: (M+1):308.1 @ 2.343min (0-30 % ACN in $H_2O$, 10 min)

$^1$H NMR:(400 MHz, DEUTERIUM OXIDE) δ 4.80-4.77 (m, 1H), 3.97 (s, 2H), 3.44-3.32 (m, 1H), 3.30-3.18 (m, 1H), 2.76-2.49 (m, 4H)

$^{13}$C NMR: (100 MHz, DMSO-$d_6$) δ 174.31, 171.86, 171.71, 171.49, 171.48, 171.47, 169.46, 120.83, 52.50, 52.22, 51.15, 41.44, 31.39, 30.91, 30.80, 12.77

**Synthesis of HKMS-0302**

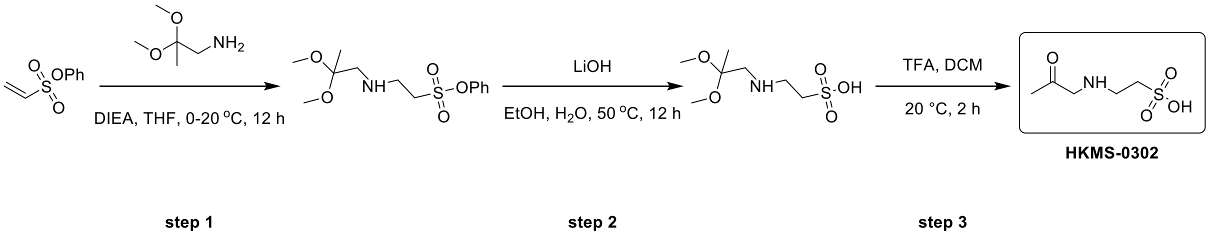


**Step 1:**

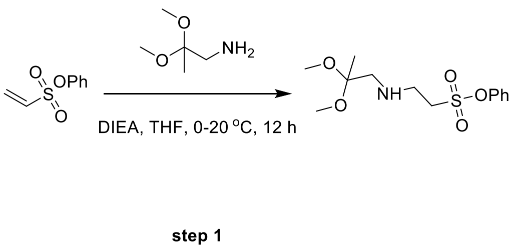


**Phenyl 2-((2,2-dimethoxypropyl)amino)ethane-1-sulfonate**

To a solution of 2,2-dimethoxypropan-1-amine (0.5 g, 4.20 mmol, 1.0 eq) in THF (5 mL) was added DIEA (1.90 g, 14.69 mmol, 2.56 mL, 3.5 eq) and phenyl ethenesulfonate (827.06 mg, 4.49 mmol, 1.07 eq) at 0°C. The mixture was stirred at 20°C for 12hr. The reaction mixture was poured into $H_2O$ (15 mL), and extracted with EtOAc(15 mLx3). The combined organic layers were washed with brine(15 mLx2), dried over $Na_2SO_4$, filtered and concentrated under reduced pressure to give a residue. The residue was purified by column chromatography ($SiO_2$, Petroleum ether/Ethyl acetate=1/0 to 3/1) to give phenyl 2-(2,2-dimethoxypropylamino) ethanesulfonate (1.2 g, 3.96 mmol, 94.27% yield) as a colourless oil.

$^1$H NMR: (400 MHz, CHLOROFORM-d) δ 7.49 - 7.40 (m, 2H), 7.38 - 7.29 (m, 3H), 3.50 - 3.43 (m, 2H), 3.31 - 3.26 (m, 2H), 3.23 (s, 6H), 2.76 (s, 2H), 1.38 (s, 3H)

**Step 2**

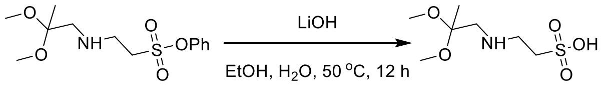


step 2

**2-((2,2-dimethoxypropyl)amino)ethane-1-sulfonic acid**

To a solution of phenyl 2-(2,2-dimethoxypropylamino)ethanesulfonate (0.5 g, 1.65 mmol, 1.0 eq) in EtOH (2 mL) was added NaOH (131.84 mg, 3.30 mmol, 2.0 eq) and $H_2O$ (2 mL). The mixture was stirred at 50 °C for 12hr. The reaction mixture was poured into $H_2O$ (10 mL), and extracted with EtOAc(10 mLx3). The aqueous phase was adjust pH to 5, and extracted with EtOAc(10 mLx3). The combined organic layers were washed with brine(10 mLx2), dried over $Na_2SO_4$, filtered and concentrated under reduced pressure to give 2-(2,2-dimethoxypropylamino) ethanesulfonic acid (0.4 g, crude) as a white solid.

**Step 3 :**

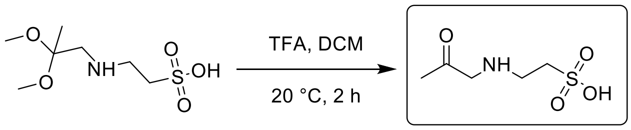


HKMS-0302

step 3

**2-((2-oxopropyl)amino)ethane-1-sulfonic acid**

To a solution of 2-(2,2-dimethoxypropylamino) ethanesulfonic acid (0.4 g, 1.76 mmol, 1.0 eq) in DCM (2 mL) was added TFA (614.00 mg, 5.38 mmol, 400.0 µL, 3.06 eq). The mixture was stirred at 20 °C for 2hr. The reaction mixture was concentrated under reduced pressure to give a residue. The residue was purified by prep-HPLC (column: PrePulite Perfect 150*30; mobile phase: [$H_2O$ (0.1% TFA)-ACN]; gradient:1%-6% B over 12.0 min) to afford 2-(acetonylamino) ethanesulfonic acid (0.01 g, 50.99 µmol, 2.90% yield, 92.4% purity) as a pink solid.

LCMS: $(M+H^+)$: 181.99 @ 4.061 min, (90_50_10min_10cm_2_PH3_Amide)

$^1H$ NMR: (400 MHz, DEUTERIUM OXIDE) δ 4.19 (s, 2H), 3.51 - 3.37 (m, 2H), 3.29 - 3.16 (m, 2H), 2.22 (s, 3H)

$^{13}C$ NMR:(100 MHz, DEUTERIUM OXIDE) δ 203.03, 55.21, 47.65, 46.47, 43.05, 26.59, 26.49

#### Synthesis of HKMS-0310

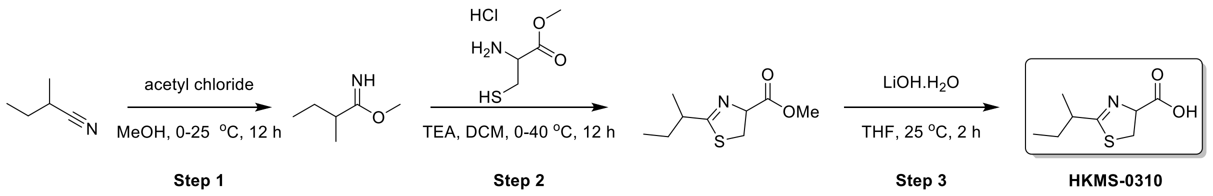


#### Step 1:

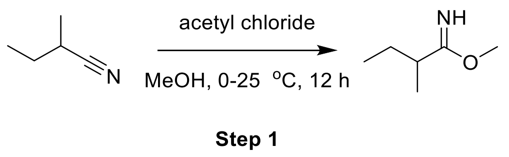


##### Methyl 2-methylbutanimidate

To a solution of 2-methylbutanenitrile (2 g, 24.06 mmol, 2.44 mL, 1 eq) in MeOH (10 mL) was added acetyl chloride (9.44 g, 120.29 mmol, 8.55 mL, 5 eq) at 0 °C. The mixture was stirred at 25 °C for 12 h. The reaction mixture was concentrated under reduced pressure to remove solvent. The crude product was purified by re-crystallization from MTBE (30 ml) at 25 ºC. The mixture was collected by filtration and the solid was concentrated under reduced pressure to give a residue. Compound methyl 2-methylbutanimidate (3 g, crude) was obtained as a white solid. The crude product methyl 2-methylbutanimidate (3 g, crude) was used into the next step without further purification.

$^1$H NMR: (400 MHz, DMSO-d6) δ 4.10 (s, 3H), 2.76 (sxt, *J* = 7.0 Hz, 1H), 1.69-1.47 (m, 2H), 1.17 (d, *J* = 7.0 Hz, 3H), 0.86 (t, *J* = 7.5 Hz, 3H)

#### Step 2 :

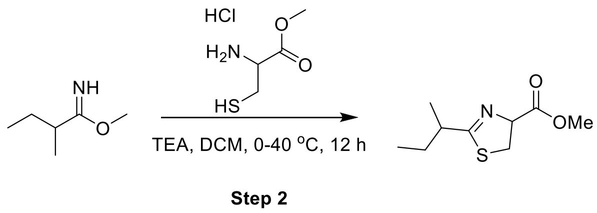


##### Methyl 2-sec-butyl-4,5-dihydrothiazole-4-carboxylate

To a solution of methyl 2-methylbutanimidate (1 g, 8.68 mmol, 1 eq) and methyl 2-amino-3-sulfanyl-propanoate;hydrochloride (1.49 g, 8.68 mmol, 1 eq) in DCM (10 mL) was added TEA (3.51 g, 34.73 mmol, 4.83 mL, 4 eq) at 0°C. The mixture was stirred at 40 °C for 12 h. The mixture was collected by filtration and the filtrate was concentrated under reduced pressure to give a residue. The residue was purified by flash silica gel chromatography (ISCO®; 4 g SepaFlash® Silica Flash Column, Eluent of

0~20% Ethyl acetate/Commercial hexanes gradient @ 40mL/min). Compound methyl 2-sec-butyl-4,5-dihydrothiazole-4-carboxylate (70 mg, 347.77 μmol, 4.01% yield) was obtained as a colorless oil.

**Step 3 :**

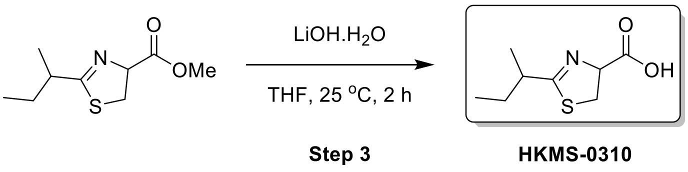


**2-sec-butyl-4,5-dihydrothiazole-4-carboxylic acid**

A mixture of methyl 2-sec-butyl-4,5-dihydrothiazole-4-carboxylate (50 mg, 248.40 μmol, 1 eq), LiOH.$H_2O$ (20.85 mg, 496.81 μmol, 2 eq) in $H_2O$ (0.2 mL) and THF (1 mL) was degassed and purged with $N_2$ for 3 times, and then the mixture was stirred at 25 °C for 2 h under $N_2$ atmosphere. The reaction mixture was concentrated under reduced pressure to remove solvent. The residue was purified by prep-HPLC (neutral condition).(column: WePure Biotech XPt C18150*40*7um;mobile phase: [$H_2O$(10mM $NH_4HCO_3$)-ACN];gradient:5%-45% B over 8.0 min). Compound 2-sec-butyl-4,5-dihydrothiazole-4-carboxylic acid (39.1 mg, 204.98 μmol, 58.94% yield, 98.17% purity) was obtained as a yellow oil.

LCMS: (M+$H^+$): 188.0 @ 1.589 min (0-60 % ACN in $H_2O$, 6 min)

$^1$H NMR:(400 MHz, DMSO-d6) δ 4.68 (dt, *J* = 5.4, 9.0 Hz, 1H), 3.45-3.27 (m, 2H), 2.59-2.51 (m, 1H), 1.63-1.36 (m, 2H), 1.09 (dd, *J* = 2.6, 6.9 Hz, 3H), 0.85 (t, *J* = 7.4 Hz, 3H)

$^{13}$C NMR:(100 MHz, DMSO-d6) δ 175.36, 175.23, 175.11, 173.08, 172.98, 172.10, 172.03, 55.69, 55.54, 53.14, 41.81, 41.70, 41.61, 41.48, 27.53, 27.23, 27.12, 18.18, 18.10, 17.88, 17.84, 12.20

**Synthesis of HKMS-0322**

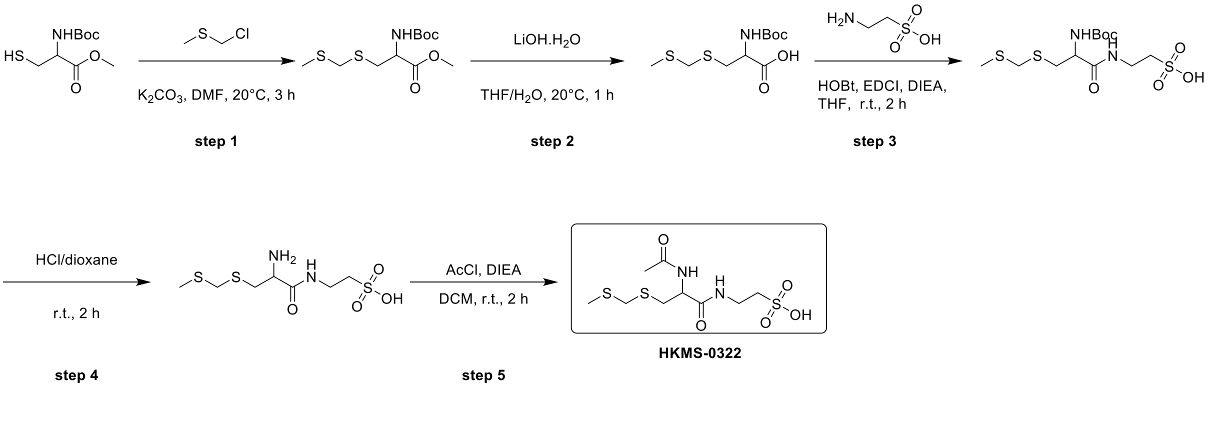

**Step 1 :**

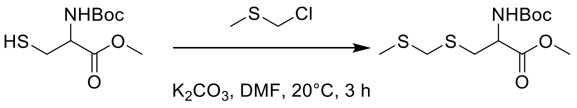


**step 1**

**Methyl N-(tert-butoxycarbonyl)-S-((methylthio)methyl) cysteinate:**

To a solution of chloro (methylsulfanyl) methane (1.23 g, 12.75 mmol, 1.07 mL, 1.5 eq) and methyl (2R)-2-(tert-butoxycarbonylamino)-3-sulfanyl-propanoate (2 g, 8.50 mmol, 1.0 eq) in DMF (20 mL) was added $K_2CO_3$ (2.35 g, 17.00 mmol, 2.0 eq). The mixture was stirred at 20 °C for 3hr. The reaction mixture was poured into $H_2O$ (30 mL), and extracted with EtOAc(30 mLx3). The combined organic layers were washed with brine(30 mLx2), dried over $Na_2SO_4$, filtered and concentrated under reduced pressure to give a residue. The residue was purified by column chromatography ($SiO_2$, Petroleum ether/Ethyl acetate=1/0 to 3/1) to afford methyl 2-(tert-butoxycarbonylamino)-3-(methylsulfanylmethylsulfanyl)propanoate (1.8 g, 6.09 mmol, 71.68% yield) as a yellow oil.

**Step 2:**

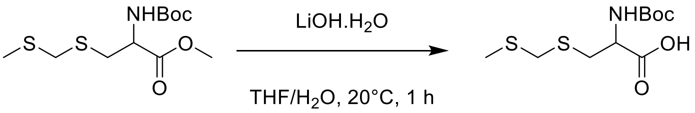


**step 2**

**N-(tert-butoxycarbonyl)-S-((methylthio)methyl)cysteine**

To a solution of methyl 2-(tert-butoxycarbonylamino)-3-(methylsulfanylmethylsulfanyl) propanoate (1 g, 3.39 mmol, 1.0 eq) in THF (10 mL) and $H_2O$ (2 mL) was added $LiOH.H_2O$ (426.14 mg, 10.16 mmol, 3.0 eq). The mixture was stirred at 20 °C for 1hr. The reaction mixture was poured into $H_2O$ (10 mL), and extracted with EtOAc (10 mLx3). The aqueous phase was adjusted pH to 3, and extracted with EtOAc(10 mLx3). The combined organic layers were washed with brine(30 mLx2), dried over $Na_2SO_4$, filtered and concentrated under reduced pressure to give 2-(tert-butoxycarbonylamino)-3-(methylsulfanylmethylsulfanyl)propanoic acid (0.36 g, 1.28 mmol, 37.79% yield) as a yellow oil.

$^{1}H$ NMR: (400 MHz, CHLOROFORM-d) δ 4.56 (br d, $J$ = 4.0 Hz, 1H), 3.69 (d, $J$ = 1.7 Hz, 2H), 3.30 - 2.95 (m, 2H), 2.17 (s, 3H), 1.47 (s, 9H)

**Step 3 :**

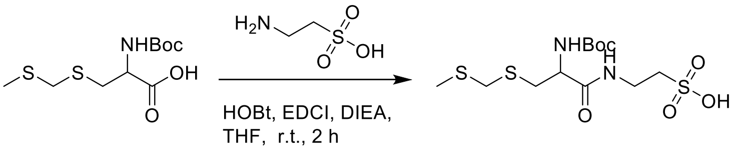


**step 3**

**2-(2-((tert-butoxycarbonyl) amino)-3-(((methylthio)methyl)thio) propanamido) ethane-1-sulfonic acid**

To a solution of 2-(tert-butoxycarbonylamino)-3-(methylsulfanylmethylsulfanyl) propanoic acid (0.31 g, 1.10 mmol, 1.0 eq) and 2-aminoethanesulfonic acid (137.87 mg, 1.10 mmol, 137.3 μL, 1.0 eq) in THF (5 mL) was added HOBt (163.75 mg, 1.21 mmol, 1.1 eq) and EDCI (232.31 mg, 1.21 mmol, 1.1 eq), DIEA (284.76 mg, 2.20 mmol, 383.8 μL, 2.0 eq). The mixture was stirred at 20 °C for 2hr. The reaction mixture was concentrated under reduced pressure to give a residue. The residue was purified by prep-HPLC (column: Waters Xbridge BEH C18 100*30mm; mobile phase: [$H_2O$ (10mM $NH_4HCO_3$)-ACN]; gradient:1%-40% B over 12.0 min) to afford 2-[[2-(tert-butoxycarbonylamino)-3-(methylsulfanylmethylsulfanyl) propanoyl] amino] ethanesulfonic acid (0.25 g, 643.46 μmol, 58.41% yield) as a white solid.

**Step 4 :**

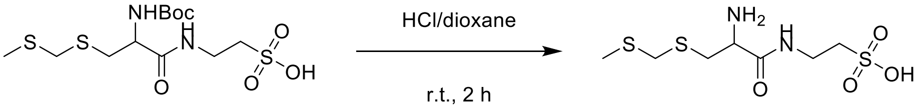


**step 4**

**2-(2-amino-3-(((methylthio)methyl) thio) propanamido) ethane-1-sulfonic acid**

To a solution of 2-[[2-(tert-butoxycarbonylamino)-3-(methylsulfanylmethylsulfanyl) propanoyl] amino] ethanesulfonic acid (0.05 g, 128.69 μmol, 1.0 eq) in HCl/dioxane (1 mL). The mixture was stirred at 20 °C for 2hr. The reaction mixture was concentrated under reduced pressure to give 2-[[2-amino-3-(methylsulfanylmethylsulfanyl) propanoyl] amino] ethanesulfonic acid (0.05 g, crude, HCl) as a white solid.

**Step 5:**

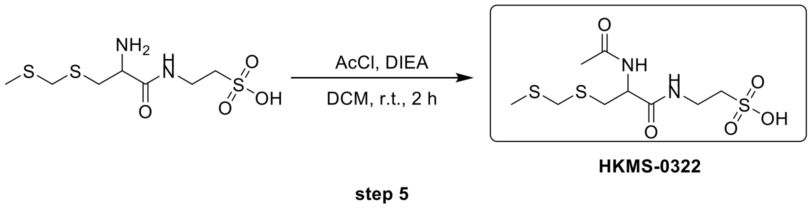

**2-(2-acetamido-3-(((methylthio)methyl) thio) propanamido) ethane-1-sulfonic acid**

To a solution of 2-[[2-amino-3-(methylsulfanylmethylsulfanyl) propanoyl] amino] ethanesulfonic acid (0.05 g, 153.91 μmol, 1.0 eq, HCl) and acetyl chloride (12.08 mg, 153.91 μmol, 10.9 μL, 1.0 eq) in DCM (1 mL) was added TEA (46.72 mg, 461.73 μmol, 64.3 μL, 3.0 eq). The mixture was stirred at 20 °C for 2hr. The reaction mixture was concentrated under reduced pressure to give a residue. The residue was purified by prep-HPLC (column: WePure Biotech XPt C18 100*30mm; mobile phase: [$H_2O$ (0.1% TFA)-ACN]; gradient:1%-25% B over 12.0 min) to give compound 2-[[2-acetamido-3-(methylsulfanylmethylsulfanyl) propanoyl] amino] ethane sulfonic acid (13.9 mg, 42.06 μmol, 27.33% yield, 99%< purity) as a white solid.

LCMS: ($M+H^+$): 331.0 @ 1.885 min, (0_30AB_6min_ELSD)

$^1$H NMR: (400 MHz, DEUTERIUM OXIDE) δ 4.47 (dd, $J$ = 5.2, 8.5 Hz, 1H), 3.68 (s, 2H), 3.53 (dt, $J$ = 2.8, 6.8 Hz, 2H), 3.12 - 2.97 (m, 3H), 2.88 (dd, $J$ = 8.5, 14.2 Hz, 1H), 2.09 (s, 3H), 1.99 (s, 3H)

$^{13}$C NMR: (100 MHz, DEUTERIUM OXIDE) δ 174.37, 172.06, 53.21, 49.46, 37.12, 35.14, 32.18, 21.71, 13.67

**Synthesis of HKMS-0078**

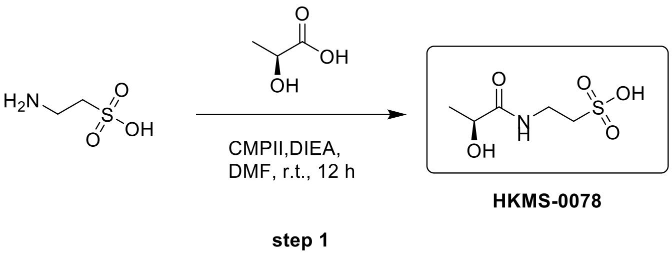

**(S)-2-(2-hydroxypropanamido) ethane-1-sulfonic acid**

To a solution of 2-aminoethanesulfonic acid (69.47 mg, 555.08 μmol, 69.2 μL, 1.0 eq) and (2S)-2-hydroxypropanoic acid (0.05 g, 555.08 μmol, 1 eq) in DMF (1 mL) was added CMPI (283.63 mg, 1.11 mmol, 2.0 eq) and DIEA (286.96 mg, 2.22 mmol, 386.7 μL, 4.0 eq). The mixture was stirred at 20 °C for 12hr. The reaction mixture was concentrated under reduced pressure to give a residue. The residue was purified by prep-HPLC (column: Kromasil C18(W) 150*30*10; mobile phase: [$H_2O$ (0.1%TFA)-ACN]; gradient:1%-30% B over 12.0 min) to afford 2-[[(2S)-2-hydroxypropanoyl] amino] ethanesulfonic acid (10.8 mg, 54.76 µmol, 9.87% yield, 99%< purity) as a colourless oil.

LCMS: ($M+H^+$): 198.1 @ 1.939 min (T3_0_30AB_10min_ELSD)

$^1$H NMR: (400 MHz, METHANOL-$d_4$) δ 4.12 (d, $J$ = 6.8 Hz, 1H), 3.66 (d, $J$ = 5.2 Hz, 2H), 2.99 (t, $J$ = 6.4 Hz, 2H), 1.35 (d, $J$ = 6.8 Hz, 3H)

$^{13}$C NMR: (100 MHz, METHANOL-$d_4$) δ 48.27, 47.85, 47.42, 47.63 (t, $J$ = 42.9 Hz, 1C), 46.99

**Synthesis of HKMS-0157**

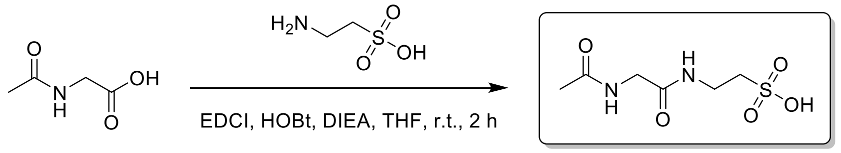


**HKMS-0157**

**2-(2-acetamidoacetamido) ethane-1-sulfonic acid**

To a solution of 2-acetamidoacetic acid (0.10 g, 853.95 µmol, 1.0 eq) and 2-aminoethanesulfonic acid (106.87 mg, 853.95 µmol, 106.4 µL, 1.0 eq) in THF (2 mL) was added HOBt (138.47 mg, 1.02 mmol, 1.2 eq) and EDCI (196.44 mg, 1.02 mmol, 1.2 eq) and DIEA (331.10 mg, 2.56 mmol, 446.2 µL, 3.0 eq). The mixture was stirred at 20 °C for 12hr. The reaction mixture was concentrated under reduced pressure to give a residue. The residue was purified by prep-HPLC (column: Welchrom CSH C18 100*30*7; mobile phase: [$H_2O$ (0.02% FA)-ACN]; gradient:1%-15% B over 12.0 min) to afford 2-[(2-acetamidoacetyl) amino] ethanesulfonic acid (63.3 mg, 282.29 µmol, 33.06% yield, 99%< purity) as a yellow oil.

LCMS: ($M+H^+$): 225.0 @ 1.958 min (T3_0_30AB_10min_ELSD)

$^1$H NMR: (400 MHz, DEUTERIUM OXIDE) δ 3.79 (s, 2H), 3.51 (t, $J$ = 6.7 Hz, 2H), 2.99 (t, $J$ = 6.7 Hz, 2H), 1.97 (s, 3H)

$^{13}$C NMR: (100 MHz, DEUTERIUM OXIDE) δ 174.90, 171.38, 49.45, 42.59, 35.13, 34.97, 21.69

**Synthesis of HKMS-0158**

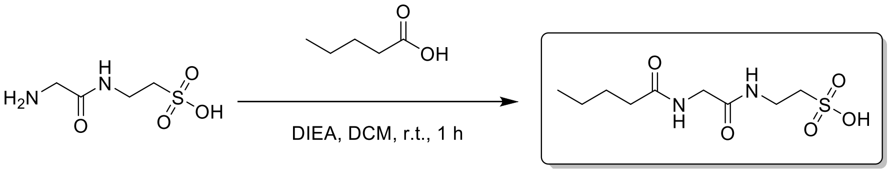

step 1

HKMS-0158

**2-(2-pentanamidoacetamido) ethane-1-sulfonic acid**

To a solution of 2- [(2-aminoacetyl) amino] ethanesulfonic acid (0.05 g, 228.67 μmol, 1.0 eq, HCl) and pentanoyl chloride (27.57 mg, 228.67 μmol, 27.7 μL, 1.0 eq) in DCM (1 mL) was added DIEA (88.66 mg, 686.00 μmol, 119.5 μL, 3.0 eq). The mixture was stirred at 20 °C for 1hr. The reaction mixture was concentrated under reduced pressure to give a residue. The residue was purified by prep-HPLC (column: PrePulite Perfect T3 150*30*5; mobile phase: [$H_2O$ (0.1% TFA)-ACN]; gradient:1%-20% B over 15.0 min) to afford 2-[[2-(pentanoylamino) acetyl] amino] ethanesulfonic acid (7 mg, 25.97 μmol, 11.36% yield, 98.81% purity) was obtained as a colorless oil.

LCMS: ($M+H^+$): 267.1 @1.746 min (0_30AB_6min_ELSD)

$^1$H NMR: (400 MHz, DEUTERIUM OXIDE) δ 3.80 (s, 2H), 3.52 (t, $J$ = 6.6 Hz, 2H), 3.00 (t, $J$ = 6.7 Hz, 2H), 2.25 (t, $J$ = 7.4 Hz, 2H), 1.50 (quin, $J$ = 7.5 Hz, 2H), 1.24 (sxt, $J$ = 7.4 Hz, 2H), 0.81 (t, $J$ = 7.3 Hz, 3H)

$^{13}$C NMR: (100 MHz, DEUTERIUM OXIDE) δ 178.09, 171.42, 49.71, 49.49, 42.53, 35.24, 35.02, 27.40, 27.27, 21.58, 21.51, 12.97

**Synthesis of HKMS-0159**

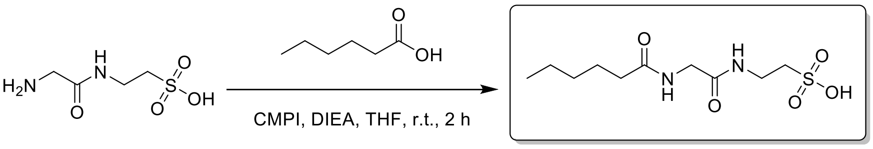

HKMS-0159

step 1

**2-(2-hexanamidoacetamido) ethane-1-sulfonic acid**

To a solution of 2-[(2-aminoacetyl) amino] ethanesulfonic acid (0.05 g, 228.67 μmol, 1.0 eq, HCl) and hexanoic acid (26.56 mg, 228.67 μmol, 28.6 μL, 1.0 eq) in THF (1 mL) was added DIEA (88.66 mg, 686.00 μmol, 119.5 μL, 3.0 eq) and CMPI (70.10 mg, 274.40 μmol, 1.2 eq). The mixture was stirred at 20 °C for 2hr. The reaction mixture was concentrated under reduced pressure to give a residue. The residue was purified by prep-HPLC (column: Welchrom CSH C18 100*30*7; mobile phase: [$H_2O$ (0.03%

TFA)-ACN]; gradient:1%-30% B over 12.0 min) to afford 2-[[2-(hexanoylamino) acetyl] amino] ethanesulfonic acid (14.2 mg, 49.43 μmol, 21.62% yield, 97.58% purity) as a yellow oil.

LCMS:(M+H$^+$): 281.1 @2.333 min (0_30AB_6min_ELSD)

$^1$H NMR: (400 MHz, DEUTERIUM OXIDE) δ 3.79 (s, 2H), 3.52 (t, $J$ = 6.7 Hz, 2H), 3.00 (t, $J$ = 6.7 Hz, 2H), 2.23 (t, $J$ = 7.5 Hz, 2H), 1.52 (br t, $J$ = 7.3 Hz, 2H), 1.30 - 1.12 (m, 4H), 0.78 (t, $J$ = 7.0 Hz, 3H)

$^{13}$C NMR: (100 MHz, DEUTERIUM OXIDE) δ 178.05, 171.37, 49.45, 42.49, 35.41, 34.98, 30.46, 24.75, 21.62, 13.14

**Synthesis of HKMS-0160**

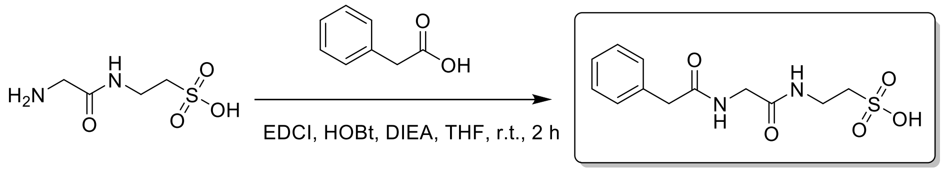


HKMS-0160

**(2-(2-phenylacetamido) acetamido) ethane-1-sulfonic acid**

To a solution of 2-[(2-aminoacetyl) amino] ethanesulfonic acid (0.05 g, 228.67 μmol, 1.0 eq, HCl) and 2-phenylacetic acid (31.13 mg, 228.67 μmol, 28.8 μL, 1.0 eq) in THF (1 mL) was added HOBt (37.08 mg, 274.40 μmol, 1.2 eq) and EDCI (52.60 mg, 274.4 μmol, 1.2 eq), DIEA (88.66 mg, 686.0 μmol, 119.5 μL, 3.0 eq). The mixture was stirred at 20 °C for 2hr. The reaction mixture was concentrated under reduced pressure to give a residue. The residue was purified by prep-HPLC (column: Waters Xbridge BEH C18 100*30mm*10um; mobile phase: [$H_2O$ (10mM $NH_4HCO_3$)-ACN]; gradient:1%-10% B over 15.0 min) to afford 2-[[2-[(2-phenylacetyl) amino] acetyl] amino] ethanesulfonic acid (9 mg, 29.97 μmol, 13.11% yield, 99%< purity) as a white solid.

LCMS: (M+H$^+$): 301.0 @1.875 min (0_30CD_6min_ELSD)

$^1$H NMR: (400 MHz, DEUTERIUM OXIDE) δ 7.37 - 7.23 (m, 5H), 3.80 (s, 2H), 3.61 (s, 2H), 3.50 (s, 2H), 2.97 (t, $J$ = 6.7 Hz, 2H)

$^{13}$C NMR: (100 MHz, DEUTERIUM OXIDE) δ 175.45, 171.25, 134.66, 129.31, 128.98, 127.41, 49.50, 42.75, 42.06, 35.04

**Synthesis of HKMS-0162**

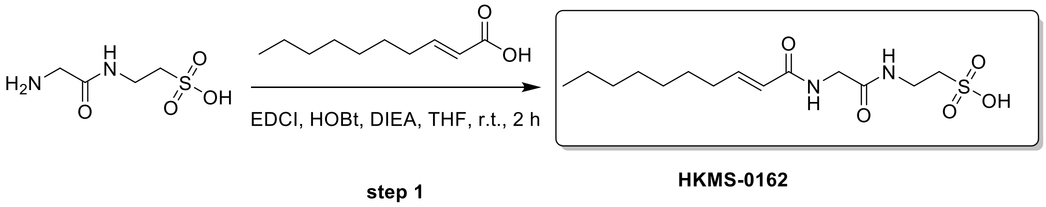


**(E)-2-(2-(dec-2-enamido) acetamido) ethane-1-sulfonic acid**

To a solution of 2-[(2-aminoacetyl) amino] ethanesulfonic acid (0.05 g, 228.67 μmol, 1.0 eq, HCl) and (E)-dec-2-enoic acid (38.93 mg, 228.67 μmol, 1 eq) in THF (2 mL) was added HOBt (37.08 mg, 274.40 μmol, 1.2 eq) and EDCI (52.60 mg, 274.40 μmol, 1.2 eq), DIEA (88.66 mg, 686.00 μmol, 119.5 μL, 3.0 eq). The mixture was stirred at 20 °C for 2hr. The reaction mixture was concentrated under reduced pressure to give a residue. The residue was purified by prep-HPLC (column: Waters Xbridge BEH C18 100*30mm*10um; mobile phase: [$H_2O$ (10mM $NH_4HCO_3$)-ACN]; gradient:1%-30% B over 15.0 min) to afford 2-[[2-[[(E)-dec-2-enoyl] amino] acetyl] amino] ethanesulfonic acid (3.7 mg, 11.02 μmol, 4.82% yield, 99.64% purity) as a white solid.

LCMS: ($M+H^+$): 335.1 @2.261 min (5_95CD_6min)

$^1$H NMR: (400 MHz, DEUTERIUM OXIDE) δ 6.85 - 6.69 (m, 1H), 5.93 (d, $J$ = 15.4 Hz, 1H), 3.86 (s, 2H), 3.51 (t, $J$ = 6.7 Hz, 2H), 2.99 (t, $J$ = 6.7 Hz, 2H), 2.15 (q, $J$ = 6.9 Hz, 2H), 1.44 - 1.12 (m, 10H), 0.82 - 0.71 (m, 3H)

$^{13}$C NMR: (100 MHz, DEUTERIUM OXIDE) δ 171.44, 171.24, 169.54, 147.96, 136.91, 121.81, 121.41, 49.47, 42.57, 39.18, 35.01, 31.78, 31.54, 31.06, 30.94, 28.28, 28.23, 27.96, 27.37, 22.00, 13.38

**Synthesis of HKMS-0163**

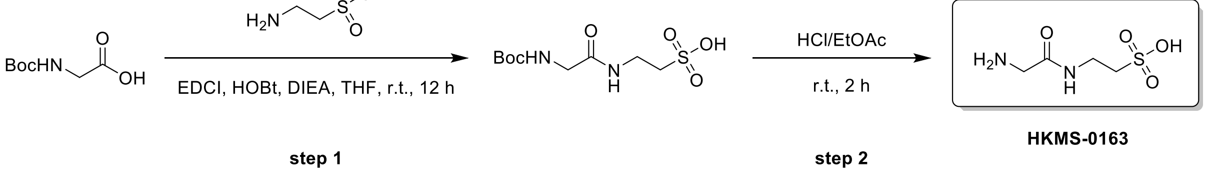


**Step 1:**

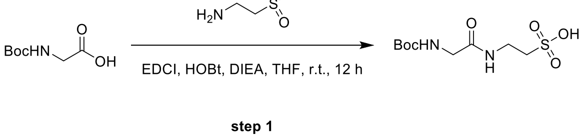

**2-(2-((tert-butoxycarbonyl) amino) acetamido) ethane-1-sulfonic acid**

To a solution of 2-(tert-butoxycarbonylamino) acetic acid (5.00 g, 28.54 mmol, 1.0 eq) and 2-aminoethanesulfonic acid (3.57 g, 28.54 mmol, 3.56 mL, 1.0 eq) in THF (70 mL) was added HOBt (4.63 g, 34.25 mmol, 1.2 eq) and EDCI (6.57 g, 34.25 mmol, 1.2 eq), DIEA (11.07 g, 85.63 mmol, 14.91 mL, 3.0 eq). The mixture was stirred at 20 °C for 12hr. The reaction mixture was concentrated under reduced pressure to give a residue. The residue was purified by prep-HPLC (column: WePure Biotech XP tC18 250*70*10um; mobile phase: [$H_2O$ (10mM $NH_4HCO_3$)-ACN]; gradient:0%-25% B over 20.0 min) to afford 2-[[2-(tert-butoxycarbonylamino) acetyl] amino] ethanesulfonic acid (3 g, 10.63 mmol, 37.23% yield) as a yellow oil.

**Step 2**

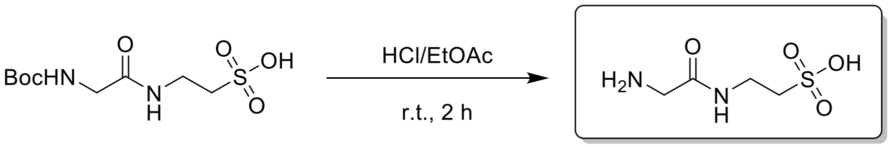


HKMS-0163

step 2

**2-(2-aminoacetamido) ethane-1-sulfonic acid**

To a solution of 2-[[2-(tert-butoxycarbonylamino) acetyl] amino] ethanesulfonic acid (1 g, 3.54 mmol, 1.0 eq) in HCl/EtOAc (10 mL). The mixture was stirred at 20 °C for 2hr. The reaction mixture was concentrated under reduced pressure to give a residue (600 mg). The residue (50 mg) was purified by prep-HPLC (column: Welchrom CSH C18 100*30*7; mobile phase: [$H_2O$ (0.025% FA)-ACN]; gradient:1%-15% B over 12.0 min) to afford 2-[(2-aminoacetyl) amino] ethanesulfonic acid (18.5 mg, 101.54 µmol, 2.87% yield, 99%< purity) as a white solid.

LCMS: ($M+H^+$): 183.1 @ 1.584 min (T3_0_30AB_10min_ELSD)

$^1$H NMR: (400 MHz, DEUTERIUM OXIDE) δ 3.65 (s, 2H), 3.49 (t, $J$ = 6.6 Hz, 2H), 2.96 (t, $J$ = 6.5 Hz, 2H)

$^{13}$C NMR: (100 MHz, DEUTERIUM OXIDE) δ 166.84, 49.47, 40.45, 35.17

**Synthesis of HKMS-0164**

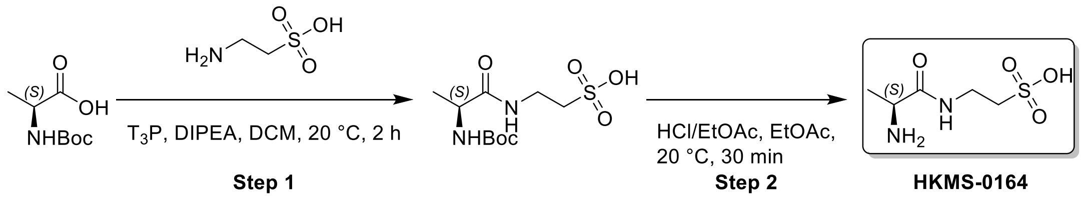


**Step 1**

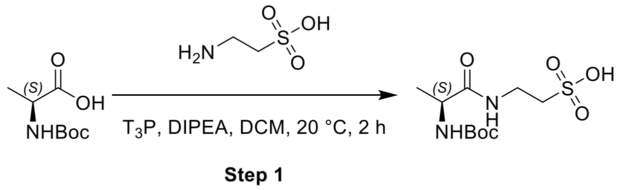


**2-[[(2S)-2-(tert-butoxycarbonylamino)propanoyl]amino]ethanesulfonic acid**

A mixture of (2S)-2-(tert-butoxycarbonylamino)propanoic acid (500 mg, 2.64 mmol, 1 *eq*), 2-aminoethanesulfonic acid (330.71 mg, 2.64 mmol, 329.4 µL, 1 *eq*), $T_3P$ (2.52 g, 3.96 mmol, 2.36 mL, 50% purity, 1.5 *eq*), DIEA (1.02 g, 7.93 mmol, 1.38 mL, 3 *eq*) in DCM (5 mL) was degassed and purged with $N_2$ for 3 times, and then the mixture was stirred at 20 °C for 2 h under $N_2$ atmosphere. The reaction mixture was diluted with $NH_4Cl$ (30 mL) and extracted with EtOAc (15 mL * 3). The combined organic layers were washed with brine (30 mL), dried over $Na_2SO_4$, filtered and concentrated under reduced pressure to give a residue. The residue was purified by prep-HPLC(neutral condition).(column: Waters Xbridge BEH C18 100*30mm*10um;mobile phase: [$H_2O$(10mM $NH_4HCO_3$)-ACN];gradient:1%-15% B over 12.0 min) Compound 2-[[(2S)-2-(tert-butoxycarbonylamino)propanoyl]amino]ethanesulfonic acid (50 mg, 165.69 µmol, 6.27% yield, 98.2% purity) was obtained as a white solid and used into the next step.

LCMS: (M-1):295.0 @ 0.773min (0-60 % ACN in $H_2O$, 2.5 min)

**Step 2**

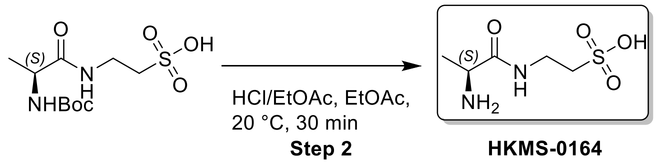


**2-[[(2S)-2-aminopropanoyl]amino]ethanesulfonic acid**

To a solution of 2-[[(2S)-2-(tert-butoxycarbonylamino)propanoyl]amino]ethanesulfonic acid (40 mg, 134.98 µmol, 1 *eq*) in EtOAc (1 mL) was added HCl/EtOAc (5 mL). The mixture was stirred at 20 °C for 30 min. The reaction was concentrated under reduced pressure to remove solvent. The residue was purified by prep-HPLC (HCl condition). (column: Welchrom CSH C18 100*30*7;mobile phase: [$H_2O$(0.02% HCl)-ACN];gradient:1%-15% B over 12.0 min) Compound 2-[[(2S)-2-aminopropanoyl]amino]ethanesulfonic acid (14.7 mg, 62.84 µmol, 42.32% yield, 99.47% purity, HCl) was obtained as a white solid.

LCMS: (M+1):197.1 @ 1.760min (0-30 % ACN in $H_2O$, 10 min)

SFC: (R.T.=2.055 min, 99%< ee)

$^1$H NMR: (400 MHz, DEUTERIUM OXIDE)

δ 4.02 (q, $J$ = 7.1 Hz, 1H), 3.60 (dt, $J$ = 2.0, 6.6 Hz, 2H), 3.07 (t, $J$ = 6.6 Hz, 2H), 1.48 (d, $J$ = 7.0 Hz, 3H)

$^{13}$C NMR: (100 MHz, DEUTERIUM OXIDE) δ 170.45, 49.36, 49.06, 35.19, 16.27

**Synthesis of HKMS-0172**

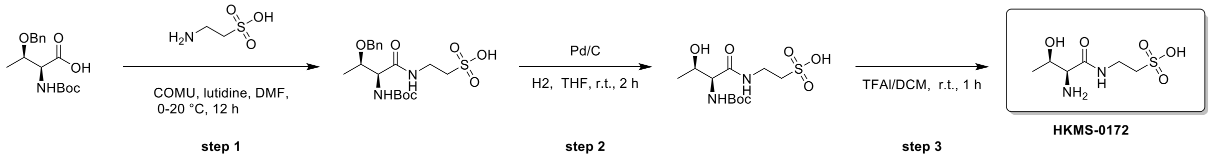


**Step 1**

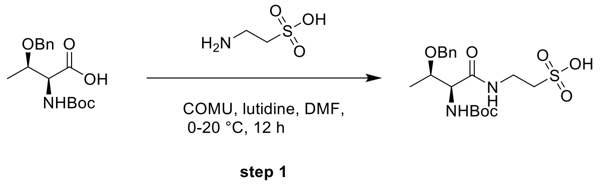


**2-((2S,3R)-3-(benzyloxy)-2-((tert-butoxycarbonyl) amino) butanamido) ethane-1-sulfonic acid**

To a solution of (2S,3R)-3-benzyloxy-2-(tert-butoxycarbonylamino) butanoic acid (0.50 g, 1.62 mmol, 1.0 eq) and 2-aminoethanesulfonic acid (202.27 mg, 1.62 mmol, 201.46 µL, 1.0 eq) in DMF (5 mL) was added COMU (692.19 mg, 1.62 mmol, 1.0 eq) and 2,6-dimethylpyridine (467.60 mg, 4.36 mmol, 508.3 µL, 2.7 eq) at 0°C. The mixture was stirred at 20°C for 12hr. The reaction mixture was concentrated under reduced pressure to give a residue. The residue was purified by prep-HPLC (column: Waters Xbridge BEH C18 100*30mm*10um; mobile phase: [$H_2O$ (10mM NH4HCO3)-ACN]; gradient:5%-35% B over 15.0 min) to afford 2-[[(2S,3R)-3-benzyloxy-2-(tert-butoxycarbonylamino) butanoyl] amino] ethanesulfonic acid (0.067 g, 160.87 µmol, 9.95% yield) as a white solid.

SFC-MS: (M+H$^+$): 317.1 @ 0.887 min (AD_EtOH_MNH3_5_50_34_35_3min)

**Step 2 :**

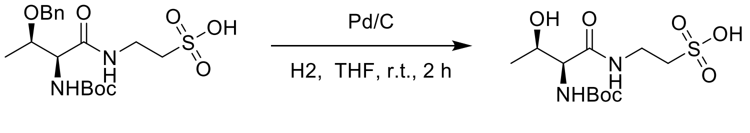


**step 2**

**2-((2S,3R)-2-((tert-butoxycarbonyl) amino)-3-hydroxybutanamido) ethane-1-sulfonic acid**

To a solution of 2-[[(2S,3R)-3-benzyloxy-2-(tert-butoxycarbonylamino) butanoyl] amino] ethanesulfonic acid (0.05 g, 120.05 µmol, 1.0 eq) in THF (2 mL) was added Pd/C (0.05 g, 46.98 µmol, 10% purity, 0.39 eq) under N2 atmosphere. The suspension was degassed and purged with H2 for 3 times. The mixture was stirred under H2 (15 Psi or atm.) at 20 °C for 2hr. It was filtrate and concentrated under reduced pressure to afford 2-[[(2S,3R)-2-(tert-butoxycarbonylamino)-3-hydroxy-butanoyl] amino] ethanesulfonic acid (0.039 g, 119.50 µmol, 99.54% yield) as a colourless oil.

**Step 3:**

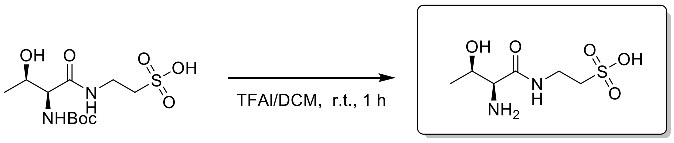


**HKMS-0172**

**step 3**

**2-((2S,3R)-2-amino-3-hydroxybutanamido) ethane-1-sulfonic acid**

To a solution of 2-[[(2S,3R)-2-(tert-butoxycarbonylamino)-3-hydroxy-butanoyl] amino] ethanesulfonic acid (0.039 g, 119.50 µmol, 1.0 eq) in DCM (1 mL) was added TFA (307.00 mg, 2.69 mmol, 0.2 mL, 22.53 eq). The mixture was stirred at 20 °C for 1hr. The reaction mixture was concentrated under reduced pressure to give a residue. The residue was purified by prep-HPLC (column: Kromasil C18(w) 250*50mm 10u; mobile phase: [$H_2O$ (0.05% HCl)-ACN]; gradient:1%-5% B over 12.0 min) to afford 2-[[(2S,3R)-2-amino-3-hydroxy-butanoyl] amino] ethanesulfonic acid (18.9 mg, 82.68 µmol, 69.19% yield, 98.98% purity) as a white solid.

LCMS: (M+H$^+$): 227.1 @ 1.873 min (T3_0_30AB_10min_ELSD)

$^1$H NMR: (400 MHz, DEUTERIUM OXIDE) δ 4.09 (t, $J$ = 6.3 Hz, 1H), 3.77 (d, $J$ = 6.0 Hz, 1H), 3.58

(t, $J = 6.5$ Hz, 2H), 3.05 (t, $J = 6.6$ Hz, 2H), 1.22 (d, $J = 6.4$ Hz, 3H)

$^{13}C$ NMR: (100 MHz, DEUTERIUM OXIDE) δ 167.70, 65.99, 58.75, 49.32, 35.32, 18.65

**Synthesis of HKMS-0174**

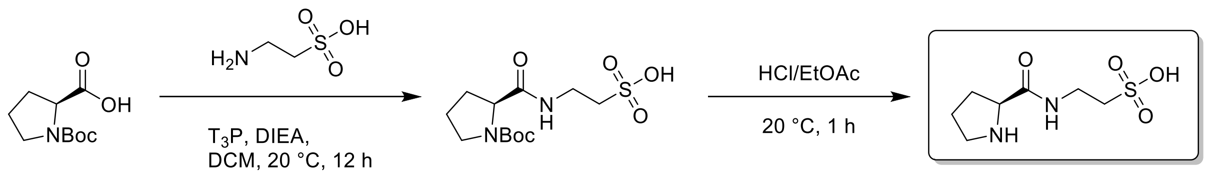


HKMS-0174

step 1 step 2

**Step 1 :**

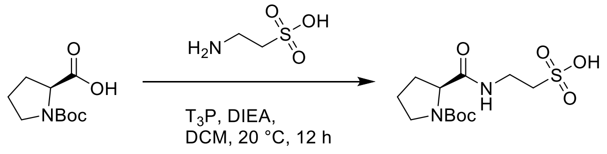


step 1

**(S)-2-(1-(tert-butoxycarbonyl) pyrrolidine-2-carboxamido) ethane-1-sulfonic acid**

To a solution of (2S)-1-tert-butoxycarbonylpyrrolidine-2-carboxylic acid (0.30 g, 1.39 mmol, 1.0 eq) and 2-aminoethanesulfonic acid (174.42 mg, 1.39 mmol, 173.7 μL, 1 eq) in DCM (3 mL) was added DIEA (540.39 mg, 4.18 mmol, 728.3 μL, 3.0 eq) and $T_3P$ (1.33 g, 2.09 mmol, 1.24 mL, 50% purity, 1.5 eq). The mixture was stirred at 20 °C for 12hr. The reaction mixture was quenched with $NaHCO_3$ (aq., 5 mL) and water (5 mL). The mixture was concentrated under reduced pressure to give a residue. The residue was purified by prep-HPLC (column: Waters Xbridge BEH C18 100*30mm*10um; mobile phase: [$H_2O$ (10mM NH4HCO3)-ACN]; gradient:1%-25% B over 12.0 min) to afford 2-[[(2S)-1-tert-butoxycarbonylpyrrolidine-2-carbonyl] amino] ethanesulfonic acid (0.2 g, crude) as a white solid.

**Step 2:**

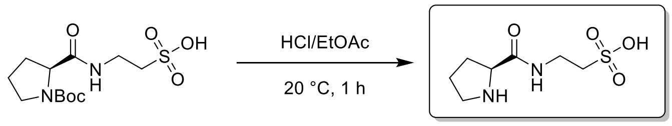


HKMS-0174

step 2

**(S)-2-(pyrrolidine-2-carboxamido) ethane-1-sulfonic acid**

To a solution of 2-[[(2S)-1-tert-butoxycarbonylpyrrolidine-2-carbonyl] amino] ethanesulfonic acid (0.05 g, 155.10 μmol, 1.0 eq) in HCl/EtOAc (1 mL). The mixture was stirred at 20 °C for 1hr. The reaction mixture was concentrated under reduced pressure to give a residue. The residue was purified by prep-HPLC (column: PrePulite Perfect T3 150*30*5; mobile phase: [$H_2O$ (0.1% TFA)-ACN]; B%:1%, isocratic elution mode) to afford 2-[[(2S)-pyrrolidine-2-carbonyl] amino] ethanesulfonic acid (23 mg, 103.48 μmol, 66.72% yield, 99%< purity) as a white solid.

SFC: @ 1.749 min (IH_EtOH_MNH3_10_50_34_35_4min)

LCMS: ($M+H^+$): 223.1 @ 2.601 min (T3_0_60AB_10min_ELSD)

$^1$H NMR: (400 MHz, DEUTERIUM OXIDE) δ 4.35 - 4.24 (m, 1H), 3.66 - 3.50 (m, 2H), 3.44 - 3.25 (m, 2H), 3.04 (t, $J$ = 6.5 Hz, 2H), 2.42 - 2.28 (m, 1H), 2.09 - 1.91 (m, 3H)

$^{13}$C NMR: (100 MHz, DEUTERIUM OXIDE) δ 169.29, 59.85, 49.37, 46.42, 35.42, 29.46, 23.73

**Synthesis of HKMS-0182**

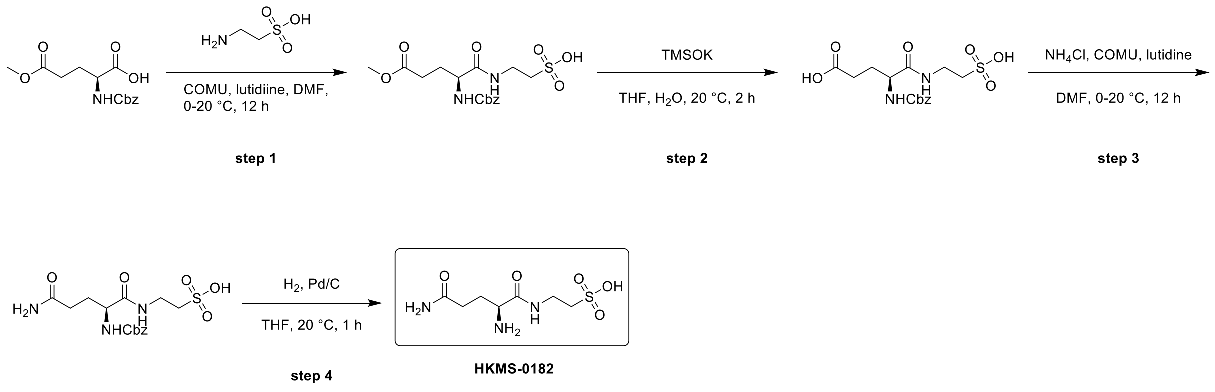


**Step 1:**

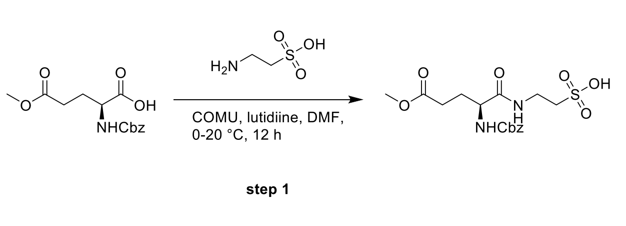


**(S)-2-(2-(((benzyloxy)carbonyl) amino)-5-methoxy-5-oxopentanamido) ethane-1-sulfonic acid**

To a solution of 2-aminoethanesulfonic acid (1.70 g, 13.55 mmol, 1.69 mL, 1.0 eq) and (2S)-2-(benzyloxycarbonylamino)-5-methoxy-5-oxo-pentanoic acid (4 g, 13.55 mmol, 1.0 eq) in DMF (40 mL) was added COMU (5.80 g, 13.55 mmol, 1.0 eq) and 2,6-dimethylpyridine (4.35 g, 40.64 mmol, 4.73 mL, 3.0 eq) at 0°C. The mixture was stirred at 20°C for 12hr. The reaction mixture was concentrated under reduced pressure to give a residue. The residue was purified by prep-HPLC (column: WePure Biotech Phenyl-Hexyl 250*70 7u; mobile phase: [$H_2O$ (10mM $NH_4HCO_3$)-ACN]; gradient:0%-30% B over 20.0 min) to afford 2-[[(2S)-2-(benzyloxycarbonylamino)-5-methoxy-5-oxo-pentanoyl] amino] ethanesulfonic acid (3.4 g, 8.45 mmol, 62.37% yield) as a yellow oil.

$^1$H NMR: (400 MHz, DEUTERIUM OXIDE) δ 7.46 - 7.26 (m, 5H), 5.06 (br d, $J$ = 6.0 Hz, 2H), 4.07 - 3.97 (m, 1H), 3.70 - 3.44 (m, 5H), 3.06 - 2.86 (m, 2H), 2.50 - 2.32 (m, 2H), 2.15 - 1.75 (m, 2H)

**Step 2 :**

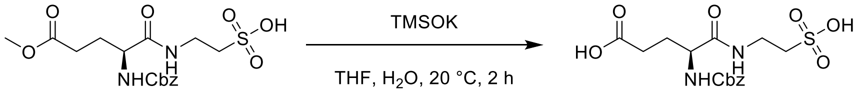


step 2

**(S)-4-(((benzyloxy)carbonyl) amino)-5-oxo-5-((2-sulfoethyl) amino) pentanoic acid**

To a solution of 2-[[(2S)-2-(benzyloxycarbonylamino)-5-methoxy-5-oxo-pentanoyl] amino] ethanesulfonic acid (1.7 g, 4.22 mmol, 1.0 eq) in THF (15 mL) and $H_2O$ (3 mL) was added TMSOK (1.08 g, 8.45 mmol, 2.0 eq). The mixture was stirred at 20 °C for 2hr. The reaction mixture was concentrated under reduced pressure to give (4S)-4-(benzyloxycarbonylamino)-5-oxo-5-(2-sulfoethylamino) pentanoic acid (1.69 g, 2.18 mmol, 51.50% yield, 50% purity) as a yellow solid.

**Step 3 :**

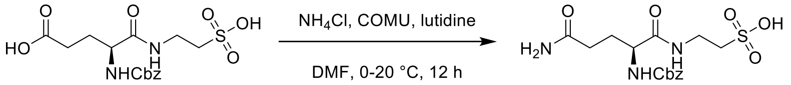


step 3

**(S)-2-(5-amino-2-(((benzyloxy)carbonyl) amino)-5-oxopentanamido) ethane-1-sulfonic acid**

To a solution of (4S)-4-(benzyloxycarbonylamino)-5-oxo-5-(2-sulfoethylamino) pentanoic acid (1 g, 1.29 mmol, 1.0 eq) in DMF (10 mL) was added COMU (551.34 mg, 1.29 mmol, 1.0 eq) and 2,6-dimethylpyridine (413.82 mg, 3.86 mmol, 449.8 µL, 3.0 eq), $NH_4Cl$ (344.31 mg, 6.44 mmol, 5.0 eq) at 0°C. The mixture was stirred at 20 °C for 12hr. The reaction mixture was concentrated under reduced pressure to give a residue. The residue was purified by prep-HPLC (column: Waters Xbridge BEH C18 100*30mm*10um; mobile phase: [$H_2O$ (10mM $NH_4HCO_3$)-ACN]; gradient:1%-15% B over 15.0 min) to afford 2-[[(2S)-5-amino-2-(benzyloxycarbonylamino)-5-oxo-pentanoyl] amino] ethanesulfonic acid

(46 mg, 103.30 μmol, 8.02% yield, 87% purity) as a white solid.

**Step 4:**

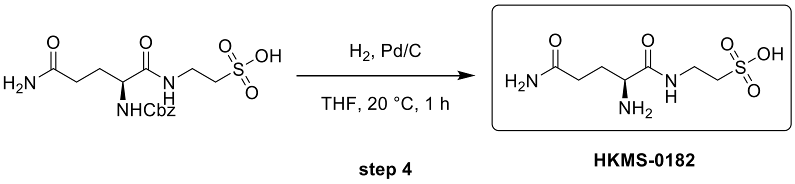


**(S)-2-(2,5-diamino-5-oxopentanamido) ethane-1-sulfonic acid**

To a solution of 2-[[(2S)-5-amino-2-(benzyloxycarbonylamino)-5-oxo-pentanoyl] amino] ethanesulfonic acid (0.036 g, 92.93 μmol, 1.0 eq) in THF (2 mL) was added 10% Pd/C (30.00 mg, 28.19 μmol, 0.3 eq) under $N_2$ atmosphere. The suspension was degassed and purged with $H_2$ for 3 times. The mixture was stirred under $H_2$ (15 Psi or atm.) at 20 °C for 1hr. It was filtrate and concentrated under reduced pressure to give a residue. The residue was purified by prep-HPLC (column: HUAPU 1010-X Amide 100*30mm*10um; mobile phase: [$H_2O$ (10mM $NH_4HCO_3$)-ACN]; gradient:95%-68% B over 14.0 min) to afford 2-[[(2S)-2,5-diamino-5-oxo-pentanoyl] amino] ethanesulfonic acid (4.6 mg, 18.16 μmol, 19.54% yield, 99%< purity) as a white solid.

SFC-MS: ($M+H^+$): 254.1 @ 1.837 min (IG_MeOH_5IPAm_10_50_34_35_4min)

LCMS: ($M+H^+$): 254.1 @ 1.861 min, (T3_0_30AB_10min_ELSD)

$^1$H NMR: (400 MHz, DEUTERIUM OXIDE) δ 3.97 (br t, $J$ = 6.4 Hz, 1H), 3.51 (br d, $J$ = 2.6 Hz, 2H), 3.02 (br t, $J$ = 6.2 Hz, 2H), 2.44 - 2.29 (m, 2H), 2.19 - 2.02 (m, 2H)

$^{13}$C NMR: (100 MHz, DEUTERIUM OXIDE) δ 173.81, 171.47, 52.26, 49.44, 35.15, 30.77, 26.48

**Synthesis of HKMS-0200**

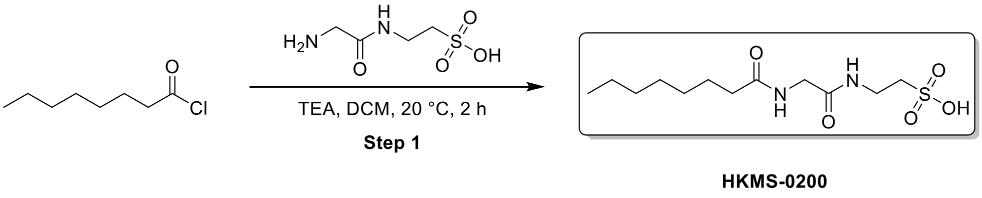


**2-[[2-(octanoylamino)acetyl]amino]ethanesulfonic acid**

To a solution of 2-[(2-aminoacetyl)amino]ethanesulfonic acid (50 mg, 274.43 μmol, 1 *eq*) in DCM (1 mL) was added TEA (55.54 mg, 548.85 μmol, 76.4 μL, 2 *eq*) and octanoyl chloride (44.64 mg, 274.43 μmol, 46.8 μL, 1 *eq*). The mixture was stirred at 20 °C for 2 h. The reaction mixture was concentrated under reduced pressure to remove solvent. The residue was purified by prep-HPLC(neutral condition). (column: Waters Xbridge BEH C18 100*30mm*10um;mobile phase: [$H_2O$(10mM $NH_4HCO_3$)-ACN];gradient:10%-40% B over 15.0 min) and compound 2-[[2-(octanoylamino)acetyl]amino]ethanesulfonic acid (1.7 mg, 5.29 μmol, 1.93% yield, 95.97% purity) was obtained as a white solid

LCMS: (M+1):309.1 @ 1.623min (5-95 % ACN in $H_2O$, 6 min)

$^1$H NMR: (400 MHz, DEUTERIUM OXIDE) δ 3.84 (s, 2H), 3.57 (t, *J* = 6.7 Hz, 2H), 3.05 (t, *J* = 6.7 Hz, 2H), 2.29 (t, *J* = 7.4 Hz, 2H), 1.57 (br t, *J* = 7.0 Hz, 2H), 1.31-1.20 (m, 8H), 0.90-0.76 (m, 3H)

$^{13}$C NMR: (100 MHz, DEUTERIUM OXIDE) δ 178.06, 171.38, 49.45, 42.50, 35.46, 34.99, 30.89, 28.14, 28.03, 25.06, 21.90, 13.32

**Synthesis of HKMS-0206**

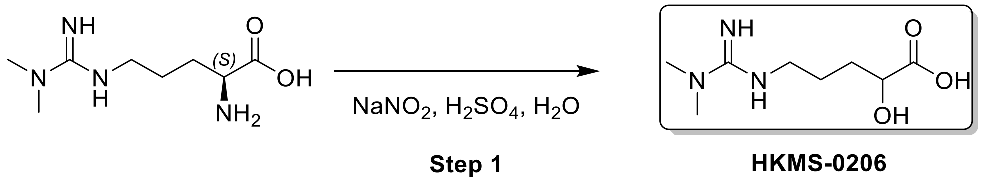


**(2S)-5-[(N,N-dimethylcarbamimidoyl)amino]-2-hydroxy-pentanoic acid**

To a solution of (2S)-2-amino-5-[(N,N-dimethylcarbamimidoyl)amino]pentanoic acid (150 mg, 741.64 μmol, 1 *eq*) in $H_2O$ (1.5 mL) was added $H_2SO_4$ (352.21 mg, 3.41 mmol, 191.4 μL, 95% purity, 4.6 *eq*) and $NaNO_2$ (235.38 mg, 3.41 mmol, 4.6 *eq*) at 0°C. The mixture was stirred at 20 °C for 12 h. The reaction mixture was filtered and concentrated under reduced pressure to give a residue. The residue was purified by prep-HPLC(FA condition). column: Kromasil C18(w) 250*50mm 10u;mobile phase: [$H_2O$(0.2% FA)-ACN];gradient:1%-15% B over 10.0 min. Compound (2S)-5-[(N,N-dimethylcarbamimidoyl)amino]-2-hydroxy-pentanoic acid (15 mg, 54.68 μmol, 7.37% yield, 90.86% purity, FA) was obtained as a yellow solid.

LCMS: (M+1):204.1 @ 4.725min (0-30 % ACN in $H_2O$, 10 min)

$^1$H NMR: (400 MHz, DEUTERIUM OXIDE) δ 4.04 (t, *J* = 5.2 Hz, 1H), 3.23 (t, *J* = 6.5 Hz, 2H), 2.97 (s, 6H), 1.76-1.55 (m, 4H)

$^{13}$C NMR: (100 MHz, DEUTERIUM OXIDE) δ 180.89, 155.94, 71.54, 54.47, 41.65, 39.68, 37.48, 35.71, 30.75, 27.64, 25.30, 23.99

#### Synthesis of HKMS-0210

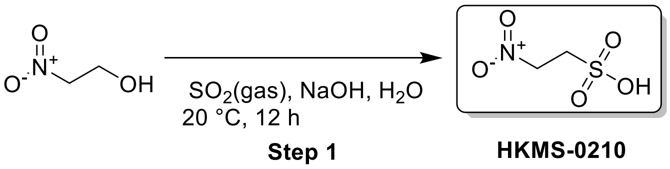


##### 2-nitroethanesulfonic acid

To a solution of NaOH (219.61 mg, 5.49 mmol, 1 *eq*) in $H_2O$ (2 mL) was added $SO_2$ (351.74 mg, 5.49 mmol, 1 *eq*) (15 Psi) and2-nitroethanol (0.5 g, 5.49 mmol, 387.0 µL, 1 *eq*). The mixture was stirred at 20 °C for 12 h. The reaction mixture was concentrated under reduced pressure to remove solvent. The residue was purified by prep-HPLC (column: HUAPU 1010-X Amide 100*30mm*10um;mobile phase: [$H_2O$(10mM $NH_4HCO_3$)-ACN];B%:95%, isocratic elution mode ) Compound 2-nitroethanesulfonic acid (2.6 mg, 16.39 µmol, 0.3% yield, 97.77% purity) was obtained as a white solid.

LCMS: (M-1):154.1 @ 1.016min (90-50 % ACN in $H_2O$, 10 min)

$^{1}$H NMR: (400 MHz, DEUTERIUM OXIDE) δ 4.88-4.83 (m, 2H), 3.62-3.55 (m, 2H)

$^{13}$C NMR: (100 MHz, DEUTERIUM OXIDE) δ 70.70, 47.49

#### Synthesis of HKMS-0215

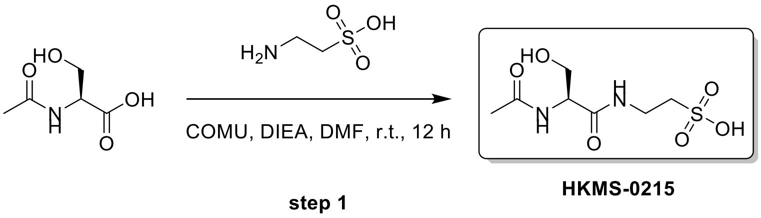


##### (S)-2-(2-acetamido-3-hydroxypropanamido) ethane-1-sulfonic acid

To a solution of (2S)-2-acetamido-3-hydroxy-propanoic acid (0.50 g, 3.40 mmol, 1.0 eq) and 2-aminoethanesulfonic acid (850.59 mg, 6.80 mmol, 847.2 µL, 2.0 eq) in DMF (5 mL) was added COMU (1.46 g, 3.40 mmol, 1 eq) and 2,6-dimethylpyridine (1.46 g, 13.59 mmol, 1.58 mL, 4.0 eq). The mixture was stirred at 20 °C for 12hr. The reaction mixture was concentrated under reduced pressure to give a residue. The residue was purified by prep-HPLC (column: HUAPU 1010-X Amide 100*30mm*10um; mobile phase: [$H_2O$ (10mM $NH_4HCO_3$)-ACN]; gradient:97%-85% B over 15.0 min) to afford 2-[[(2S)-2-acetamido-3-hydroxy-propanoyl] amino] ethanesulfonic acid (1.1 mg, 4.28 µmol, 0.1% yield, 98.9% purity) as a white solid.

LCMS: $(M+H^+)$: 253.27 @ 6.377 min (97_80CD_10_HILIC_Amide_PH3_2)

$^1$H NMR: (400 MHz, DEUTERIUM OXIDE) δ 4.30 (s, 1H), 3.77 (dd, $J$ = 1.8, 5.1 Hz, 2H), 3.59 - 3.47 (m, 2H), 3.01 (t, $J$ = 6.7 Hz, 2H), 1.99 (s, 3H)

$^{13}$C NMR: (100 MHz, DEUTERIUM OXIDE) δ 174.62, 171.64, 61.12, 55.83, 49.50, 35.14, 21.82

**Synthesis of HKMS-0223**

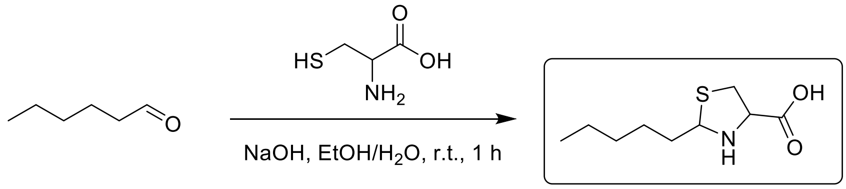


HKMS-0223

step 1

**2-pentylthiazolidine-4-carboxylic acid**

To a solution of hexanal (0.1 g, 998.42 μmol, 119.9 μL, 1.0 eq) and 2-amino-3-sulfanyl-propanoic acid (120.97 mg, 998.42 μmol, 1.0 eq) in EtOH (1 mL) and $H_2O$ (1 mL) was added NaOH (39.93 mg, 998.42 μmol, 1.0 eq). The mixture was stirred at 20 °C for 1hr. The reaction mixture was concentrated under reduced pressure to give a residue. The residue was purified by prep-HPLC (column: WePure Biotech XP tC18 100*30*7um; mobile phase: [$H_2O$ (10mM $NH_4HCO_3$)-ACN]; gradient:10%-40% B over 8.0 min) to afford 2-pentylthiazolidine-4-carboxylic acid (80.4 mg, 384.79 μmol, 38.5% yield, 97.3% purity) as a white solid.

LCMS: (M+H$^+$): 204.1 @ 2.497 min (0_60CD_6min_ELSD)

$^1$H NMR: (400 MHz, DEUTERIUM OXIDE) δ 4.77 (br s, 1H), 4.47 - 4.27 (m, 1H), 3.46 - 3.34 (m, 1H), 3.32 - 3.20 (m, 1H), 2.11 - 1.92 (m, 1H), 1.91 - 1.70 (m, 1H), 1.44 - 1.31 (m, 2H), 1.31 - 1.17 (m, 4H), 0.89 - 0.69 (m, 3H)

$^{13}$C NMR: (100 MHz, DMSO-$d_6$) δ 173.37, 172.83, 71.57, 70.83, 65.76, 64.65, 37.49, 37.19, 37.14, 35.29, 31.53, 31.49, 27.72, 27.52, 22.50, 22.44, 14.33, 14.33

**Synthesis of HKMS-0230**

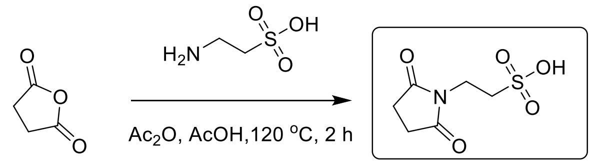


step 1

HKMS-0230

**2-(2,5-dioxopyrrolidin-1-yl) ethane-1-sulfonic acid**

To a solution of tetrahydrofuran-2,5-dione (0.1 g, 999.27 μmol, 1.0 eq) and 2-aminoethanesulfonic acid (93.79 mg, 749.46 μmol, 93.4 μL, 0.75 eq) in AcOH (0.6 mL) was added KOAc (98.07 mg, 999.27 μmol, 1.0 eq). The mixture was stirred at 120 °C for 2h. Then was added $Ac_2O$ (0.05 mL). The mixture was stirred at 120 °C for 2h. reaction mixture was filtered and the cake was dried under reduced pressure to afford the crude product (120 mg). The crude product (60 mg) residue was purified by prep-HPLC (column: Kromasil C18(w) 250*70mm 10u; mobile phase: [$H_2O$ (0.1% TFA)-ACN]; gradient:1%-6% B over 12.0 min) to afford 2-(2,5-dioxopyrrolidin-1-yl) ethanesulfonic acid (42.3 mg, 198.23 μmol, 19.84% yield, 97.1% purity) as a white solid.

LCMS: (M+H$^+$): 208.1 @ 2.601 min (T3_0_30AB_10min)

$^1$H NMR: (400 MHz, DEUTERIUM OXIDE) δ 3.81 (t, $J$ = 6.9 Hz, 2H), 3.08 (t, $J$ = 6.9 Hz, 2H), 2.70 (s, 4H)

$^{13}$C NMR: (100 MHz, DEUTERIUM OXIDE) δ 180.92, 46.84, 34.07, 27.98

**Synthesis of HKMS-0235**

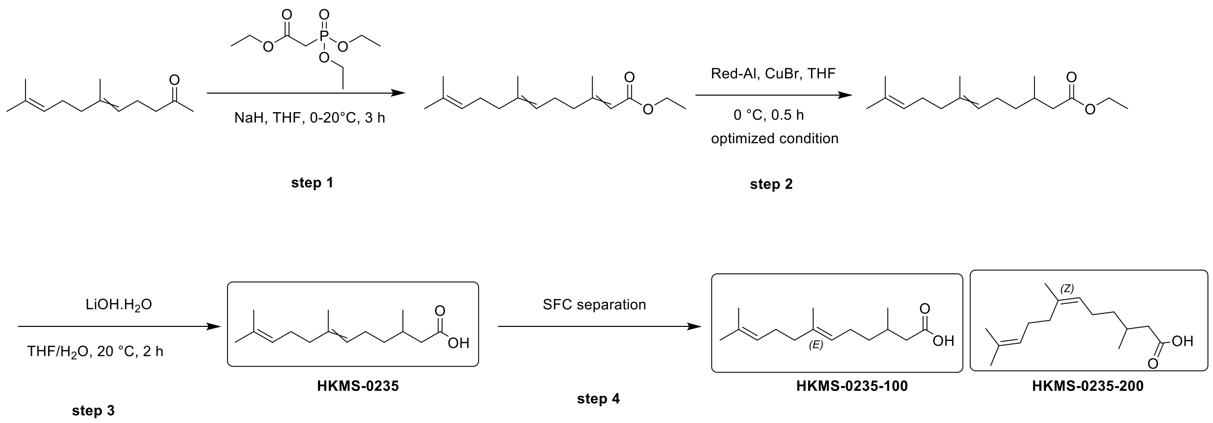


**Step 1 :**

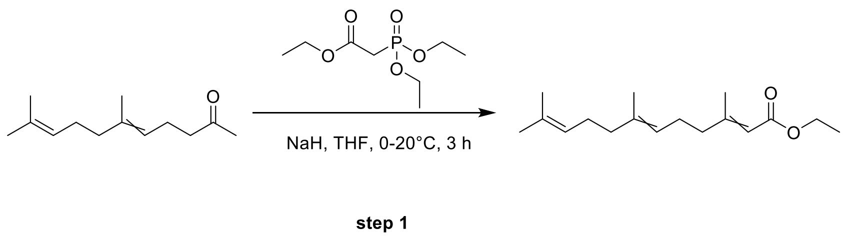


**Ethyl 3,7,11-trimethyldodeca-2,6,10-trienoate**

To a solution of ethyl 2-diethoxyphosphorylacetate (5 g, 22.30 mmol, 4.42 mL, 1.0 eq) in THF (50 mL) was added NaH (981.21 mg, 24.53 mmol, 60% purity, 1.1 eq) at 0 °C for 1h. Then was added (5E)-6,10-

dimethylundeca-5,9-dien-2-one (3.90 g, 20.07 mmol, 0.9 eq). The mixture was stirred at 20°C for 2 h. The reaction mixture was quenched with $NH_4Cl$ (aq., 50 mL) and water (50 mL). The mixture was extracted with EtOAc (50 mLx2) and the combined extracts were dried over $Na_2SO_4$, filtered and concentrated under reduced pressure to give a residue.

The residue was purified by column chromatography ($SiO_2$, Petroleum ether/Ethyl acetate=1/0 to 80/1) to afford ethyl (2E,6E)-3,7,11-trimethyldodeca-2,6,10-trienoate (3.9 g, 14.75 mmol, 66.14% yield) as a yellow oil.

$^1H$ NMR: (400 MHz, CHLOROFORM-d) δ 5.59 (s, 1H), 5.02 (br s, 2H), 4.18 - 3.96 (m, 2H), 2.13 - 2.06 (m, 5H), 2.03 - 1.86 (m, 4H), 1.61 (d, *J* = 2.5 Hz, 3H), 1.57 - 1.51 (m, 4H), 1.47 (s, 2H), 1.24 - 1.18 (m, 3H)

**Step 2**

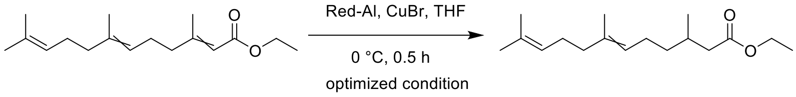


**step 2**

**Ethyl 3,7,11-trimethyldodeca-6,10-dienoate**

To a solution of ethyl (2E,6E)-3,7,11-trimethyldodeca-2,6,10-trienoate (1 g, 3.78 mmol, 1.0 eq) in THF (50 mL) was added CuBr (5.43 g, 37.82 mmol, 10.0 eq) and sodium; bis(2-methoxyethoxy) alumanylium;hydride (3.6 M, 8.40mL, 8 eq) at 0°C. The mixture was stirred at 0 °C for 0.5 h. The reaction mixture was quenched with water (50 mL). The mixture was extracted with EtOAc (50 mLx2) and the combined extracts were dried over $Na_2SO_4$, filtered and concentrated under reduced pressure to give a residue. The residue was purified by column chromatography ($SiO_2$,Petroleum ether/Ethyl acetate=1/0 to 40/1) to afford ethyl (6E)-3,7,11-trimethyldodeca-6,10-dienoate (0.39 g, 1.46 mmol, 38.70% yield) as a yellow oil.

**Step 3:**

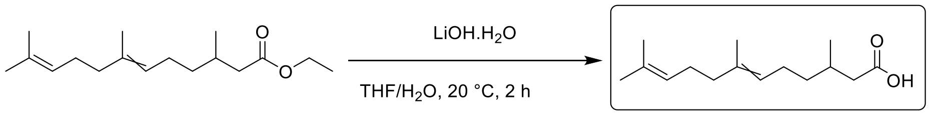


**HKMS-0235**

**step 3**

**3,7,11-trimethyldodeca-6,10-dienoic acid**

To a solution of ethyl (6E)-3,7,11-trimethyldodeca-6,10-dienoate (0.06 g, 225.21 μmol, 1 eq) in THF (0.5 mL) and $H_2O$ (0.1 mL) was added $LiOH.H_2O$ (18.90 mg, 450.42 μmol, 2 eq). The mixture was stirred at 20 °C for 2hr. The reaction mixture was poured into $H_2O$ (5 mL), and extracted with EtOAc(5 mLx3). Then the aqueous phase was adjusted pH to 2, and extracted with EtOAc(5 mLx3) The combined organic layers were washed with brine(5 mLx2), dried over $Na_2SO_4$, filtered and concentrated under reduced pressure to afford 3,7,11-trimethyldodec-10-enoic acid (10.9 mg, 44.35 μmol, 19.69% yield, 97.8% purity) as a yellow oil.

LCMS: (M-H$^-$): 239.2 @ 2.696 min, (5_95CD_6min_220)

$^1$H NMR: (400 MHz, METHANOL-$d_4$) δ 5.18 - 5.05 (m, 2H), 2.34 - 2.24 (m, 1H), 2.13 - 1.84 (m, 8H), 1.74 - 1.54 (m, 9H), 1.43 - 1.31 (m, 1H), 1.31 - 1.19 (m, 1H), 0.96 (d, *J* = 6.6 Hz, 3H)

$^{13}$C NMR: (100 MHz, METHANOL-$d_4$) δ 175.71, 134.83, 134.68, 130.95, 130.70, 124.80, 124.14, 124.01, 48.24, 47.81, 47.39, 47.60 (t, *J* = 42.9 Hz, 1C), 46.96, 41.18, 39.46, 36.78, 36.44, 31.48, 29.79, 29.66, 24.90, 24.46, 22.23, 18.62, 16.33, 14.62

**Step 4:**

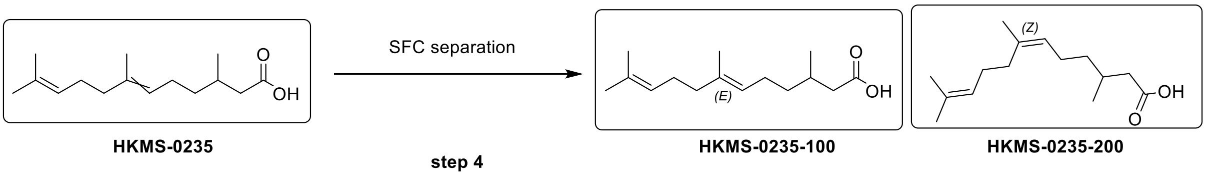


**(E)-3,7,11-trimethyldodeca-6,10-dienoic acid (HKMS-0235-100)**

The 3,7,11-trimethyldodec-10-enoic acid (0.04 g, 166.4 μmol, 1 eq) was purified by SFC purification (column: DAICEL CHIRALPAK AD (250mm*30mm,10um); mobile phase: [$CO_2$-IPA (0.1% $NH_3H_2O$)]; B%:13%, isocratic elution mode) to afford (6E)-3,7,11-trimethyldodeca-6,10-dienoic acid (15.6 mg, 63.13 μmol, 37.9% yield, 96.5% purity) as a colourless oil.

SFC: (M-H$^-$): 239.3 @ 1.310 min (AD_IPA_MNH3_10_50_25_35_5min)

LCMS: (M-H$^-$): 239.2 @ 2.695 min, (5_95CD_6min_220)

$^1$H NMR: (400 MHz, DMSO-$d_6$) δ 12.08 - 11.69 (m, 1H), 5.16 - 4.97 (m, 2H), 2.20 (dd, *J* = 5.9, 14.9 Hz, 1H), 2.09 - 1.88 (m, 7H), 1.87 - 1.77 (m, 1H), 1.63 (s, 3H), 1.56 (s, 6H), 1.38 - 1.25 (m, 1H), 1.23 - 1.10 (m, 1H), 0.88 (d, *J* = 6.6 Hz, 3H)

**(Z)-3,7,11-trimethyldodeca-6,10-dienoic acid (HKMS-0235-200)**

The 3,7,11-trimethyldodec-10-enoic acid (0.04 g, 166.4 μmol, 1 eq) was purified by SFC purification (column: DAICEL CHIRALPAK AD (250mm*30mm,10um); mobile phase: [$CO_2$-IPA (0.1% $NH_3H_2O$)]; B%:13%, isocratic elution mode) to afford (6Z)-3,7,11-trimethyldodeca-6,10-dienoic acid (8.5 mg, 34.56 μmol, 20.8% yield, 96.92% purity) as a colourless oil.

SFC: (M-$H^-$): 239.3 @ 1.158 min (AD_IPA_MNH3_10_50_25_35_5min)

LCMS: (M-$H^-$): 239.2 @ 2.667 min, (5_95CD_6min_220)

$^1$H NMR: (400 MHz, DMSO-$d_6$) δ 12.23 - 11.53 (m, 1H), 5.17 - 4.99 (m, 2H), 2.19 (dd, $J$ = 5.9, 14.9 Hz, 1H), 2.06 - 1.88 (m, 7H), 1.87 - 1.75 (m, 1H), 1.64 (s, 6H), 1.57 (s, 3H), 1.36 - 1.22 (m, 1H), 1.21 - 1.08 (m, 1H), 0.88 (d, $J$ = 6.7 Hz, 3H)

**Synthesis of HKMS-0238**

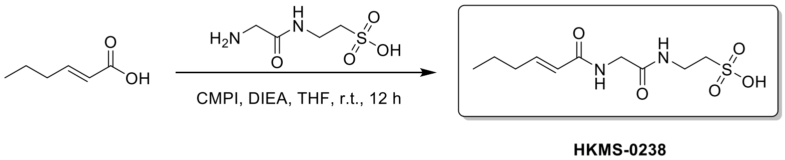


**2-[[2-[[(E)-hex-2-enoyl] amino] acetyl] amino] ethanesulfonic acid**

To a solution of (E)-hex-2-enoic acid (31.32 mg, 274.43 μmol, 1.0 eq) and 2-[(2-aminoacetyl) amino] ethanesulfonic acid (0.05 g, 274.43 μmol, 1.0 eq) in DMF (1 mL) was added CMPI (77.12 mg, 301.87 μmol, 1.1 eq) and DIEA (106.40 mg, 823.28 μmol, 143.40 μL, 3.0 eq). The mixture was stirred at 20 °C for 12 hr. The reaction mixture was concentrated under reduced pressure to give a residue. The residue was purified by prep-HPLC (column: WePure Biotech PHS XPt C18 100*30mm*7um; mobile phase: [$H_2O$ (0.05% TFA)-ACN]; gradient:1%-25% B over 12.0 min) to afford 2-[[2-[[(E)-hex-2-enoyl] amino] acetyl] amino] ethanesulfonic acid (37.6 mg, 135.09 μmol, 49.23% yield, 99%< purity) as a yellow solid.

LCMS: (M+$H^+$): 279.1 @ 5.321 min, (T3_0_60AB_10min)

$^1$H NMR: (400 MHz, DEUTERIUM OXIDE) δ 6.86 - 6.66 (m, 1H), 5.93 (d, $J$ = 15.6 Hz, 1H), 3.85 (s, 2H), 3.51 (t, $J$ = 6.6 Hz, 2H), 2.99 (t, $J$ = 6.8 Hz, 2H), 2.12 (q, $J$ = 7.1 Hz, 2H), 1.46 - 1.29 (m, 2H), 0.81 (t, $J$ = 7.4 Hz, 3H)

$^{13}$C NMR: (100 MHz, DEUTERIUM OXIDE) δ 173.50, 171.47, 169.64, 169.55, 147.72, 147.62, 121.95, 121.92, 49.46, 47.39, 42.58, 41.06, 35.34, 35.01, 33.57, 20.79, 18.47, 12.87

#### Synthesis of HKMS-0241

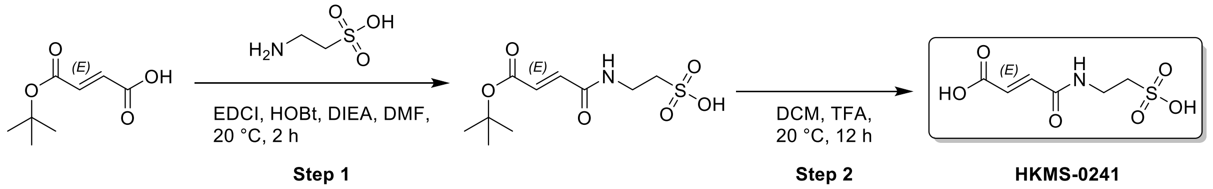


#### Step 1

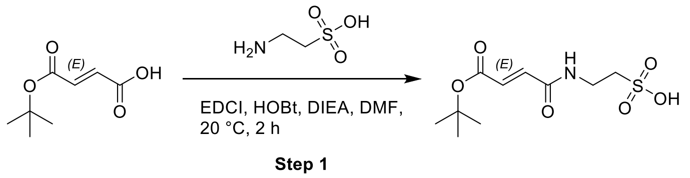


##### 2-[[(E)-4-tert-butoxy-4-oxo-but-2-enoyl]amino]ethanesulfonic acid

A mixture of (E)-4-tert-butoxy-4-oxo-but-2-enoic acid (100 mg, 580.79 µmol, 1 *eq*), 2-aminoethanesulfonic acid (72.68 mg, 580.79 µmol, 72.4 µL, 1 *eq*), EDCI (133.61 mg, 696.95 µmol, 1.2 *eq*), HOBt (102.02 mg, 755.03 µmol, 1.3 *eq*), DIEA (150.13 mg, 1.16 mmol, 202.33 µL, 2 *eq*) in DMF (1 mL) was degassed and purged with $N_2$ for 3 times, and then the mixture was stirred at 20 °C for 2 h under $N_2$ atmosphere. The reaction mixture was concentrated under reduced pressure to remove solvent. Compound 2-[[(E)-4-tert-butoxy-4-oxo-but-2-enoyl]amino]ethanesulfonic acid (160 mg, crude) was obtained as a yellow solid and used directly into the next step.

#### Step 2

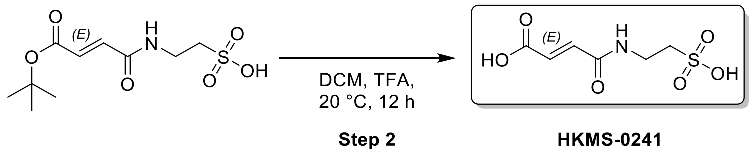


##### (E)-4-oxo-4-(2-sulfoethylamino)but-2-enoic acid

To a solution of 2-[[(E)-4-tert-butoxy-4-oxo-but-2-enoyl]amino]ethanesulfonic acid (160 mg, 572.84 µmol, 1 *eq*) in DCM (2 mL) was added TFA (0.6 mL). The mixture was stirred at 20 °C for 12 h. The reaction mixture was concentrated under reduced pressure to remove solvent. The residue was purified by prep-HPLC (HCl condition). (column: Kromasil C18(W) 150*30*10;mobile phase: [$H_2O$(0.05% HCl)-ACN];gradient:1%-10% B over 12.0 min ) Compound (E)-4-oxo-4-(2-sulfoethylamino)but-2-enoic acid (40.4 mg, 154.06 µmol, 26.89% yield, 99.02% purity, HCl) was obtained as a yellow solid.

LCMS: (M+1):224.0 @ 3.585min (0-30 % ACN in $H_2O$, 10 min)

[1]H NMR: (400 MHz, DEUTERIUM OXIDE) δ 8.47 (br s, 1H), 6.86 (br d, $J$ = 15.5 Hz, 1H), 6.48 (br d, $J$ = 15.5 Hz, 1H), 3.40 (br d, $J$ = 6.6 Hz, 2H), 2.59-2.54 (m, 2H)

[13]C NMR: (100 MHz, DEUTERIUM OXIDE) δ 169.13, 168.62, 165.52, 139.80, 136.65, 132.04, 52.98, 50.24, 42.80, 42.59, 42.38, 41.96, 42.17 (br t, $J$ = 42.0 Hz, 1C), 41.55, 38.57

**Synthesis of HKMS-0245**

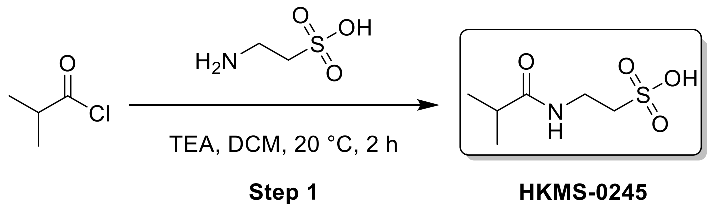


**2-(2-methylpropanoylamino)ethanesulfonic acid**

To a solution of 2-aminoethanesulfonic acid (300 mg, 2.40 mmol, 298.8 µL, 1 *eq*), TEA (485.14 mg, 4.79 mmol, 667.3 µL, 2 *eq*) in DCM (3 mL) was added2-methylpropanoyl chloride (255.42 mg, 2.40 mmol, 251.1 µL, 1 *eq*). The mixture was stirred at 20 °C for 2 h. The reaction mixture was concentrated under reduced pressure to remove solvent. The residue was purified by prep-HPLC(HCl condition). (column: WePure Biotech PHS XPt C18 100*30mm*7um;mobile phase: [$H_2O$(0.05% HCl)-ACN];gradient:1%-20% B over 12.0 min ) Compound 2-(2-methylpropanoylamino)ethanesulfonic acid (100.5 mg, 514.76 µmol, 21.47% yield, 99%< purity) was obtained as a white solid.

LCMS: (M+1):196.0 @ 3.924min (0-30 % ACN in $H_2O$, 10 min)

[1]H NMR: (400 MHz, DEUTERIUM OXIDE) δ 3.52 (t, $J$ = 6.6 Hz, 2H), 3.08-2.98 (m, 2H), 2.44 (td, $J$ = 6.9, 13.7 Hz, 1H), 1.05 (dd, $J$ = 0.8, 6.9 Hz, 6H)

[13]C NMR: (100 MHz, DEUTERIUM OXIDE) δ 180.92, 49.61, 34.97, 34.90, 18.44

**Synthesis of HKMS-0257**

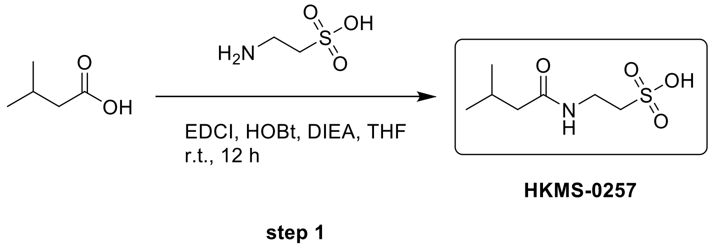


**2-(3-methylbutanamido) ethane-1-sulfonic acid**

To a solution of 3-methylbutanoic acid (0.1 g, 979.13 µmol, 107.5 µL, 1.0 eq) and 2-aminoethanesulfonic acid (122.54 mg, 979.13 µmol, 122.0 µL, 1.0 eq) in THF (1 mL) was added HOBt (145.53 mg, 1.08 mmol, 1.1 eq) and EDCI (206.47 mg, 1.08 mmol, 1.1 eq), DIEA (379.64 mg, 2.94 mmol, 511.6 µL, 3.0 eq). The mixture was stirred at 20 °C for 12hr. The reaction mixture was concentrated under reduced pressure to give a residue. The residue was purified by prep-HPLC (column: Welch Utimate Hilic Amide 250*50mm*10um; mobile phase: [$H_2O$ (0.1% TFA)-ACN]; gradient:95%-50% B over 15.0 min) to afford 2-(3-methylbutanoylamino) ethanesulfonic acid (47.3 mg, 226.03 µmol, 23.08% yield, 99%< purity) as a white solid.

LCMS: ($M+H^+$): 210.1 @ 5.166 min (T3_0_30AB_10min_ELSD)

$^1$H NMR: (400 MHz, DEUTERIUM OXIDE) δ 3.49 (t, $J$ = 6.8 Hz, 2H), 3.00 (t, $J$ = 6.9 Hz, 2H), 2.03 (d, $J$ = 7.4 Hz, 2H), 1.96 - 1.82 (m, 1H), 0.83 (d, $J$ = 6.6 Hz, 6H)

$^{13}$C NMR: (100 MHz, DEUTERIUM OXIDE) δ 176.49, 49.75, 44.96, 34.96, 26.03, 21.46

**Synthesis of HKMS-0260**

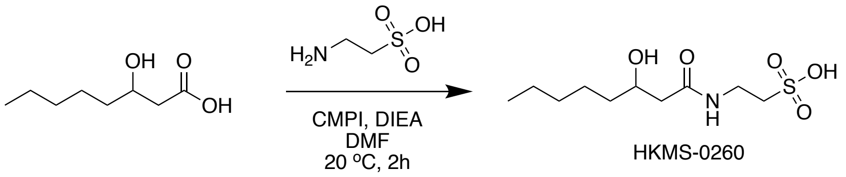


**2-(3-hydroxyoctanoylamino)ethanesulfonic acid**

A mixture of 3-hydroxyoctanoic acid (150 mg, 936.27 µmol, 1 *eq*), 2-aminoethanesulfonic acid (117.17 mg, 936.27 µmol, 116.7 µL, 1 *eq*), CMPI (358.80 mg, 1.40 mmol, 1.5 *eq*), DIEA (363.02 mg, 2.81 mmol, 489.2 µL, 3 *eq*) in DMF (2 mL) was degassed and purged with $N_2$ for 3 times, and then the mixture was stirred at 20 °C for 2 h under $N_2$ atmosphere. The reaction mixture was filtered and concentrated under reduced pressure to give a residue. The residue was purified by prep-HPLC(TFA condition). (column: WePure Biotech PHS XPt C18 100*30mm*7um;mobile phase: [$H_2O$(0.1% TFA)-ACN];gradient:1%-40% B over 14.0 min ) Compound 2-(3-hydroxyoctanoylamino)ethanesulfonic acid (32.9 mg, 86.27 µmol, 8.38% yield, 99%< purity, TFA) was obtained as a white solid.

LCMS: (M+1):268.2 @ 5.778min (0-30 % ACN in $H_2O$, 10 min)

$^1$H NMR: (400 MHz, DEUTERIUM OXIDE) δ 4.04-3.91 (m, 1H), 3.63-3.46 (m, 2H), 3.05 (t, $J$ = 6.8 Hz, 2H), 2.48-2.27 (m, 2H), 1.51-1.41 (m, 2H), 1.37-1.19 (m, 6H), 0.94-0.75 (m, 3H)

$^{13}$C NMR: (100 MHz, DEUTERIUM OXIDE) δ 174.20, 68.72, 49.70, 43.43, 36.00, 35.06, 30.87, 24.30,

21.85, 13.28

**Synthesis of HKMS-0263**

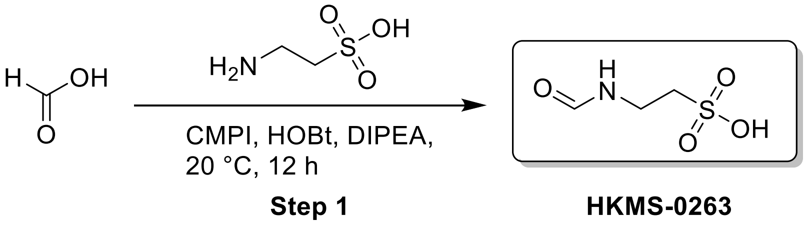


**2-formamidoethanesulfonic acid**

A mixture of formic acid (551.66 mg, 11.99 mmol, 452.2 µL, 3 *eq*), 2-aminoethanesulfonic acid (500 mg, 4.00 mmol, 498.0 µL, 1 *eq*), CMPI (1.53 g, 5.99 mmol, 1.5 *eq*), DIEA (1.03 g, 7.99 mmol, 1.39 mL, 2 *eq*) in DMF (5 mL) was degassed and purged with $N_2$ for 3 times, and then the mixture was stirred at 20 °C for 12 h under $N_2$ atmosphere. The reaction mixture was filtered and concentrated under reduced pressure to give a residue. The residue was purified by prep-HPLC(neutral condition). (column: Welch Utimate Hilic 100*25mm;mobile phase: [$H_2O$(10mM $NH_4HCO_3$)-ACN];gradient:95%-50% B over 15.0 min ) Compound 2-formamidoethanesulfonic acid (122 mg, 796.6 µmol, 19.94% yield, >99%purity) was obtained as a white solid. (DIEA salt)

LCMS: (M+1):154.1 @ 3.41min (90-50 % ACN in $H_2O$, 10 min)

$^1$H NMR: (400 MHz, DEUTERIUM OXIDE) δ 8.05 (s, 1H), 3.62 (t, *J* = 6.7 Hz, 2H), 3.11 (t, *J* = 6.7 Hz, 2H)

$^{13}$C NMR: (100 MHz, DEUTERIUM OXIDE) δ 164.30, 49.69, 42.61

**Synthesis of HKMS-0273**

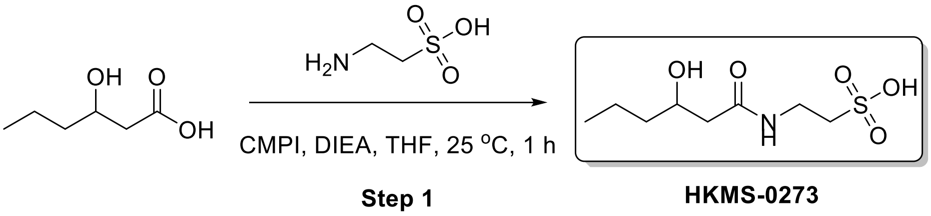


**2-(3-hydroxyhexanoylamino)ethanesulfonic acid**

A mixture of 3-hydroxyhexanoic acid (30 mg, 227.0 µmol, 1 eq), 2-aminoethanesulfonic acid (28.41 mg, 227.0 µmol, 28.3 µL, 1 eq), CMPI (86.99 mg, 340.5 µmol, 1.5 eq), DIEA (58.68 mg, 454.00 µmol, 79.1 µL, 2 eq) in THF (1 mL) was degassed and purged with $N_2$ for 3 times, and then the mixture was stirred at 25 °C for 1 h under $N_2$ atmosphere. The reaction mixture was concentrated under reduced pressure to remove solvent. The residue was purified by prep-HPLC (neutral condition)(column: WePure Biotech XPt C18150*40*7um;mobile phase: [$H_2O$(10mM $NH_4HCO_3$)-ACN];gradient:1%-20% B over 8.0 min) to give desired compound (10 mg, purity 88%) as a colorless oil, which was further separated by prep-HPLC(TFA condition)(column: Kromasil C18(W) 150*30*10; mobile phase: [$H_2O$(0.1% TFA)-ACN]; gradient:1%-15% B over 12.0 min). Compound 2-(3-hydroxyhexanoylamino)ethane sulfonic acid (4.7 mg, 19.64 μmol, 5.21% yield, 99%< purity) was obtained as a colorless oil.

LCMS: ($M+H^+$): 240.2 @ 4.134 min (T3-0-60 % ACN in $H_2O$, 10 min)

$^1$H NMR: (400 MHz, DEUTERIUM OXIDE) δ 4.05-3.94 (m, 1H), 3.62-3.50 (m, 2H), 3.06 (t, $J$ = 6.8 Hz, 2H), 2.47-2.27 (m, 2H), 1.50-1.23 (m, 4H), 0.91-0.83 (m, 3H)

$^{13}$C NMR: (100 MHz, DEUTERIUM OXIDE) δ 174.20, 68.38, 49.66, 43.43, 38.25, 35.04, 18.07, 13.11

**Synthesis of HKMS-0281**

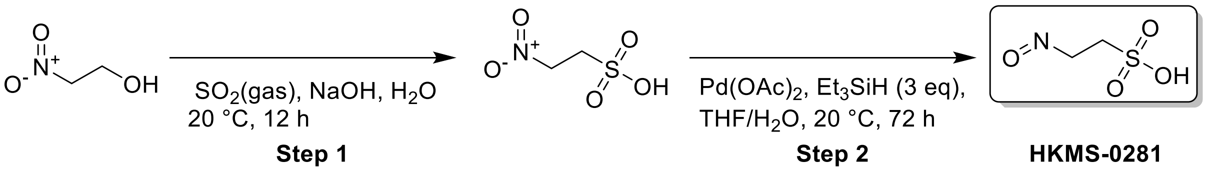


Step 1:

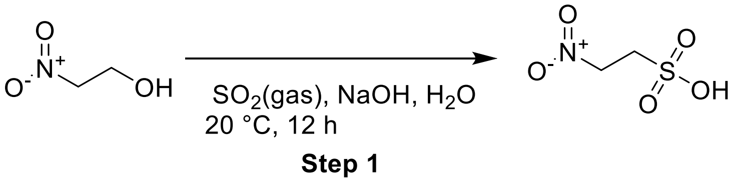


**2-nitroethanesulfonic acid**

To a solution of NaOH (2.20 g, 54.91 mmol, 1 *eq*) in $H_2O$ (20 mL) was added $SO_2$ (3.52 g, 54.91 mmol, 1 *eq*) (15 Psi) and2-nitroethanol (5 g, 54.91 mmol, 3.87 mL, 1 *eq*). The mixture was stirred at 20 °C for 12 h. The reaction mixture was concentrated under reduced pressure to remove solvent. The residue was purified by prep-HPLC(neutral condition). (column: Welch Utimate Hilic Amide 100*25mm*10um;mobile phase: [$H_2O$(10mM $NH_4HCO_3$)-ACN];gradient:95%-50% B over 15.0 min) Compound 2-nitroethanesulfonic acid (250 mg, 1.61 mmol, 2.94% yield) was obtained as a white solid and used into the next step.

$^{1}H$ NMR: (400 MHz, DEUTERIUM OXIDE) δ 4.88-4.83 (m, 2H), 3.63-3.56 (m, 2H)

**Step 2:**

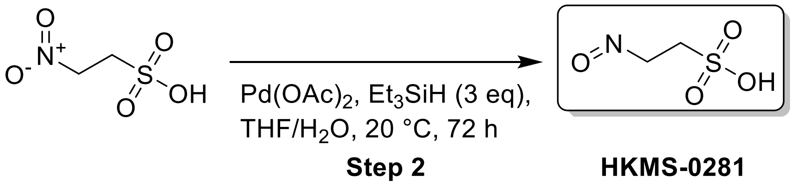


**2-nitrosoethanesulfonic acid**

A mixture of 2-nitroethanesulfonic acid (50 mg, 322.31 μmol, 1 *eq*), $Et_3SiH$ (112.43 mg, 966.93 μmol, 154.44 μL, 3 *eq*), $Pd(OAc)_2$ (3.62 mg, 16.12 μmol, 0.05 *eq*) in THF (2 mL) and $H_2O$ (0.5 mL) was degassed and purged with $N_2$ for 3 times, and then the mixture was stirred at 20 °C for 4 h under $N_2$ atmosphere. The reaction mixture was filtered and concentrated under reduced pressure to give a residue. The residue was purified by prep-HPLC(neutral condition). (column: Welch Utimate Hilic Amide 100*25mm*10um;mobile phase: [$H_2O$(10mM $NH_4HCO_3$)-ACN];gradient:95%-60% B over 15.0 min ) Compound 2-nitrosoethanesulfonic acid (2.7 mg, 14.50 μmol, 4.50% yield, 74.74% purity) was obtained as a pink solid.

LCMS: ET100754-306-P1B (M-1):138.2 @ 2.725min (90-50 % ACN in $H_2O$, 10 min)

$^{1}H$ NMR: ET100754-306-P1H (400 MHz, DEUTERIUM OXIDE) δ 4.04 (d, $J$ = 5.8 Hz, 2H), 3.79 (d, $J$ = 6.6 Hz, 2H)

13 C NMR: (100 MHz, DEUTERIUM OXIDE) δ 144.59, 142.90, 51.30, 46.75

**Synthesis of HKMS-0242-200**

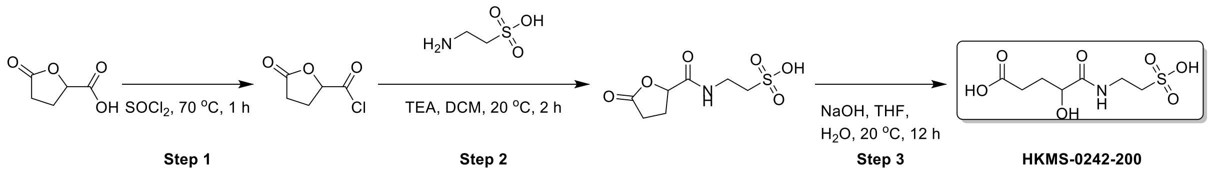


**Step 1 :**

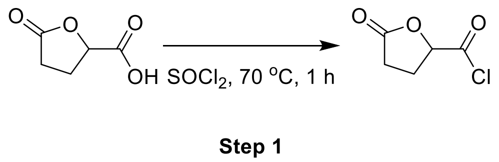


**5-oxotetrahydrofuran-2-carbonyl chloride**

A mixture of 5-oxotetrahydrofuran-2-carboxylic acid (2 g, 15.37 mmol, 1 *eq*) in $SOCl_2$ (20 mL) was degassed and purged with $N_2$ for 3 times, and then the mixture was stirred at 70 °C for 1 h under $N_2$ atmosphere. The reaction mixture was concentrated under reduced pressure to remove solvent. Compound 5-oxotetrahydrofuran-2-carbonyl chloride (2.28 g, crude) was obtained as a yellow oil and used into the next step.

**Step 2 :**

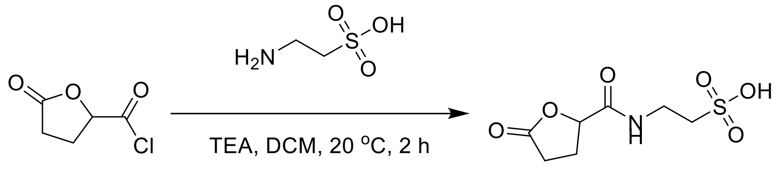


**Step 2**

**2-[(5-oxotetrahydrofuran-2-carbonyl)amino]ethanesulfonic acid**

To a solution of 2-aminoethanesulfonic acid (750 mg, 5.99 mmol, 747.01 µL, 1 *eq*), TEA (667.07 mg, 6.59 mmol, 917.57 µL, 1.1 *eq*) in DCM (10 mL) was added5-oxotetrahydrofuran-2-carbonyl chloride (2.23 g, 14.98 mmol, 2.5 *eq*). The mixture was stirred at 20 °C for 2 h. The reaction mixture was concentrated under reduced pressure to give a residue. The crude product 2-[(5-oxotetrahydrofuran-2-carbonyl)amino]ethanesulfonic acid (4 g, crude) was ontained as a black solid and used into the next step without further purification.

**Step 3:**

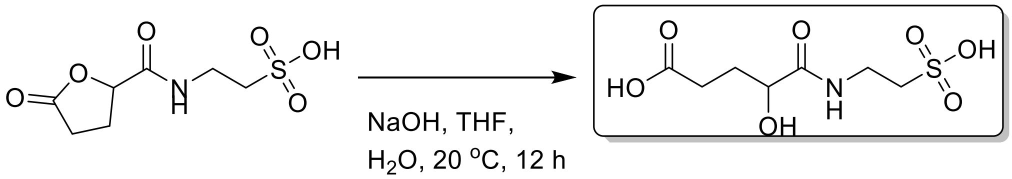


**Step 3** **HKMS-0242-200**

**4-hydroxy-5-oxo-5-(2-sulfoethylamino)pentanoic acid**

To a solution of 2-[(5-oxotetrahydrofuran-2-carbonyl)amino]ethanesulfonic acid (1.4 g, 5.90 mmol, 1 *eq*) in THF (10 mL) was added NaOH (259.64 mg, 6.49 mmol, 1.1 *eq*) and $H_2O$ (2 mL). The mixture was stirred at 20 °C for 12 h. The reaction mixture was concentrated under reduced pressure to remove solvent. The residue was purified by prep-HPLC(neutral condition). (column: Welch Utimate Hilic Amide 100*25mm*10um;mobile phase: [$H_2O$(10mM $NH_4HCO_3$)-ACN];gradient:95%-65% B over 15.0 min) Compound 4-hydroxy-5-oxo-5-(2-sulfoethylamino)pentanoic acid (34.6 mg, 135.56 µmol,

2.30% yield, 99%< purity) was obtained as a white solid.

LCMS: (M+1):256.1 @ 2.202min (0-30 % ACN in $H_2O$, 10 min)

[1]H NMR: (400 MHz, DEUTERIUM OXIDE) δ 4.12 (dd, *J* = 3.9, 8.2 Hz, 1H), 3.60 (dt, *J* = 1.8, 6.8 Hz, 2H), 3.08 (t, *J* = 6.8 Hz, 2H), 2.32-2.25 (m, 2H), 2.03 (dtd, *J* = 3.9, 7.9, 14.3 Hz, 1H), 1.87-1.74 (m, 1H)

13 C NMR: (100 MHz, DEUTERIUM OXIDE) δ 180.54, 176.23, 78.08, 70.86, 49.59, 49.30, 34.93, 34.71, 31.67, 29.53, 27.44, 25.45

**Synthesis of HKMS-0277**

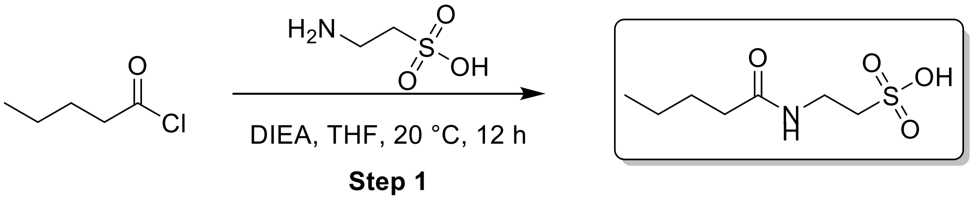


**HKMS-0277**

**2-(pentanoylamino)ethanesulfonic acid**

To a solution of 2-aminoethanesulfonic acid (1 g, 7.99 mmol, 996.02 µL, 1 *eq*) in THF (10 mL) was added DIPEA (1.14 g, 8.79 mmol, 1.53 mL, 1.1 *eq*) and pentanoyl chloride (963.49 mg, 7.99 mmol, 968.33 µL, 1 *eq*). The mixture was stirred at 20 °C for 12 h. The reaction mixture was filtered and concentrated under reduced pressure to give a residue. The residue was purified by prep-HPLC(TFA condition). (column: WePure Biotech XPt C18 100*30mm;mobile phase: [$H_2O$(0.1% TFA)-ACN];gradient:5%-30% B over 12.0 min ) Compound 2-(pentanoylamino)ethanesulfonic acid (38.7 mg, 178.37 µmol, 2.23% yield, 96.45% purity) was obtained as a white oil.

LCMS: (M+1):210.1 @ 0.564min (0-60 % ACN in $H_2O$, 6 min)

[1]H NMR: (400 MHz, DEUTERIUM OXIDE) δ 3.46 (t, *J* = 6.8 Hz, 2H), 2.97 (t, *J* = 6.8 Hz, 2H), 2.15 (t, *J* = 7.4 Hz, 2H), 1.45 (quin, *J* = 7.4 Hz, 2H), 1.20 (sxt, *J* = 7.4 Hz, 2H), 0.77 (t, *J* = 7.4 Hz, 3H)

13 C NMR: (100 MHz, DEUTERIUM OXIDE) δ 177.29, 49.70, 35.48, 35.01, 27.40, 21.51, 12.96

**Synthesis of HKMS-0284**

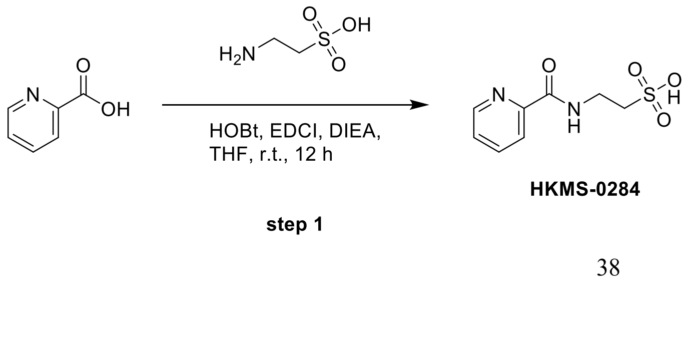

**2-(picolinamido) ethane-1-sulfonic acid**

To a solution of 2-aminoethanesulfonic acid (203.31 mg, 1.62 mmol, 202.50 μL, 1.0 eq) and pyridine-2-carboxylic acid (0.2 g, 1.62 mmol, 1.0 eq) in THF (3 mL) was added DIEA (629.89 mg, 4.87 mmol, 848.91 μL, 3.0 eq), HOBt (241.46 mg, 1.79 mmol, 1.1 eq), EDCI (342.57 mg, 1.79 mmol, 1.1 eq). The mixture was stirred at 20 °C for 12hr. The reaction mixture was concentrated under reduced pressure to give a residue. The residue was purified by prep-HPLC (column: Welch Utimate Hilic Amide 100*25mm*10um; mobile phase: [$H_2O$ (0.1% TFA)-ACN]; gradient:95%-50% B over 15.0 min) to 2-(pyridine-2-carbonylamino) ethanesulfonic acid (95.9 mg, 416.52 μmol, 25.64% yield, 99%< purity) as a yellow solid.

LCMS:($M+H^+$): 231.1 @ 4.454 min (T3_0_30AB_10min_220)

$^1$H NMR: (400 MHz, DEUTERIUM OXIDE) δ = 8.74 (d, *J* = 5.4 Hz, 1H), 8.43 (dt, *J* = 1.4, 7.9 Hz, 1H), 8.24 (d, *J* = 8.0 Hz, 1H), 8.07 - 7.86 (m, 1H), 3.79 (t, *J* = 6.7 Hz, 2H), 3.16 (t, *J* = 6.6 Hz, 2H)

$^{13}$C NMR: (100 MHz, DEUTERIUM OXIDE) δ = 161.11, 146.26, 143.71, 143.41, 129.26, 124.56, 49.16, 35.98

**Synthesis of HKMS-0275**

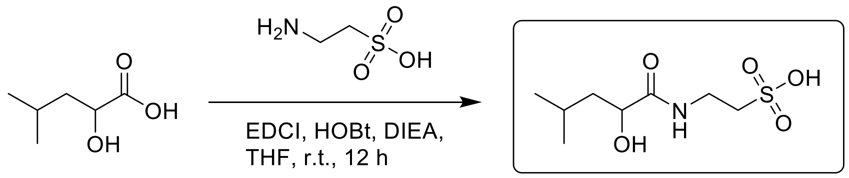


**HKMS-0275**

**2-(2-hydroxy-4-methylpentanamido) ethane-1-sulfonic acid**

To a solution of 2-hydroxy-4-methyl-pentanoic acid (0.2 g, 1.51 mmol, 1.0 eq) and 2-aminoethanesulfonic acid (189.39 mg, 1.51 mmol, 188.64 μL, 1.0 eq) in THF (3 mL) was added DIEA (586.77 mg, 4.54 mmol, 790.79 μL, 3.0 eq) and HOBT (224.94 mg, 1.66 mmol, 1.1 eq), EDCI (319.12 mg, 1.66 mmol, 1.1 eq). The mixture was stirred at 20 °C for 12hr. The reaction mixture was concentrated under reduced pressure to give a residue. The residue was purified by prep-HPLC (column: Welch Utimate Hilic Amide 100*25mm*10um; mobile phase: [$H_2O$ (0.1% TFA)-ACN]; gradient:95%-50% B over 15.0 min) to afford 2-[(2-hydroxy-4-methyl-pentanoyl) amino] ethanesulfonic acid (29.1 mg, 121.61 μmol, 8.04% yield, 99%< purity) as a white solid.

LCMS: ($M+H^+$): 240.2 @ 5.391 min (T3_0_30AB_10min_ELSD)

$^{1}H$ NMR: (400 MHz, DEUTERIUM OXIDE) δ = 4.08 (t, *J* = 6.7 Hz, 1H), 3.54 (t, *J* = 6.6 Hz, 2H), 3.01 (t, *J* = 6.7 Hz, 2H), 1.74 - 1.60 (m, 1H), 1.45 (t, *J* = 6.9 Hz, 2H), 0.83 (d, *J* = 6.7 Hz, 6H)

$^{13}C$ NMR: (100 MHz, DEUTERIUM OXIDE) δ = 177.45, 70.05, 49.59, 42.49, 34.72, 23.87, 22.61, 20.59

**Synthesis of HKMS-0282**

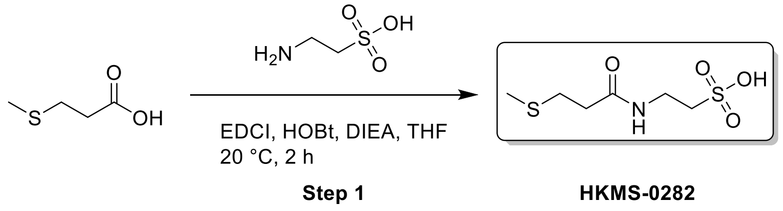


**2-(3-methylsulfanylpropanoylamino)ethanesulfonic acid**

A mixture of 3-methylsulfanylpropanoic acid (300 mg, 2.50 mmol, 1 *eq*), 2-aminoethanesulfonic acid (312.42 mg, 2.50 mmol, 311.18 µL, 1 *eq*), EDCI (574.29 mg, 3.00 mmol, 1.2 *eq*), HOBt (438.53 mg, 3.25 mmol, 1.3 *eq*) and DIPEA (483.97 mg, 3.74 mmol, 652.26 µL, 1.5 *eq*) in THF (5 mL) was degassed and purged with $N_2$ for 3 times, and then the mixture was stirred at 20 °C for 2 h under $N_2$ atmosphere. The reaction mixture was filtered and concentrated under reduced pressure to give a residue. The residue was purified by prep-HPLC(TFA condition). (column: Kromasil C18(w) 250*70mm 10u;mobile phase: [$H_2O$(0.1% TFA)-ACN];gradient:1%-15% B over 12.0 min ) Compound 2-(3-methylsulfanylpropanoylamino)ethanesulfonic acid (111.5 mg, 323.57 µmol, 12.96% yield, 99.05% purity, TFA) was obtained as a white solid.

LCMS: (M+1):228.1 @ 4.264min (0-30 % ACN in $H_2O$, 10 min)

$^{1}H$ NMR: (400 MHz, DEUTERIUM OXIDE) δ 3.53 (t, *J* = 6.8 Hz, 2H), 3.04 (t, *J* = 6.8 Hz, 2H), 2.76-2.68 (m, 2H), 2.55-2.47 (m, 2H), 2.06 (s, 3H)

13 C NMR: (100 MHz, DEUTERIUM OXIDE) δ 174.61, 49.69, 47.43, 35.25, 35.11, 33.67, 29.10, 28.22, 14.17

**Synthesis of HKMS-0283**

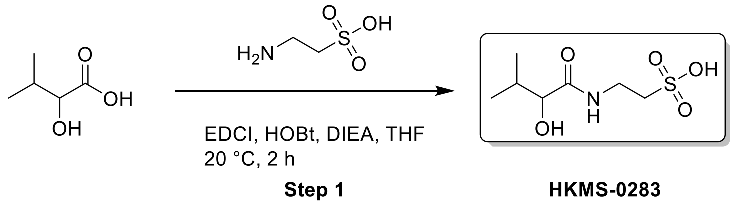


**2-[(2-hydroxy-3-methyl-butanoyl)amino]ethanesulfonic acid**

A mixture of 2-hydroxy-3-methyl-butanoic acid (500 mg, 4.23 mmol, 1 *eq*), 2-aminoethanesulfonic acid (529.70 mg, 4.23 mmol, 527.59 μL, 1 *eq*), EDCI (973.67 mg, 5.08 mmol, 1.2 *eq*), HOBt (743.50 mg, 5.50 mmol, 1.3 *eq*) and DIPEA (820.55 mg, 6.35 mmol, 1.11 mL, 1.5 *eq*) in THF (5 mL) was degassed and purged with $N_2$ for 3 times, and then the mixture was stirred at 20 °C for 2 h under $N_2$ atmosphere. The reaction mixture was filtered and concentrated under reduced pressure to give a residue. The residue was purified by prep-HPLC(TFA condition). (column: Kromasil C18(w) 250*70mm 10u;mobile phase: [$H_2O$(0.1% TFA)-ACN];gradient:1%-15% B over 12.0 min ) Compound 2-[(2-hydroxy-3-methyl-butanoyl)amino]ethanesulfonic acid (8.3 mg, 24.46 μmol, 5.78e-1% yield, 99%< purity, TFA) was obtained as a white solid.

LCMS: ET100754-282-P1B1 (M+1):226.2 @ 4.275min (0-30 % ACN in $H_2O$, 10 min)

$^1$H NMR: (400 MHz, DEUTERIUM OXIDE) δ 3.95 (d, $J$ = 3.9 Hz, 1H), 3.60 (t, $J$ = 6.7 Hz, 2H), 3.11-3.03 (m, 2H), 2.04 (dtd, $J$ = 3.9, 7.0, 13.8 Hz, 1H), 0.94 (d, $J$ = 7.0 Hz, 3H), 0.80 (d, $J$ = 6.8 Hz, 3H)

13 C NMR: (100 MHz, DEUTERIUM OXIDE) δ 176.28, 75.92, 49.63, 34.58, 31.08, 18.14, 15.18

**Synthesis of HKMS-0285**

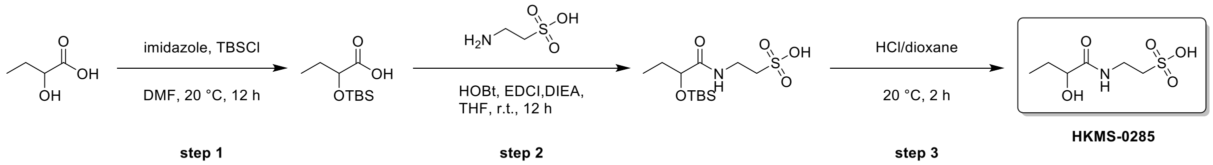


#### Step 1

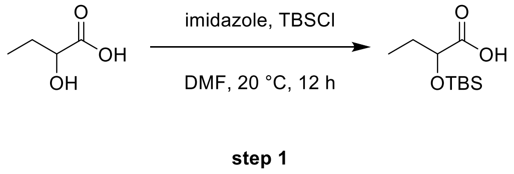


**2-((tert-butyldimethylsilyl)oxy)butanoic acid**

To a solution of 2-hydroxybutanoic acid (1 g, 9.61 mmol, 1.0 eq) in DMF (9 mL) was added IMIDAZOLE (1.96 g, 28.82 mmol, 3.0 eq) and TBSCl (4.34 g, 28.82 mmol, 3.55 mL, 3.0 eq). The mixture was stirred at 20 °C for 12hr. The reaction mixture was poured into $H_2O$ (30 mL), adjust PH to 3 and extracted with EtOAc(30 mLx3). The combined organic layers were washed with brine(30 mLx2), dried over $Na_2SO_4$, filtered and concentrated under reduced pressure to give 2-[tert-butyl(dimethyl)silyl]oxybutanoic acid (3.6 g, crude) as a yellow oil.

$^1$H NMR: (400 MHz, CHLOROFORM-d) δ 4.09 - 4.04 (m, 1H), 1.79 - 1.63 (m, 2H), 0.92 - 0.88 (m, 11H), 0.06 - -0.03 (m, 6H)

**Step 2**

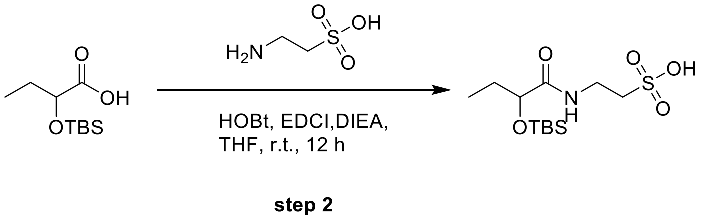


**2-(2-((tert-butyldimethylsilyl) oxy)butanamido) ethane-1-sulfonic acid**

To a solution of 2-[tert-butyl(dimethyl)silyl]oxybutanoic acid (1.5 g, 6.87 mmol, 1.0 eq) and 2-aminoethanesulfonic acid (859.66 mg, 6.87 mmol, 856.24 µL, 1.0 eq) in THF (10 mL) was added HOBt (1.02 g, 7.56 mmol, 1.1 eq) and EDCI (1.45 g, 7.56 mmol, 1.1 eq), DIEA (2.66 g, 20.61 mmol, 3.59 mL, 3.0 eq). The mixture was stirred at 20 °C for 12hr. The reaction mixture was concentrated under reduced pressure to give a residue. The residue was purified by prep-HPLC (column: Waters Xbridge BEH C18 100*30mm*10um; mobile phase: [$H_2O$ (10mM $NH_4HCO_3$)-ACN]; gradient:15%-45% B over 15.0 min) to afford 2-[2-[tert-butyl(dimethyl)silyl] oxybutanoylamino] ethanesulfonic acid (0.43 g, 1.32 mmol, 19.23% yield) as a white solid.

**Step 3**

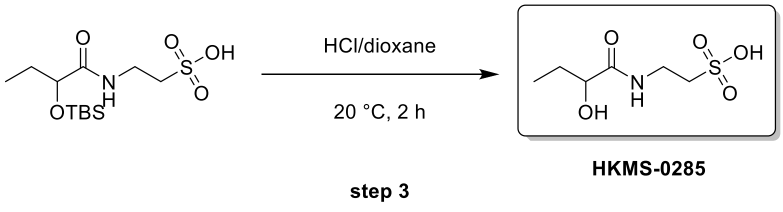


**2-(2-hydroxybutanamido)ethane-1-sulfonic acid**

To a solution of 2-[2-[tert-butyl(dimethyl)silyl] oxybutanoylamino] ethanesulfonic acid (0.2 g, 614.45 μmol, 1.0 eq) in HCl/dioxane (2 mL). The mixture was stirred at 20 °C for 2hr. The reaction mixture was concentrated under reduced pressure to give a residue. The residue was purified by prep-HPLC (column: Kromasil C18(w) 250*70mm 10u; mobile phase: [$H_2O$ (0.1% TFA)-ACN]; gradient:1%-8% B over 12.0 min) to afford 2-(2-hydroxybutanoylamino) ethanesulfonic acid (36.4 mg, 172.32 μmol, 28.04% yield, 99%< purity) as a colourless oil.

LCMS: (M+H$^+$): 245.1 @ 2.877 min, (T3_0_30AB_10min_ELSD)

$^1$H NMR: (400 MHz, DEUTERIUM OXIDE) δ 4.04 (dd, *J* = 4.3, 7.3 Hz, 1H), 3.57 (t, *J* = 6.7 Hz, 2H), 3.04 (t, *J* = 6.7 Hz, 2H), 1.81 - 1.67 (m, 1H), 1.58 (quind, *J* = 7.4, 14.5 Hz, 1H), 0.86 (t, *J* = 7.5 Hz, 3H)

$^{13}$C NMR: (100 MHz, DEUTERIUM OXIDE) δ 176.68, 72.45, 49.59, 34.63, 26.60, 8.38

**Synthesis of HKMS-0286**

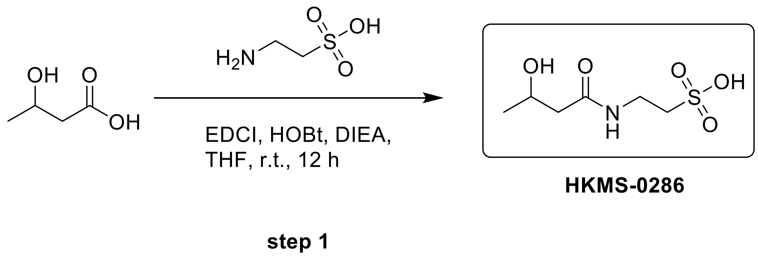


**2-(3-hydroxybutanamido) ethane-1-sulfonic acid**

To a solution of 3-hydroxybutanoic acid (0.1 g, 960.58 μmol, 88.81 μL, 1.0 eq) and 2-aminoethanesulfonic acid (120.21 mg, 960.58 μmol, 119.73 μL, 1.0 eq) in THF (2 mL) was added EDCI (202.56 mg, 1.06 mmol, 1.1 eq) and HOBt (142.78 mg, 1.06 mmol, 1.1 eq), DIEA (248.29 mg, 1.92 mmol, 334.63 μL, 2.0 eq). The mixture was stirred at 20 °C for 12hr. The reaction mixture was concentrated under reduced pressure to give a residue. The residue was purified by prep-HPLC (column: Kromasil C18(W) 150*30*10; mobile phase: [$H_2O$ (0.1% TFA)-ACN]; gradient:1%-5% B over 8.0 min) to afford 2-(3-hydroxybutanoylamino) ethanesulfonic acid (30 mg, 142.02 μmol, 14.79% yield, 99%< purity) as a yellow solid.

LCMS: (M+H$^+$): 212.1 @ 2.332 min (T3_0_30AB_10min_ELSD)

$^1$H NMR (400 MHz, DEUTERIUM OXIDE) δ = 4.10 (q, *J* = 6.4 Hz, 1H), 3.51 (dt, *J* = 1.4, 6.8 Hz, 2H), 3.02 (t, *J* = 6.8 Hz, 2H), 2.33 (d, *J* = 6.7 Hz, 2H), 1.13 (d, *J* = 6.3 Hz, 3H)

$^{13}$C NMR: (100 MHz, DEUTERIUM OXIDE) δ 173.87, 64.95, 49.60, 44.78, 35.02, 21.83

#### Synthesis of HKMS-0213

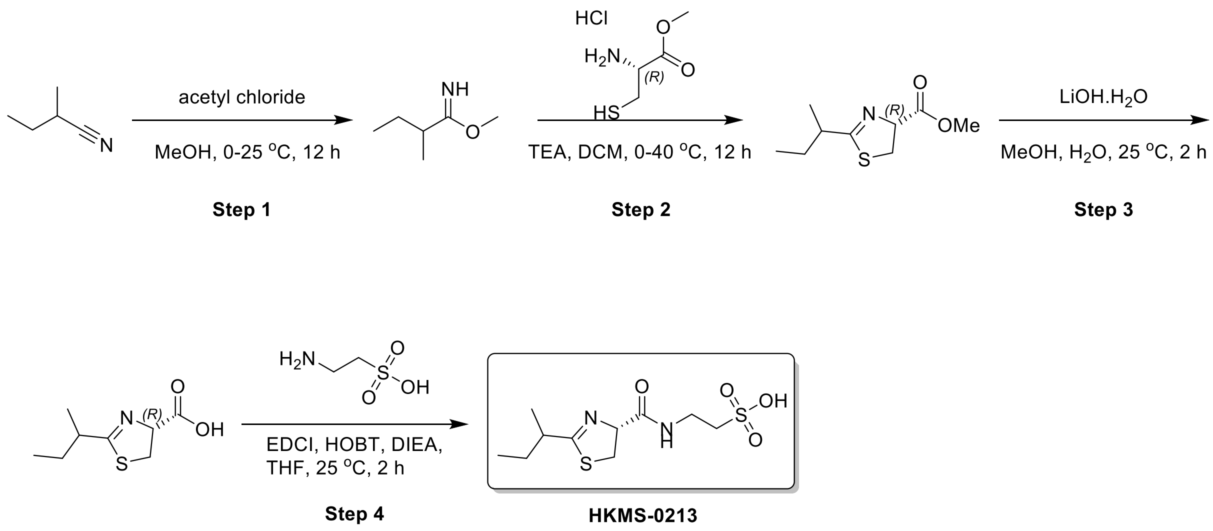


#### Step 1 (ET100716-220):

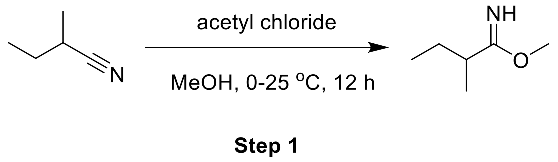


#### Methyl 2-methylbutanimidate

To a solution of 2-methylbutanenitrile (2 g, 24.06 mmol, 2.44 mL, 1 eq) in MeOH (20 mL) was added acetyl chloride (9.44 g, 120.29 mmol, 8.55 mL, 5 eq) at 0 °C. The mixture was stirred at 25 °C for 12 h. The reaction mixture was concentrated under reduced pressure to remove solvent. The crude product was purified by re-crystallization from MTBE (30 ml) at 25 °C. The mixture was collected by filtration and the solid was concentrated under reduced pressure to give a residue. Compound methyl 2-methylbutanimidate (3 g, crude) was obtained as a white solid. The crude product methyl 2-methylbutanimidate (3 g, crude) was used into the next step without further purification.

**$^1$H NMR:** (400 MHz, DMSO-d6) δ 4.11 (s, 3H), 2.79 (sxt, $J$ = 7.0 Hz, 1H), 1.69-1.45 (m, 2H), 1.17 (d, $J$ = 6.8 Hz, 3H), 0.86 (t, $J$ = 7.5 Hz, 3H)

#### Step 2

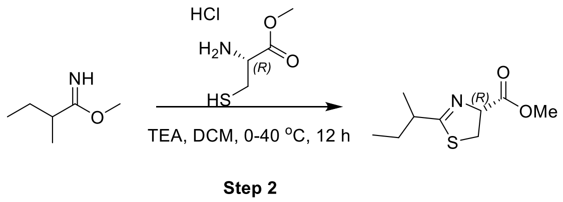

**Methyl (4R)-2-sec-butyl-4,5-dihydrothiazole-4-carboxylate**

To a solution of methyl 2-methylbutanimidate (1 g, 8.68 mmol, 1 eq) and methyl (2R)-2-amino-3-sulfanyl-propanoate;hydrochloride (1.49 g, 8.68 mmol, 1 eq) in DCM (10 mL) was added TEA (3.51 g, 34.73 mmol, 4.83 mL, 4 eq) at 0 °C. The mixture was stirred at 40 °C for 12 h. The mixture was poured into water (20 mL) and extracted with DCM (30 mL*3). The combined organic phase was washed with brine (30 mL), dried with anhydrous $Na_2SO_4$, filtered and concentrated in vacuum. The residue was purified by flash silica gel chromatography (ISCO®; 4 g SepaFlash® Silica Flash Column, Eluent of 0~5% Ethyl acetate/Commercial hexanes gradient @ 40 mL/min). Compound methyl (4R)-2-sec-butyl-4,5-dihydrothiazole-4-carboxylate (270 mg, 725.69 μmol, 8.36% yield, 54.10% purity) was obtained as a colorless oil.

**Step 3**

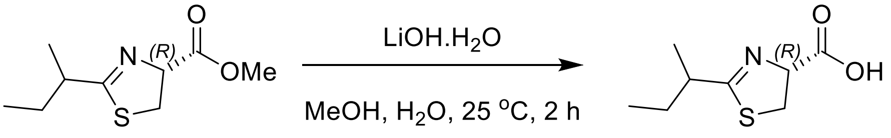


**Step 3**

**(4R)-2-sec-butyl-4,5-dihydrothiazole-4-carboxylic acid**

A mixture of methyl (4R)-2-sec-butyl-4,5-dihydrothiazole-4-carboxylate (250 mg, 1.24 mmol, 1 eq), $LiOH.H_2O$ (104.23 mg, 2.48 mmol, 2 eq) in MeOH (1 mL) and $H_2O$ (0.2 mL) was degassed and purged with $N_2$ for 3 times, and then the mixture was stirred at 25 °C for 2 h under $N_2$ atmosphere. The reaction mixture was concentrated under reduced pressure to remove solvent. Compound (4R)-2-sec-butyl-4,5-dihydrothiazole-4-carboxylic acid (210 mg, 1.12 mmol, 90.29% yield) was obtained as a yellow solid. The crude product (4R)-2-sec-butyl-4,5-dihydrothiazole-4-carboxylic acid (210 mg, 1.12 mmol, 90.29% yield) was used into the next step without further purification.

#### Step 4 (ET100716-242):

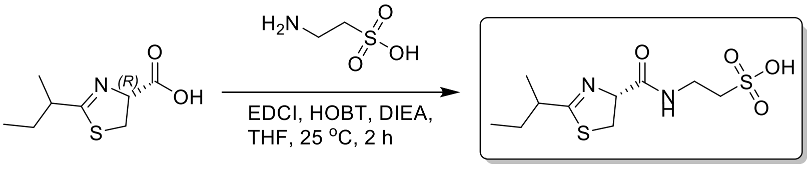


**Step 4** **HKMS-0213**

**2-[[(4R)-2-sec-butyl-4,5-dihydrothiazole-4-carbonyl]amino]ethanesulfonic acid**

A mixture of (4R)-2-sec-butyl-4,5-dihydrothiazole-4-carboxylic acid (70 mg, 373.81 μmol, 1 eq), 2-aminoethanesulfonic acid (46.78 mg, 373.81 μmol, 46.60 μL, 1 eq), EDCI (85.99 mg, 448.58 μmol, 1.2 eq), HOBt (65.66 mg, 485.96 μmol, 1.3 eq) and DIEA (96.63 mg, 747.63 μmol, 130.22 μL, 2 eq) in THF (1 mL) was degassed and purged with $N_2$ for 3 times, and then the mixture was stirred at 25 °C for 2 h under $N_2$ atmosphere. The reaction mixture was concentrated under reduced pressure to remove solvent. The residue was purified by prep-HPLC (neutral condition). (column: Waters Xbridge BEH C18 100*30mm*10um;mobile phase: [$H_2O$(10mM $NH_4HCO_3$)-ACN];gradient:1%-25% B over 15.0 min). Compound 2-[[(4R)-2-sec-butyl-4,5-dihydrothiazole-4-carbonyl]amino]ethanesulfonic acid (10.3 mg, 34.99 μmol, 9.36% yield, 99%< purity) was obtained as a colorless oil.

**LCMS:** ($M+H^+$): 295.1 @ 2.438 min (0-60 % ACN in $H_2O$, 6 min)

**$^1$H NMR:** (400 MHz, DEUTERIUM OXIDE) δ 5.01 (ddd, *J* = 4.3, 7.8, 9.8 Hz, 1H), 3.66- 3.53 (m, 3H), 3.41 (dd, *J* = 7.8, 11.4 Hz, 1H), 3.07 (dt, *J* = 2.6, 6.6 Hz, 2H), 2.69 (sxt, *J* = 7.0 Hz, 1H), 1.66-1.45 (m, 2H), 1.16 (dd, *J* = 2.1, 6.8 Hz, 3H), 0.85 (dt, *J* = 3.7, 7.5 Hz, 3H)

**$^{13}$C NMR:** (100 MHz, DEUTERIUM OXIDE) δ 186.13, 185.96, 173.47, 173.47, 77.01, 76.78, 49.42, 40.74, 40.54, 35.55, 35.14, 34.52, 34.52, 28.46, 28.27, 18.32, 18.24, 16.69, 10.95, 10.82

### HKMS-0291

#### 1H NMR

**Compound ID: HKMS-0291**

ET92536-1014-P1Z DMSO Bruker_02_C_400MHz

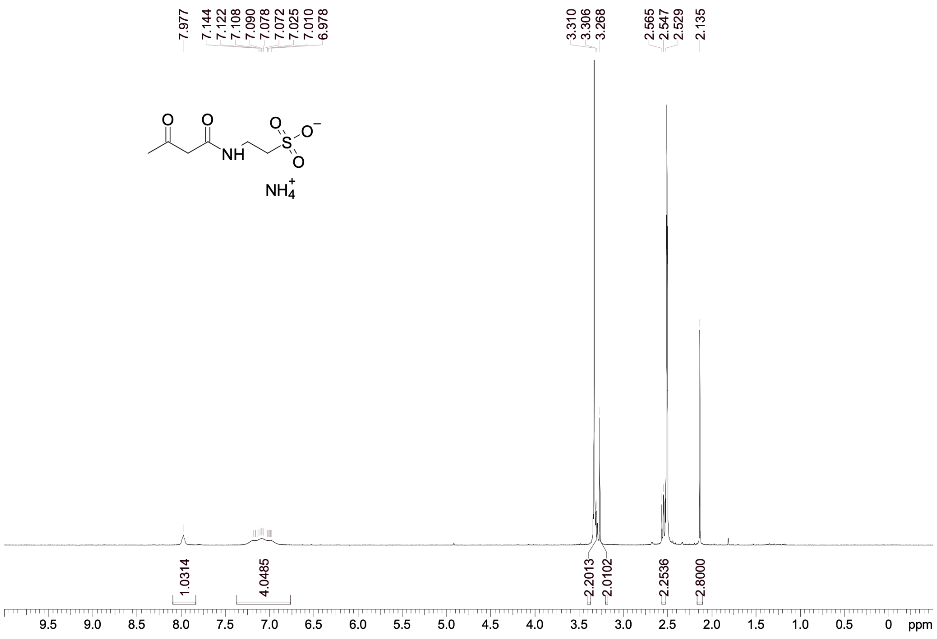


#### 13C NMR

**Compound ID: HKMS-0291**

ET92536-1170-P1Y DMSO Bruker_02_A_400MHz 13C-NMR

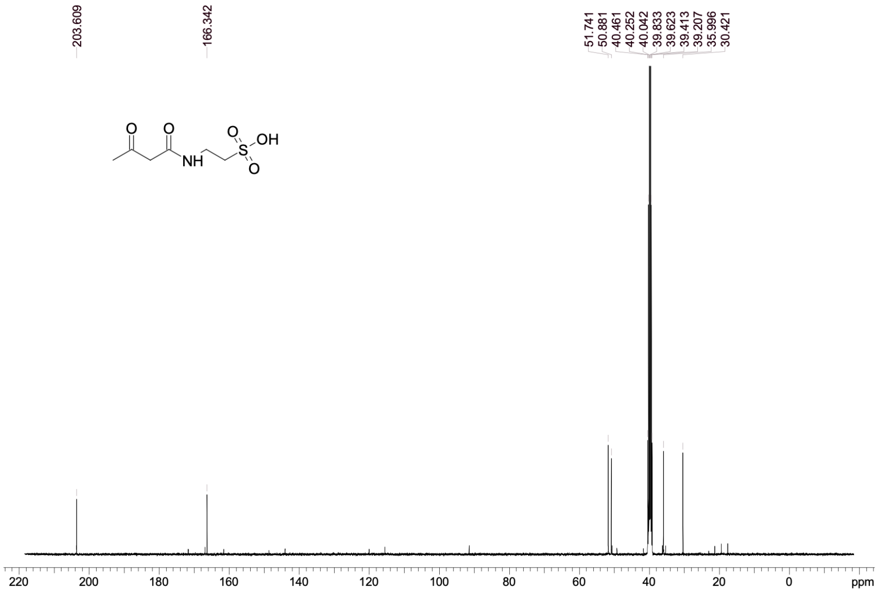

### HKMS-0300

#### 1H NMR

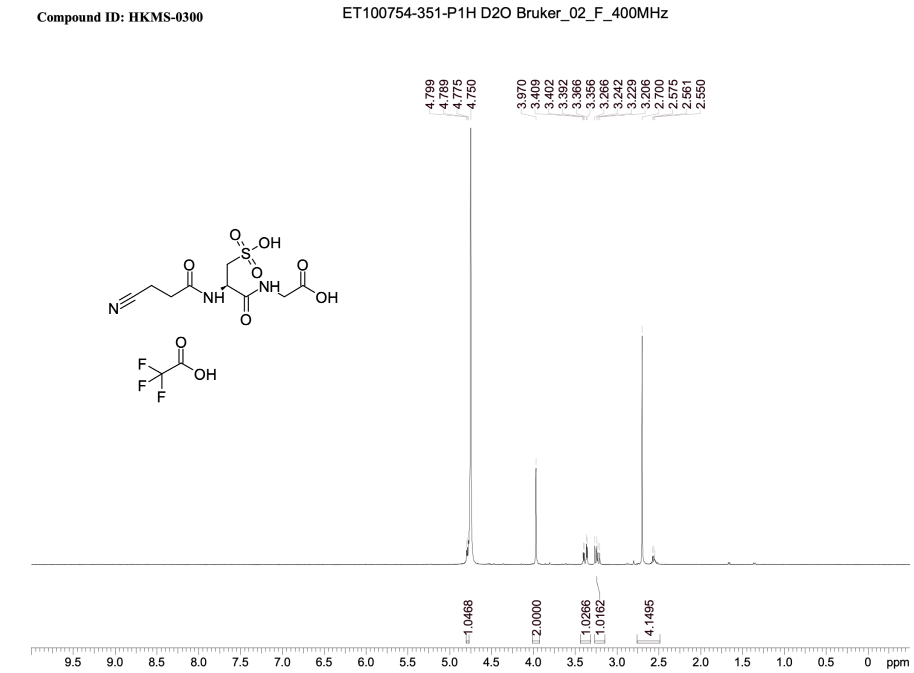


#### 13C NMR

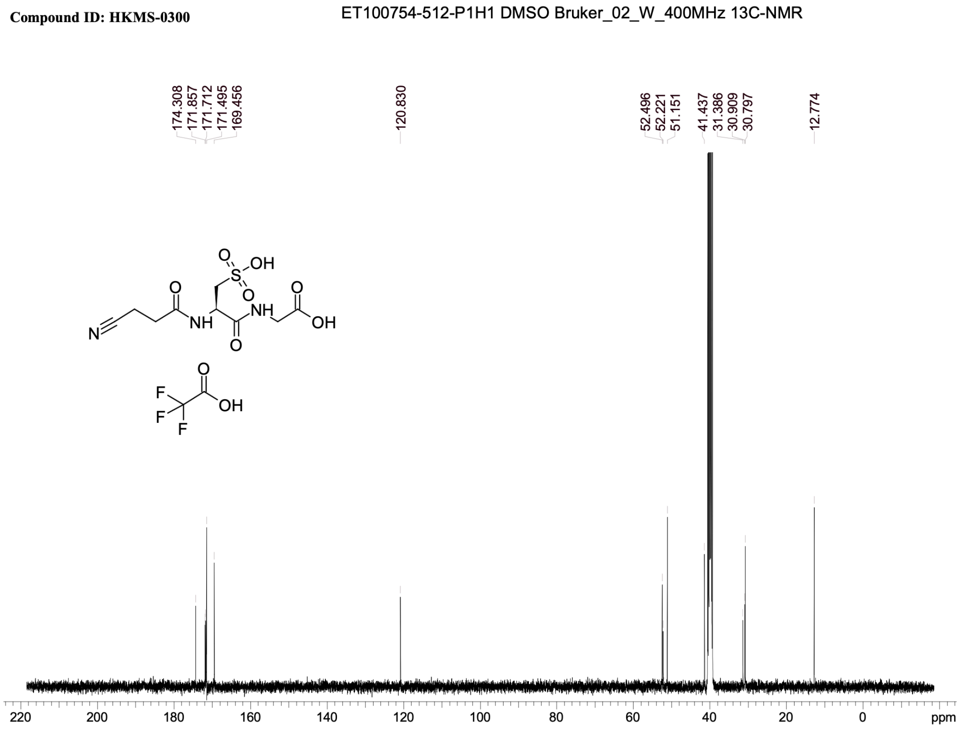

### HKMS-0302

#### 1H NMR

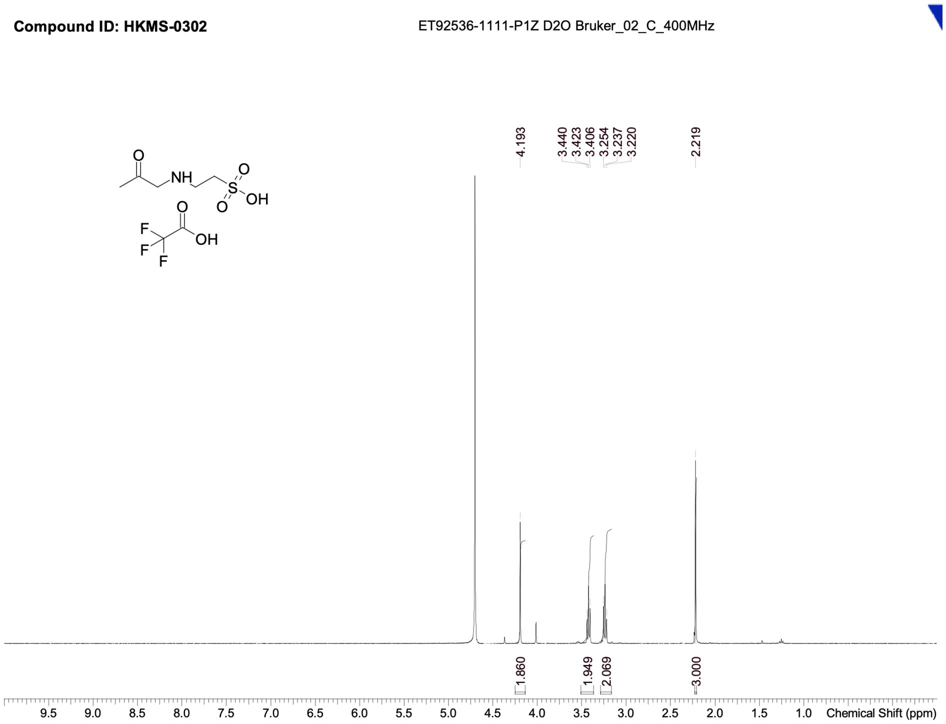


#### 13C NMR

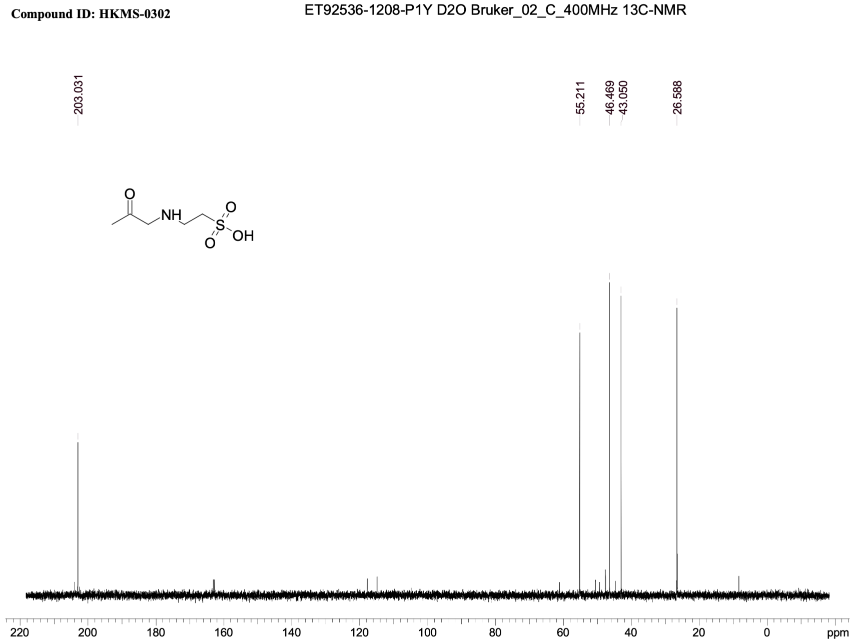

### HKMS-0303

#### 1H NMR

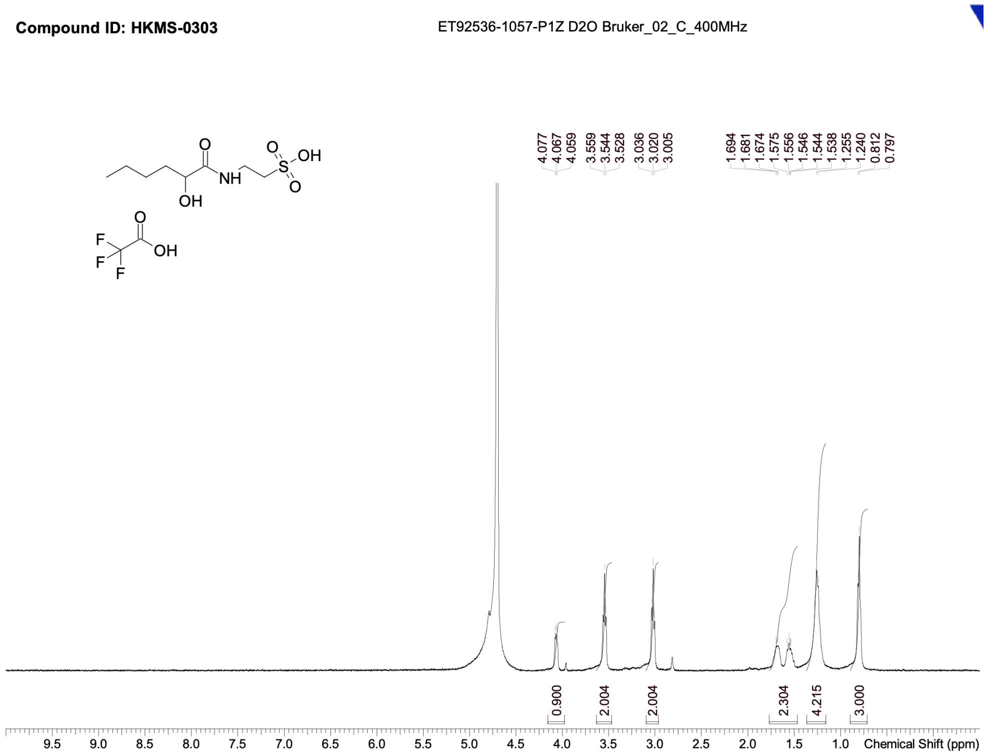


#### 13C NMR

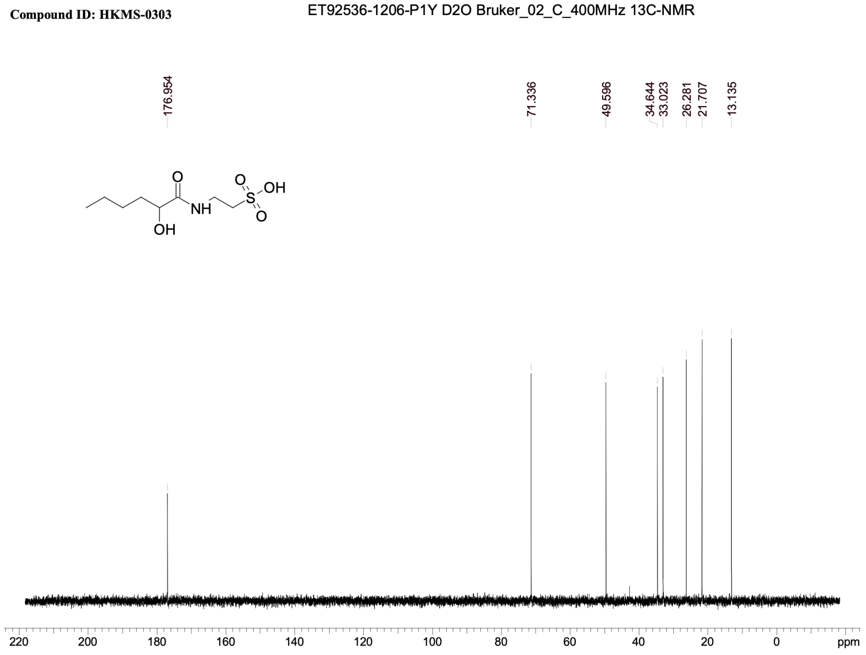

### HKMS-0305

#### 1H NMR

**Compound ID: HKMS-0305** ET92536-1079-P1Z1 D2O Bruker_02_C_400MHz

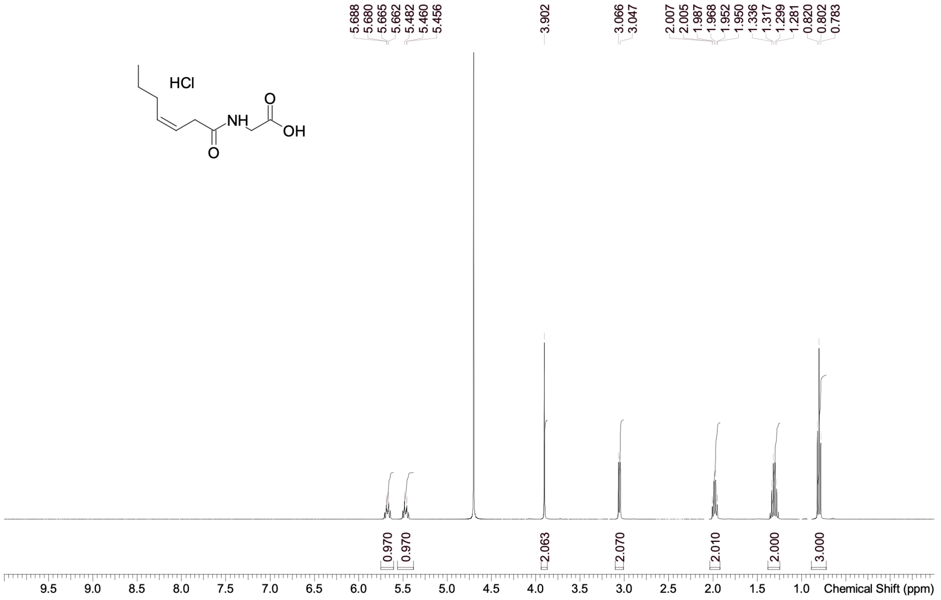


#### 13C NMR

**Compound ID: HKMS-0305** ET92536-1207-P1Y D2O Bruker_02_C_400MHz 13C-NMR

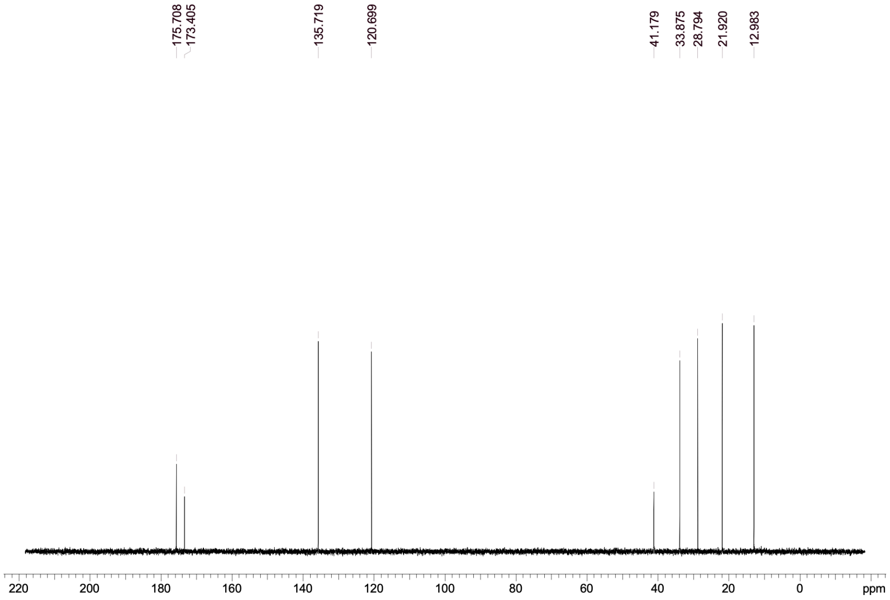

### HKMS-0310

#### 1H NMR

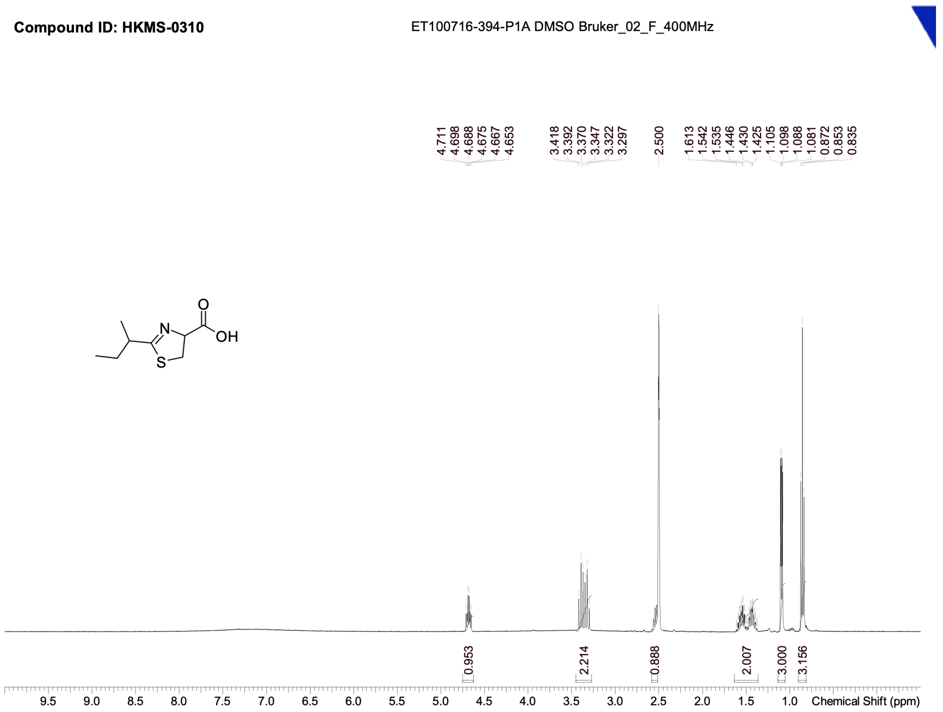


#### 13C NMR

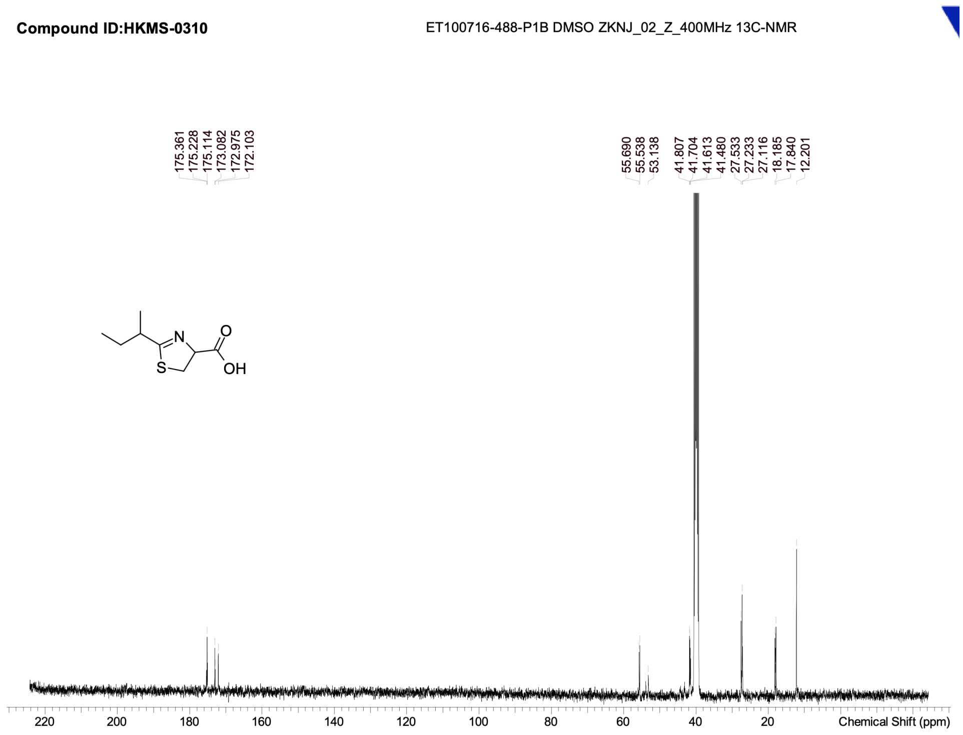

### HKMS-0322

#### 1H NMR

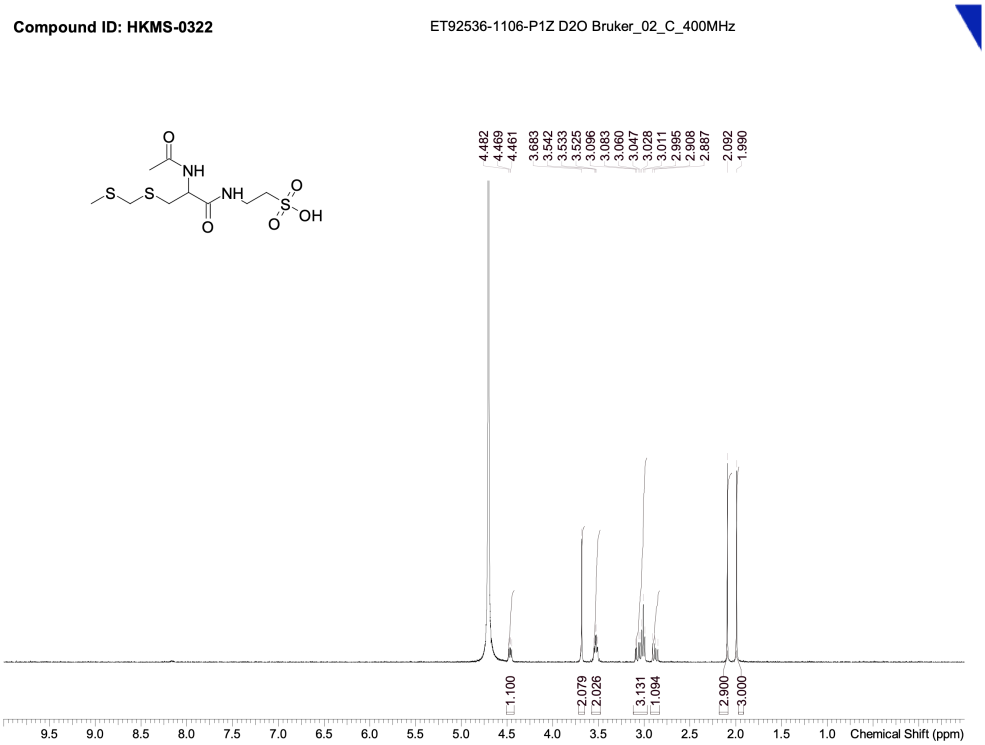


#### 13C NMR

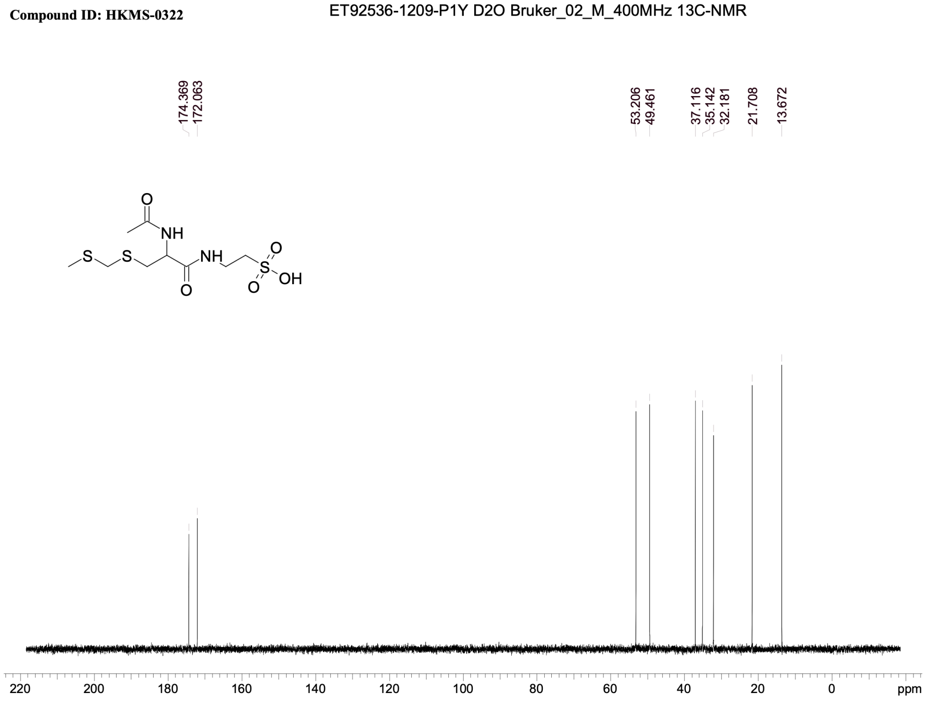

HKMS-0078

1H NMR

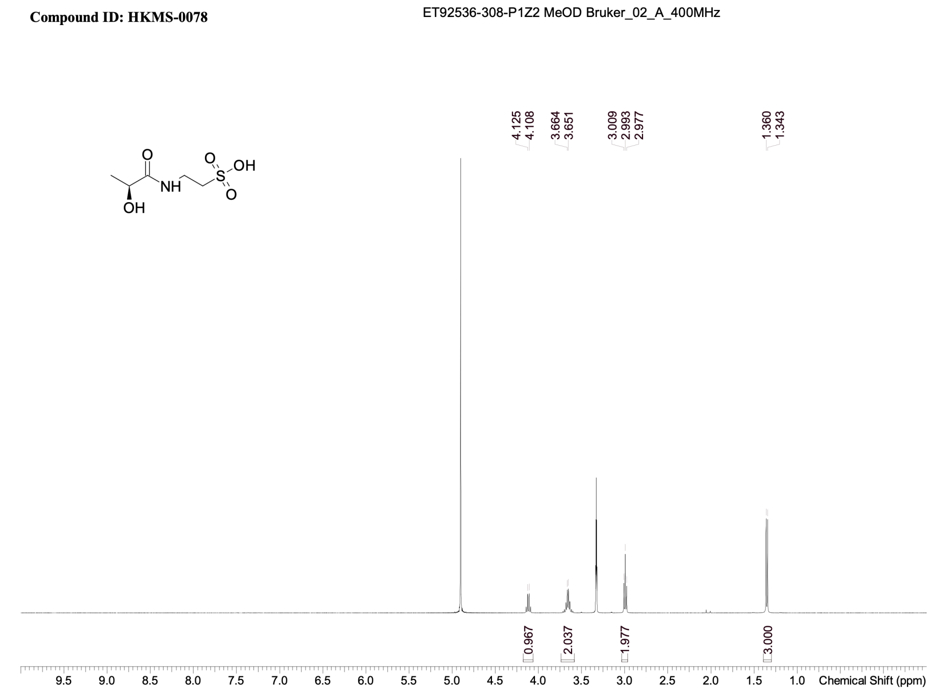


13C NMR

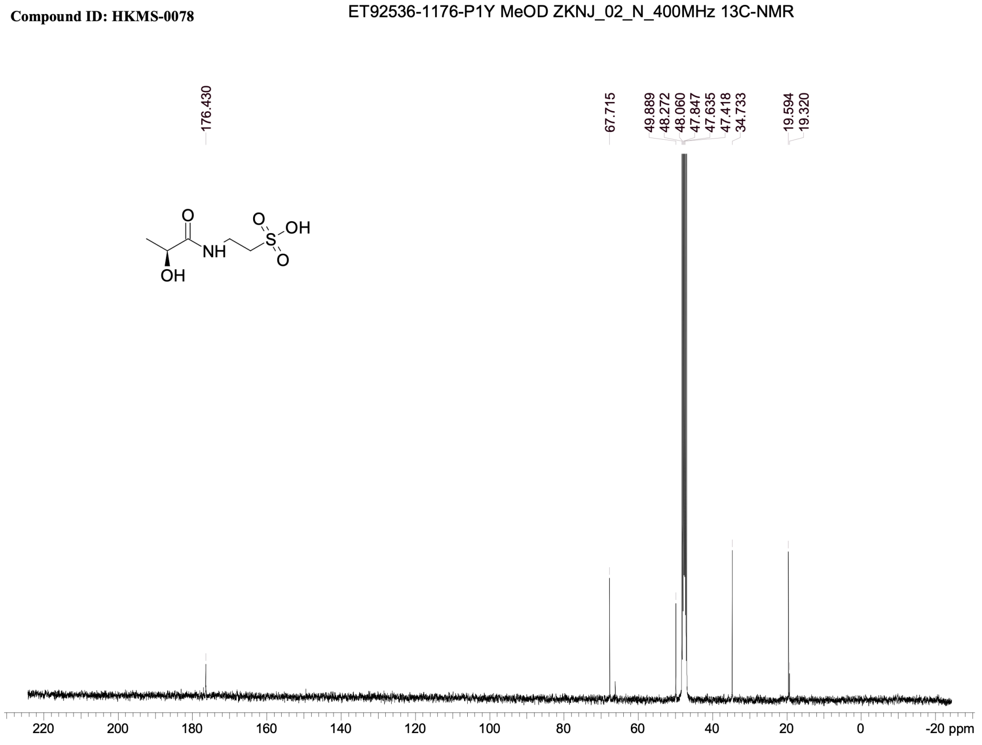

### HKMS-0157

#### 1H NMR

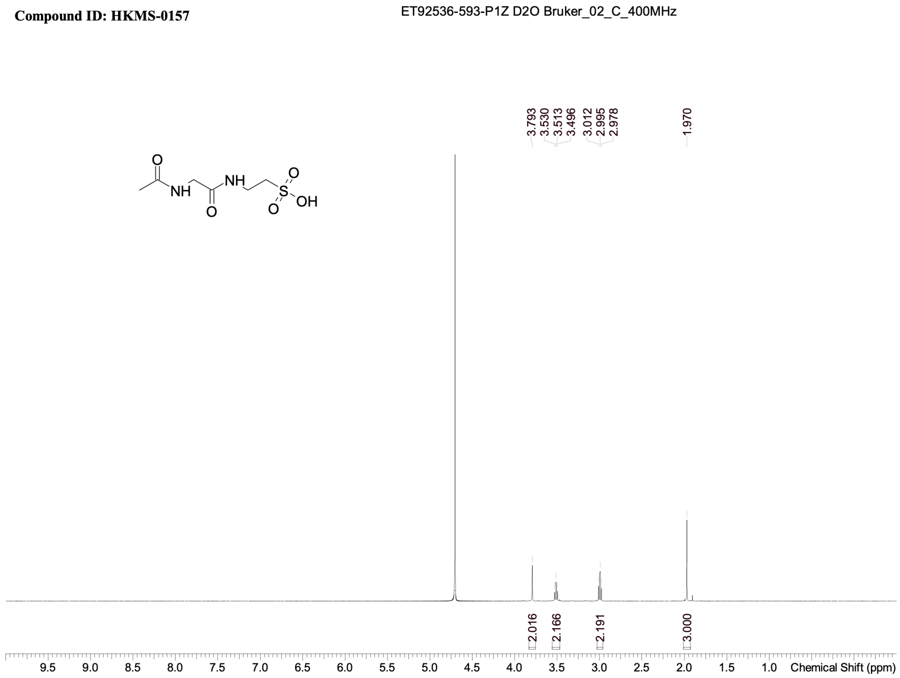


#### 13C NMR

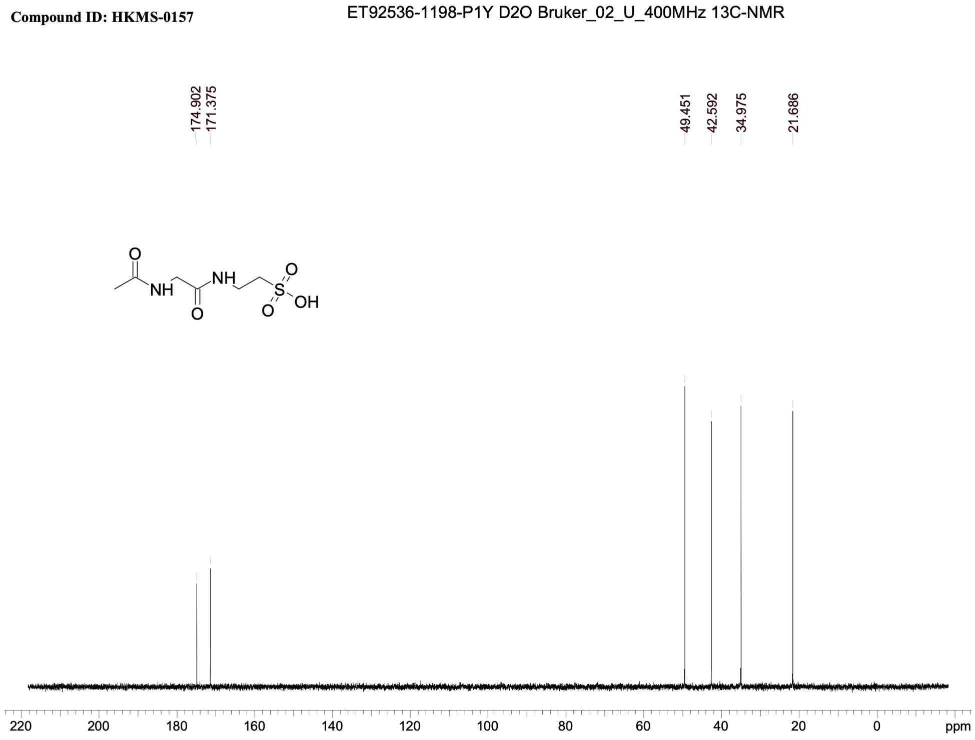

### HKMS-0158

#### 1H NMR

**Compound ID: HKMS-0158**

ET92536-648-P1Z1 D2O Bruker_02_L_400MHz

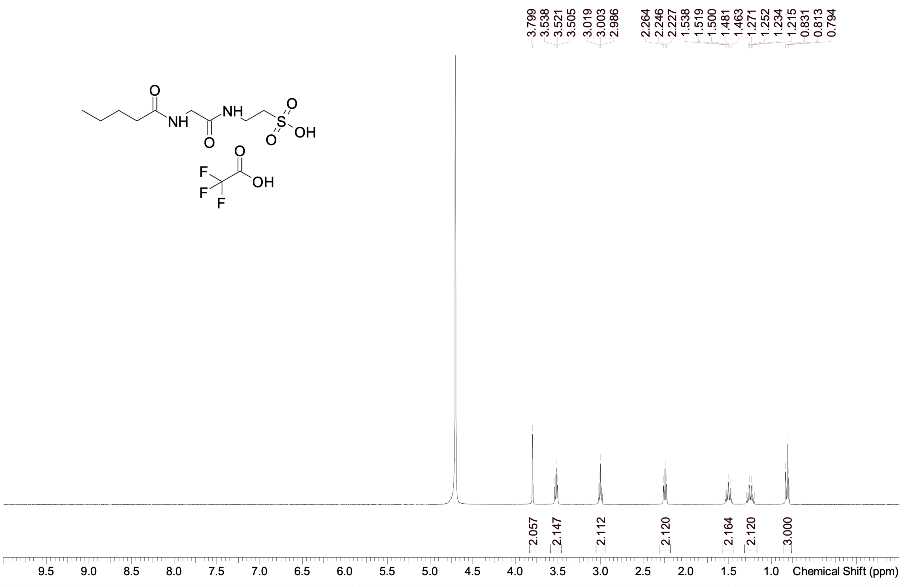


#### 13C NMR

**Compound ID: HKMS-0158**

ET92536-1173-P1Y D2O Bruker_02_F_400MHz 13C-NMR

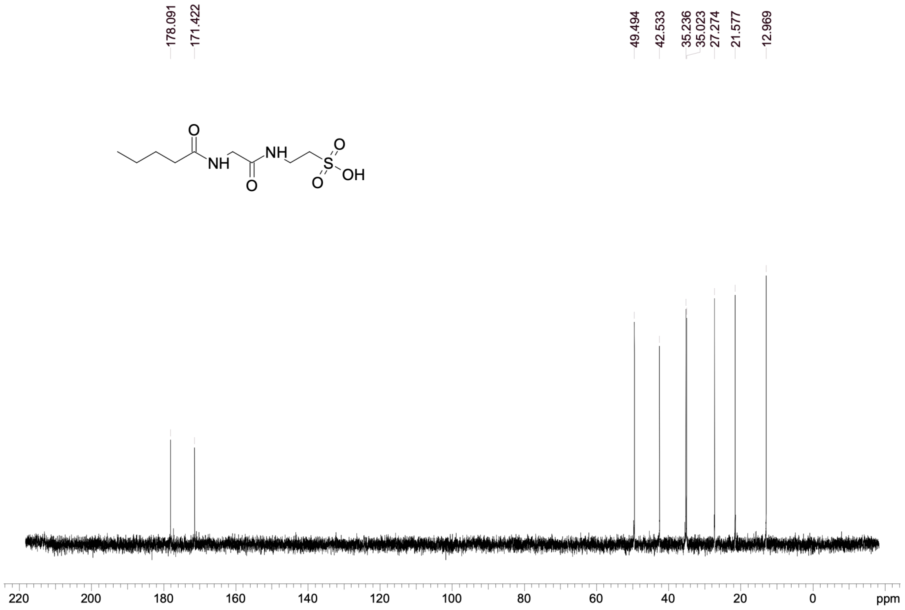

### HKMS-0159

#### 1H NMR

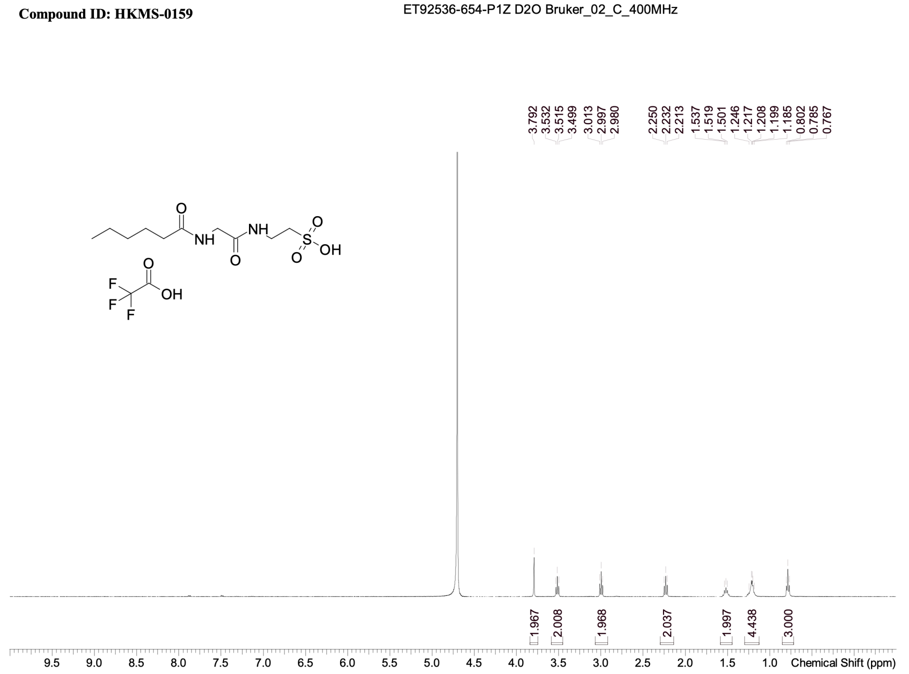


#### 13C NMR

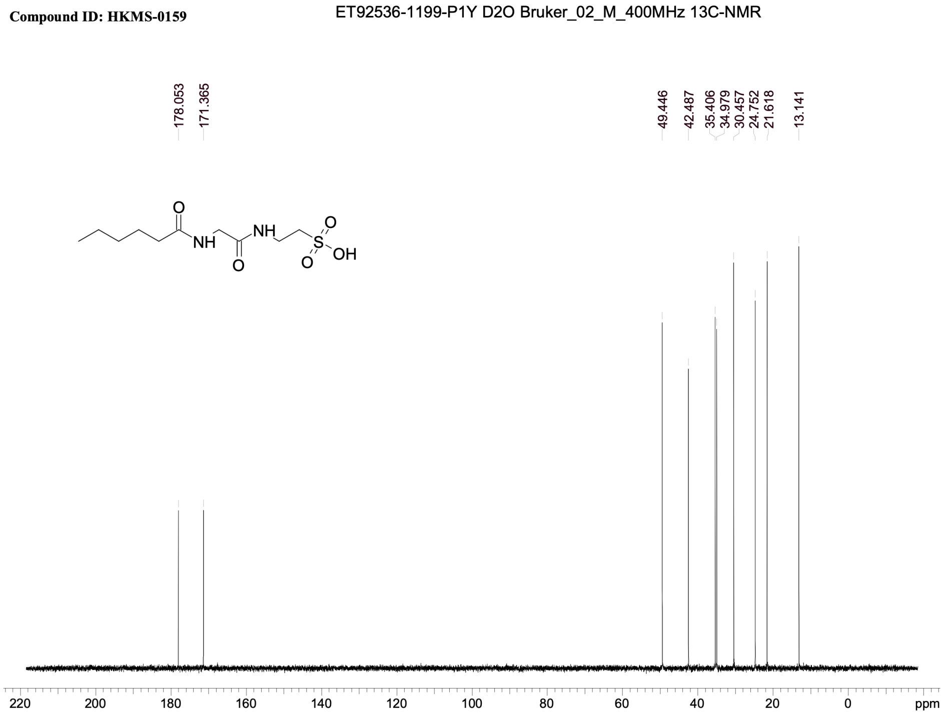

### HKMS-0160

#### 1H NMR

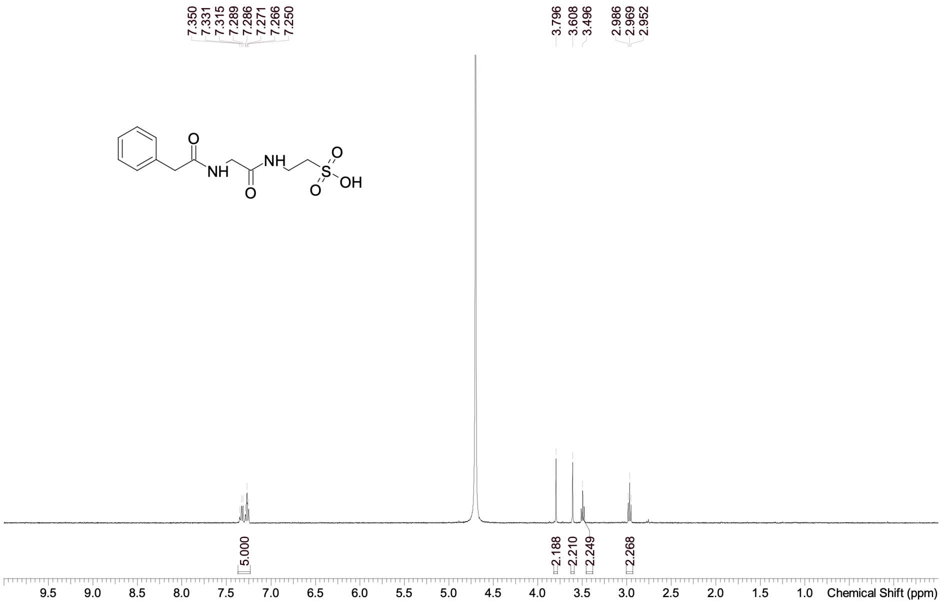


#### 13C NMR

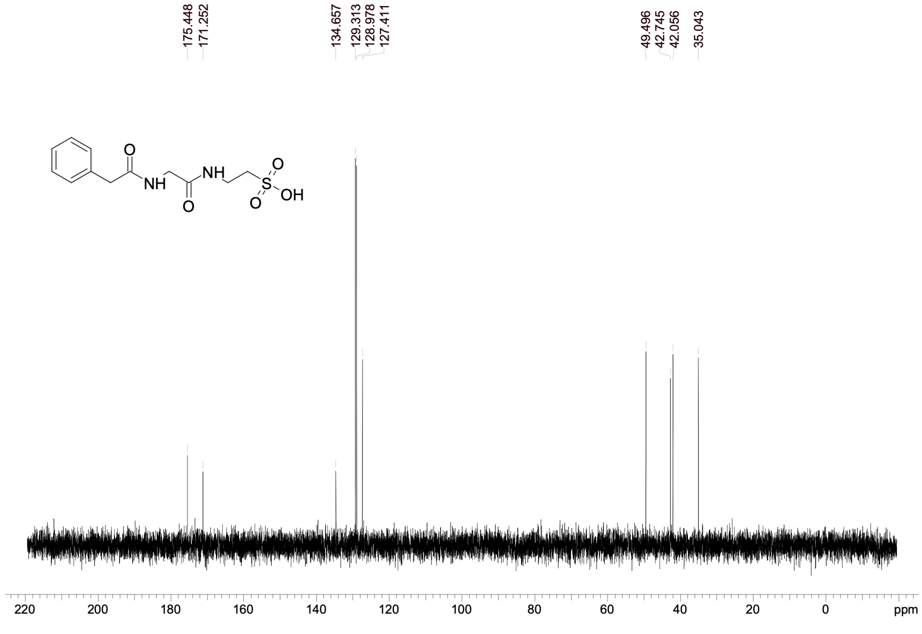

### HKMS-0162

#### 1H NMR

**Compound ID: HKMS-0162**

ET92536-651-P1Z D2O Bruker_02_C_400MHz

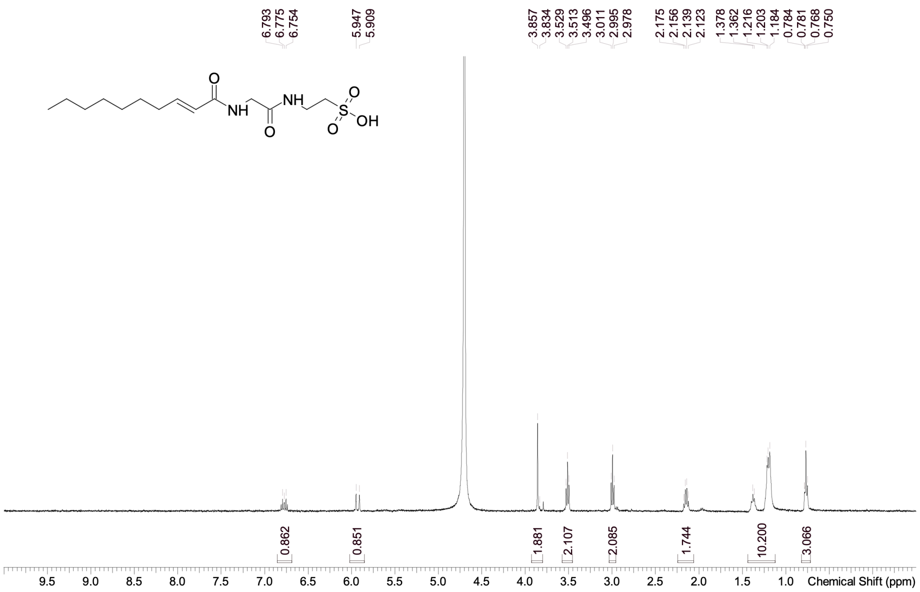


#### 13C NMR

**Compound ID: HKMS-0162**

ET100754-502-P1H2 D2O Bruker_02_M_400MHz 13C-NMR

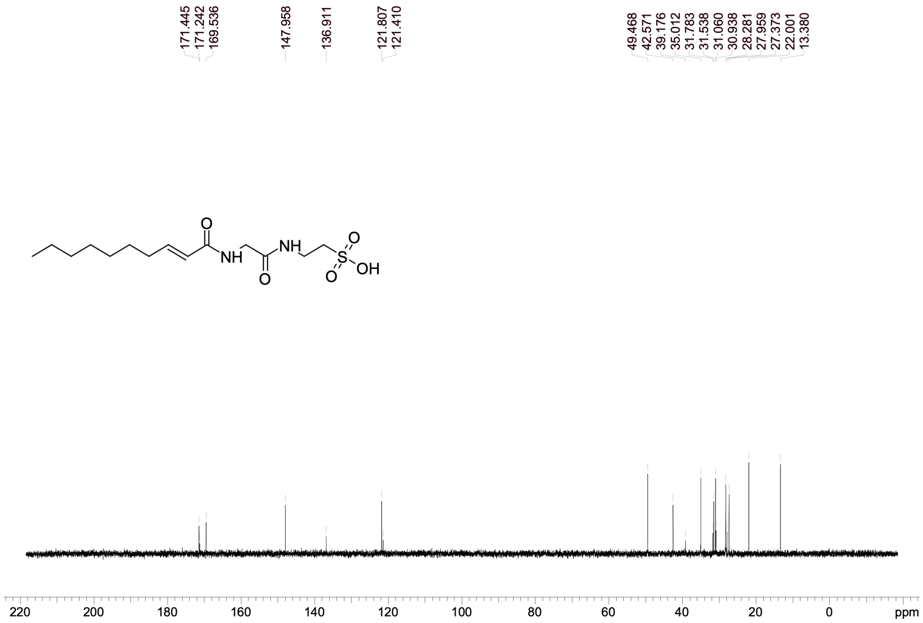

### HKMS-0163

#### 1H NMR

**Compound ID: HKMS-0163**

ET92536-640-P1Z2 D2O Bruker_02_L_400MHz

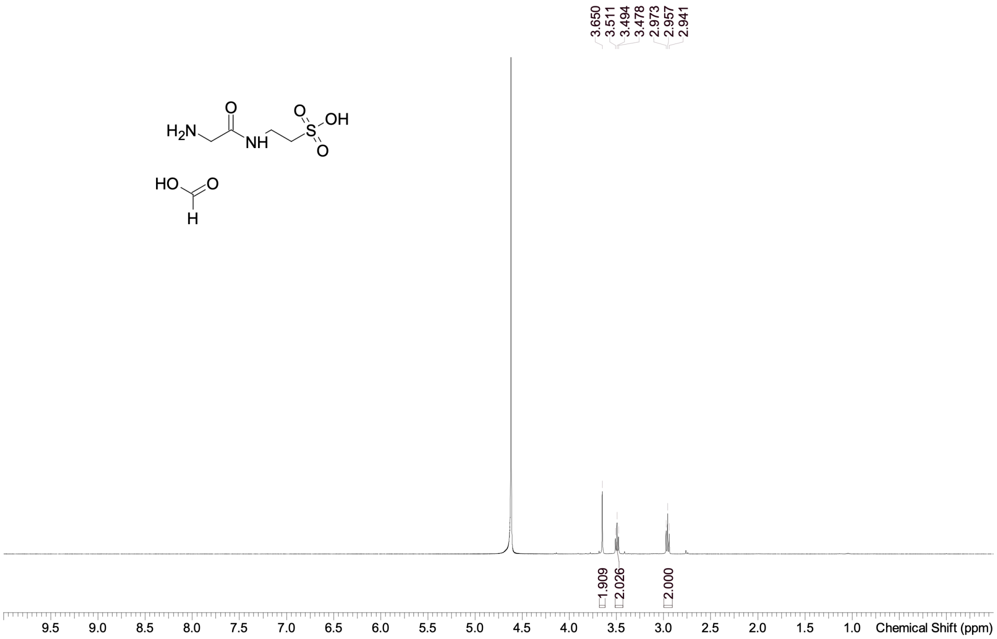


#### 13C NMR

**Compound ID: HKMS-0163**

ET92536-1181-PIY D2O Bruker_02_L_400MHz 13C-NMR

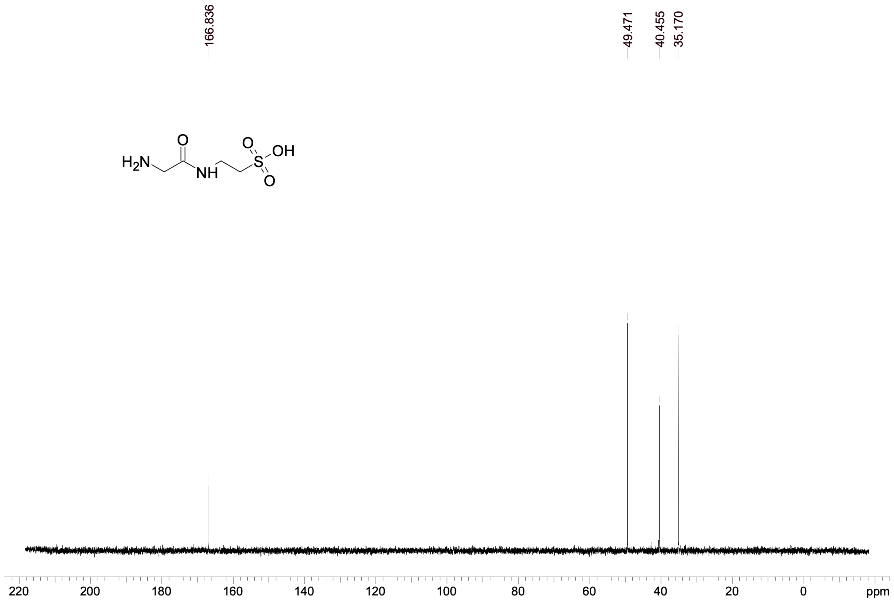

### HKMS-0164

#### 1H NMR

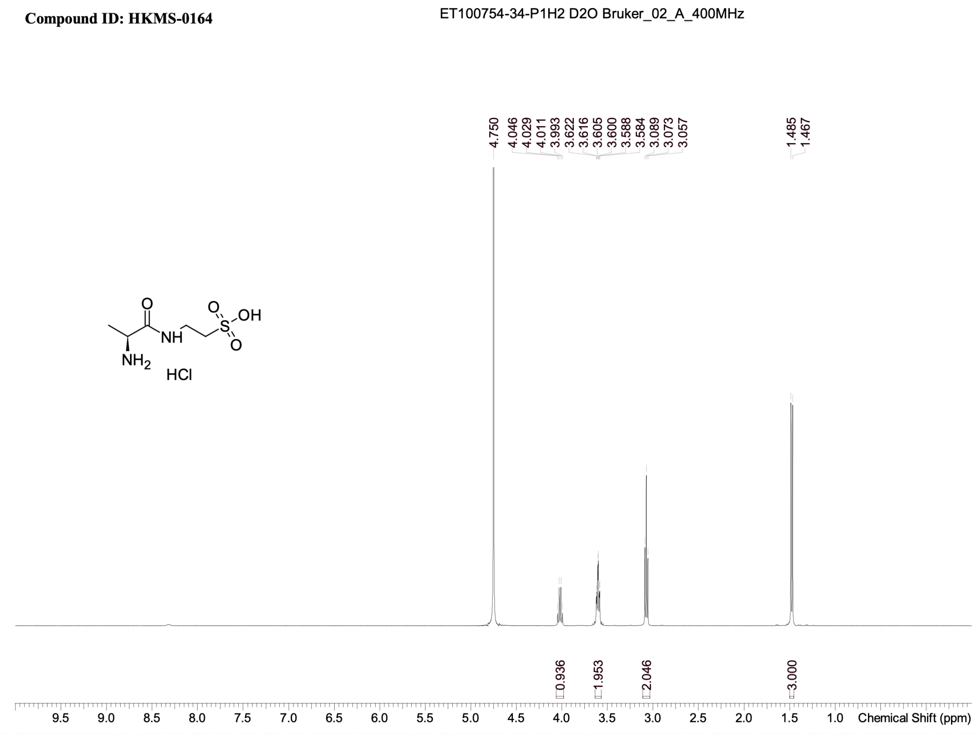


#### 13C NMR

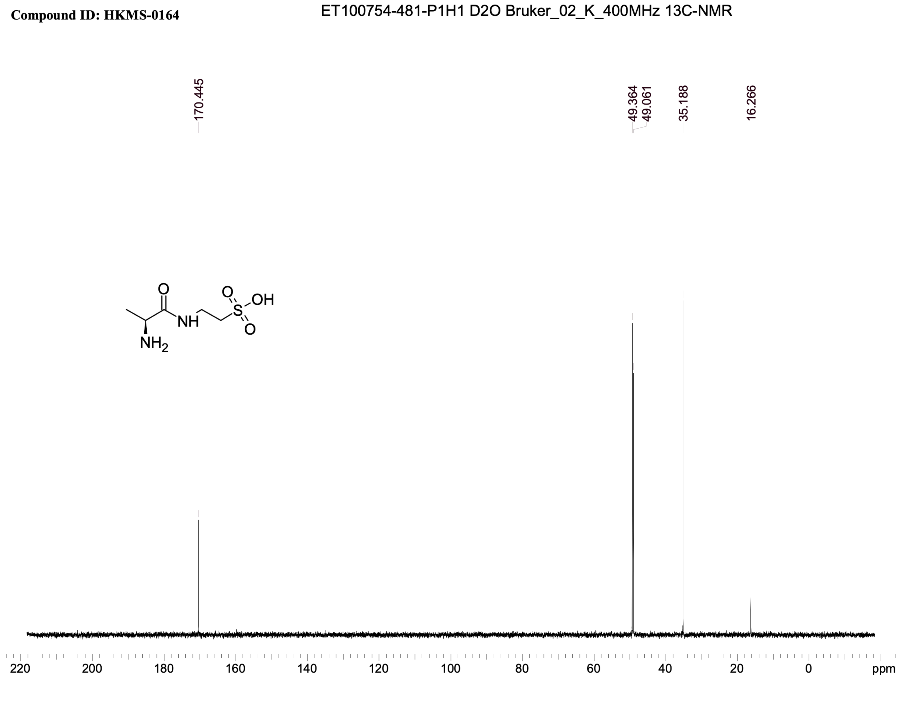

### HKMS-0172

#### 1H NMR

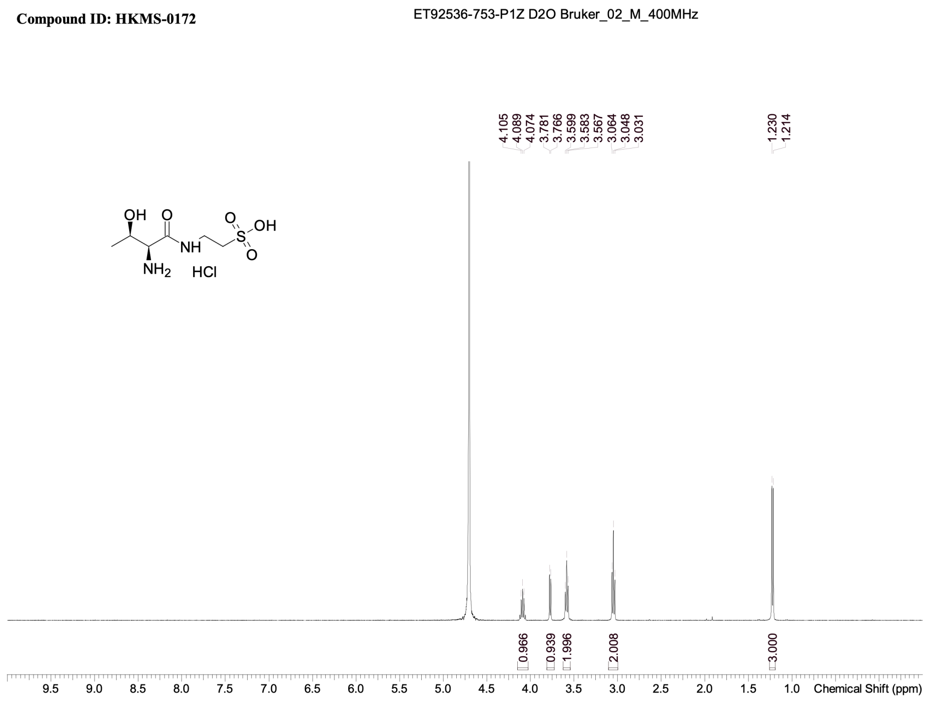


#### 13C NMR

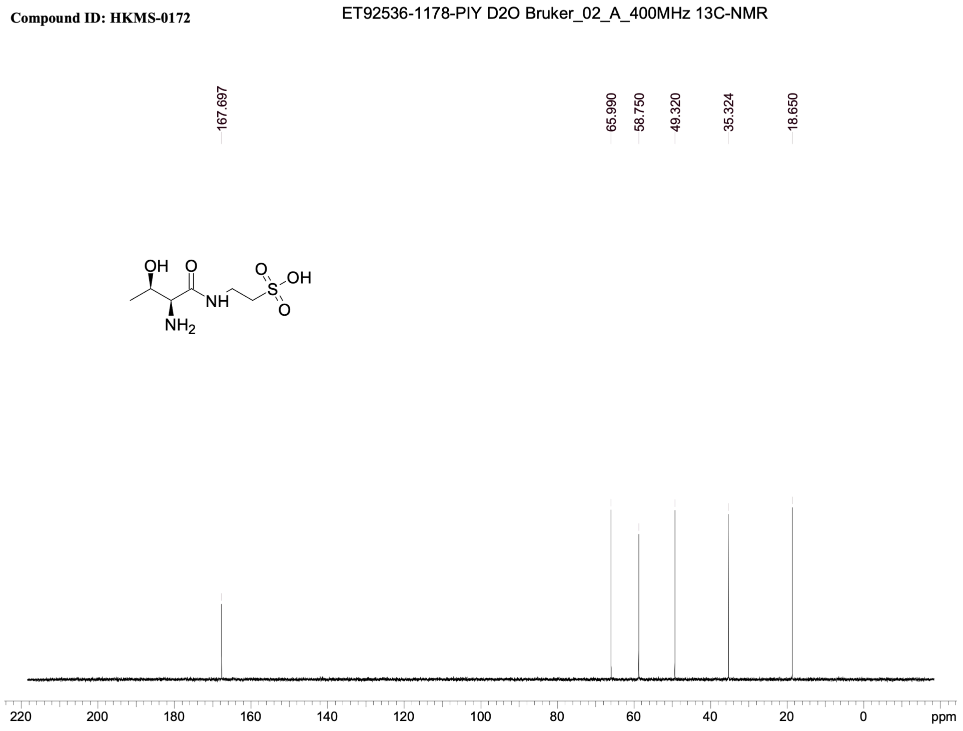

### HKMS-0174

#### 1H NMR

**Compound ID: HKMS-0174**

ET92536-667-P1Z1 D2O Bruker_02_L_400MHz

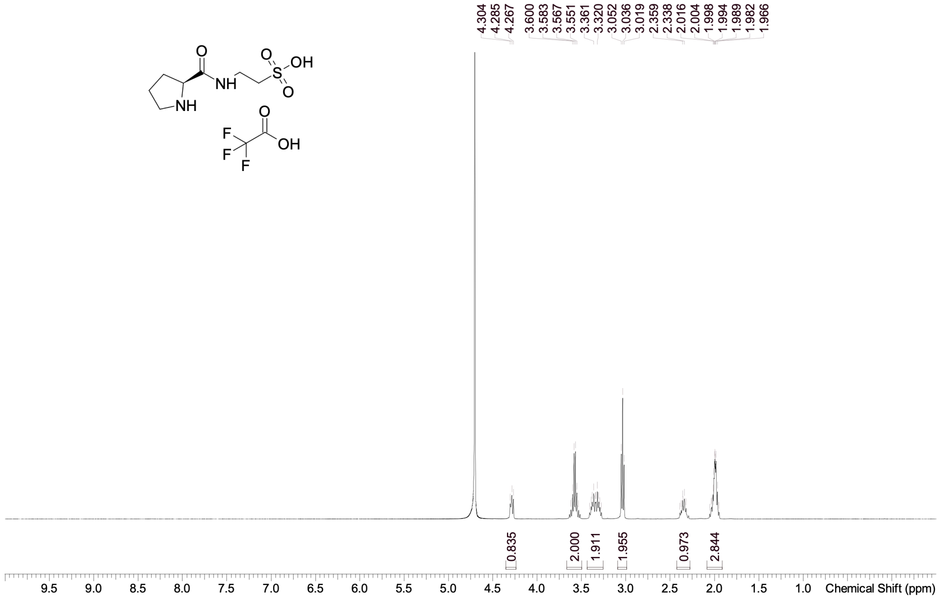


#### 13C NMR

**Compound ID: HKMS-0174**

ET92536-1182-PIY D2O Bruker_02_004_400MHz 13C-NMR

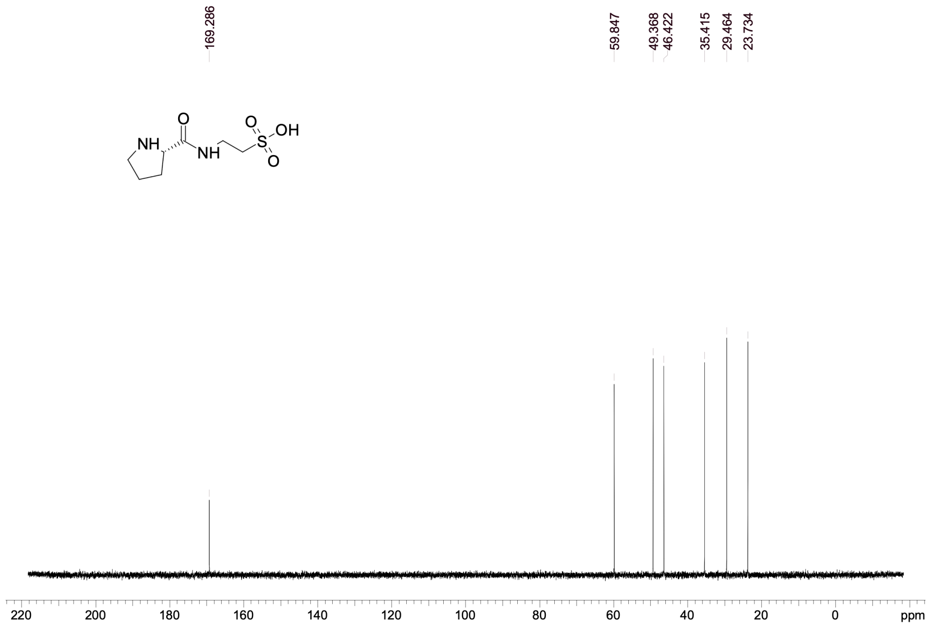

### HKMS-0182

#### 1H NMR

#### 13C NMR

**Compound ID: HKMS-0182**

ET92536-1184-PIY D2O Bruker_02_003_400MHz 13C-NMR

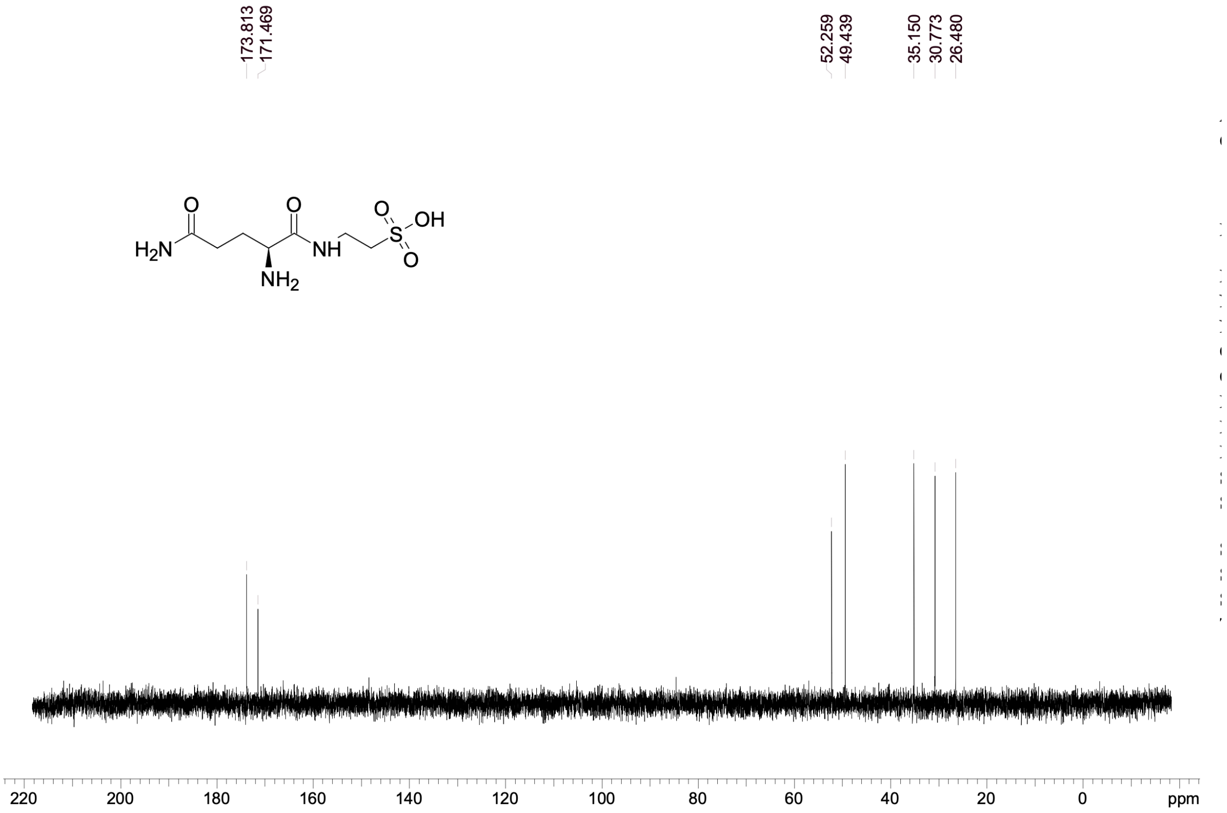

### HKMS-0200

#### 1H NMR

**Compound ID: HKMS-0200**

ET100754-98-P1H D2O Bruker_02_M_400MHz

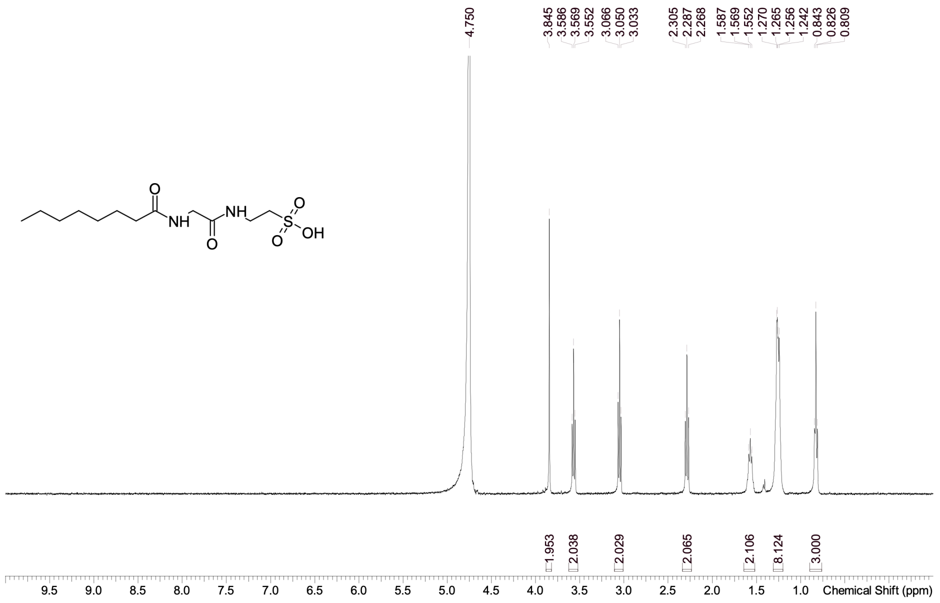


#### 13C NMR

**Compound ID: HKMS-0200**

ET92536-1174-P1Y D2O Bruker_02_W_400MHz 13C-NMR

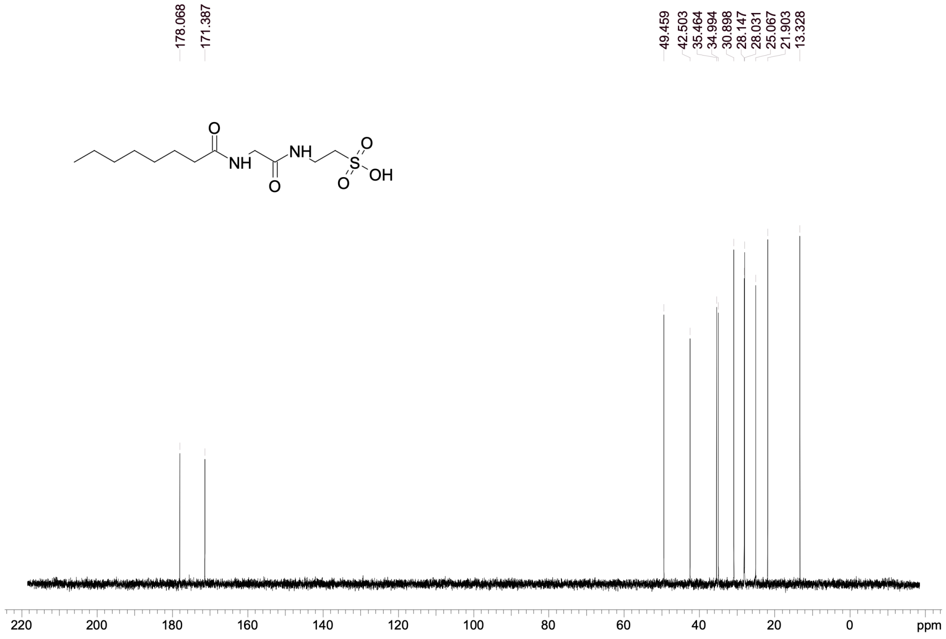

### HKMS-0206

#### 1H NMR

**Compound ID: HKMS-0206**

ET100754-239-P1H D2O Bruker_02_C_400MHz

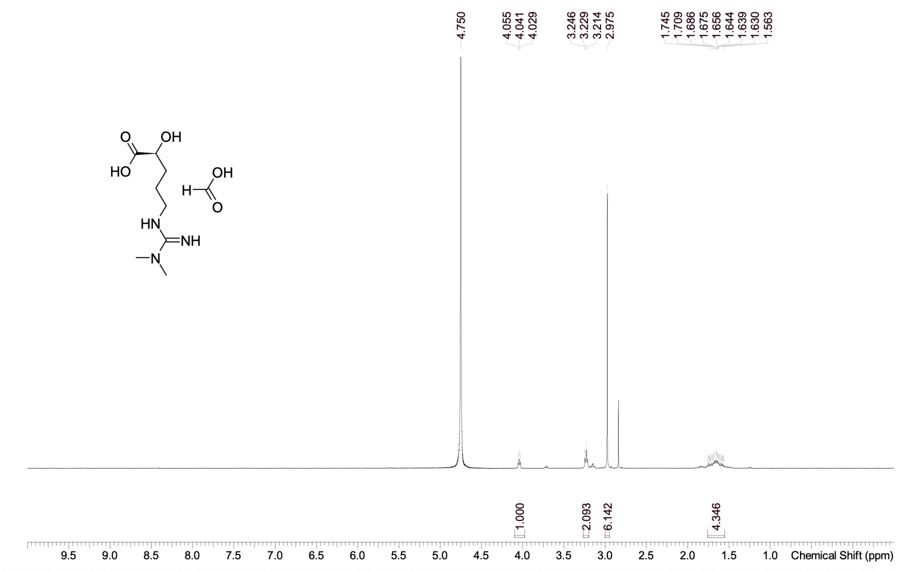


#### 13C NMR

**Compound ID: HKMS-0206**

ET100754-483-P1H1 D2O Bruker_02_K_400MHz 13C-NMR

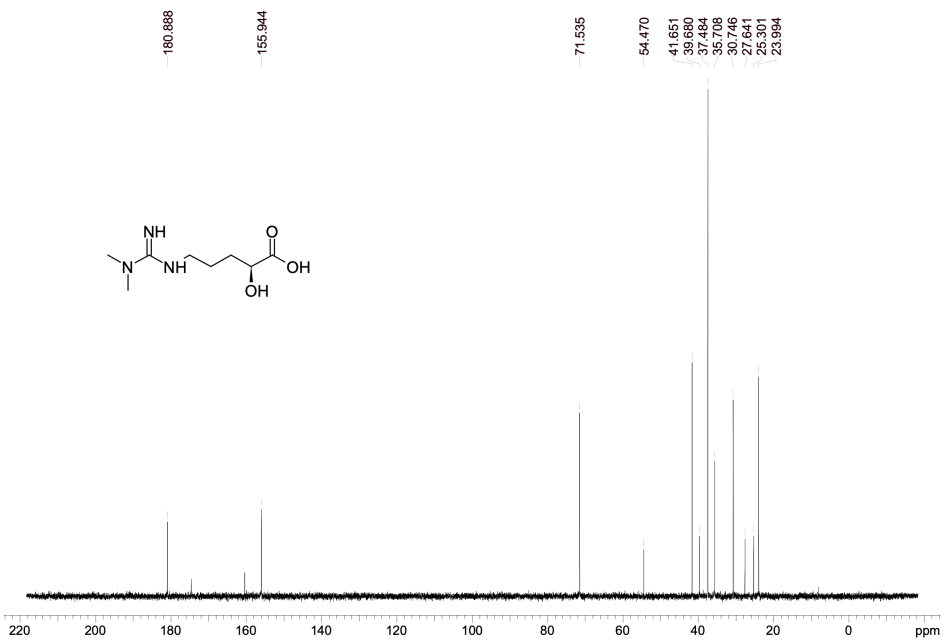

### HKMS-0210

#### 1H NMR

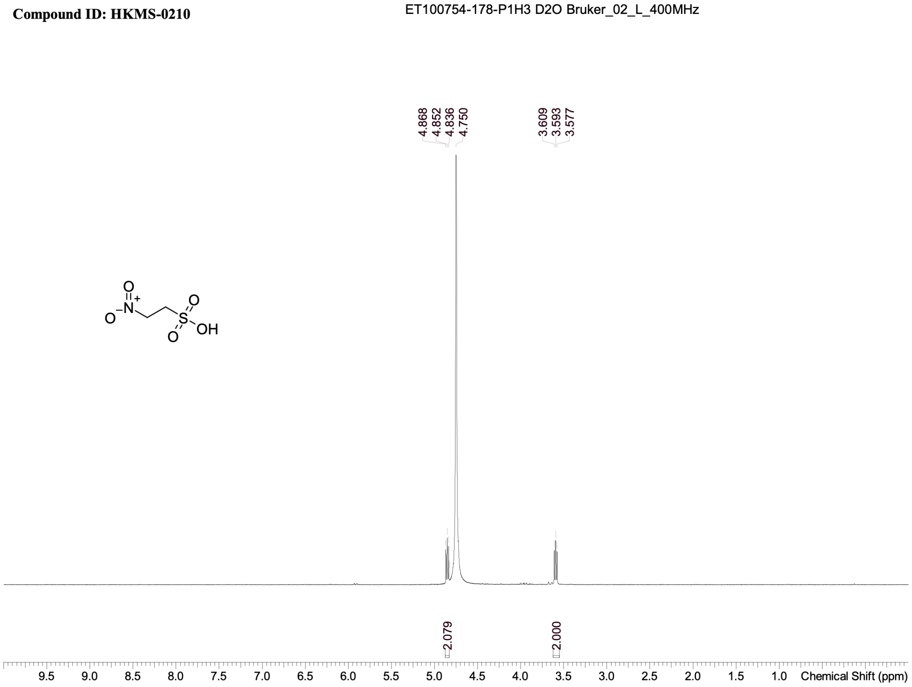


#### 13C NMR

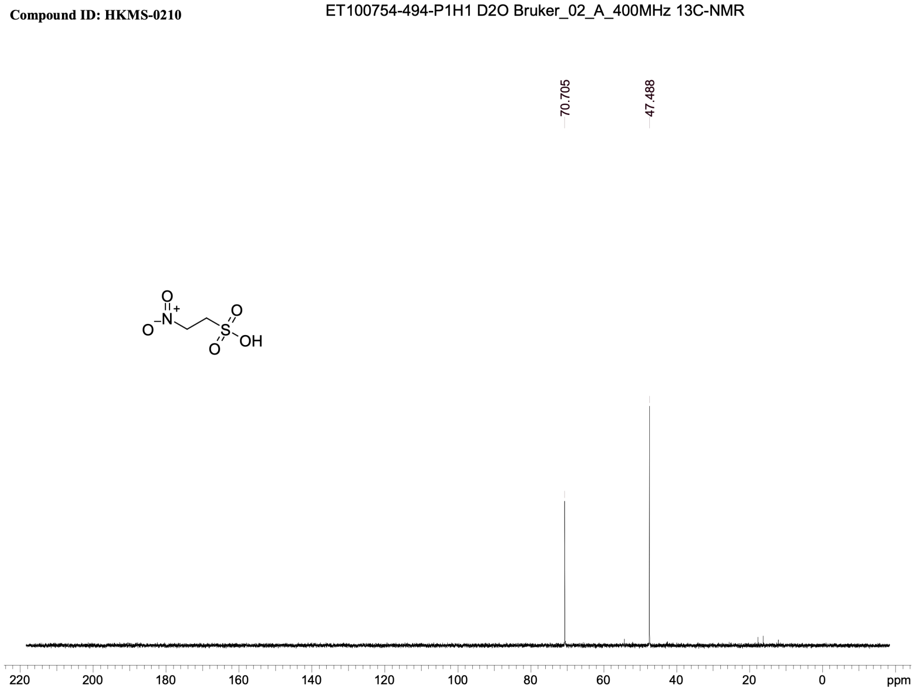

### HKMS-0223

#### 1H NMR

**Compound ID: HKMS-0223**

ET92536-921-P1Z D2O Bruker_02_M_400MHz

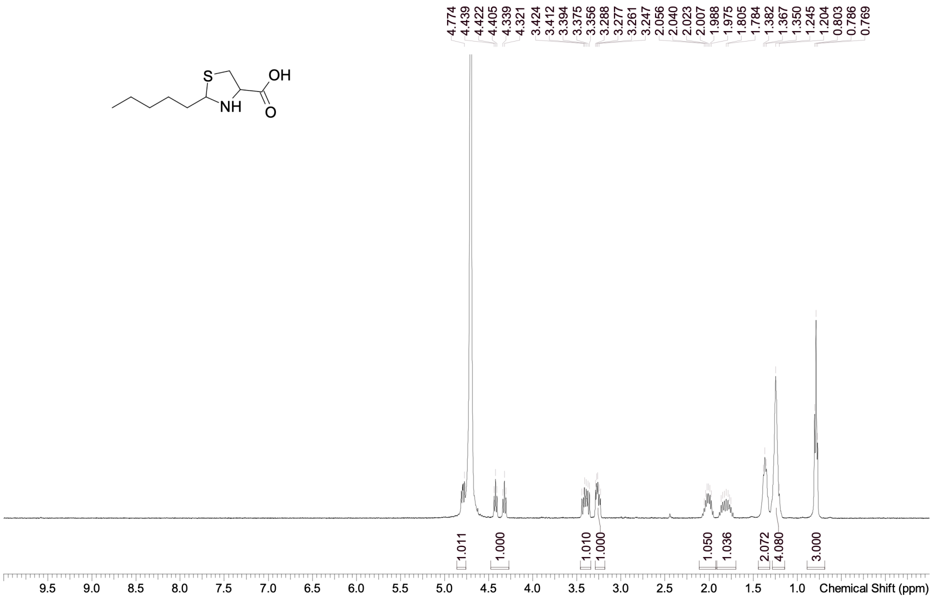


#### 13C NMR

**Compound ID: HKMS-0223**

ET92536-1189-P1Y DMSO Bruker_02_E_400MHz 13C-NMR

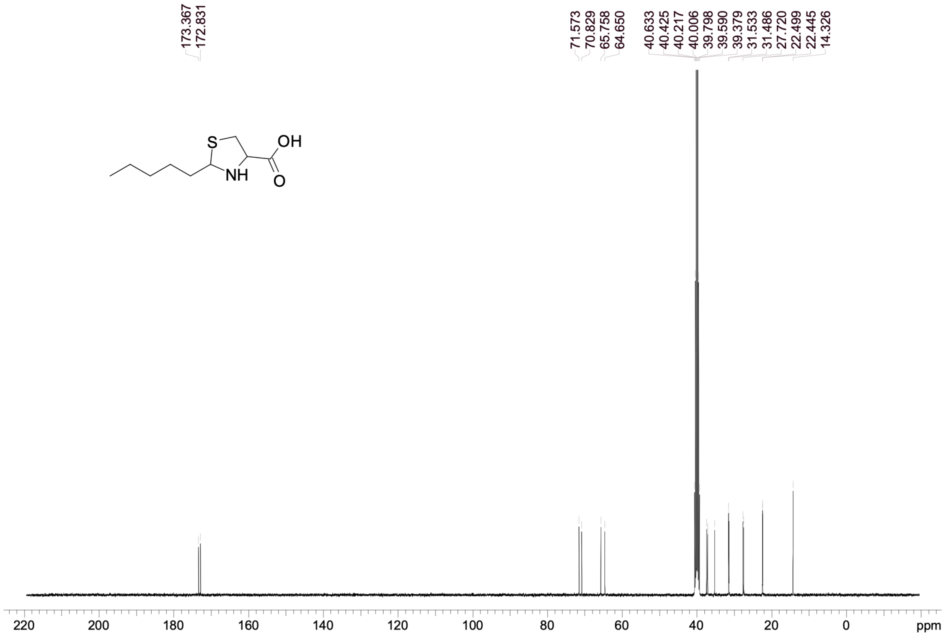

### HKMS-0230

#### 1H NMR

**Compound ID: HKMS-0230**

ET92536-919-P1Z1 D2O Bruker_02_M_400MHz

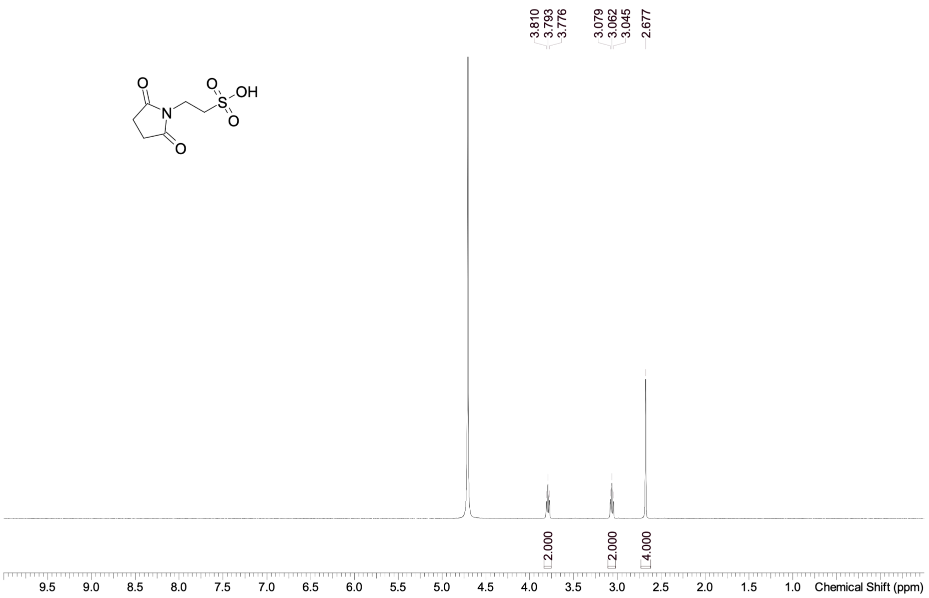


#### 13C NMR

**Compound ID: HKMS-0230**

ET92536-1190-P1Y D2O Bruker_02_L_400MHz 13C-NMR

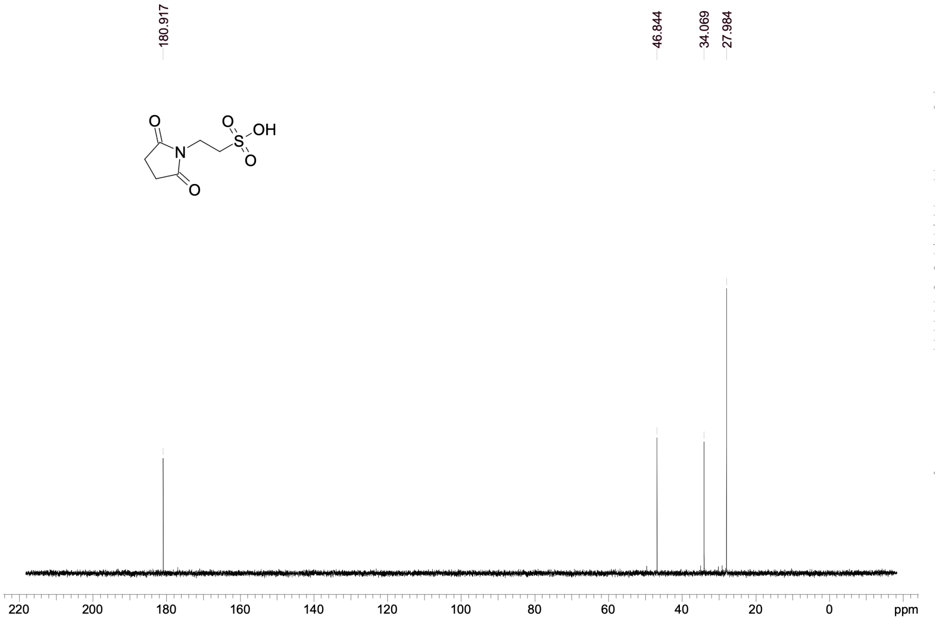

### HKMS-0235

#### 1H NMR

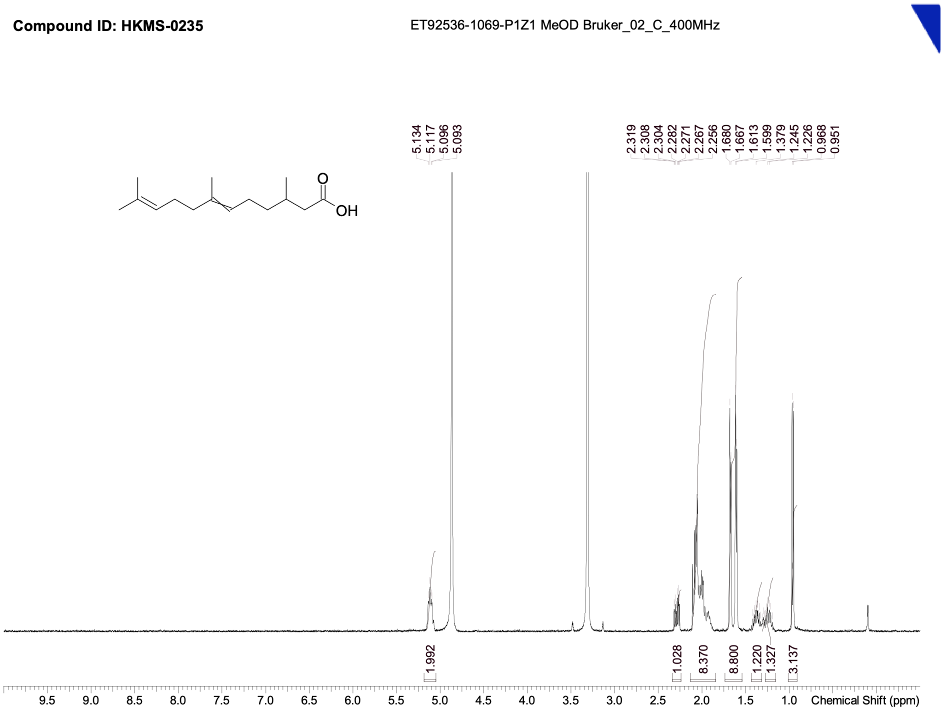


#### 13C NMR

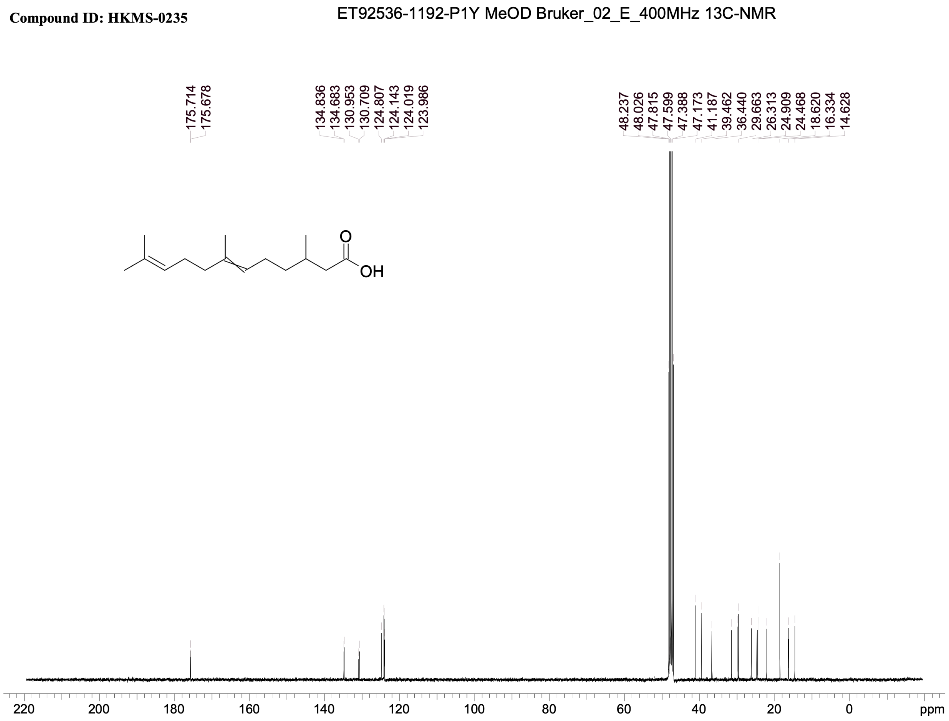

### HKMS-0238

#### 1H NMR

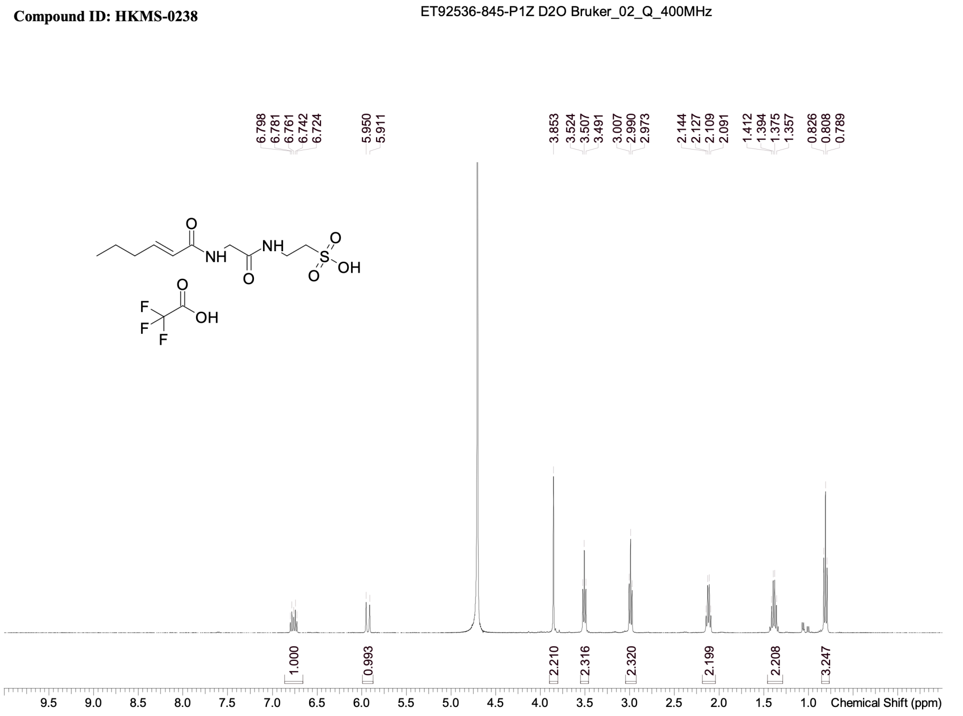


#### 13C NMR

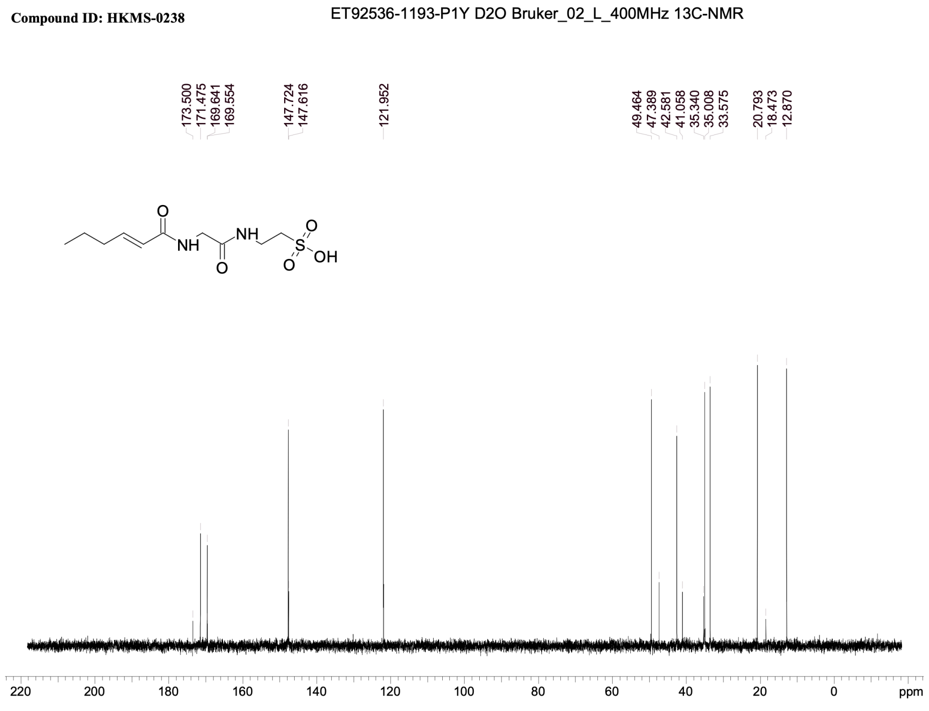

### HKMS-0241

#### 1H NMR

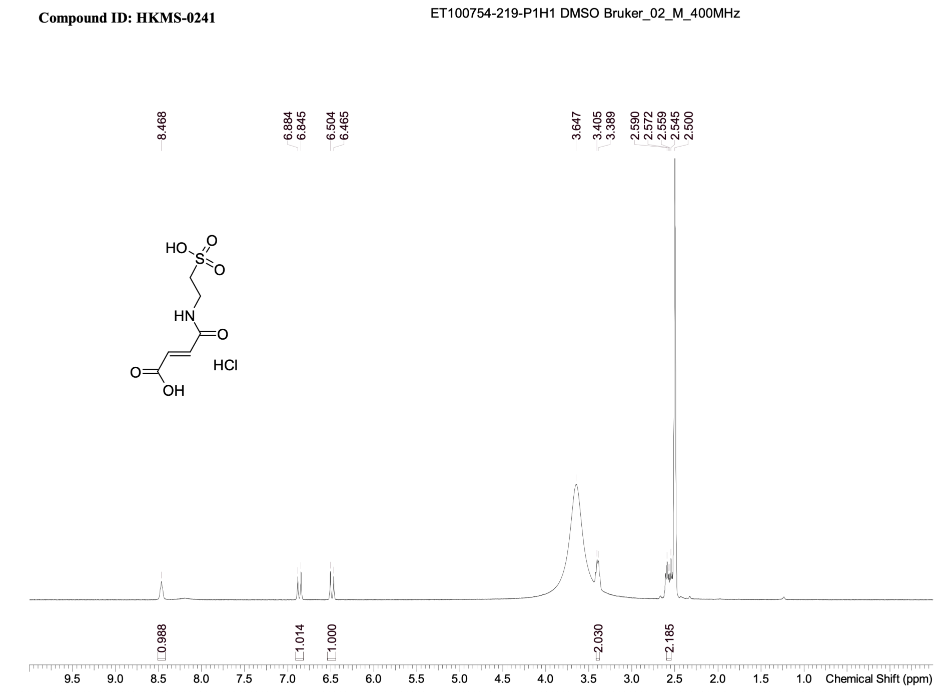


#### 13C NMR

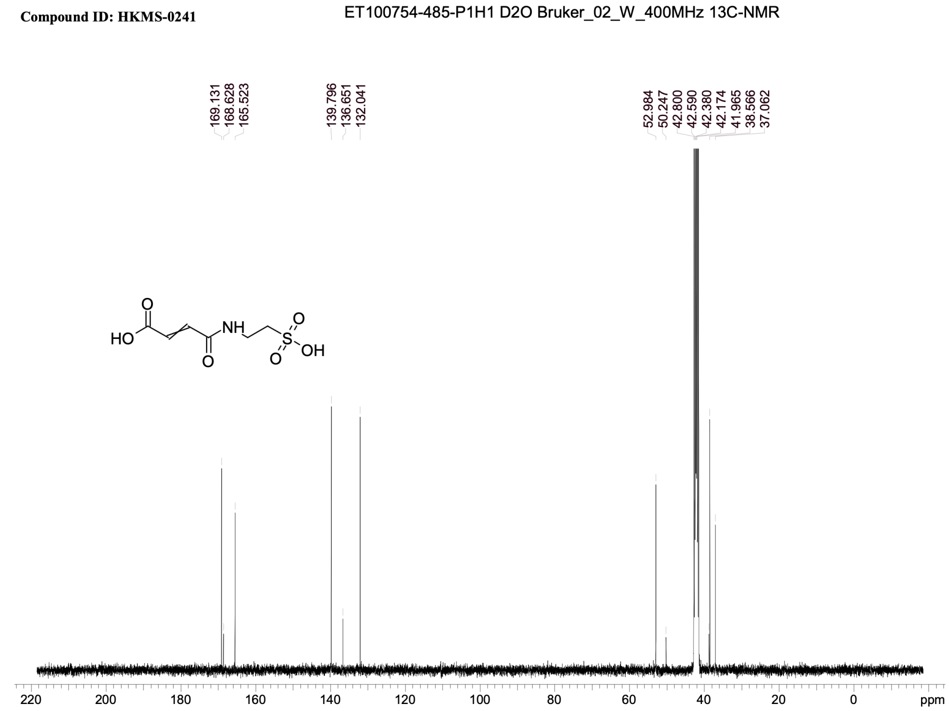

### HKMS-0245

#### 1H NMR

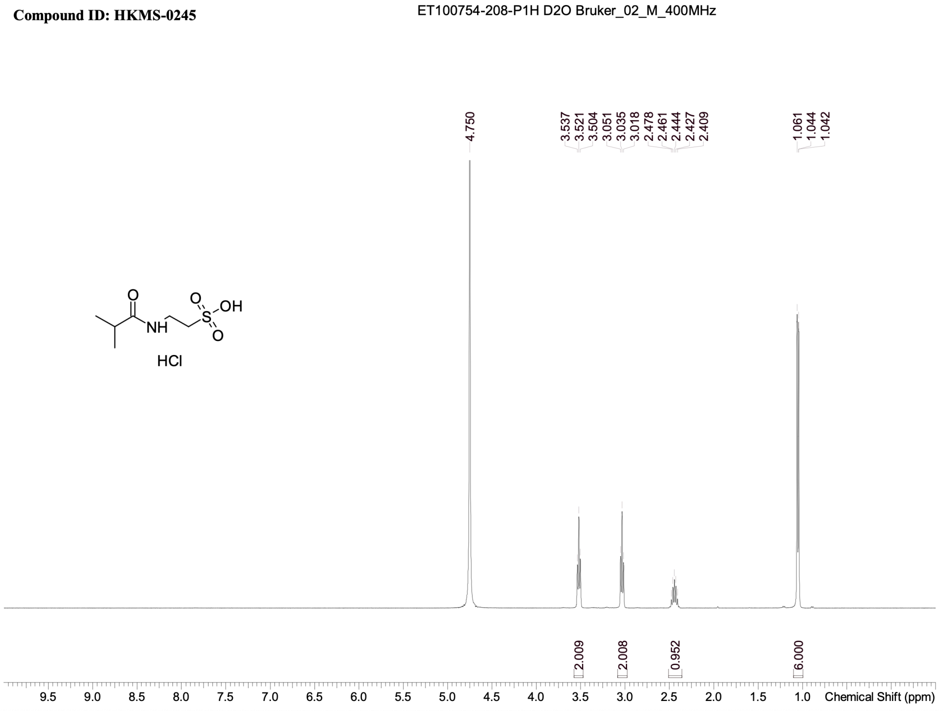


#### 13C NMR

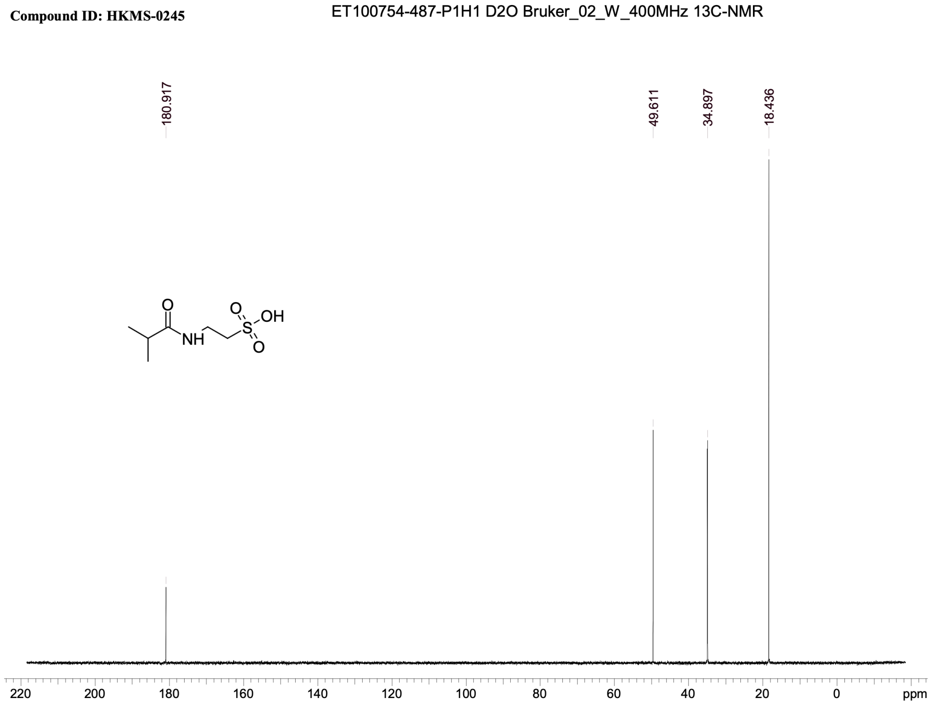

### HKMS-0257

#### 1H NMR

**Compound ID: HKMS-0257**

ET92536-913-P1Z2 D2O Bruker_02_M_400MHz

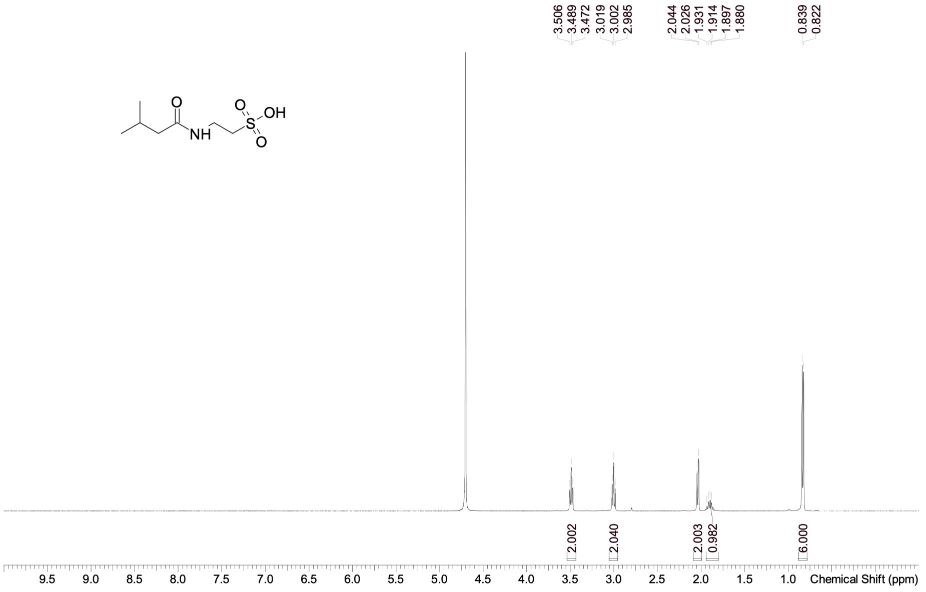


#### 13C NMR

**Compound ID: HKMS-0257**

ET92536-1194-p1y D2O Bruker_02_L_400MHz 13C-NMR

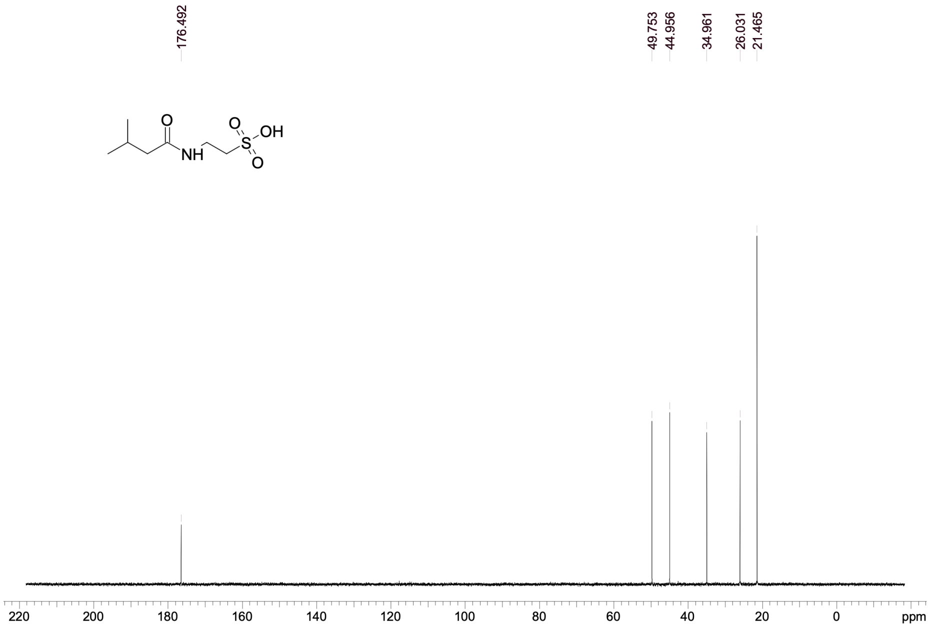

### HKMS-0260

#### 1H NMR

**Compound ID: HKMS-0260**

ET100754-277-P1H D2O Bruker_02_C_400MHz

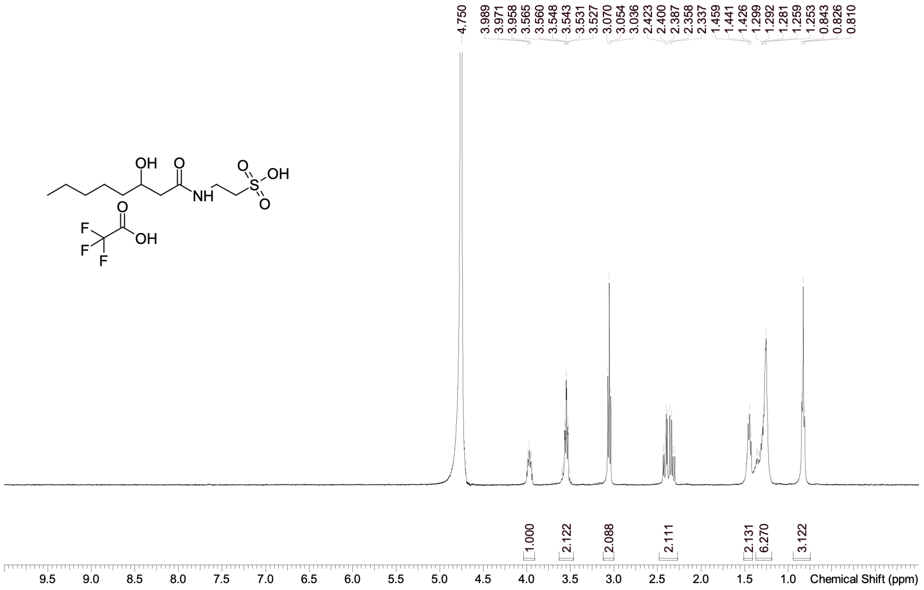


#### 13C NMR

**Compound ID: HKMS-0260**

ET100754-488-P1H1 D2O Bruker_02_F_400MHz 13C-NMR

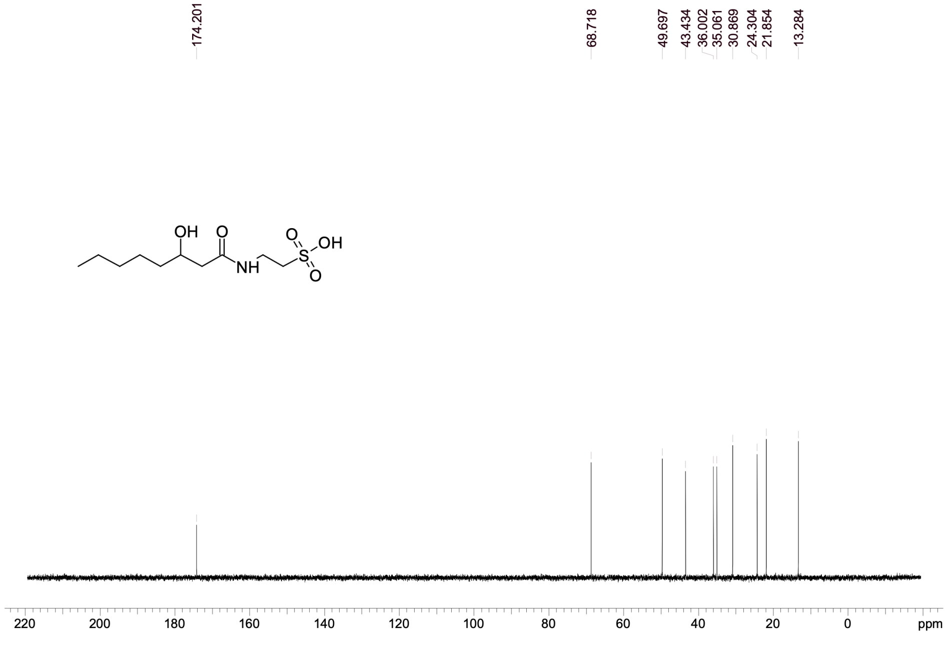

### HKMS-0263

#### 1H NMR ( + residual DIPEA)

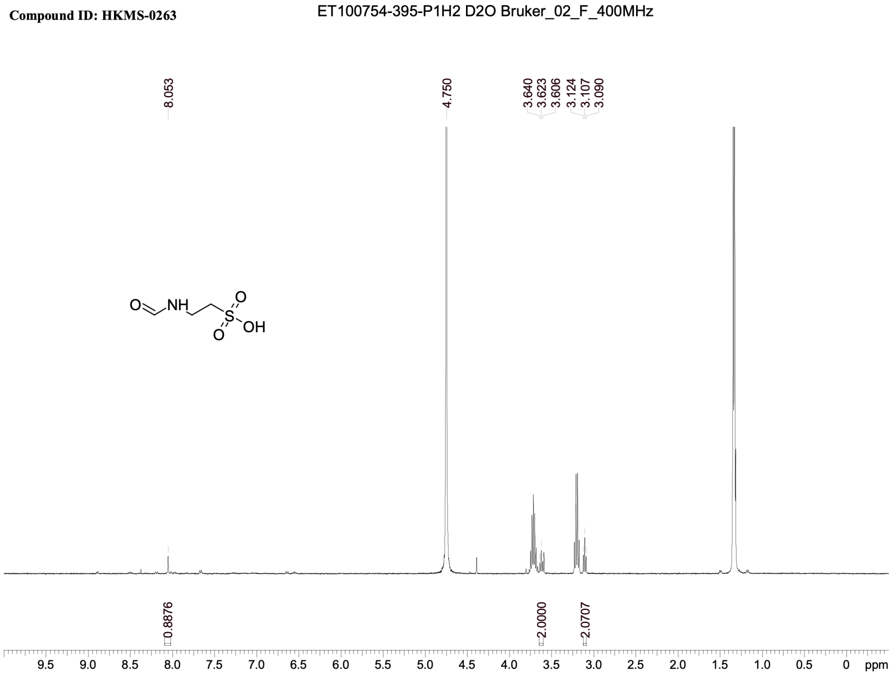


#### 13C NMR ( + residual DIPEA)

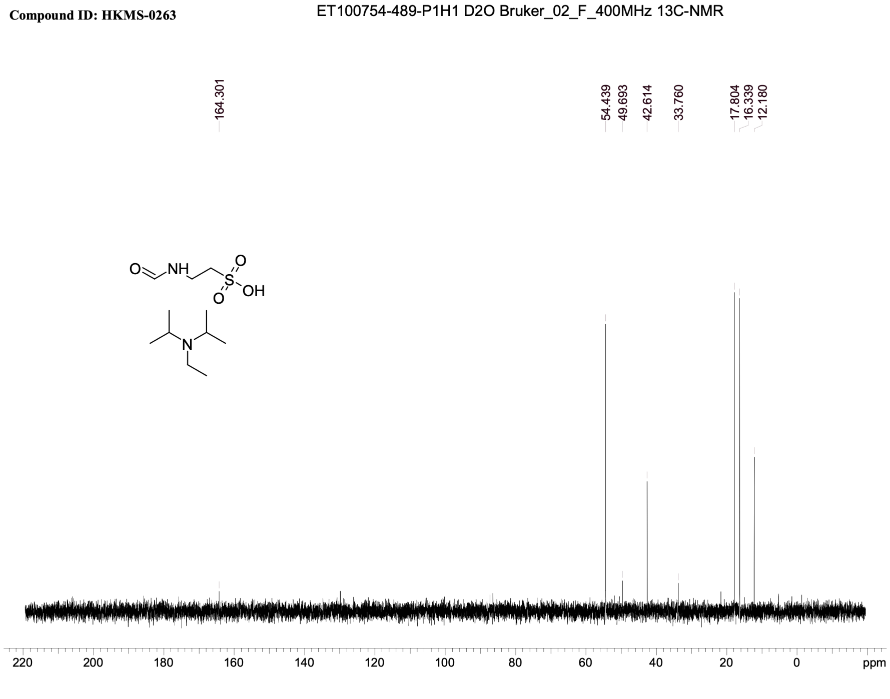

### HKMS-0273

#### 1H NMR

**Compound ID:HKMS-0273**

ET100716-291-P1B D2O Bruker_02_L_400MHz

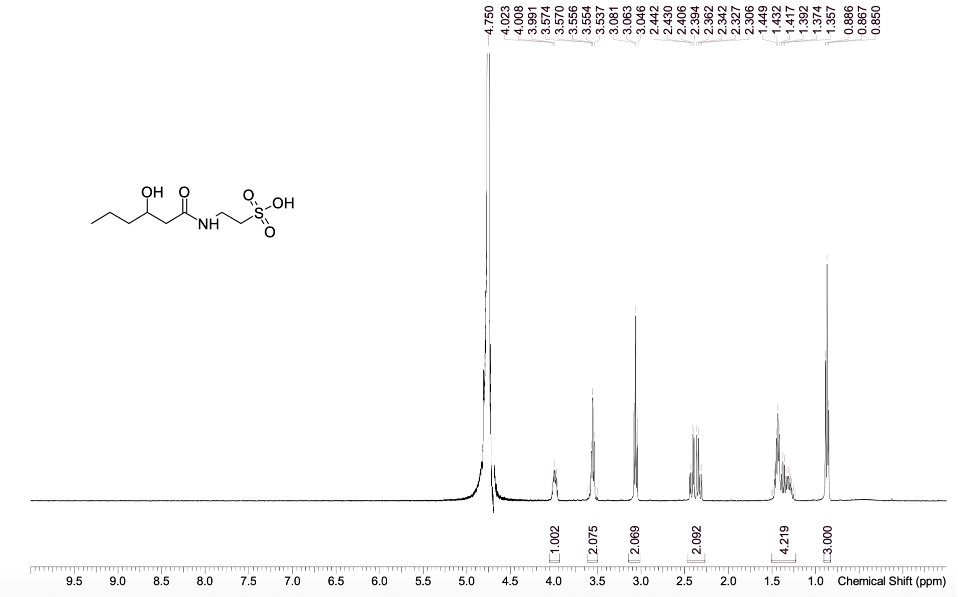


#### 13C NMR

**Compound ID:HKMS-0273**

ET100716-523-P1B D2O Bruker_02_F_400MHz 13C-NMR

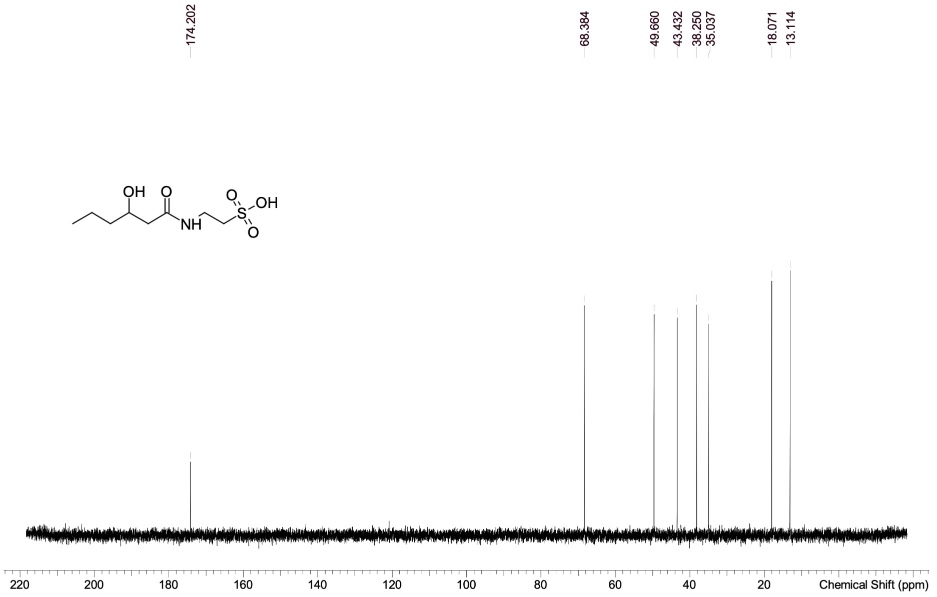

### HKMS-0281

#### 1H NMR

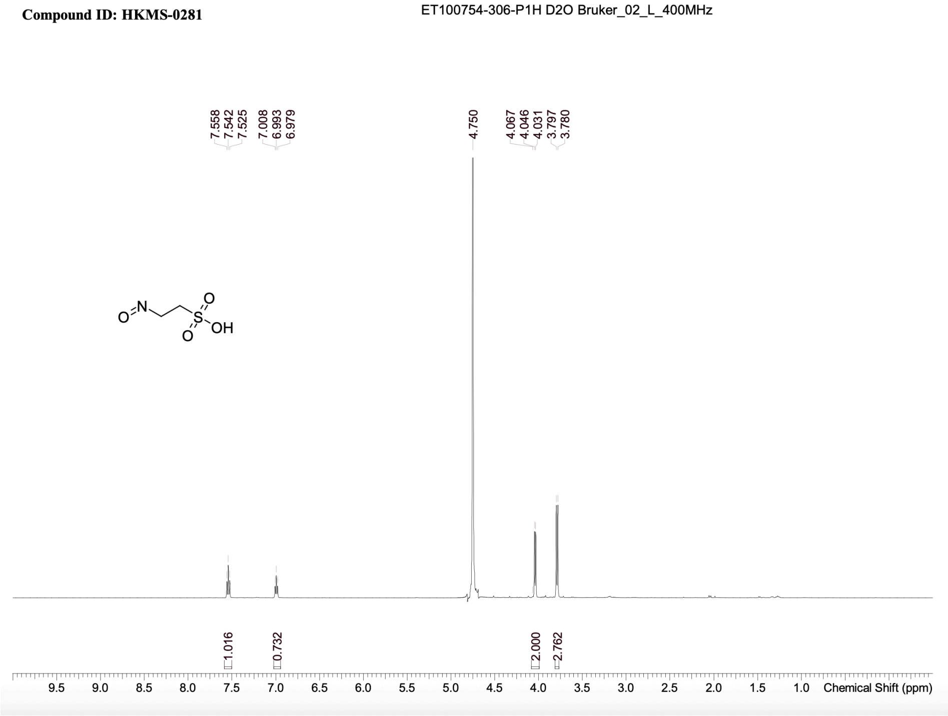


#### 13C NMR

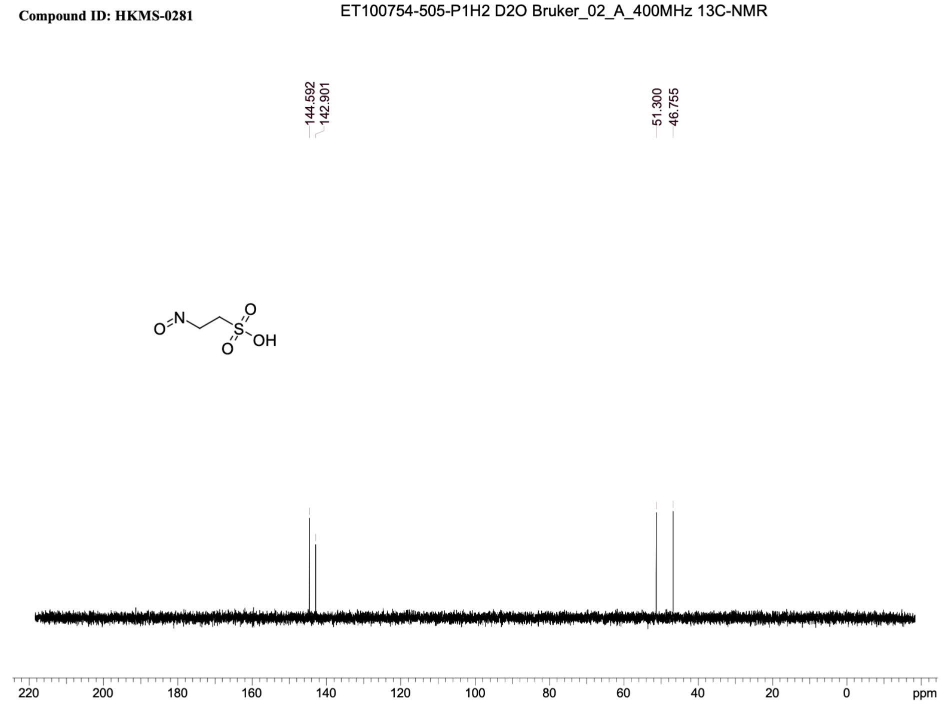

### HKMS-0242_200

#### 1H NMR

**Compound ID: HKMS-242-200**

ET100754-262-P1H1 D2O Bruker_02_M_400MHz

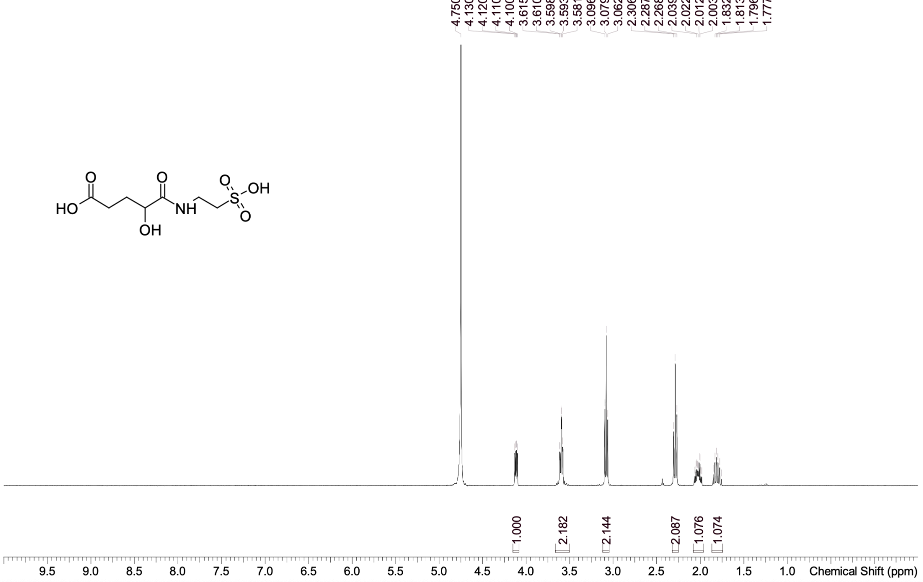


#### 13C NMR

**Compound ID: HKMS-0242-200**

ET100754-493-P1H2 D2O Bruker_02_F_400MHz 13C-NMR

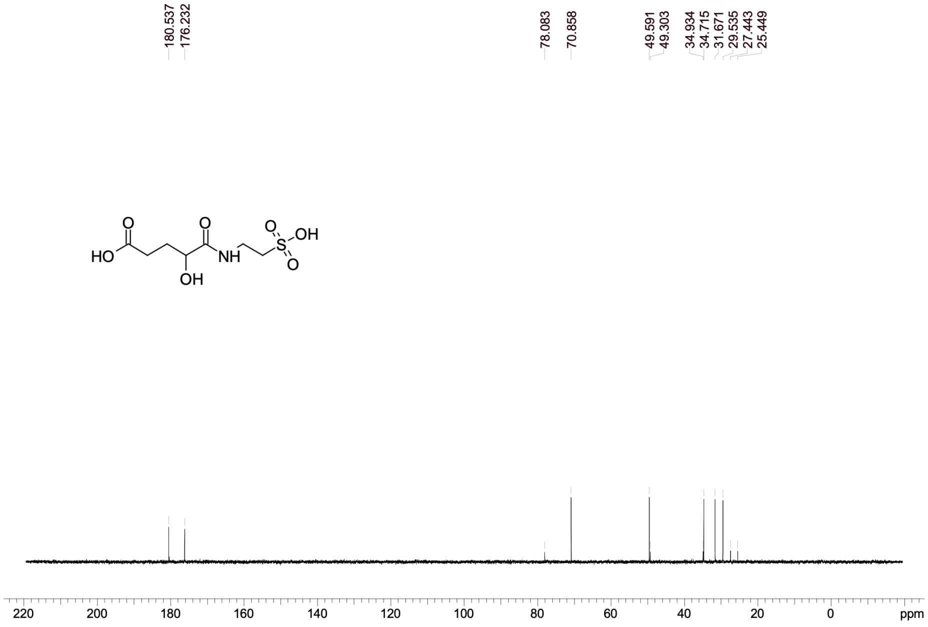

### HKMS-0277

#### 1H NMR

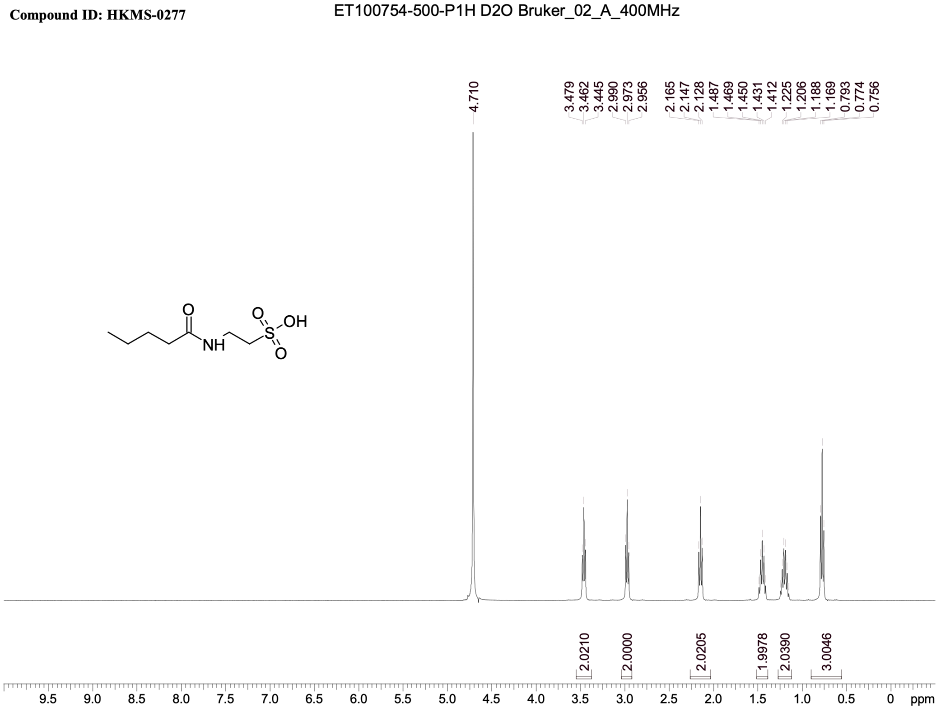


#### 13C NMR

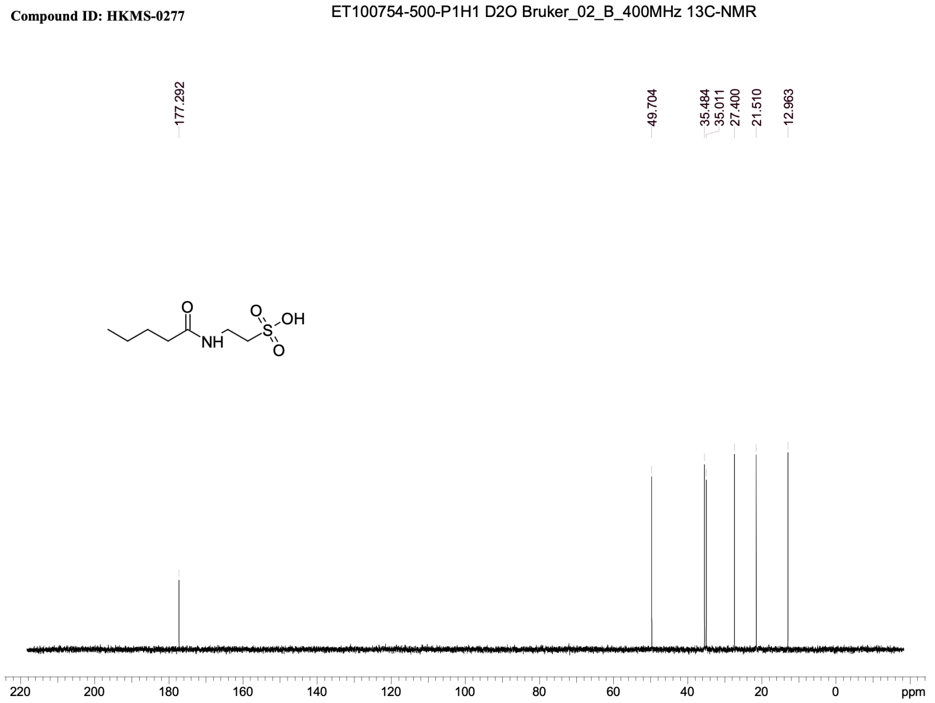

### HKMS-0284

#### 1H NMR

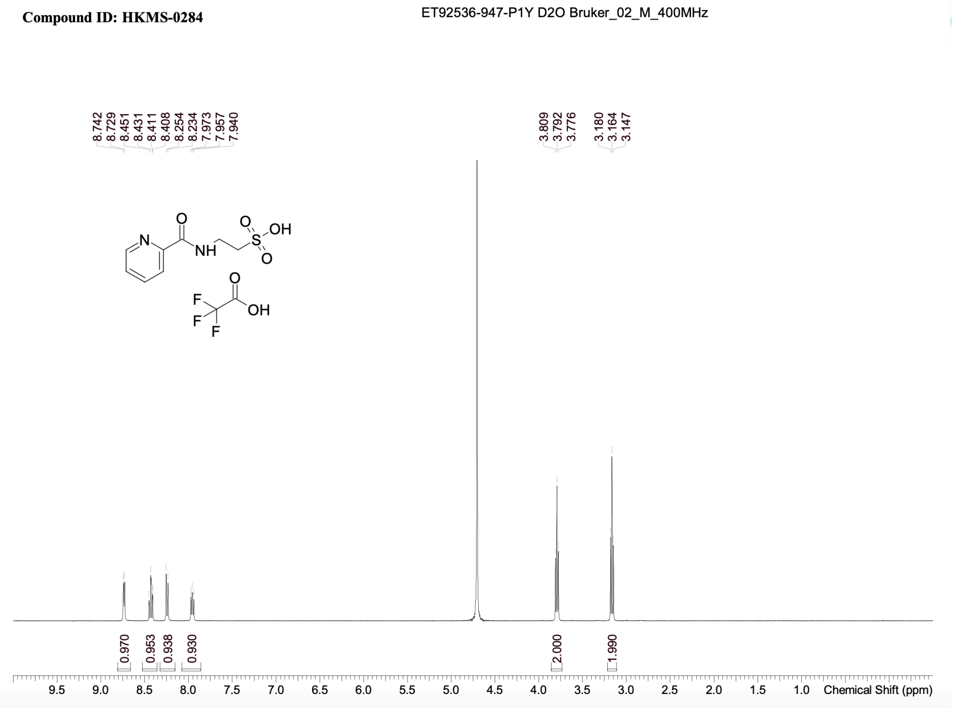


#### 13C NMR

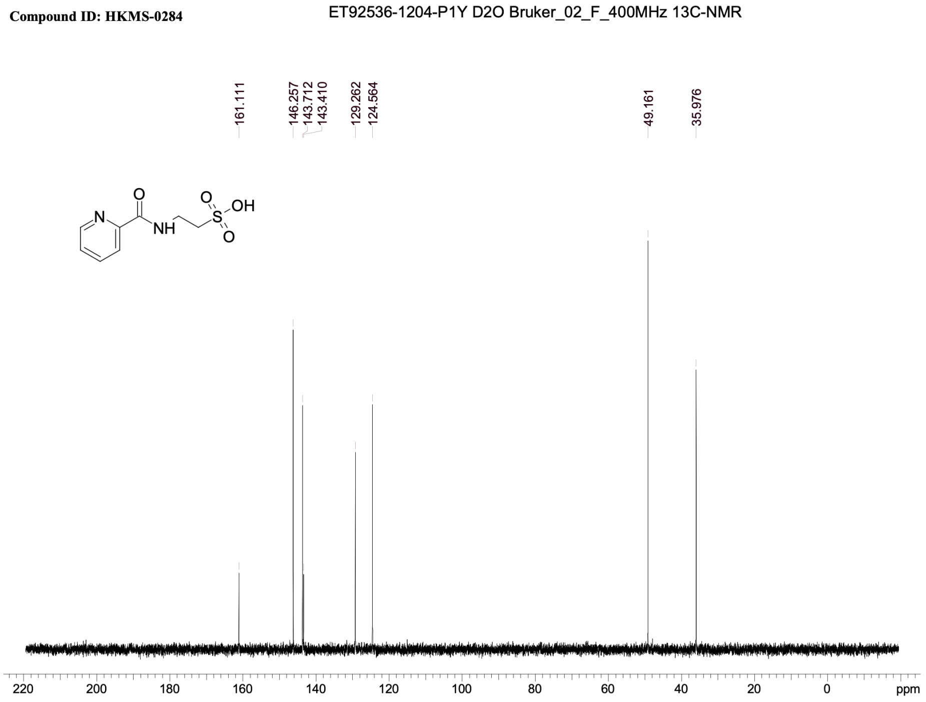

### HMS-0275

#### 1H NMR

**Compound ID: HKMS-0275**

ET92536-938-P1Z1 D2O Bruker_02_M_400MHz

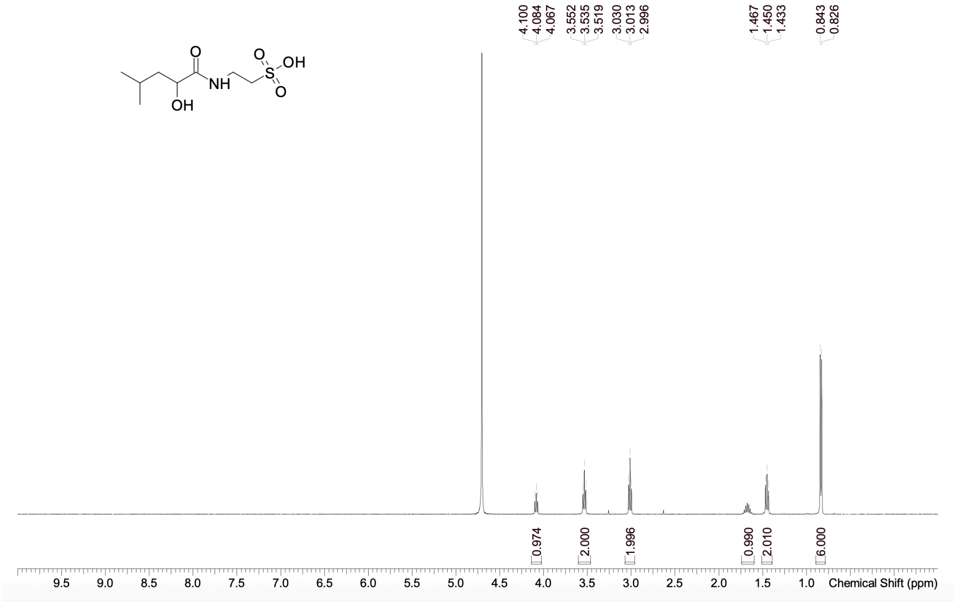


#### 13C NMR

**Compound ID: HKMS-0275**

ET92536-1203-P1Y D2O Bruker_02_C_400MHz 13C-NMR

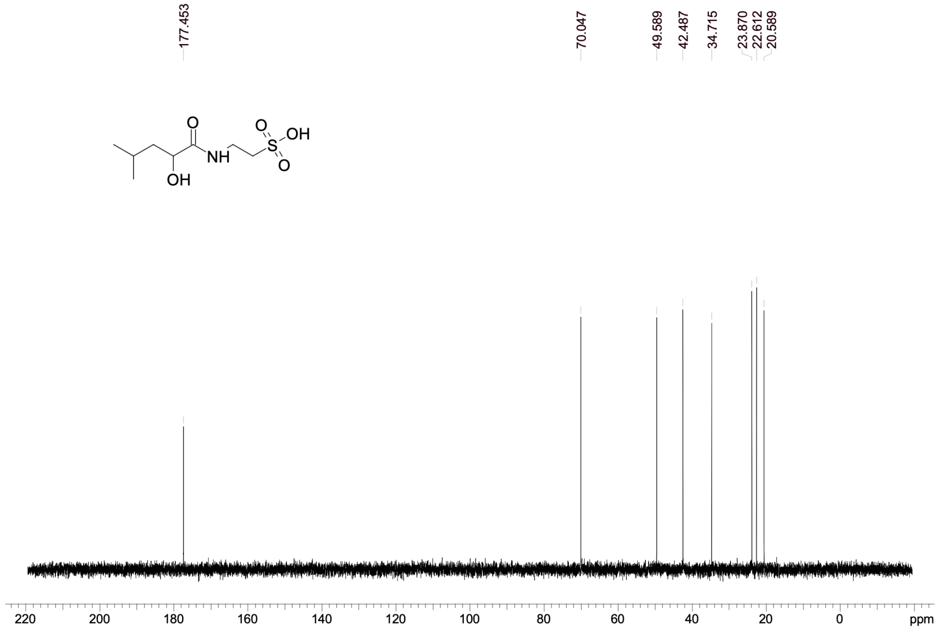

### HKMS-0282

#### 1H NMR

**Compound ID: HKMS-0282**

ET100754-280-P1H D2O Bruker_02_C_400MHz

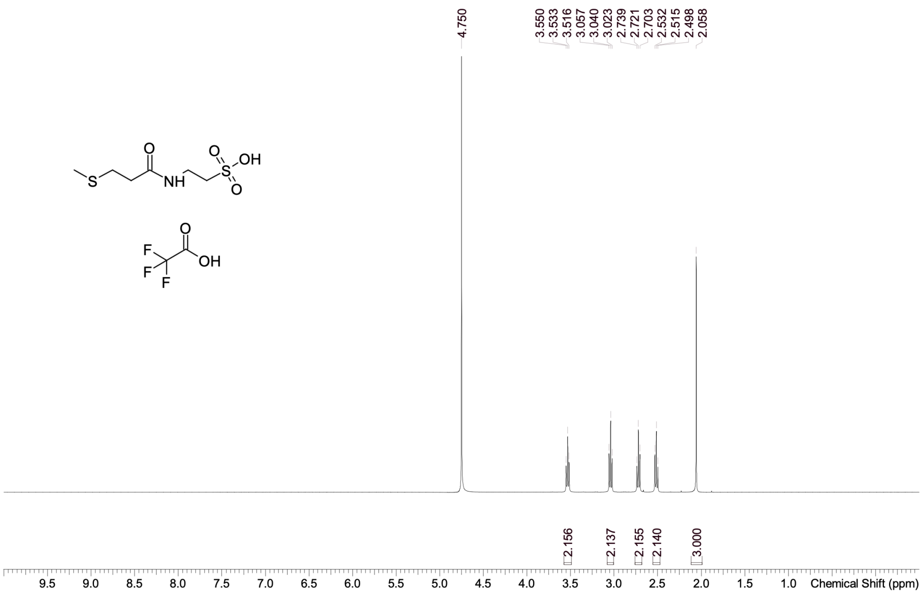


#### 13C NMR

**Compound ID: HKMS-0282**

ET100754-491-P1H1 D2O Bruker_02_F_400MHz 13C-NMR

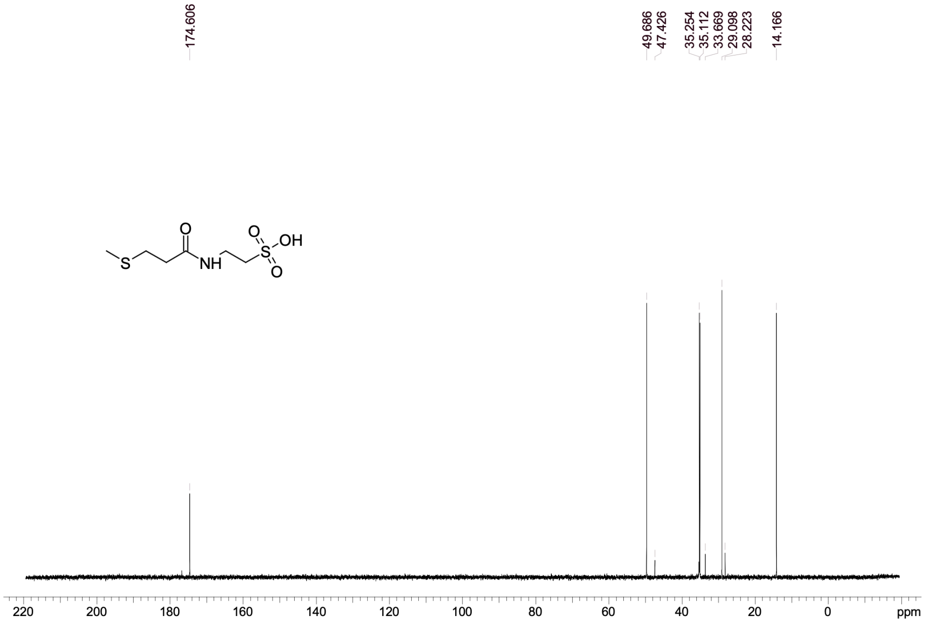

### HKMS-0283

#### 1H NMR

**Compound ID: HKMS-0283**

ET100754-282-P1H D2O Bruker_02_C_400MHz

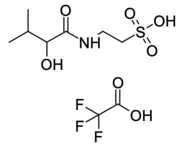


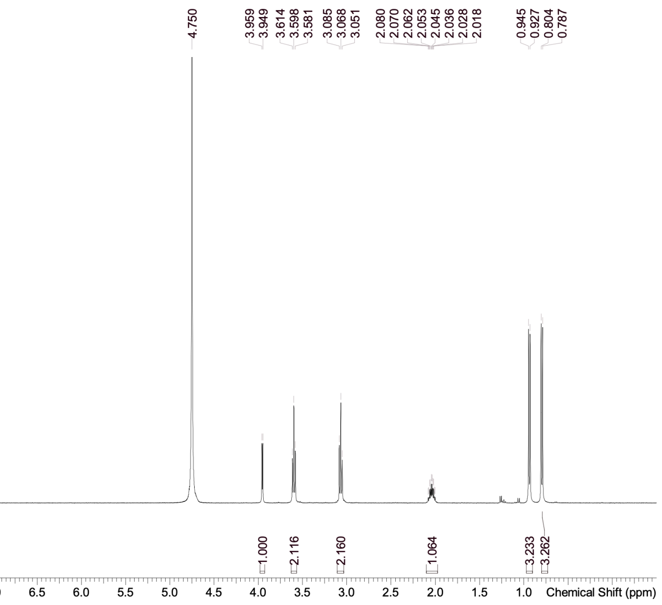


#### 13C NMR

**Compound ID: HKMS-0283**

ET100754-492-P1H1 D2O Bruker_02_A_400MHz 13C-NMR

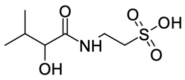


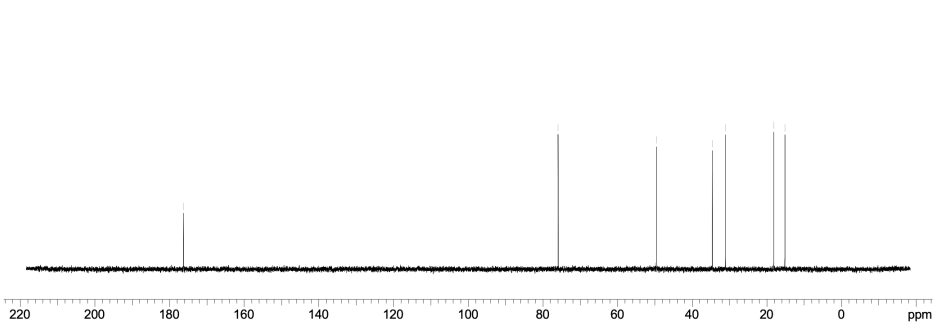

### HKMS-0285

#### 1H NMR

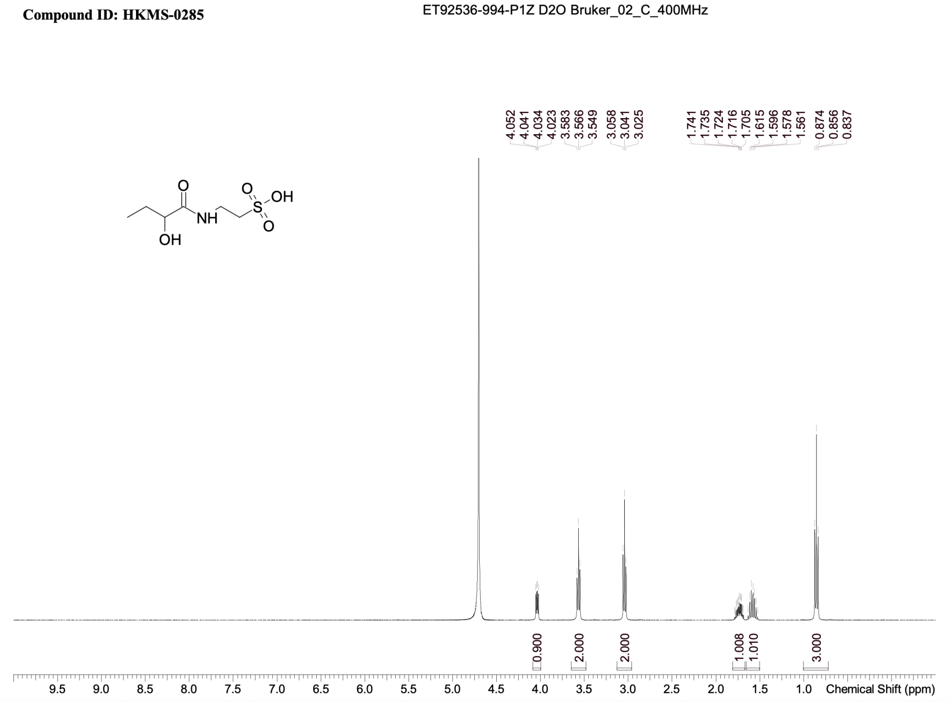


#### 13C NMR

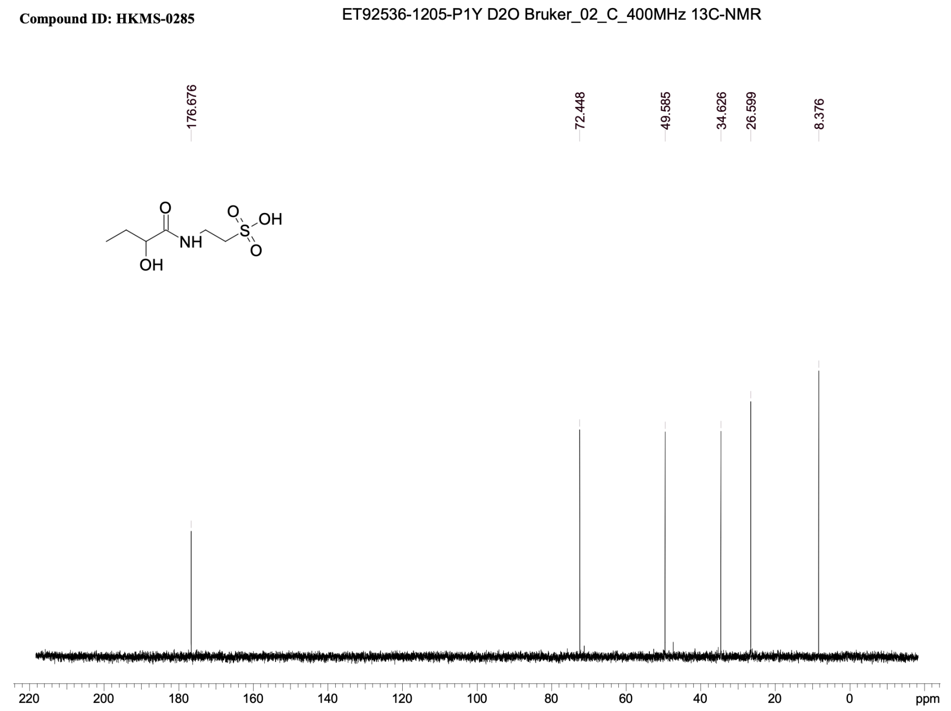

### HKMS-0286

#### 1H NMR

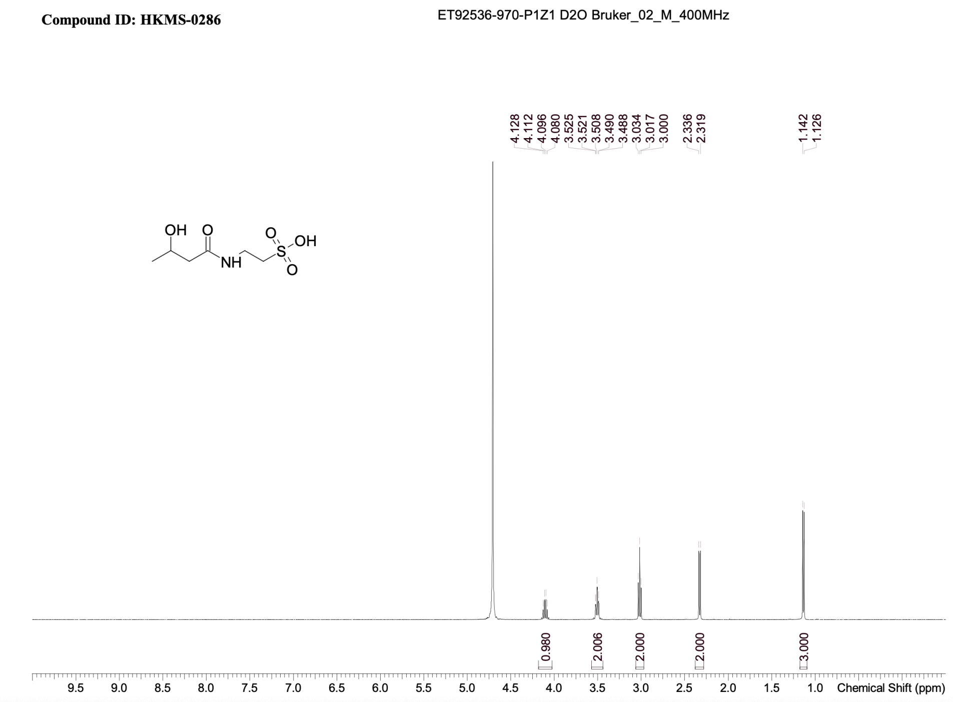


#### 13C NMR

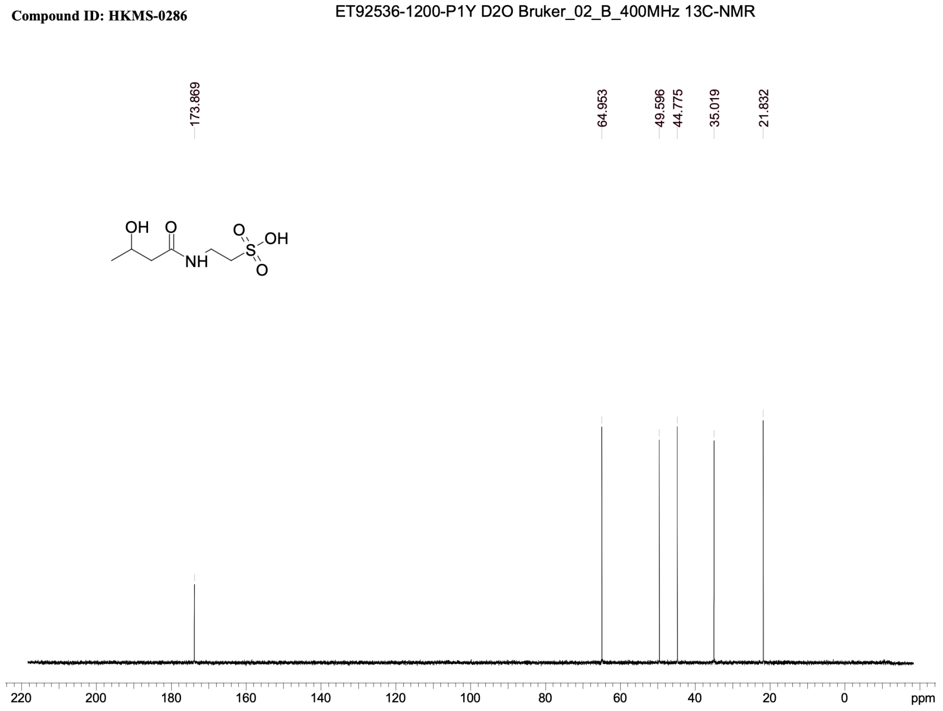

### HKMS-0213

#### 1H NMR

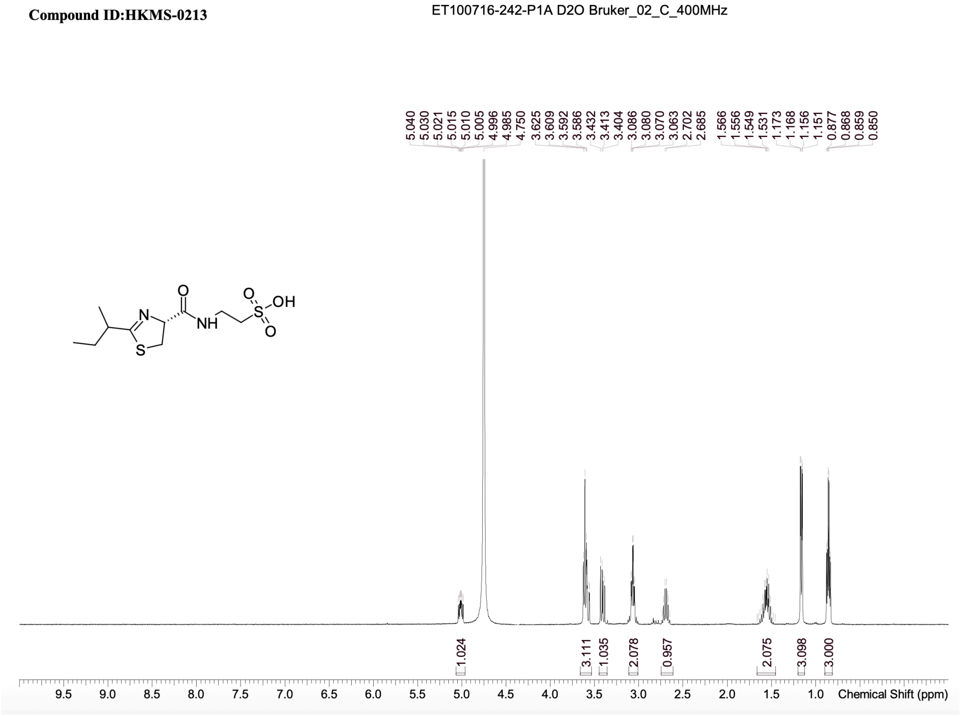


#### 13C NMR

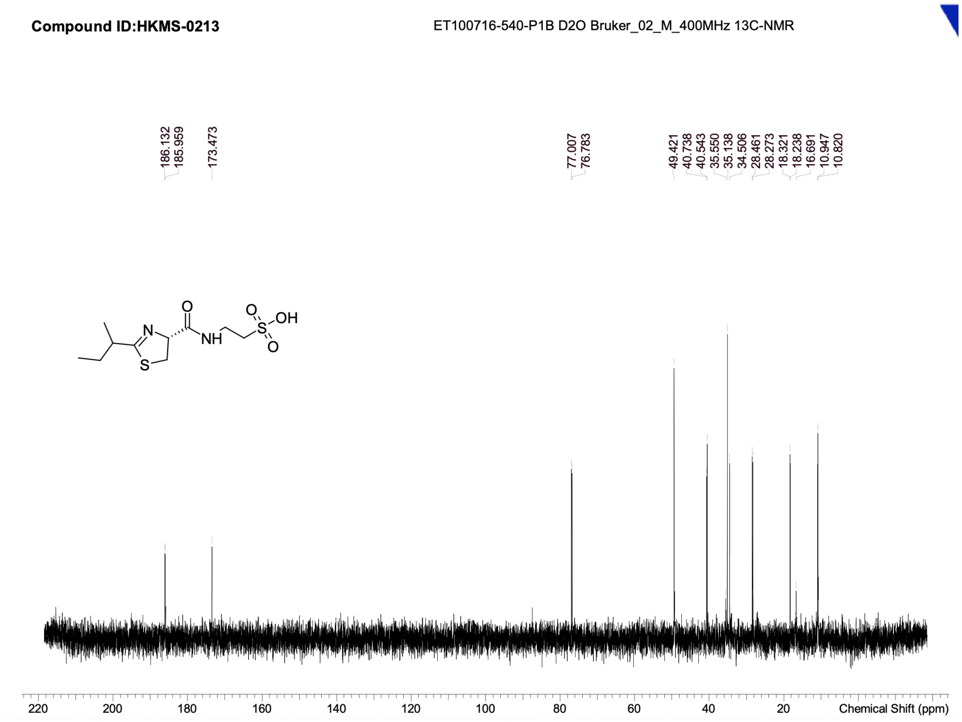
