## Supplementary Note 2 for "Mapping the mammalian dark metabolome by *in vivo* isotope tracing"

**Supplementary Note 2 | Overview of the existing literature for previously unrecognized metabolites.**

Throughout the main text, we use the terminology of "previously unrecognized metabolite" to refer to small molecules that, to the best of our knowledge, had not previously been recognized as mammalian metabolites. We employed a multi-tiered process involving extensive manual review to support this categorization, as described in the Methods section ("Terminology"). Here, we provide additional context for metabolites that have previously appeared in the literature, in particular when these have been identified in non-mammalian species, *in vitro*, or tentatively identified (but not definitively confirmed).

**N,N-dimethylargininic acid.** We were unable to locate any report of the identification of this compound as a metabolite in any species.

**S-methylthiomethyl mercapturic acid.** We were unable to locate any report of the identification of this compound as a metabolite in any species.

**(3-cyanopropanoyl)(sulfo)alanylglycine.** This compound is not found in PubChem. We were unable to locate any report of its identification as a metabolite in any species.

**2-propylthiazolidine-4-carboxylic acid.** An isomer of this compound, 2-isopropylthiazolidine-4-carboxylic acid and 2-isobutylthiazolidine-4-carboxylic acid, has been identified in beer[1–3]. This compound has been described as a component of the human exposome using a literature mining approach[4], possibly because it is sometimes referred to by the acronym PCTA, which also refers to a medical procedure, percutaneous transluminal coronary angioplasty, and PubChem lists several literature references to the compound from cardiological journals. It was previously associated with a HMDB identifier, HMDB0245245, that has been retired, and, perhaps as a result, has been reported as a metabolite in peritoneal fluid, pomegranates, and mung beans, apparently through tentative identifications based on m/z matching only[5–7]. In view of the apparently spurious nature of its identification as a component of the human exposome, the fact that no analytical data was made available to corroborate recent purported identifications, and the growing scrutiny of unreliable annotations in metabolomic studies[8], we considered it appropriate to categorize as a previously unrecognized metabolite.

**2-butylthiazolidine-4-carboxylic acid.** Isomers of this compound, 2-sec-butylthiazolidine-4-carboxylic acid and 2-isobutylthiazolidine-4-carboxylic acid, have been identified in beer[1–3]. We were unable to locate any report of its identification in mammals.

**2-pentylthiazolidine-4-carboxylic acid.** This compound has been identified in certain foods, including orange juice[9] and beer[1,2]. We were unable to locate any report of its identification in mammals.

**2-hexylthiazolidine-4-carboxylic acid.** We were unable to locate any report of the identification of this compound as a metabolite in any species.

**2-heptylthiazolidine-4-carboxylic acid.** This compound has been identified in orange juice[9]. We were unable to locate any report of its identification in mammals.

**2-octylthiazolidine-4-carboxylic acid.** This compound is not found in PubChem, but has been identified in orange juice[9]. We were unable to locate any report of its identification in mammals.

**2-nonylthiazolidine-4-carboxylic acid.** This compound has been identified in orange juice[9]. We were unable to locate any report of its identification in mammals.

**2-(sec-butyl)-4,5-dihydrothiazole-4-carboxylic acid.** We were unable to locate any report of the identification of this compound as a metabolite in any species.

**(2-pentylthiazolidine-4-carbonyl)glycine.** This compound is not found in PubChem. We were unable to locate any report of its identification as a metabolite in any species.

**(2-heptylthiazolidine-4-carbonyl)glycine.** This compound is not found in PubChem. We were unable to locate any report of its identification as a metabolite in any species.

**(2-octylthiazolidine-4-carbonyl)glycine.** This compound is not found in PubChem. We were unable to locate any report of its identification as a metabolite in any species.

**2,3-dihydrofarnesoic acid.** This compound has been reported as a terpene produced by hairy tomatoes, *Lycopersicon hirsutum*[10], and as a pheromone produced by multiple insect species including European beewolves, *Philanthus triangulum*[11] and *Heliconius* butterflies[12]. We were unable to locate any report of its identification in mammals.

**Taurine-C4:0.** We were unable to locate any report of the identification of this compound as a metabolite in any species.

**Taurine-C5:0.** While this work was underway, taurine-C5:0 was tentatively identified based on a MS/MS search of the GNPS repository through comparison to the MS/MS acquired from a synthetic standard; however, the identification was not confirmed by retention time with biological samples[13]. We were unable to locate any other report of the identification of this compound as a metabolite in any species.

**Taurine-C6:0;O1($\alpha$).** Labelling and analytical data was consistent with three (leucine-, isoleucine-, or linoleic acid-derived) isomers; we were unable to locate any report of the identification of any of these compounds as a metabolite in any species.

**Taurine-C6:0;O1($\beta$).** This compound is not found in PubChem. We were unable to locate any report of its identification as a metabolite in any species.

**Taurine-C6:1;O1($\alpha$).** Neither branched (leucine- or isoleucine-derived) or linear isomers of this compound could be found in PubChem, and we were unable to locate any report of the identification of any isomer as a metabolite in any species.

**Taurine-C8:0.** While this work was underway, taurine-C8:0 was tentatively identified based on a MS/MS search of the GNPS repository through comparison to the MS/MS acquired from a synthetic standard; however, the identification was not confirmed by retention time with biological samples[13]. We were unable to locate any other report of the identification of this compound as a metabolite in any species.

**Taurine-C8:0;O1.** This compound is not found in PubChem. We were unable to locate any report of its identification as a metabolite in any species.

**Dodecanedioyltaurine.** This compound is not found in PubChem. We were unable to locate any report of its identification as a metabolite in any species.

**$\alpha$-hydroxybutyroyltaurine.** This compound is not found in PubChem. We were unable to locate any report of its identification as a metabolite in any species.

**$\beta$-hydroxybutyroyltaurine.** This compound is not found in PubChem. We were unable to locate any report of its identification as a metabolite in any species.

**$\alpha$-hydroxyisovaleroyltaurine.** Neither this compound, nor the isomer $\alpha$-hydroxyvaleroyltaurine, is found on PubChem. We were unable to locate any report of the identification of either compound as a metabolite in any species.

**Acetoacetoyltaurine.** We were unable to locate any report of the identification of this compound as a metabolite in any species.

**Lactoyltaurine.** This compound has previously been described as a component of oyster glycolipids[14]. We were unable to locate any report of its identification in mammals.

**Fumaryltaurine.** We were unable to locate any report of the identification of this compound as a metabolite in any species.

**2-hydroxyglutaroyltaurine.** This compound is not found in PubChem. We were unable to locate any report of its identification as a metabolite in any species.

**Taurine succinimide.** We were unable to locate any report of the identification of this compound as a metabolite in any species.

**2-(2-oxopropylamino)ethanesulfonic acid.** We were unable to locate any report of the identification of this compound as a metabolite in any species.

**Glycyltaurine.** The fate of synthetic glycyltaurine has been studied after its administration to rabbits[15], and glycyltaurine conjugates of deoxycholic acid[16] and quinaldic acid[17] have been described in mammals; however, we were unable to locate any report of glycyltaurine itself as a metabolite in any species.

**Alanyltaurine.** Alanyltaurine was considered, but discredited, as a match to an unknown signal detected in rat heart by $^{1}$H NMR[18]. We were unable to locate any report of its identification as a metabolite in any species.

**Prolyltaurine.** We were unable to locate any report of the identification of this compound as a metabolite in any species.

**Threonyltaurine.** We were unable to locate any report of the identification of this compound as a metabolite in any species.

**Glutaminyltaurine.** We were unable to locate any report of the identification of this compound as a metabolite in any species.

**N-acetylseryltaurine.** This compound is not found in PubChem. We were unable to locate any report of the identification of this compound as a metabolite in any species.

**N-acetylphenylalanyltaurine.** This compound is not found in PubChem. We were unable to locate any report of the identification of this compound as a metabolite in any species.

**Glycyltaurine-C5:0.** This compound is not found in PubChem. We were unable to locate any report of the identification of this compound as a metabolite in any species.

**Glycyltaurine-C6:0.** This compound is not found in PubChem. We were unable to locate any report of the identification of this compound as a metabolite in any species.

**Glycyltaurine-C6:1.** This compound is not found in PubChem. We were unable to locate any report of the identification of this compound as a metabolite in any species.

**Glycyltaurine-C8:0.** This compound is not found in PubChem. We were unable to locate any report of the identification of this compound as a metabolite in any species.

**Glycyltaurine-C10:1.** This compound is not found in PubChem. We were unable to locate any report of the identification of this compound as a metabolite in any species.

**Acetylglycyltaurine.** This compound is not found in PubChem. We were unable to locate any report of its identification as a metabolite in any species.

**Phenylacetylglycyltaurine.** This compound is not found in PubChem. We were unable to locate any report of its identification as a metabolite in any species.

**Picolinoyltaurine.** This compound is not found in PubChem. We were unable to locate any report of the identification of this compound as a metabolite in any species.

**3-methylthiopropionyltaurine.** We were unable to locate any report of the identification of this compound as a metabolite in any species.

**N-formyltaurine.** We were unable to locate any report of the identification of this compound as a metabolite in any species.

**(2-(sec-butyl)-4,5-dihydrothiazole-4-carbonyl)taurine.** This compound is not found in PubChem. We were unable to locate any report of the identification of this compound as a metabolite in any species.

**S-methylthiomethyl mercapturoyltaurine.** This compound is not found in PubChem. We were unable to locate any report of the identification of this compound as a metabolite in any species.

**Phloretoyltaurine.** This compound is not found in PubChem. We were unable to locate any report of the identification of this compound as a metabolite in any species.

**Nitrotaurine.** We were unable to locate any report of the identification of this compound as a metabolite in any species.

**Nitrosotaurine.** We were unable to locate any report of the identification of this compound as a metabolite in any species.
